# An mRNA Display Approach for Covalent Targeting of a *Staphylococcus aureus* Virulence Factor

**DOI:** 10.1101/2024.11.06.622387

**Authors:** Sijie Wang, Emily C. Woods, Jeyun Jo, Jiyun Zhu, Althea Hansel-Harris, Matthew Holcomb, Nichole J. Pedowitz, Tulsi Upadhyay, John Bennett, Matthias Fellner, Ki Wan Park, Anna Zhang, Tulio A. Valdez, Stefano Forli, Alix I Chan, Christian N. Cunningham, Matthew Bogyo

## Abstract

Staphylococcus aureus (*S. aureus*) is an opportunistic human pathogen that causes over one million deaths around the world each year. We recently identified a family of serine hydrolases termed fluorophosphonate binding hydrolases (Fphs) that play important roles in lipid metabolism and colonization of a host. Because many of these enzymes are only expressed in *Staphylococcus* bacteria, they are valuable targets for diagnostics and therapeutics. Here we developed and screened highly diverse cyclic peptide libraries using mRNA display with a genetically encoded oxadiazolone (Ox) electrophile that was previously shown to potently and covalently inhibit multiple Fph enzymes. By performing multiple rounds of counter selections with WT and catalytic dead FphB, we were able to tune the selectivity of the resulting selected cyclic peptides containing the Ox residue towards the desired target. From our mRNA display hits, we developed potent and selective fluorescent probes that label the active site of FphB at single digit nanomolar concentrations in live *S. aureus* bacteria. Taken together, this work demonstrates the potential of using direct genetically encoded electrophiles for mRNA display of covalent binding ligands and identifies potent new probes for FphB that have the potential to be used for diagnostic and therapeutic applications.

## INTRODUCTION

Covalent inhibitors have become indispensable in probe development and drug discovery due to their unique mechanism of action.^1,2^ Unlike traditional non-covalent inhibitors, covalent inhibitors function through a two-step process that includes an initial reversible binding to their target protein followed by the formation of a covalent bond between an electrophilic moiety, known as the “warhead,” and a specific amino acid residue on the protein.^3,4^ This formation of a durable covalent bond not only enhances overall drug efficacy by providing a prolonged effect, but can also drive high specificity while potentially reducing the likelihood of drug resistance.^2,4–6^

The rational design of covalent inhibitors often relies heavily on structural information of the target protein to position a warhead on an existing high affinity ligand and yield a potent and irreversible binding molecule.^2,7,8^ However, in cases where structural information is lacking, alternative approaches are necessary for *de novo* development of covalent inhibitors. High-throughput screening strategies, including the use of fragment-based electrophilic libraries^9–12^ and genetically encoded libraries (GELs)^13–19^ with high diversity, are effective methods for this purpose. Structurally diverse electrophilic libraries have proven to be highly valuable for discovering,^20^ profiling,^21,22^ and screening of disease-associated biomarkers.^9–11,23^ GELs, such as those based on phage display,^24^ mRNA display,^25^ or DNA-encoded libraries,^26^ offer even greater diversity (10^9^–10^15^) and are emerging as powerful tools for covalent drug discovery. Particularly, mRNA and phage display, which can incorporate warheads during or after translation of peptides or proteins,^18,19,27^ are exceptionally powerful tools for covalent ligand discovery. The dual-function roles of the genetic template, serving as both a barcode for hit deconvolution and template for library resynthesis in display methods, allow progressive rounds of covalent probe screening. In addition, for mRNA display methods, it is possible to use genetic code reprogramming with flexizymes, a method known as FIT (flexible in vitro translation),^28–30^ to expand the chemical space from natural amino acids to a broad range of non-natural amino acids. This overall method of mRNA display using non-natural amino acids is termed RaPID (random nonstandard peptide integrated discovery).^31,32^

The ability to introduce non-natural amino acids into ribosomally synthesized peptides enables the *in situ* generation of macrocyclic peptides using a thiol-reactive N-terminal capping group. Macrocycles are therapeutically attractive moieties due to their enhanced proteolytic stability,^33^ bioavailability,^34^ increased cell membrane permeability,^35^ and conformational restrictions that reduce the entropic cost of binding protein surfaces.^36^ Therefore, macrocycles are becoming a core scaffold for drug development efforts targeting an ever increasing scope of proteins (cell surface receptors,^37–39^ enzymes,^40–44^ and PPI^45–50^). Furthermore, recently approved macrocyclic drugs suggest that cyclic peptides will continue to find broad applications in antibacterial, antiviral, and oncology drug develoment.^33^ However, despite the advantages of display methods, there remain only limited examples of screening campaigns using covalent binding peptide libraries,^19,27^ and yet fewer examples where the covalent electrophile is directly encoded into the peptides.^51^

Here we describe the use of a genetically encoded non-canonical analog of phenylalanine containing a reactive oxadiazolone electrophile to screen for potent covalent cyclic probes for a *Staphylococcus aureus* serine hydrolase, FphB.^52^ *Staphylococcus aureus* is a highly versatile and virulent bacterium, causing a range of infections from mild skin conditions to severe diseases like infective endocarditis (IE) or life-threatening bacteremia.^53,54^ The emergence of antibiotic-resistant strains, such as methicillin-resistant *S. aureus* (MRSA) and vancomycin-resistant *S. aureus* (VRSA), has significantly increased morbidity and mortality rates and effective treatments for these resistant strains are urgently needed.^54^ We previously used activity-based protein profiling (ABPP) with a fluorophosphonate (FP) probe to identify a family of serine hydrolases, termed Fph (fluorophosphonate binding) proteins.^52^ For this study, we chose to focus on FphB, because it is confirmed to be a membrane protein that is important for effective bacterial colonization of specific tissues inside a host.^52,55^ In addition, while we have reported on small molecule inhibitors of this target and used these molecules to label FphB on live bacteria, these compounds lack stability and showed cross reactivity when used at high concentrations.^52,56^ Recently, our group^55^ and others^57^ reported that the oxadiazolone (Ox) electrophile potently and irreversibly labels multiple Fph proteins in live *S. aureus*, making it a potentially ideal functional group for use in imaging and therapeutic agents targeting the Fph proteins with high sequence homology (Figure S1). However, this electrophile showed reactivity towards other enzymes including the lipid biosynthesis enzyme FabH by targeting a catalytic cysteine residue. In addition, the use of the Ox electrophile attached to a long lipid tail resulted in a highly FphE-specific probe.^55^ Therefore, we reasoned that further efforts would be required to direct the reactivity of the Ox electrophile away from FphE and FabH and towards FphB.

To accomplish this goal, we leveraged a modified oxadiazolone derivative for genetic encoding into large libraries of cyclic peptides using the flexible *in vitro* translation mRNA display approach. Using multiple rounds of positive selection for covalent FphB binding and counter selections using catalytically dead FphB, we identified potent covalent inhibitors of FphB targeting serine at the active site. These molecules show highly selective cellular labeling of FphB in live *S. aureus* with effective target modification at concentrations as low as single digit nanomolar. This study showcases the power of employing a genetic encoding strategy to incorporate covalent warheads into mRNA display, which enabled tuning of the reactivity of the covalent electrophile towards a specific enzyme target, even when the starting electrophile is highly potent for related off targets. The success of covalent cyclic probe development to this bacterial enzyme target highlights the potential of this mRNA display approach. We believe this approach can be more generally applied to enable the discovery of covalent probes for the treatment, diagnosis, and profiling of diverse disease-associated biomarkers.

## RESULTS AND DISCUSSION

### Design, Synthesis and Selections of mRNA Display Libraries to FphB

#### Warhead selection and design for mRNA display libraries

In a previous study,^55^ we determined that the oxadiazolone-based molecule JJ-OX-004 is a potent and highly selective probe for targeting FphE in the cytosol of live *S. aureus* bacteria, making it a potentially ideal electrophile for further development. Furthermore, this probe demonstrated moderately potent binding to FphB (IC_50_ = 0.7 µM; Figure S2) suggesting that it could potentially be used and modified for a selection approach to target FphB with increased potency and selectivity. We therefore designed an oxadiazolone-based warhead fragment, JJ-OX-009 (Ox; Figure 1A), that could be easily incorporated into a modified amino acid containing a ketone through oxime formation for encoding into peptides using mRNA display methods. The parent electrophile, JJ-OX-009, showed significantly reduced potency (IC_50_ > 20 µM) for FphB compared to the original JJ-OX-004 containing the lipid backbone (Figure S2). This same fragment that lost two orders of magnitude potency for FphB showed a 10-fold drop in potency for the related FphE enzyme (30 nM to 300 nM; Figure S2). Therefore, our starting Ox electrophile has high potency and selectivity for the off target FphE. We therefore wanted to determine if mRNA selections with this electrophile would allow targeting of the electrophile towards FphB.

**Figure 1.**
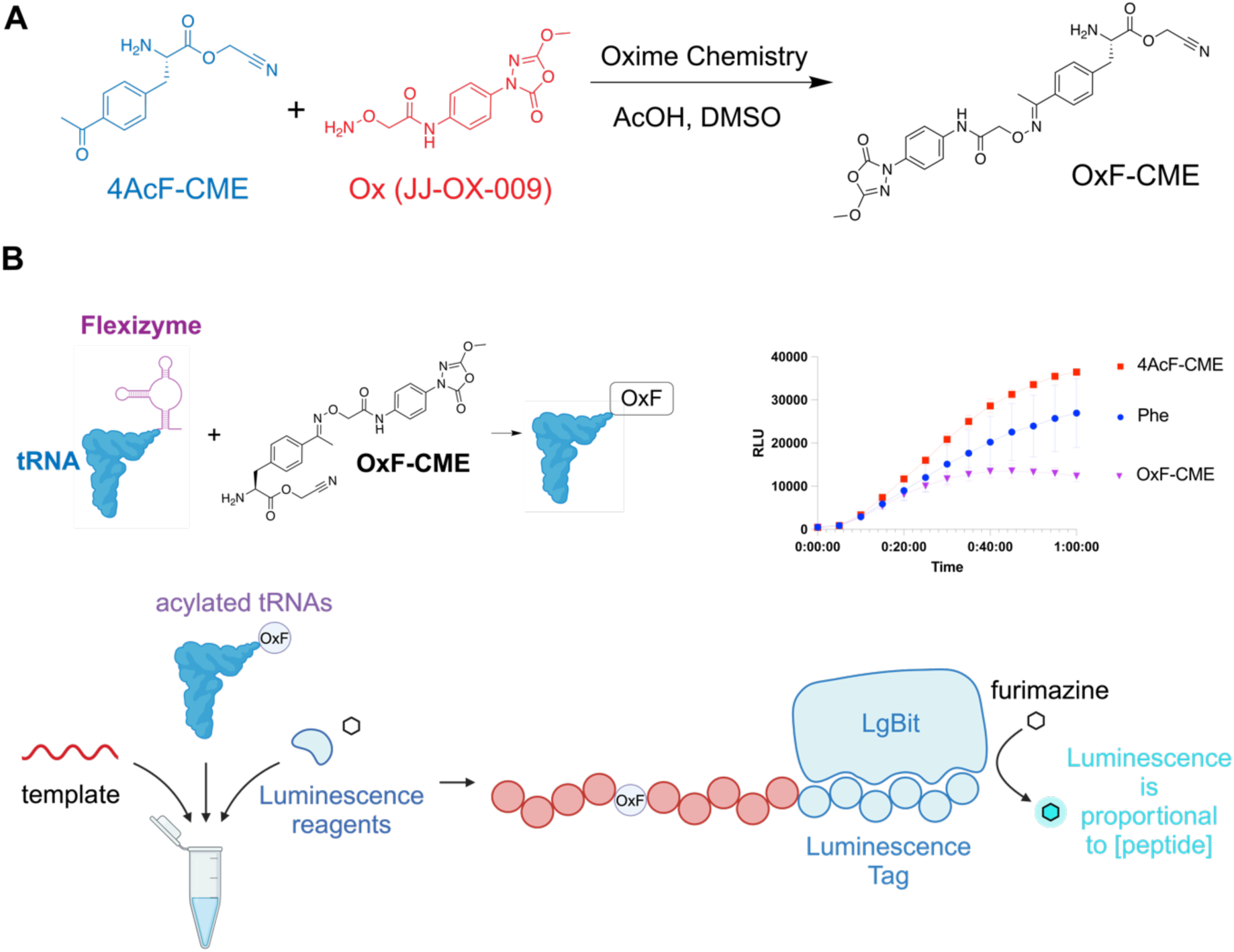
Design strategy of covalent mRNA display libraries by genetic encoding of the oxadiazolone electrophile. (A) Synthesis of 4-acetyl-phenylalanine cyanomethyl ester (OxF-CME). The parent oxadiazolone electrophile fragment containing a hydroxylamine was conjugated to the cyanomethyl ester of 4-acyl-phenylalanine (4AcF-CME) using oxime chemistry to obtain the final oxadiazolone phenylalanine cyanomethyl ester (OxF-CME) used for flexizyme-mediated acylation of tRNA. The synthesis of this compound is described in the Supporting Information. (B) Schematic of the NanoBiT luminescence-based assay to evaluate the in vitro translation (IVT) efficiency of non-natural amino acid oxadiazolone phenylalanine (OxF). The modified OxF-CME was used to acylate a tRNA using the flexizyme system followed by *in vitro* translation of a peptide containing the OxF amino acid and a luminescent peptide tag. Addition of luminescence reagents allows real-time monitoring of peptide translation. Plot shows the resulting translation rates for the natural Phe (phenylalanine; blue), compared to 4Acetyl phenylalanine (4AcF-CME; red), and the oxadiazolone modified phenylalanine (OxF-CME; purple).

#### Design and synthesis of highly diverse mRNA display libraries

To incorporate the covalent warhead into ribosomally synthesized peptides, we coupled the Ox fragment JJ-OX-009 to 4-acetyl-phenylalanine cyanomethyl ester (4AcF-CME) to yield the OxF-CME amino acid ester (Figure 1A). We tested acylation and translation of this amino acid using the NanoBiT luminescence assay^58^ and found that the amino acid was incorporated into peptides with nearly 50% translation efficiency, compared with benchmark control phenylalanine (Figure 1B). We also confirmed that the resulting peptides contained the OxF non-natural amino acid using mass spectrometry (Figure S3).^58^To screen for specific and potent binders of FphB, we used a genetically reprogrammed *in vitro* translation (IVT) system and mRNA display methods (Figure 2A). Our highly diverse libraries were comprised of 6-10 amino acids in each peptide macrocycle (Figure 2A), with each peptide initiated by an N-chloroacetylated amino acid which cyclizes with a downstream cysteine sulfhydryl group (Figure 2B), encoded by the initiator (Ini) AUG and UGG codons, respectively. In addition, each peptide displayed a single oxadiazolone warhead which is varied through each of the positions in the macrocycle (Figure 2B), encoded by the CAG codon for the OxF electrophile amino acid (Figure 2B, Figure S4). To ensure only one OxF electrophile per peptide, we used the NNU degenerate codon for all variable positions. OxF-CME was acylated onto the corresponding tRNA^Asn^ _CUG_ for translation of the electrophile OxF (Figure 2B).

**Figure 2.**
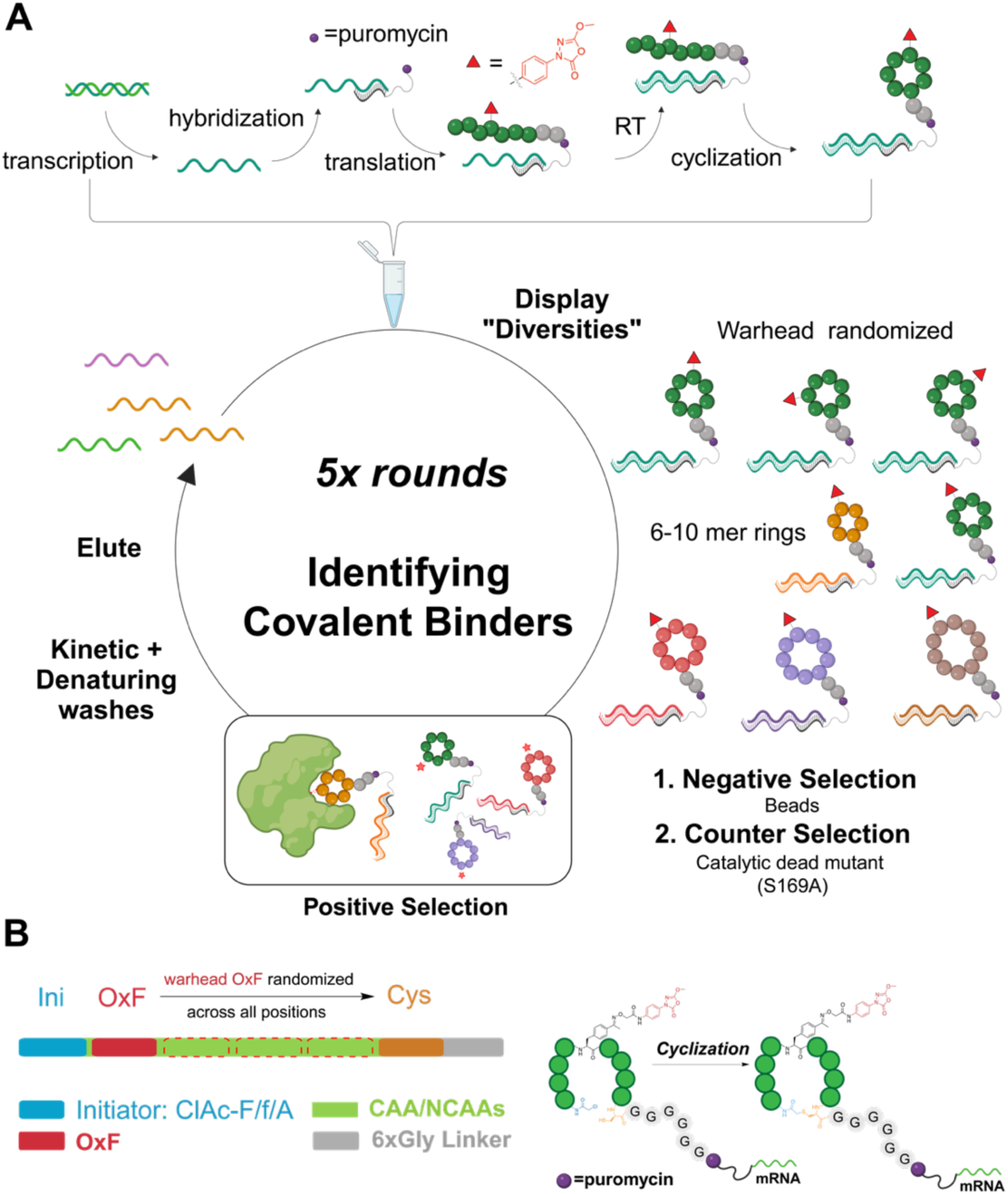
Synthesis and selections of highly diverse mRNA display libraries with a genetically encoded oxadiazolone warhead to FphB (A) The mRNA libraries were prepared following five main steps (1) DNA transcription to mRNA (2) mRNA templates hybridized to a puromycin linked cDNA (3) flexible *in vitro* translation and cyclization of peptides from mRNA template (4) reverse transcription of mRNA to generate a cDNA strand. After library synthesis, highly diverse libraries (6-10mer ring sizes with warhead randomized at all positions) were selected against WT rFphB as positive selection, and catalytic dead mutant rFphB (S169A) and beads as negative selections. Different washing stringencies (kinetic long-time washes and 5M guanidine washes) were applied at various rounds of selections. DNA elutes after each round of selection were reamplified and prepared for the next round of selection. (B) Each peptide in the library is composed of N terminal initiator ClAc-R (R=F/f/A; blue) encoded by ATG, variable region (green) and C-terminal cysteine (orange) flanked with a six-glycine linker (grey). In the variable regions, a single warhead OxF encode by CAG was used at all positions. Other canonical amino acids (CAA) and non-canonical amino acids (NCAA) were fully randomized at all other positions using the NNT codon. Peptides were automatically cyclized post translationally through the reaction of the N-terminal chloroacetylated amino acid with the single cysteine side chain thiol in each peptide.

#### Positive/Negative selections to screen for hits targeting the active site pocket

For each round of positive selection, we incubated biotinylated recombinantly expressed FphB protein (rFphB) with *in vitro* translated macrocyclic peptide mRNA display libraries at room temperature. To avoid bias in selections that favors larger ring sizes, we used cyclic peptides with 6-8 or 9-10 residues in the cycle in separate selections. Binders were isolated by affinity purification of the biotinylated rFphB along with bound library members using magnetic neutravidin beads. To increase the selection stringency and bias our hits for covalent binders, we elevated the selection stringency over subsequent rounds. Beginning in round two, we increased the wash time (kinetic wash) to disfavor binders with faster off-rates, as well as performed negative selections against neutravidin beads alone to remove non-specific bead binders (Figure 2A). In addition, from round three onwards, we performed denaturing washes in 5M guanidine-HCl after library incubation, to remove any non-covalent binding molecules. Finally, in our last round of selection, we performed parallel selections against rFphB and a catalytically inactive S169A rFphB mutant (Figure 2A). After each round of selection, we performed qPCR to quantify the total amount of recovered mRNA and then used PCR to amplify the selected sequences for the next round of screening. We performed deep sequencing of the PCR amplified DNA for each round of selection to calculate the peptide frequency and then used the complete data set to cluster hits, remove non-specific binding sequences and identify the most enriched sequences.

#### Selection of top hit sequences using informatic analysis

The results from NGS of all of the selection rounds indicated predominant enrichment of small-ring peptides (6-mers) rather than larger rings. This could be due to the fact that our target enzyme favors short chain lipid substrates^52,55^ and therefore may lack sufficient space near the active site for larger cyclic peptide, which is also indicated by the Alphafold predicted structure of FphB (data not shown). It may also reflect the fact that the covalent binding mechanism drives the association and therefore only limited interactions with the peptide scaffold are required or the fact that small ring libraries have overall lower diversity and therefore more copies of each sequence in the initial round of screening.

Using a previously reported chemoinformatic method,^59^ we clustered enriched hits from FphB screening based on peptide atomic similarity (Figure 3A). The screening results showed several enriched peptide family clusters grouped by their shared conserved peptide motifs, with the top two families depicted in Figure 3B. The most enriched binder in family 1, fOYNYC, with 2.2% enrichment in round five selection elutes, exhibited a conserved binding motif, “fOYXYC” (where f = D-Phe, O = Oxadiazolone, X = variable residue), along with other enriched members. The second family displayed the motif “FFDOXC,” with the most abundant binder, FFDOIC, showing a 0.34% enrichment post-selection. The third family contained motifs “FOXYXC” or “FONXXC.” We ultimately selected twenty-one hits (Figure S5, named as FphB-1 to FphB-21, highlighted in the cluster in Figure 3A), including two hits fOYNYC (FphB-14) and FFDOIC (FphB-5) from the top two enriched family clusters (Figure 3B), based on their representation in cluster analysis, high enrichment, sequencing profile in counter selections, and scaffold diversity.

**Figure 3.**
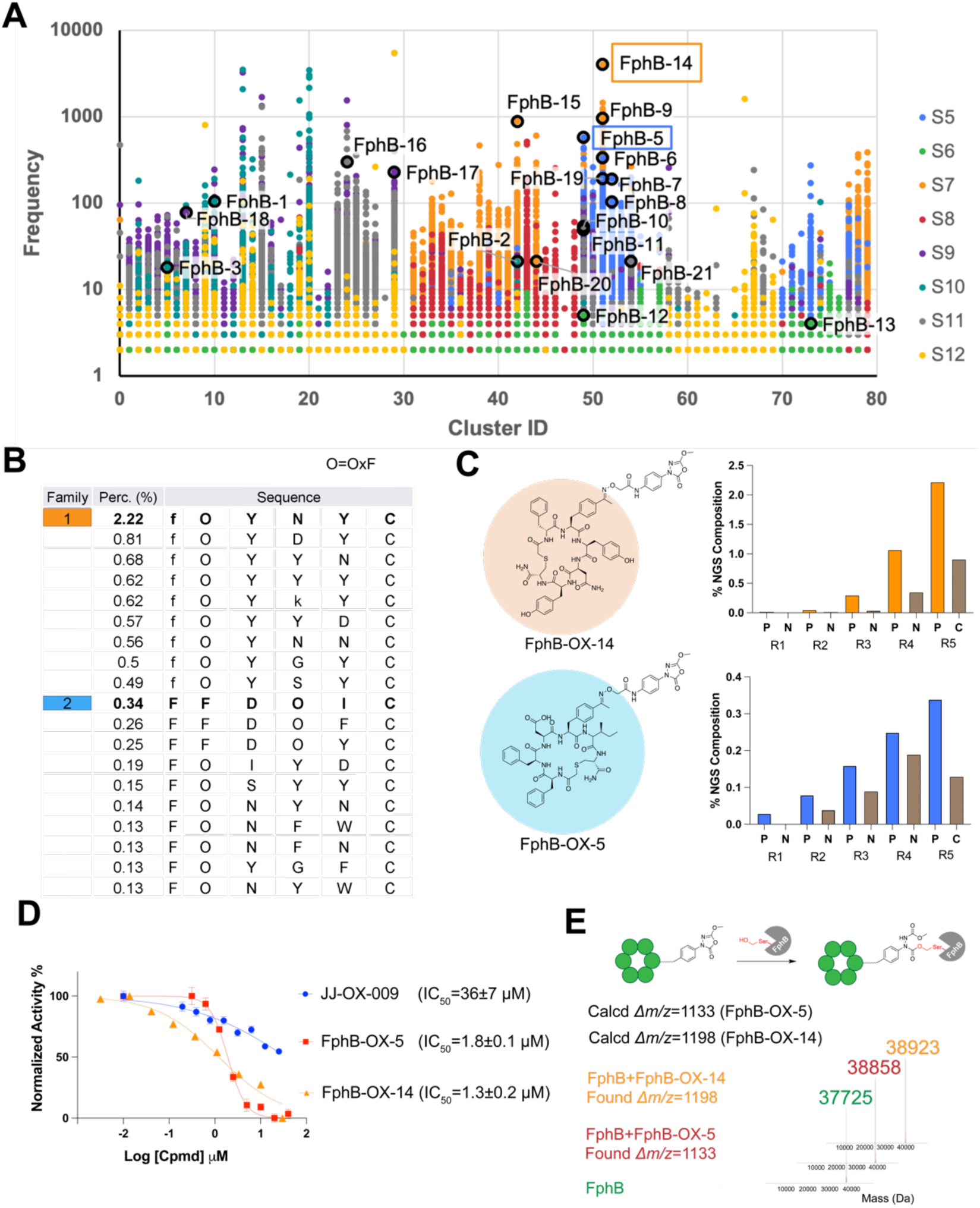
Family cluster profiling and hits from mRNA display screening to FphB. (A) Cluster analysis based on atomic similarity of enriched peptide hits. The top 21 hits selected for validation studies are circled in black in the cluster, with FphB-OX-5 and FphB-OX-14 highlighted. S5-12 represents different libraries with different codon tables (see Figure S3 for codon use). S5: 6-8mers, codon table 1; S6: 9-10mers, codon table 1; S7: 6-8mers, codon table 2; S8: 9-10mers, codon table 2; S9: 6-8mers, codon table 3; S10: 9-10mers, codon table 3; S11: 6-8mers, codon table 4; S12: 9-10mers, codon table 4. (B) Table showing sequences and enrichment of two clusters (O=OxF warhead, lowercase letters are d-amino acids, first reside is N-chloroacetyl amino acid). The percentage (Perc. (%)) of DNA copy numbers of each unique macrocycle in total reads of the NGS pool is shown. (C) Structures of the top two lead molecules and their enrichment profile in positive and negative selections in all five rounds. Round 1 uses only positive selection, rounds 2-4, beads were used as negative selections (annotated as N), round 5, S169A catalytic dead mutant rFphB was used for counter selection (annotated as C). (D) IC_50_ quantifications of JJ-OX-009, FphB-OX-5 and FphB-OX-14 for rFphB (100 nM), measured by the cleavage of the 4-MUB substrate (20 µM). The activity was normalized to a DMSO control, with points and bars representing means ± standard deviation (n=4). (E) Intact rFphB labeling with FphB-OX-5 and FphB-OX-14, validated by deconvoluted mass spectra of rFphB before (green) and after (OX-5: red, OX-14: orange) treatment. MS deconvolution analysis confirmed a single covalent modification of the protein.

### Synthesis and Direct Assay of High-yield Crude Hits at Nanomole Scale

To test the selected top twenty-one cyclic peptide hits, we first synthesized linear peptides using solid-phase peptide synthesis, followed by “N-terminus-to-side-chain” cyclization using base.^60^ Following cyclization, we conjugated the JJ-OX-009 warhead to the cyclic peptides through an oxime bond using optimized acidic conditions (Figure S6). The resulting products exhibited high purity, with yields ranging from 80% to 90%. Given the high purity and yield of the final products, we opted to scale down the synthesis for the initial round of test screening and use crude compounds for the enzymatic assays to confirm inhibition of the target. We synthesized each cyclic peptide on a nanomole scale (approximately 20-50 nmols of peptide based on resin load) and prepared the crude samples by removal of the acetic acid used for the oxime reaction by lyophilization. We then resuspended the resulting crude compounds in DMSO to achieve a ∼1 mM stock for each compound (based on resin load, product yield and purified benchmark peptides) that could be used in enzymatic assays. To evaluate the impact of impurities in the crude samples, we tested negative controls that contained residual reagents such as DIPEA (for cyclization) and acetic acid as well as positive controls (JJ-OX-004 and JJ-OX-009) prepared in a DIPEA/AcOH/DMSO alongside the hit compounds (Figure S7). The negative controls showed minimal inhibition (less than 10%), while the positive control, JJ-OX-009 and JJ-OX-004, exhibited medium-to-high activity, similar to the pure compound activities (Figure S7, Figure S2).

To further assess the robustness and feasibility of using crude compounds for initial screening, we purified several hits and compared the inhibition results with crude samples (Figure S8). The IC_50_ values of the purified and crude forms of the hits were comparable, confirming overall high yield and purity of the crude products. Because the JJ-OX-009 electrophile fragment has only weak activity for the target and constitutes less than 5% of the crude mixture (Figure S6) after oxime chemistry, this residual starting material does not create a significant background signal for the activity screens. We screened all 21 crude hits in full-dose enzymatic assays, with the majority of peptides showing promising high nanomolar to low micromolar activities (Figure S7).

### Biochemical Validations of Lead Molecules

#### FphB-OX-14 and FphB-OX-5 are Potent, Irreversible Covalent Probes, Targeting the Serine in the Active Site of FphB

We synthesized and purified several hits on a larger scale based on their diversity in cluster analysis, potency and selectivity in activity assays, and diversity of structural scaffolds. We chose the top two enriched hits from two family clusters (Figure 3A), FphB-OX-14 and FphB-OX-5 (Figure 3C), as lead compounds for further characterization. Both compounds demonstrated progressive enrichment through five rounds of positive selections over negative selections (Figure 3C), especially at round 5 with the catalytic dead mutant as counter selection. To validate the activity of the two purified hits, we conducted full-dose-dependent rFphB inhibition assays. FphB-OX-14 exhibited an IC_50_ value of 1.3 ± 0.2 µM, and FphB-OX-5 showed an IC_50_ of 1.8 ± 0.1 µM (Figure 3D). In comparison, the weakly active warhead molecule JJ-OX-009 had a IC_50_ of 36 ± 7 (or >20) µM. The compounds without warhead of (FphB-14 and FphB-5) had only weak inhibition activities (IC_50_ >20 µM; Figure S9).

To confirm the irreversible covalent binding mechanism of the two top hits, we performed both intact protein labeling using mass spectrometry and a jump dilution assay in which protein and inhibitor are pre-incubated and then diluted before measuring residual activity. Mass spectrometry analysis of the labeled protein confirmed the expected mass shift upon covalent bond formation with the Ox electrophile at the active site serine-169 (Figure 3E). Furthermore, the results of the jump dilution assay confirmed sustained irreversible inhibition of the enzyme even after dilution (Figure S10). Finally, incubation of the probes with the catalytically dead mutant rFphB in which the active site serine was replaced by alanine (rFphB S169A) confirmed that the labeling was activity-dependent and site specific.

#### Identified cyclic peptide scaffolds tune activity and selectivity of the Ox electrophile for FphB

To determine how effectively the selected cyclic peptides targeted the Ox electrophile towards rFphB, we evaluated the full-dose inhibitory potencies of FphB-OX-14 and FphB-OX-5 for rFphE (Figure 4A) and compared FphB/FphE selectivity ratios with the OX linker JJ-OX-009 and our previously reported FphE probe JJ-OX-004 (Figure S11). Notably, FphB-OX-5, demonstrated a 7-fold improvement in FphB/FphE selectivity compared to JJ-OX-004, and nearly a 50-fold improvement compared to the OX linker JJ-OX-009 (Figure S11). To assess selectivity using protein labeling, we incubated a 1:1 mixture of rFphB and rFphE proteins with FphB-OX-5, alongside control molecules JJ-OX-009 and JJ-OX-004 (Figure S12). This competition labeling was performed using the broad-spectrum serine hydrolase activity probe FP-TAMRA. All tested compounds demonstrated dose-dependent competition for rFphB and rFphE labeling, consistent with the enzymatic assays. In contrast, JJ-OX-009 and JJ-OX-004 exhibited a preference for binding to rFphE, while FphB-OX-5 comparably compete rFphB and rFphE, consistent with its improved FphB/FphE selectivity index.

**Figure 4:**
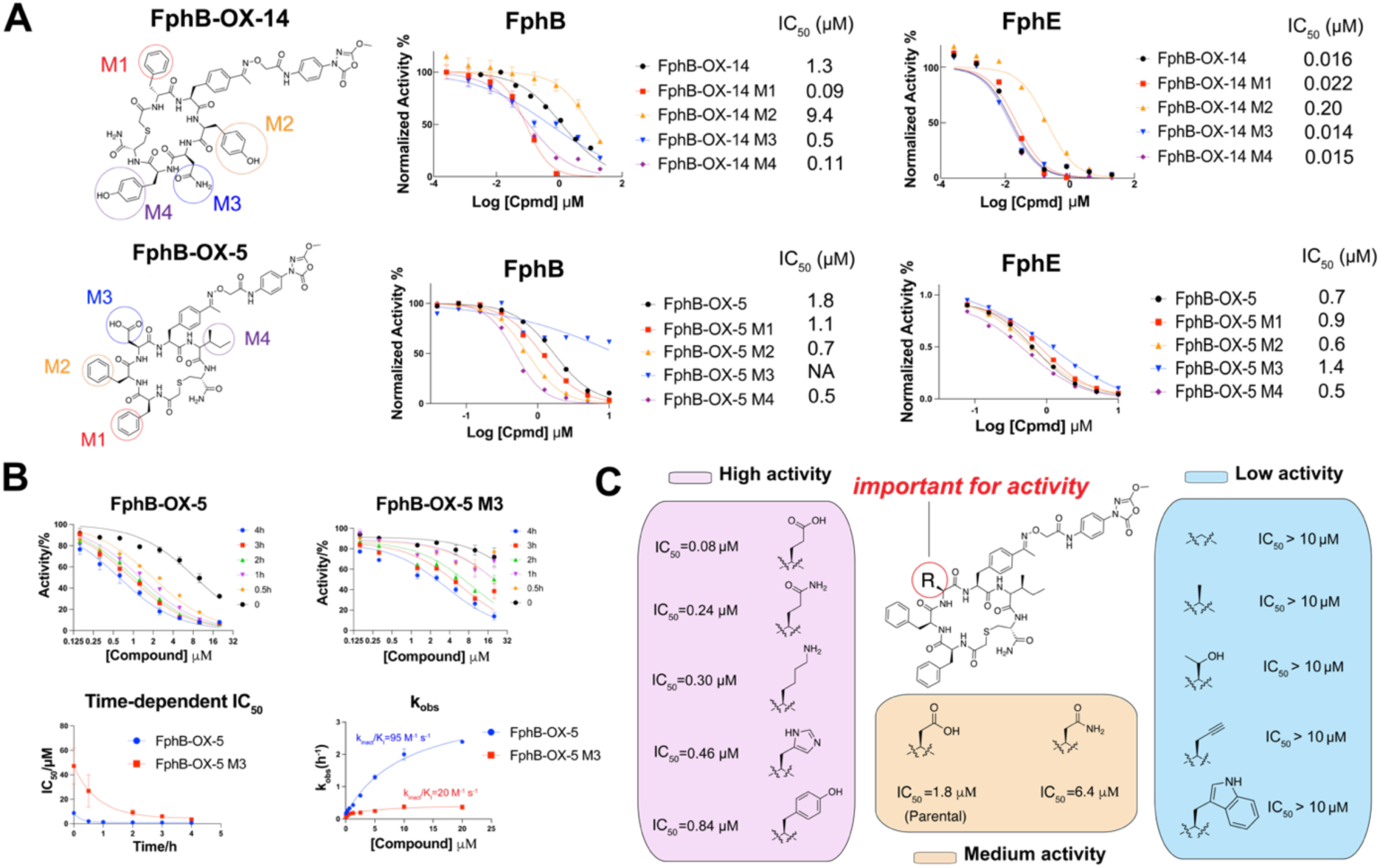
Structure-dependent activity validations of hits. (A) Each amino acid (highlighted as M1/M2/M3/M4) of FphB-OX-14 and FphB-OX-5 was individually replaced with propargylglycine to generate positional scanning alkyne mutants. Inhibitory activity of alkyne mutants with parental molecules to rFphB were quantified. (B) Time dependent enzymatic activities of OX-5 and OX-5 M3 to rFphB were quantified at six time points (0, 0.5, 1, 2, 3, and 4 hours). Second order rate constants k_inact_/K_i_ were determined using time-dependent IC_50_ values. (C) Structure-activity relationship (SAR) data for FphB-OX-5 at the M3 position. The indicated amino acids were used to individually replace the aspartic acid in the parental structure. The IC_50_ values of each compound in enzymatic assays against rFphB are listed for each compound.

#### Sequence dependent activity of the cyclic peptide FphB inhibitors

To confirm that the selected cyclic peptides direct binding to the FphB target through specific interactions between the peptide and protein, we performed a mutational scan. Instead of using the standard alanine residue, we chose to use the non-natural propargylglycine which has an alkyne sidechain. We chose this residue as it allows direct labeling studies of all the resulting mutants and the alkyne side chain has similar overall properties to alanine but can be used for labeling by CLICK chemistry. We synthesized the FphB-OX-14 (FphB-OX-14 M1-M4) and FphB-OX-5 (FphB-OX-5 M1-M4) alkyne mutants and evaluated their inhibition activities against rFphB (Figure 4A). The alkyne variant, FphB-OX-14 M3 showed a modest increase in activity (IC_50_ = 0.5 µM) while FphB-OX-14 M1 and FphB-OX-14 M4 exhibited significantly improved activities (∼100 nM IC_50_), with a nearly 10-fold increase in potency compared to the parental compound. This indicates the M1/M4 positions can be used for further chemical derivatizations and perhaps for increasing potency and selectivity for FphE. In contrast, FphB-OX-14 M2 demonstrated a 7-fold decrease in activity (IC_50_ = 9.1 µM), highlighting the importance of tyrosine at the M2 position for optimal activity. The overall large drop in potency for FphB-OX-14 M2 compared to FphB-OX-14 M4 (100-fold difference) for two macrocycles that are both modified by a single tyrosine-to-alkyne mutation, highlights the structure-dependent activities of the inhibitors.

To further confirm that the sequence specific effects of the alkyne mutants were specific to target binding and not a general effect, we tested all of the alkyne mutants and parental compounds against the off target rFphE. We found that, with the exception of FphB-OX-14 M2, all of the mutants exhibited similar potencies towards rFphE (IC_50_=10-20 nM). This result confirms that the high potency of the Ox electrophile for rFphE drives the potency of the cyclic peptides, and explains why our top cyclic peptides containing the Ox electrophile still retain potent rFphE activity.

For the second hit molecule, FphB-OX-5, the M1, M2 and M4 alkyne mutants exhibited only slight increases in activities (2-4 fold) compared to the parental compound. However, the M3 mutant, with the alkyne replacing aspartic acid, showed a dramatic loss of activity (IC_50_ > 20 µM; Figure 4A, Figure S13A). This drop in activity was due to overall slow kinetics as was confirmed by measuring the kinetic rate constant for this mutant (Figure 4B). Time-course protein labeling using in-gel fluorescence analysis and mass spec also confirmed the slower kinetics of the FphB-OX-5 M3 compound (Figure S13C, D).

Due to the significant role of this M3 position for inhibition activity against rFphB, we further made and tested a set of 10 compounds using diverse amino acids (Figure 4C). All small amino acids (G/A/T and propargylglycine) and tryptophan resulted in a substantial drop in activity (IC_50_>10 µM), while Amino acids with longer, flexible side chains that contain hydrogen bond donor/acceptors (H/Y/K) showed slightly improved activities compared to the parent hit. Interestingly, the E/Q analogs demonstrated a more than 10-fold increase in potency compared to the D/N analogs, indicating that the binding interaction between this residue and the target is likely dependent on both hydrogen bonding and side chain length. Importantly, as observed for the FphB-OX-14 mutants, all of the FphB-OX-5 alkyne mutants, including the M3 mutant, showed potencies for the off target rFphE that were virtually independent of the sequence mutations, again supporting the notion that the selected cyclic peptides form specific binding interactions with rFphB but not rFphE. We also tested the ability of all OX-14 and OX-05 alkyne mutants to label recombinant rFphB and rFphE (Figure S13A, B). These results were consistent with the activity data (Figure 4A). Overall, these results confirm the importance of specific residues on the cyclic peptide scaffold for binding to the target for which the molecules were selected.

Finally, to better understand the interactions between our selected cyclic peptides and target proteins, we carried out X-ray structural studies. Unfortunately, we were unable to obtain a structure for the intended target FphB as this protein fails to form stable crystals. However, we solved the structure of FphE bound to the FphB-OX-5 cyclic peptide (Figure S14). This structure confirmed the formation of the covalent bond between the oxadiazolone warhead and the serine residue in the active site. However, the electron density quality for the ligand atoms dramatically decreases with distance from the serine residue. This data suggests that FphB-OX-5 is bound in multiple conformations to FphE, resulting in no clear structure of the peptide part. Therefore, only the oxadiazolone warhead part was modelled in the crystal structure omitting the remaining atoms. This result is consistent with the mutational analysis that suggested little involvement of the cyclic peptide backbone in binding and inhibition of the off target FphE.

### Covalent Docking of Cyclic Peptide Probes to FphB

Because we were unable to obtain a structure for FphB we used the modeled structure from AlphaFold2 for docking simulations. In the initial structure the active site was closed, and cosolvent molecular dynamics (MD) was performed with the oxadiazolone as the cosolvent, to represent the reactive warhead. This led to a conformation with a significantly more open pocket into which the covalent warhead could be docked. The macrocyclic peptide conformations were similarly pre-sampled by MD and a diverse set of states were extracted by clustering analysis. In the case of FphB-OX-14, the addition of ethanol as a co-solvent was necessary to produce diverse conformational states. These distinct states were docked, with the macrocycle kept rigid but side chains treated flexibly to the receptor using AutoDock-GPU following the reactive docking protocol. To further distinguish between states, these complexes were then subjected to further MD simulations, with the most stable by RMSD being selected as final poses.

Overall, both predicted complexes feature the catalytic pocket occupied solely by the covalent handle, up through the oxime which connects the warhead to the peptide, while the macrocyclic peptide itself sits outside the entrance (Figure 5). For FphB-OX-05 (Figure 5A,B), the model suggests that the macrocycle conformation is stabilized by two major intramolecular hydrogen bonds: between the M1backbone carbonyl and M4 NH, and between the M2 backbone carbonyl and post-M4 NH. The M1 phenylalanine sidechain packs against I117 and P306 with K116 also lying above the phenyl ring and possibly sampling a cation-p interaction. In this conformation, the M3 aspartate is solvent exposed, and is located near K29, potentially stabilized through long range electrostatics. The polar environment predicted around M3 is supported by the activity of derivatives at this site, which largely benefits from polar or charged residues and prohibits nonpolar residues. The M4 and M2 sidechains are mostly solvent exposed, with the M2 phenyl stacking against the sidechain of M1.

**Figure 5.**
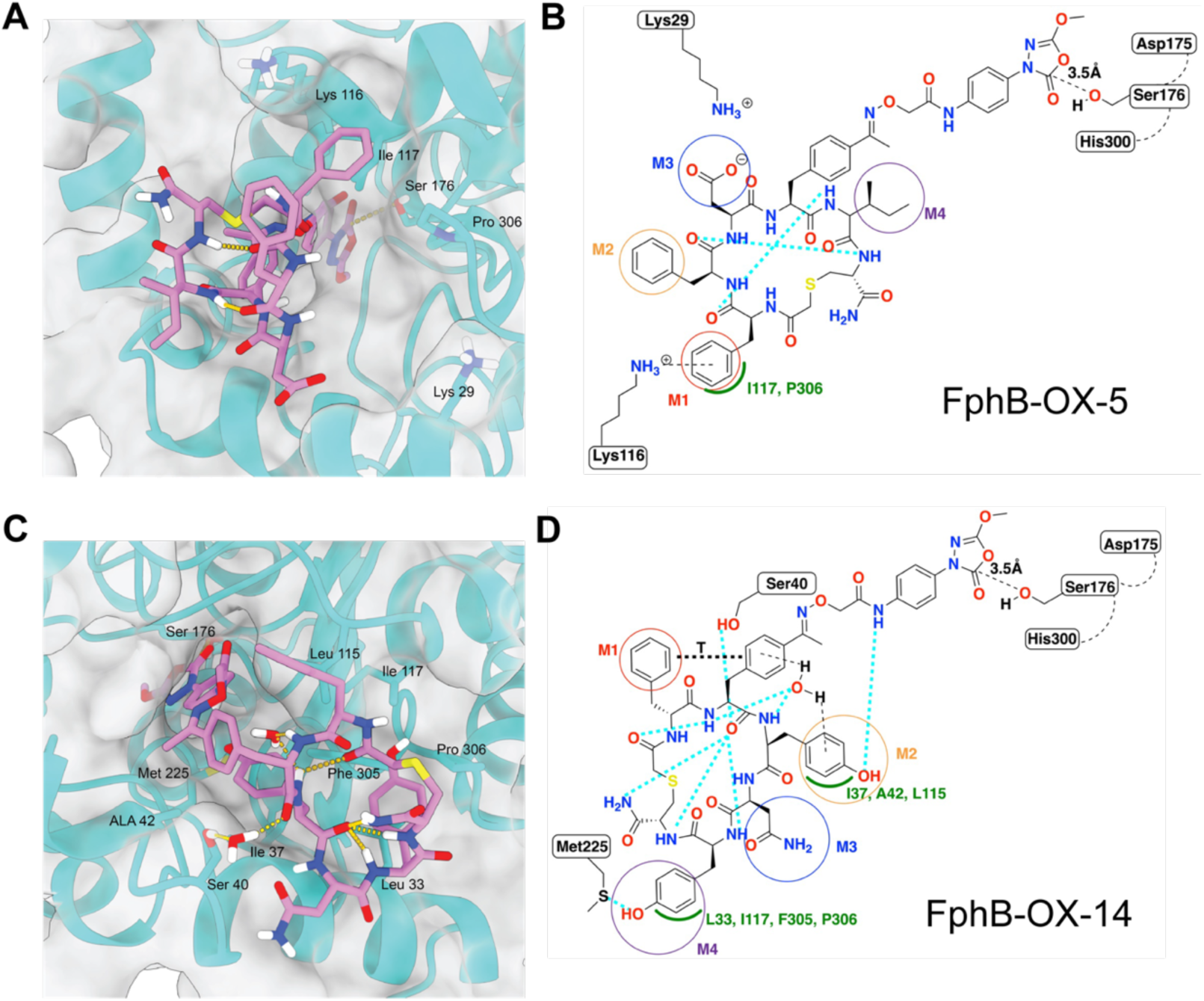
Covalent docking of FphB-OX-5 and FphB-OX-14 to FphB. (A) FphB-OX-5 simulated binding pose modeled in predicted FphB structure. Hydrogen bonds and reactive interaction with S176 shown as gold pseudobonds. (B) Interaction diagram for FphB-OX-5. Hydrogen bonds shown as cyan dashed lines. Van der Waals interactions shown by green arc. (C) FphB-OX-14 simulated binding pose modeled in predicted FphB structure. Hydrogen bonds and reactive interaction with S176 shown as gold pseudobonds. (D) Interaction diagram for FphB-OX-14. Hydrogen bonds shown as cyan dashed lines. Van der Waals interactions shown by green arc. T-stacking interaction shown as black dashed line.

For FphB-OX-14 (Figure 5C,D), the macrocycle backbone is stabilized by intramolecular hydrogen bonds between the pre-M1 carbonyl and the M2 NH, and a tridentate hydrogen bonding network between the M2 carbonyl and M4 NH, post-M4 NH, and terminal amide NH. The M1 sidechain also tends to form a T-stacking interaction with the proximal phenyl ring of the covalent handle, while the M2 phenol sidechain has a water-bridged p-HOH-p interaction with that same phenyl ring, with the water being stabilized by hydrogen bonds from the covalent handle backbone NH and M2 NH. The M2 sidechain hydroxyl also forms a hydrogen bond with the amide NH of the covalent handle. Among the interactions between the macrocycle and the protein, we observe a water-mediated hydrogen bond between the covalent handle backbone carbonyl and the S40 hydroxyl. The M2 sidechain, meanwhile, packs in the bottom of the pocket against I37, A42, and L115. M4 phenol is packed nearby that of M2, against L33, I117, F305, and P306. The M4 hydroxyl may also form an interaction with the M225 sulfur.

For both macrocycles, we computed the root-mean-squared fluctuation (RMSF) of each atom over the whole simulation time as a metric of their stability. We found the atoms of the FphB-OX-5 M1 and M3 sidechains to be most stable (Figure S15), corroborating their roles as the most essential sidechains in the mutagenesis experiments (Figure 4A) and despite their predicted high level of solvent exposure. Likewise, we also found the FphB-OX-14 M2 sidechain to be the most stable in terms of RMSF (Figure S15) and most essential by mutagenesis (Figure 4A). In both cases, the electrophilic carbon of the oxadiazalone ring was proximal to the nucleophilic oxygen of Ser176, consistent with a near attack conformation competent for covalent ligation.

### Labeling of Native FphB in Live *S. aureus* Using Covalent Cyclic Peptide Probes

The final test of the newly identified cyclic peptide probes was to assess their overall potential for imaging of live *S. aureus* bacteria. We therefore first performed direct labeling studies of live cells followed by SDS-PAGE analysis of labeled proteins to measure overall potency and selectivity of the probes for the native expressed FphB. For these studies, we added probes to bacterial cultures followed by washing of cells, lysis and click chemistry to attach a fluorescent tag and finally SDS-PAGE analysis to resolve labeled proteins. These initial labeling studies for the full set of alkyne mutants for FphB-OX-14 and -05 demonstrated that all of the OX-14 mutants primarily labeled FphB at low probe concentrations but also showed labeling of FphE as probe concentrations were increased (Figure S16A). For the OX-05 alkyne series, only the M3 mutant showed labeling of the intact bacteria (Figure S16B). This was a surprising result given that the M3 mutant was the only mutant that showed significant loss of potency for FphB. Since this mutant replaced a negatively charged aspartate residue with the uncharged alkyne, we reasoned that perhaps the negative charge on all the other probes was preventing labeling due to repulsive interactions with the negatively charge cell membranes, which is primarily attributable to the negatively charged phosphates in the lipid bilayer.^61,62^ To test this hypothesis, we synthesized analogs of the M1, M2, M4 alkyne mutants in which the aspartate at the M3 position was mutated to asparagine to remove the negative change without impacting the *in vitro* enzymatic activities and overall properties (Figure S16C). Labeling with this series of probes confirmed that elimination of the negative charge at the M3 position restored labeling of FphB by the probes in intact cells (Figure S16D) but had no effect on labeling in lysates, confirming our hypothesis. We therefore used the M1, M2, and M4 mutants of OX-05 with an asparagine in the M3 position (named M1(N), M2(N), M4(N)) going forward. For the resulting probes, we also confirmed that these compounds do not inhibit bacterial growth (Figure S17).

### Fluorescently Labeled Cyclic Peptides Can be Used to Image Live *S. aureus* Cells

We next synthesized fluorescent versions of the OX-05 and OX-14 alkyne series molecules by conjugating Cy5-azide via the CuAAC click reaction. Dose-dependent labeling of live *S. aureus* cells with these Cy5 imaging probes demonstrated selective labeling of FphB, confirmed by the *fphB* transposon (Tn; Figure S18A, B). Notably, at concentrations of 10 nM, the probes exhibited highly selective labeling of FphB (Figure 6A) that contrasted the labeling by JJ-OX-012, a selective FphE probe derivatized from JJ-OX-004^55^, which selectively labeled FphE. From the set of 10 probes tested, we selected 5 with promising potency or selectivity: FphB-OX-14 M2 Cy5, FphB-OX-14 M3 Cy5, FphB-OX-14 M4 Cy5, FphB-OX-5 M1(N) Cy5, and FphB-OX-5 M4(N) Cy5. We further evaluated these probes for dose-dependency across a range of concentrations from 3-100 nM (Figure 6B). Compared to JJ-OX-012, these probes showed high dose dependent selectivity and potency for FphB. Among these probes, FphB-OX-14 M4 Cy5 and FphB-OX-5 M4(N) Cy5 were identified as the most potent and selective based on their labeling efficiencies and potency (Figure S19A).

**Figure 6.**
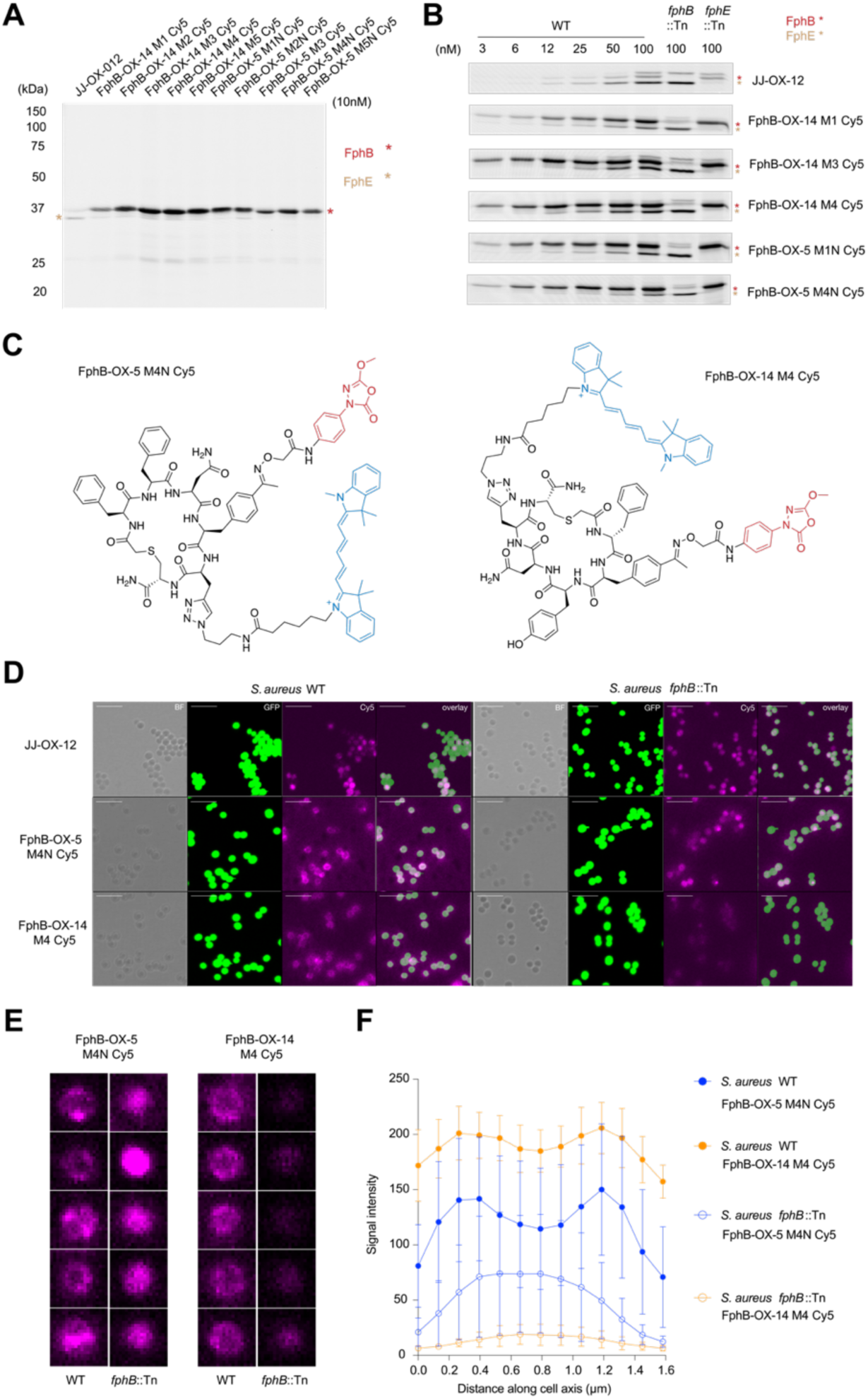
Live *S. aureus* labeling and microscope confocal imaging with fluorescent imaging probes. (A) SDS-PAGE image of live USA300 *S. aureus* cells (both wild-type and *fphB* transposon (*fphB*::Tn) mutant strains)) labeled with 10 nM of JJ-OX-012, as well as the FphB-OX-14 and FphB-OX-5 alkyne scanning mutants. Cells were treated with probes at 37 °C for 1 h before cell lysis, click chemistry labeling with Cy5-azide, SDS-PAGE and fluorescence imaging. (B) Dose-dependent labeling of USA300 *S. aureus* cells with the indicated probes at concentrations from 3 nM to 100 nM. The location of the FphB protein is shown in red, FphE shown in beige. The *fphB* transposon mutant strain (*fphB*::Tn) and *fphE* transposon mutant strain (*fphE*::Tn) are shown at the 100 nM probe concentration. (C) Structures of the two fluorescent probes FphB-OX-14 M4 Cy5 and FphB-OX-5 M4(N) Cy5. (D) Confocal micrographs of stationary phase *S.aureus* USA300-GFP cells labeled with 100 nM JJ-OX-12, FphB-OX-5 M4(N) Cy5 and FphB-OX-14 M4 Cy5. Panel 1:bright-field (BF); panel 2:GFP imaging; panel 3:Cy5 imaging; panel 4:overlay of panels GFP and Cy5. Scale bar: 10 μm (E) Representative single-cell confocal images of WT *S. aureus* USA300 (left) and *fphB* transposon (*fphB*::Tn) mutant cells (right) treated with FphB-OX-5-M4(N) Cy5 and FphB-OX-14 M4 Cy5. Scale bar: 1 μm. (F) Quantified average fluorescence intensity distribution across the cell for each probe in WT and transposon mutant cells. Microscopy images of cells were analyzed by implementing an agile script. The image was converted to a binary format, cells were identified, and the major axis length along with the mean intensity of each cell was calculated. The mean intensity was plotted against the major axis length for each identified cell in the images. Bars represent means ± standard deviation (n =6).

Based on specificity and potency, we selected the top two probes (FphB-OX-14 M4 Cy5 and FphB-OX-5 M4(N) Cy5 for imaging FphB in live cells using confocal microscopy (Figure 6C, Figure S19B). We labeled WT and *fphB*::Tn *S. aureus* cells expressing an intracellular GFP reporter with the two probes, and then imaged the cells (Figure 6D). Notably, the labeling pattern in the WT *S. aureus* strain showed the characteristic punctate staining that was enriched on the cell membrane and in the division septum consistent with FphB labeling^52,55^ (Figure 6D, E, F). For the *fphB*::Tn mutants lacking FphB expression, the probes showed broad and diffuse intracellular staining which overlapped with the GFP signal (Figure 6D, E, F), consistent with FphE labeling^55^ (Figure 6B). In support of this assessment, the selective FphE probe JJ-OX-12 showed the same broad and diffuse intracellular staining which overlapped with the GFP signal, in both WT and *fphB*::Tn *S. aureus* live cells (Figure 6D).

To evaluate any potential off-target labeling of mammalian proteins by the optimal probes, we labeled cell lysates derived from human embryonic kidney 293T (HEK293T) cells (Figure S20). When rFphB was added back to the lysates (Figure S20), it was the dominantly labeled protein. Even at probe concentrations of 1 μM, no significant, distinct proteins were labeled in HEK293T cells, with only the supplemented rFphB labeled (Figure S20). In addition, we tested the toxicity of probes in HEK293T cells and found that they were non-toxic at concentrations as high as 50 uM (Figure S21). These results demonstrate that the probes are selective for rFphB with few or no significant mammalian off-targets, highlighting their potential for future *in vivo* applications.

## CONCLUSIONS

Covalent inhibitors are powerful pharmacological tools that require optimization to be potent and selective within a biological milieu. Here, we demonstrate that it is possible to tune a relatively potent but promiscuous oxadiazolone electrophile by using genetic code reprogramming to directly incorporate the warhead into diverse libraries of macrocyclic peptide scaffolds using mRNA display. We screened diverse Ox-cyclic peptide libraries against FphB, a previously identified virulence factor in *S. aureus*. By leveraging affinity- and reactivity-driven positive and negative selections, we identified two macrocyclic hit molecules, FphB-OX-14 and FphB-OX-5, with potent covalent binding of the target enzyme and confirmed their on-target covalent binding using biochemical and cellular assays. We also used alkyne positional scanning to identify residues in both hit molecules that were critical for activity against the intended target, FphB. This enabled us to identify a suitable site for labeling with fluorescent tags for cellular labeling studies. Evaluations of optimal imaging probes in live *S. aureus* bacterial cells demonstrated that they exhibit dose-dependent covalent labeling of FphB with globally high selectivity and sensitivity. Compared with JJ-OX-12, a selective FphE imaging probe with the same oxadiazolone warhead, our macrocyclic peptide hits demonstrated dramatically increased targeting and labeling of FphB. In addition, while our hits retained activity against FphE in biochemical assays, the macrocyclic structures were able to direct binding towards FphB over FphE in live cells. In the future, counter selections against FphE could be employed to further improve the FphB/FphE selectivity ratio.

Overall, our approach of combining non-selective covalent “fragments” with focused macrocyclic libraries provides a general blueprint for the rapid generation of irreversibly binding chemical probes. Given the overall potency and selectivity of our newly optimized FphB-targeting peptides, they will likely be valuable for *S. aureus* biofilm detection and other *in vivo* applications such as site-specific delivery of antibiotics or imaging dyes that enable phototherapy applications. Covalent macrocyclic probes are also uniquely suited for targeting membrane-localized proteins beyond FphB. Given the broad spectrum of enzymatic targets (i.e. proteases, hydrolases, kinases) involved in bacterial/viral infections, immune disease, and cancer, genetically encoded electrophile libraries have great potential to identify new classes of molecular tools for the study and treatment of diverse disease conditions.

## ASSOCIATED CONTENT

### Supporting Information

Supplementary tables and figures, materials and methods for biological evaluation, detailed synthesis procedures and compound characterization, LC-MS, and NMR spectra (PDF)

### Accession Codes

The data set generated and analyzed in this study is available in the worldwide Protein Data Bank (PDB) under PDB IDs (FphE bound with FphB-OX-5). The authors will release the atomic coordinates and experimental data upon article publication.

### Funding Sources

This work was funded by an NIH grant R01 EB026332 to MB. This work was supported in part by a Sir Charles Hercus Health Research Fellowship (to M.F.). This research was undertaken in part using the MX2 beamline at the Australian Synchrotron, part of ANSTO, and made use of the ACRF detector.

## ACKNOWLEDGMENT

The authors thank Dr. Shiyu Chen and Dr. Chenzou Hao for helpful discussions. We thank Dr. Brandon Lam for helping the data export in MSD assays. We thank Sam Jamieson and Peter Mace at the University of Otago for preliminary work purifying FphB. We thank Gabriel Berger at Stanford University for the help in enzymatic assays.

## Notes

### Competing Interest Statement

The authors have declared no competing interest.

### Summary of Updates

The original file conversion created images that are low resolution and blurry.

## REFERENCES

(1) Singh, J.; Petter, R. C.; Baillie, T. A.; Whitty, A. The Resurgence of Covalent Drugs. Nat Rev Drug Discov 2011, 10 (4), 307–317. 10.1038/nrd3410.

(2) Boike, L.; Henning, N. J.; Nomura, D. K. Advances in Covalent Drug Discovery. Nat Rev Drug Discov 2022, 21 (12), 881–898. 10.1038/s41573-022-00542-z.

(3) Baillie, T. A. Targeted Covalent Inhibitors for Drug Design. Angewandte Chemie International Edition 2016, 55 (43), 13408–13421. 10.1002/anie.201601091.

(4) Ghosh, A. K.; Samanta, I.; Mondal, A.; Liu, W. R. Covalent Inhibition in Drug Discovery. ChemMedChem 2019, 14 (9), 889–906. 10.1002/cmdc.201900107.

(5) Schwartz, P. A.; Kuzmic, P.; Solowiej, J.; Bergqvist, S.; Bolanos, B.; Almaden, C.; Nagata, A.; Ryan, K.; Feng, J.; Dalvie, D.; Kath, J. C.; Xu, M.; Wani, R.; Murray, B. W. Covalent EGFR Inhibitor Analysis Reveals Importance of Reversible Interactions to Potency and Mechanisms of Drug Resistance. Proceedings of the National Academy of Sciences 2014, 111 (1), 173–178. 10.1073/pnas.1313733111.

(6) Ward, R. A.; Fawell, S.; Floc’h, N.; Flemington, V.; McKerrecher, D.; Smith, P. D. Challenges and Opportunities in Cancer Drug Resistance. Chem. Rev. 2021, 121 (6), 3297–3351. 10.1021/acs.chemrev.0c00383.

(7) Anand, K.; Ziebuhr, J.; Wadhwani, P.; Mesters, J. R.; Hilgenfeld, R. Coronavirus Main Proteinase (3CLpro) Structure: Basis for Design of Anti-SARS Drugs. Science 2003, 300 (5626), 1763–1767. 10.1126/science.1085658.

(8) Yoo, D. Y.; Hauser, A. D.; Joy, S. T.; Bar-Sagi, D.; Arora, P. S. Covalent Targeting of Ras G12C by Rationally Designed Peptidomimetics. ACS Chem. Biol. 2020, 15 (6), 1604–1612. 10.1021/acschembio.0c00204.

(9) Kathman, S. G.; Xu, Z.; Statsyuk, A. V. A Fragment-Based Method to Discover Irreversible Covalent Inhibitors of Cysteine Proteases. J. Med. Chem. 2014, 57 (11), 4969–4974. 10.1021/jm500345q.

(10) Resnick, E.; Bradley, A.; Gan, J.; Douangamath, A.; Krojer, T.; Sethi, R.; Geurink, P. P.; Aimon, A.; Amitai, G.; Bellini, D.; Bennett, J.; Fairhead, M.; Fedorov, O.; Gabizon, R.; Gan, J.; Guo, J.; Plotnikov, A.; Reznik, N.; Ruda, G. F.; Díaz-Sáez, L.; Straub, V. M.; Szommer, T.; Velupillai, S.; Zaidman, D.; Zhang, Y.; Coker, A. R.; Dowson, C. G.; Barr, H. M.; Wang, C.; Huber, K. V. M.; Brennan, P. E.; Ovaa, H.; von Delft, F.; London, N. Rapid Covalent-Probe Discovery by Electrophile-Fragment Screening. J. Am. Chem. Soc. 2019, 141 (22), 8951–8968. 10.1021/jacs.9b02822.

(11) Imhoff, R. D.; Patel, R.; Safdar, M. H.; Jones, H. B. L.; Pinto-Fernandez, A.; Vendrell, I.; Chen, H.; Muli, C. S.; Krabill, A. D.; Kessler, B. M.; Wendt, M. K.; Das, C.; Flaherty, D. P. Covalent Fragment Screening and Optimization Identifies the Chloroacetohydrazide Scaffold as Inhibitors for Ubiquitin C-Terminal Hydrolase L1. J. Med. Chem. 2024, 67 (6), 4496–4524. 10.1021/acs.jmedchem.3c01661.

(12) Lu, W.; Kostic, M.; Zhang, T.; Che, J.; Patricelli, M. P.; Jones, L. H.; Chouchani, E. T.; Gray, N. S. Fragment-Based Covalent Ligand Discovery. RSC Chem Biol 2 (2), 354–367. 10.1039/d0cb00222d.

(13) Zambaldo, C.; Daguer, J.-P.; Saarbach, J.; Barluenga, S.; Winssinger, N. Screening for Covalent Inhibitors Using DNA-Display of Small Molecule Libraries Functionalized with Cysteine Reactive Moieties. *Med*. Chem. Commun. 2016, 7 (7), 1340–1351. 10.1039/C6MD00242K.

(14) Daguer, J.-P.; Zambaldo, C.; Abegg, D.; Barluenga, S.; Tallant, C.; Müller, S.; Adibekian, A.; Winssinger, N. Identification of Covalent Bromodomain Binders through DNA Display of Small Molecules. Angewandte Chemie International Edition 2015, 54 (20), 6057–6061. 10.1002/anie.201412276.

(15) Guilinger, J. P.; Archna, A.; Augustin, M.; Bergmann, A.; Centrella, P. A.; Clark, M. A.; Cuozzo, J. W.; Däther, M.; Guié, M.-A.; Habeshian, S.; Kiefersauer, R.; Krapp, S.; Lammens, A.; Lercher, L.; Liu, J.; Liu, Y.; Maskos, K.; Mrosek, M.; Pflügler, K.; Siegert, M.; Thomson, H. A.; Tian, X.; Zhang, Y.; Konz Makino, D. L.; Keefe, A. D. Novel Irreversible Covalent BTK Inhibitors Discovered Using DNA-Encoded Chemistry. Bioorganic & Medicinal Chemistry 2021, 42, 116223. 10.1016/j.bmc.2021.116223.

(16) Li, L.; Su, M.; Lu, W.; Song, H.; Liu, J.; Wen, X.; Suo, Y.; Qi, J.; Luo, X.; Zhou, Y.-B.; Liao, X.-H.; Li, J.; Lu, X. Triazine-Based Covalent DNA-Encoded Libraries for Discovery of Covalent Inhibitors of Target Proteins. ACS Med. Chem. Lett. 2022, 13 (10), 1574–1581. 10.1021/acsmedchemlett.2c00127.

(17) Wilson, D. S.; Keefe, A. D.; Szostak, J. W. The Use of mRNA Display to Select High-Affinity Protein-Binding Peptides. Proceedings of the National Academy of Sciences 2001, 98 (7), 3750–3755. 10.1073/pnas.061028198.

(18) Chen, S.; Lovell, S.; Lee, S.; Fellner, M.; Mace, P. D.; Bogyo, M. Identification of Highly Selective Covalent Inhibitors by Phage Display. Nat Biotechnol 2021, 39 (4), 490–498. 10.1038/s41587-020-0733-7.

(19) Iskandar, S. E.; Chiou, L. F.; Leisner, T. M.; Shell, D. J.; Norris-Drouin, J. L.; Vaziri, C.; Pearce, K. H.; Bowers, A. A. Identification of Covalent Cyclic Peptide Inhibitors in mRNA Display. J. Am. Chem. Soc. 2023. 10.1021/jacs.3c04833.

(20) Barglow, K. T.; Cravatt, B. F. Discovering Disease-Associated Enzymes by Proteome Reactivity Profiling. Chemistry & Biology 2004, 11 (11), 1523–1531. 10.1016/j.chembiol.2004.08.023.

(21) Barglow, K. T.; Cravatt, B. F. Activity-Based Protein Profiling for the Functional Annotation of Enzymes. Nat Methods 2007, 4 (10), 822–827. 10.1038/nmeth1092.

(22) Berger, A. B.; Vitorino, P. M.; Bogyo, M. Activity-Based Protein Profiling. Am J Pharmacogenomics 2004, 4 (6), 371–381. 10.2165/00129785-200404060-00004.

(23) Imhoff, R. D.; Rosenthal, M. R.; Ashraf, K.; Bhanot, P.; Ng, C. L.; Flaherty, D. P. Identification of Covalent Fragment Inhibitors for *Plasmodium Falciparum* UCHL3 with Anti-Malarial Efficacy. Bioorganic & Medicinal Chemistry Letters 2023, 94, 129458. 10.1016/j.bmcl.2023.129458.

(24) Carle, V.; Kong, X.-D.; Comberlato, A.; Edwards, C.; Díaz-Perlas, C.; Heinis, C. Generation of a 100-Billion Cyclic Peptide Phage Display Library Having a High Skeletal Diversity. *Protein Engineering*, Design and Selection 2021, 34, gzab018. 10.1093/protein/gzab018.

(25) Takahashi, T. T.; Austin, R. J.; Roberts, R. W. mRNA Display: Ligand Discovery, Interaction Analysis and Beyond. Trends in Biochemical Sciences 2003, 28 (3), 159–165. 10.1016/S0968-0004(03)00036-7.

(26) Franzini, R. M.; Neri, D.; Scheuermann, J. DNA-Encoded Chemical Libraries: Advancing beyond Conventional Small-Molecule Libraries. Acc. Chem. Res. 2014, 47 (4), 1247–1255. 10.1021/ar400284t.

(27) Lan, T.; Peng, C.; Yao, X.; Chan, R. S. T.; Wei, T.; Rupanya, A.; Radakovic, A.; Wang, S.; Chen, S.; Lovell, S.; Snyder, S. A.; Bogyo, M.; Dickinson, B. C. Discovery of Thioether-Cyclized Macrocyclic Covalent Inhibitors by mRNA Display. J. Am. Chem. Soc. 2024. 10.1021/jacs.4c07851.

(28) Goto, Y.; Katoh, T.; Suga, H. Flexizymes for Genetic Code Reprogramming. Nat Protoc 2011, 6 (6), 779–790. 10.1038/nprot.2011.331.

(29) Shimizu, Y.; Inoue, A.; Tomari, Y.; Suzuki, T.; Yokogawa, T.; Nishikawa, K.; Ueda, T. Cell-Free Translation Reconstituted with Purified Components. Nat Biotechnol 2001, 19 (8), 751–755. 10.1038/90802.

(30) Bessho, Y.; Hodgson, D. R. W.; Suga, H. A tRNA Aminoacylation System for Non-Natural Amino Acids Based on a Programmable Ribozyme. Nat Biotechnol 2002, 20 (7), 723–728. 10.1038/nbt0702-723.

(31) Hipolito, C. J.; Suga, H. Ribosomal Production and in Vitro Selection of Natural Product-like Peptidomimetics: The FIT and RaPID Systems. Current Opinion in Chemical Biology 2012, 16 (1), 196–203. 10.1016/j.cbpa.2012.02.014.

(32) Goto, Y.; Suga, H. The RaPID Platform for the Discovery of Pseudo-Natural Macrocyclic Peptides. Acc. Chem. Res. 2021, 54 (18), 3604–3617. 10.1021/acs.accounts.1c00391.

(33) Garcia Jimenez, D.; Poongavanam, V.; Kihlberg, J. Macrocycles in Drug Discovery─Learning from the Past for the Future. J. Med. Chem. 2023, 66 (8), 5377–5396. 10.1021/acs.jmedchem.3c00134.

(34) Nielsen, D. S.; Shepherd, N. E.; Xu, W.; Lucke, A. J.; Stoermer, M. J.; Fairlie, D. P. Orally Absorbed Cyclic Peptides. Chem. Rev. 2017, 117 (12), 8094–8128. 10.1021/acs.chemrev.6b00838.

(35) Kwon, Y.-U.; Kodadek, T. Quantitative Comparison of the Relative Cell Permeability of Cyclic and Linear Peptides. Chemistry & Biology 2007, 14 (6), 671–677. 10.1016/j.chembiol.2007.05.006.

(36) Vinogradov, A. A.; Yin, Y.; Suga, H. Macrocyclic Peptides as Drug Candidates: Recent Progress and Remaining Challenges. J. Am. Chem. Soc. 2019, 141 (10), 4167–4181. 10.1021/jacs.8b13178.

(37) Ross, N. C.; Reilley, K. J.; Murray, T. F.; Aldrich, J. V.; McLaughlin, J. P. Novel Opioid Cyclic Tetrapeptides: Trp Isomers of CJ-15,208 Exhibit Distinct Opioid Receptor Agonism and Short-Acting κ Opioid Receptor Antagonism. Br J Pharmacol 2012, 165 (4b), 1097–1108. 10.1111/j.1476-5381.2011.01544.x.

(38) Saito, T.; Hirai, H.; Kim, Y.-J.; Kojima, Y.; Matsunaga, Y.; Nishida, H.; Sakakibara, T.; Suga, O.; Sujaku, T.; Kojima, N. CJ-15,208, a Novel Kappa Opioid Receptor Antagonist from a Fungus, Ctenomyces Serratus ATCC15502. J Antibiot (Tokyo) 2002, 55 (10), 847–854. 10.7164/antibiotics.55.847.

(39) Cai, M.; Stankova, M.; Muthu, D.; Mayorov, A.; Yang, Z.; Trivedi, D.; Cabello, C.; Hruby, V. J. An Unusual Conformation of γ-Melanocyte-Stimulating Hormone Analogues Leads to a Selective Human Melanocortin 1 Receptor Antagonist for Targeting Melanoma Cells. Biochemistry 2013, 52 (4), 752–764. 10.1021/bi300723f.

(40) Liu, T.; Liu, Y.; Kao, H.-Y.; Pei, D. Membrane Permeable Cyclic Peptidyl Inhibitors against Human Peptidylprolyl Isomerase Pin1. J. Med. Chem. 2010, 53 (6), 2494–2501. 10.1021/jm901778v.

(41) Naumann, T. A.; Tavassoli, A.; Benkovic, S. J. Genetic Selection of Cyclic Peptide Dam Methyltransferase Inhibitors. ChemBioChem 2008, 9 (2), 194–197. 10.1002/cbic.200700561.

(42) Lin, K.-H.; Ali, A.; Rusere, L.; Soumana, D. I.; Kurt Yilmaz, N.; Schiffer, C. A. Dengue Virus NS2B/NS3 Protease Inhibitors Exploiting the Prime Side. J Virol 2017, 91 (10), e00045–17. 10.1128/JVI.00045-17.

(43) Takagi, Y.; Matsui, K.; Nobori, H.; Maeda, H.; Sato, A.; Kurosu, T.; Orba, Y.; Sawa, H.; Hattori, K.; Higashino, K.; Numata, Y.; Yoshida, Y. Discovery of Novel Cyclic Peptide Inhibitors of Dengue Virus NS2B-NS3 Protease with Antiviral Activity. Bioorg Med Chem Lett 2017, 27 (15), 3586–3590. 10.1016/j.bmcl.2017.05.027.

(44) Tambunan, U. S. F.; Alamudi, S. Designing Cyclic Peptide Inhibitor of Dengue Virus NS3-NS2B Protease by Using Molecular Docking Approach. Bioinformation 2010, 5 (6), 250–254.

(45) Murugan, R. N.; Park, J.-E.; Lim, D.; Ahn, M.; Cheong, C.; Kwon, T.; Nam, K.-Y.; Choi, S. H.; Kim, B. Y.; Yoon, D.-Y.; Yaffe, M. B.; Yu, D.-Y.; Lee, K. S.; Bang, J. K. Development of Cyclic Peptomer Inhibitors Targeting the Polo-Box Domain of Polo-like Kinase 1. Bioorg Med Chem 2013, 21 (9), 2623–2634. 10.1016/j.bmc.2013.02.020.

(46) Baek, S.; Kutchukian, P. S.; Verdine, G. L.; Huber, R.; Holak, T. A.; Lee, K. W.; Popowicz, G. M. Structure of the Stapled P53 Peptide Bound to Mdm2. J. Am. Chem. Soc. 2012, 134 (1), 103–106. 10.1021/ja2090367.

(47) Chang, Y. S.; Graves, B.; Guerlavais, V.; Tovar, C.; Packman, K.; To, K.-H.; Olson, K. A.; Kesavan, K.; Gangurde, P.; Mukherjee, A.; Baker, T.; Darlak, K.; Elkin, C.; Filipovic, Z.; Qureshi, F. Z.; Cai, H.; Berry, P.; Feyfant, E.; Shi, X. E.; Horstick, J.; Annis, D. A.; Manning, A. M.; Fotouhi, N.; Nash, H.; Vassilev, L. T.; Sawyer, T. K. Stapled α-Helical Peptide Drug Development: A Potent Dual Inhibitor of MDM2 and MDMX for P53-Dependent Cancer Therapy. Proc Natl Acad Sci U S A 2013, 110 (36), E3445–3454. 10.1073/pnas.1303002110.

(48) Krzyzanowski, A.; Esser, L. M.; Willaume, A.; Prudent, R.; Peter, C.; ‘t Hart, P.; Waldmann, H. Development of Macrocyclic PRMT5–Adaptor Protein Interaction Inhibitors. J. Med. Chem. 2022, 65 (22), 15300–15311. 10.1021/acs.jmedchem.2c01273.

(49) Cardote, T. A. F.; Ciulli, A. Cyclic and Macrocyclic Peptides as Chemical Tools To Recognise Protein Surfaces and Probe Protein–Protein Interactions. ChemMedChem 2016, 11 (8), 787–794. 10.1002/cmdc.201500450.

(50) Gavenonis, J.; Sheneman, B. A.; Siegert, T. R.; Eshelman, M. R.; Kritzer, J. A. Comprehensive Analysis of Loops at Protein-Protein Interfaces for Macrocycle Design. Nat Chem Biol 2014, 10 (9), 716–722. 10.1038/nchembio.1580.

(51) Mathiesen, I.; Calder, E.; Kunzelmann, S.; Walport, L. Discovering Covalent Cyclic Peptide Inhibitors of Peptidyl Arginine Deiminase 4 (PADI4) Using mRNA-Display with a Genetically Encoded Electrophilic Warhead. ChemRxiv August 16, 2024. 10.26434/chemrxiv-2024-w8nbl.

(52) Lentz, C. S.; Sheldon, J. R.; Crawford, L. A.; Cooper, R.; Garland, M.; Amieva, M. R.; Weerapana, E.; Skaar, E. P.; Bogyo, M. Identification of a S. Aureus Virulence Factor by Activity-Based Protein Profiling (ABPP). Nat Chem Biol 2018, 14 (6), 609–617. 10.1038/s41589-018-0060-1.

(53) Reddy, P. N.; Srirama, K.; Dirisala, V. R. An Update on Clinical Burden, Diagnostic Tools, and Therapeutic Options of Staphylococcus Aureus. Infect Dis (Auckl) 2017, 10, 1179916117703999. 10.1177/1179916117703999.

(54) Tong, S. Y. C.; Davis, J. S.; Eichenberger, E.; Holland, T. L.; Fowler, V. G. Staphylococcus Aureus Infections: Epidemiology, Pathophysiology, Clinical Manifestations, and Management. Clinical Microbiology Reviews 2015, 28 (3), 603–661. 10.1128/cmr.00134-14.

(55) Jo, J.; Upadhyay, T.; Woods, E. C.; Park, K. W.; Pedowitz, N. J.; Jaworek-Korjakowska, J.; Wang, S.; Valdez, T. A.; Fellner, M.; Bogyo, M. Development of Oxadiazolone Activity-Based Probes Targeting FphE for Specific Detection of Staphylococcus Aureus Infections. J. Am. Chem. Soc. 2024, 146 (10), 6880–6892. 10.1021/jacs.3c13974.

(56) Faucher, F.; Bennett, J. M.; Bogyo, M.; Lovell, S. Strategies for Tuning the Selectivity of Chemical Probes That Target Serine Hydrolases. Cell Chemical Biology 2020, 27 (8), 937–952. 10.1016/j.chembiol.2020.07.008.

(57) Bakker, A. T.; Kotsogianni, I.; Mirenda, L.; Straub, V. M.; Avalos, M.; van den Berg, R. J. B. H. N.; Florea, B. I.; van Wezel, G. P.; Janssen, A. P. A.; Martin, N. I.; van der Stelt, M. Chemical Proteomics Reveals Antibiotic Targets of Oxadiazolones in MRSA. J. Am. Chem. Soc. 2023, 145 (2), 1136–1143. 10.1021/jacs.2c10819.

(58) Chan, A. I.; Sawant, M. S.; Burdick, D. J.; Tom, J.; Song, A.; Cunningham, C. N. Evaluating Translational Efficiency of Noncanonical Amino Acids to Inform the Design of Druglike Peptide Libraries. ACS Chem. Biol. 2023, acschembio.2c00712. 10.1021/acschembio.2c00712.

(59) Lee, M.-L.; Farag, S.; Del Cid, J. S.; Bashore, C.; Hallenbeck, K. K.; Gobbi, A.; Cunningham, C. N. Identification of Macrocyclic Peptide Families from Combinatorial Libraries Containing Noncanonical Amino Acids Using Cheminformatics and Bioinformatics Inspired Clustering. ACS Chem. Biol. 2023, 18 (6), 1425–1434. 10.1021/acschembio.3c00159.

(60) Goto, Y.; Ohta, A.; Sako, Y.; Yamagishi, Y.; Murakami, H.; Suga, H. Reprogramming the Translation Initiation for the Synthesis of Physiologically Stable Cyclic Peptides. ACS Chem. Biol. 2008, 3 (2), 120–129. 10.1021/cb700233t.

(61) Gross, M.; Cramton, S. E.; Götz, F.; Peschel, A. Key Role of Teichoic Acid Net Charge inStaphylococcus Aureus Colonization of Artificial Surfaces. Infection and Immunity 2001, 69 (5), 3423–3426. 10.1128/iai.69.5.3423-3426.2001.

(62) Clarke, A. J.; Dupont, C. *O* -Acetylated Peptidoglycan: Its Occurrence, Pathobiological Significance, and Biosynthesis. Can. J. Microbiol. 1992, 38 (2), 85–91. 10.1139/m92-014.

