## Supporting Information for "An mRNA Display Approach for Covalent Targeting of a *Staphylococcus aureus* Virulence Factor"

**Table of Contents** **Page**

1. **Supplementary Tables and Figures**


**Table S1**: FphE data collection and processing S5

**Table S2**: FphE structure solution and refinement S6

**Figure S1**: Amino acid sequence alignment for Fph proteins S7

**Figure S2**: Enzymatic inhibition activity and covalent labeling of JJ-OX-004 and JJ-OX-009 to FphB and FphE S8

**Figure S3**: *In vitro* translation (IVT) efficiency evaluation of OxF-CME using NanoBiT luminescence assay S9

**Figure S4**: Designed codon tables for mRNA display libraries S10

**Figure S5**: Linear sequences of twenty-one hits from mRNA display screening to FphB S11

**Figure S6**: Synthesis of 21 compounds and final product yield validation using LCMS S12

**Figure S7**: Direct enzymatic assay of 21 crude hits S13

**Figure S8**: Comparison of purified and crude compounds' activities in enzymatic assays to rFphB S14

**Figure S9**: Enzymatic assays of non-warhead control molecules FphB-14 and FphB-5 S15

**Figure S10**: Jump dilution assay of FphB-OX-5 and FphB-OX-14 S16

**Figure S11:** Normalized selectivity index of FphB-OX-14, FphB-OX-5 and their alkyne scanning mutants to FphB over FphE, compared with JJ-OX-009 and JJ-OX-004 S17

**Figure S12:** Dose-dependent competition labeling of FphB-OX-5, FphB-OX-2 to rFphB/rFphE mixture S18

**Figure S13:** Protein labeling validations of FphB-OX-5 and FphB-OX-14 alkyne mutants to FphB using in-gel fluorescence and mass spectrometry S19

**Figure S14:** X-ray crystal structure of FphE in complex with FphB-OX-5 S20

**Figure S15:** Covalent docking of FphB with FphB-OX-14 and FphB-OX-5 S21

**Figure S16:** SDS-PAGE analysis of live S. aureus labeling with FphB-OX-5 and FphB-OX-14 alkyne probes S22
**Figure S17:** Evaluating FphB-OX-14, FphB-OX-5 and their alkyne derivatives in S. aureus bacterial growth inhibition assay S24
**Figure S18:** Live S. aureus cell labeling using JJ-OX-012, FphB-OX-14 and FphB-OX-5 fluorescent probes S25
**Figure S19:** Quantifications of dose-dependent live cell labeling using the imaging probes S27
**Figure S20:** SDS-PAGE analysis of rFphB labeling with FphB-OX-14 M4 Cy5 and FphB-OX-5 M4N Cy5 in HEK293T cell lysate S28
**Figure S21:** Evaluation of cytotoxicity of FphB-OX-14 and FphB-OX-5 compounds to HEK293T cells S29

**2. Materials and Methods** **S30**
FphB cloning, expression and purification S30
Luminescence-based assay evaluating in vitro translation efficiency of non-canonical amino acids S31
MALDI-TOF mass spectrometry of in vitro translated peptides S31
Synthesis of Aminoacyl-amino acids-tRNA S32
Ribosomal synthesis of mRNA displayed peptide macrocycle library S32
Affinity selection of mRNA display libraries against FphB S33
Enzyme inhibition assay S33
In-gel fluorescence labeling of purified recombinant proteins S35
Mass spectrometry validation of covalent FphB labeling with chemical probes S35
Jump dilution assay S35
FphE X-ray Crystallography S36
FphE data collection, processing, refinement, deposition and analysis S36
FphB structure modeling S37
FphB-OX-5 and FphB-OX-14 model predictions using covalent docking S37
Bacterial culture and bacterial growth inhibition assay S39
Live *S. aureus* or lysate labeling S39

Confocal microscopy S40
Quantitative analysis of microscopy images S41
HEK293T cell labeling S41
HEK293T cytotoxicity assay S42

**3. Chemical Synthesis S43**
General methods S43
**Scheme S1**: Synthesis of JJ-OX-009 S44
**Scheme S2**: Synthesis of OxF-CME S44
**Scheme S3**: Synthesis of 21 macrocyclic hits from mRNA screening S44
Synthesis of JJ-OX-009 S45
Synthesis of OxF-CME S46
Synthesis of macrocyclic hits S47
Synthesis of fluorescent imaging probes S48

**4. LC-MS of OxF-CME, JJ-OX-009 and FphB chemical probes S50**
Compound Summary Table S50
LC-MS traces S52

**5. $^{1}$H NMR Spectra S87**

**6. $^{13}$C NMR Spectra S88**

**7. References S89**

#### 1. Supplementary Figures and Tables

##### Table S1: FphE data collection and processing

Values for the outer shell are given in parentheses.

| | Compound FphB-OX-5 bound |
|---|---|
| PDB ID | 9DRO |
| Diffraction source | Australian synchrotron MX2 |
| Wavelength (Å) | 0.954 |
| Detector | DECTRIS EIGER X 16M |
| Space group | $P\ 1\ 2_1\ 1$ |
| a, b, c (Å) | 47.1, 74.6, 73.6 |
| α, β, γ (°) | 90.0, 91.0, 90.0 |
| Resolution range (Å) | 47.10 – 1.54 (1.57 – 1.54) |
| Total No. of reflections | 513296 (23000) |
| No. of unique reflections | 75123 (3496) |
| Completeness (%) | 99.7 (94.1) |
| Redundancy | 6.8 (6.6) |
| $\langle I/\sigma(I)\rangle$ | 11.9 (1.6) |
| $CC_{1/2}$ | 0.998 (0.560) |
| $R_{merge.}$ | 0.071 (1.149) |
| $R_{p.i.m.}$ | 0.045 (0.726) |

**Table S2: FphE structure solution and refinement**

Values for the outer shell are given in parentheses.

| | Compound FphB-OX-5 bound |
|---|---|
| PDB ID | 9DRO |
| Resolution range (Å) | 39.82 – 1.54 (1.56 – 1.54) |
| Final R-work | 0.156 (0.266) |
| Final R-free | 0.183 (0.269) |
| Protein residues | 554 |
| Ligands | 2 |
| Water | 323 |
| Magnesium atoms | 4 |
| R.m.s. deviations | |
| Bonds (Å) | 0.010 |
| Angles (°) | 1.067 |
| Average *B* factors (Å$^2$) | 34.8 |
| Ligands | 55.5 |
| Water | 37.1 |
| Magnesium atoms | 30.1 |
| Ramachandran plot | |
| Most favored (%) | 98.4 |
| Outlier (%) | 0.4 |

```
FphE      ----------------------------------------------------METLE---  5
FphB      MRKKWSTLAFGFLVAAYAHIRIKEKRSVKSYMLEQGIRLSRAKRRFMYKEEAMKALEKMA  60
FphH      ----------------------MQIKLPKPFFFEEGKRAVLLLHGFTGNSSDVRQLGRFL  38
                                                              :. *

FphE      ------LQGAKLR-----------------------------------------------  12
FphB      PQTAGEYEGTNYQFKMPVKVDKHFGSTVYTVND-KQDKHQRVVLYAHGGAWFQDPLKIHF  119
FphH      ---------------------QKKGYTSYAPQYEGHAAPPDEILKSSPFVWFKDALDGYD  77


FphE      -------------------------------------YH---QVGQGPVLIFIPGANGTG  32
FphB      EFIDELAETLNAKVIMPVYPKIPHQDYQATYVLFEKLYHDLLNQVADSKQIVVMGDSAGG  179
FphH      -YL-------------------VE-----------------------QGYDEIVVAGLSLGG  96
                                                    .   *.: * .  *

FphE      DIFLPLAEQLKDHFTVVAVDRR-DYGESELTEPLPDSASN-----------PDSDYRVKR  80
FphB      QIALSFAQLLKEKHIVQP--------------GH--------------------------  199
FphH      DFALKLSLNRDVKGIVTMCAPMGGKTEGAIYEGFLEYARNFKKYEGKDQETIDNEMDHFK  156
          :: * ::   . :  *

FphE      DAQDIAELAKSL---------SDEPVYILGSSSGSIVAMHVLKDYPEVVKKIAFHEPPIN  131
FphB      ----I---------------VLISPVL-----DA-------------------TMQHPEIP  217
FphH      PTETLKELSEALDTIKEQVDEVLDPILVIQAEND-------------------NMIDPQSA  198
              :                  .*:      .                    : .*

FphE      TFLPDSTYWKDKNDDIVHQILTEGLEKGMKTFGETLNIAPIDAKMMSQPADTEEGRIEQY  191
FphB      DYLKKDP-----------------------------------------MVGVDGSV---  232
FphH      NYIYDHV-----------------------------------------DSD--------  208
           :: .

FphE      KRTMFWL-EFEIRQYTHSNITLDDFTKYSDKITLLNGTDSRGSFPQDVNFYI-NKETGIP  249
FphB      FLAEQWAGDTPLDNYKVSPINGD-L-DGLGRITLTVGTK-EVLYPDALNLSQLLSAKGIE  289
FphH      DKNIKWYSE-------------------SGHVITIDKEK-EQVFEDI-------------  235
               *  :                    .::      . .  : :

FphE      IVDIPGGHLGYIQKPEGFADVLLNMWG----------------   276
FphB      HDFIPG----Y------YQFHIYPVFPIPERRRFLYQVKNIIN   322
FphH      -----------------YQFLESLDWS--E-------------   246
                           :       :
```

**Figure S1:** Amino acid sequence alignment for FphB (UniProt number A0A0H2XJG5_STAA3), FphE (Y2518_STAA3), and FphH (A0A0H2XJL0_STAA3) from *S. aureus* strain USA300 as determined by the UniProt CLUSTAL O(1.2.4) multiple sequence alignment tool. Overall, FphB has 25.64% sequence identity with FphE and 22.65% identity with FphH. * indicates fully conserved residues, : indicates conservation of residues with strongly similar properties, and . indicates conservation of residues with weakly similar properties.

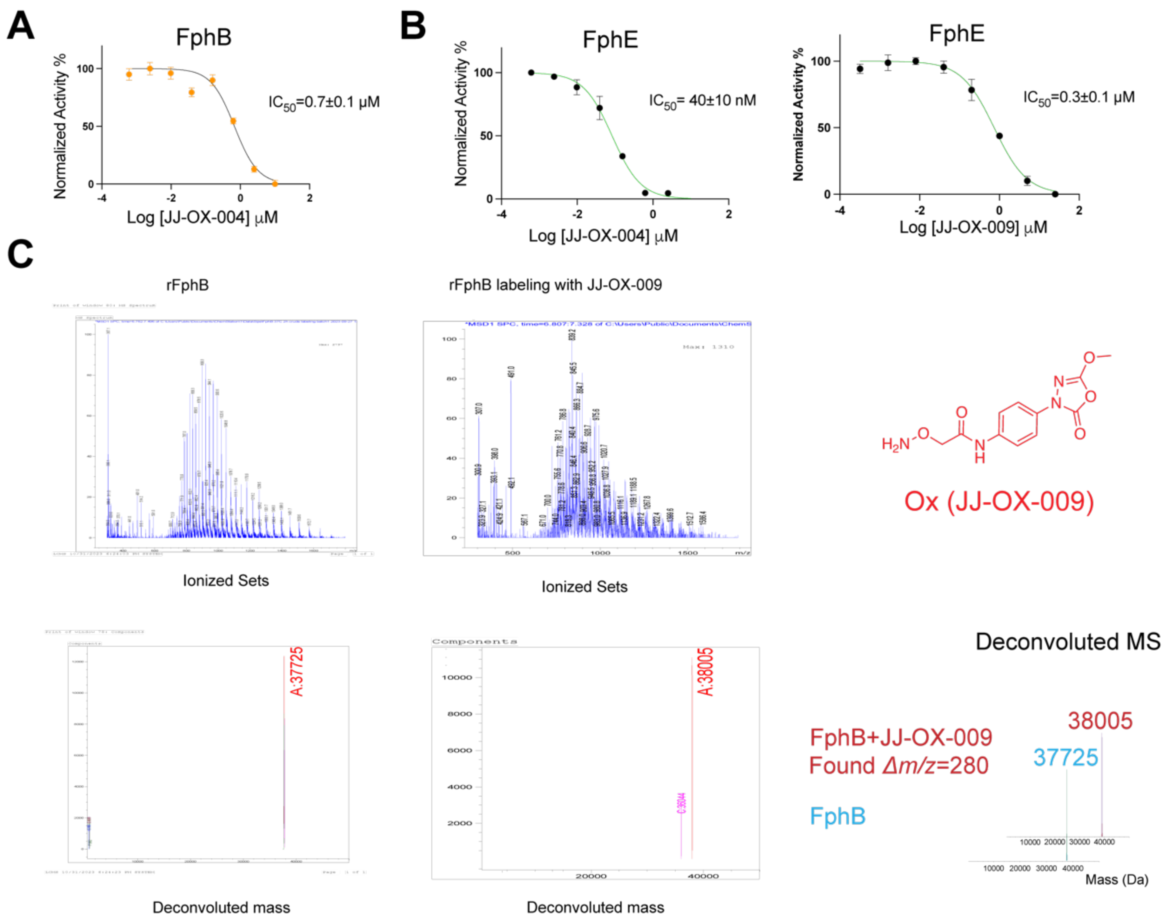


**Figure S2**: Enzymatic inhibition activities and covalent labeling of FphB and FphE by JJ-OX-004, JJ-OX-009. (A) Enzymatic assay of JJ-OX-004 for FphB. rFphB (100 nM) was incubated with inhibitor for 1 hour followed by measurment of the cleavage rate of the 4-MUB substrate (20 µM). The activity was normalized to a DMSO control, with points and bars representing means ± standard deviations (n=4). (B) Plot of inhibition of activity of FphE by JJ-OX-004 and JJ-OX-009 as performed in (A). The activity was normalized to a DMSO control, with points and bars representing means ± standard deviations (n=4). (C) Mass spectra of intact rFphB treated with DMSO (blue) or 100 µM JJ-OX-009 (red). The deconvoluted mass spectra of each is shown to indicate the mass shift resulting from covalent modification by the JJ-OX-009 probe.

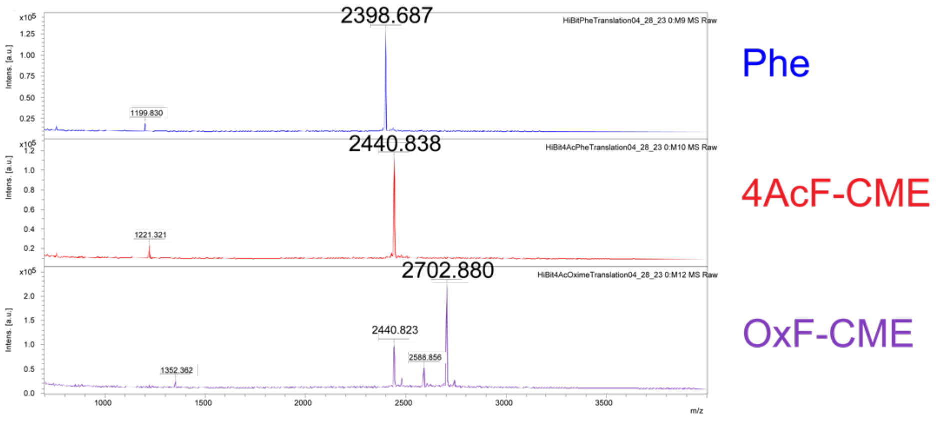


**Figure S3**: Confirmation of incorporation of non-canonical amino acids tested from the in NanoBiT assay using MALDI-MS. Mass spectra of *in vitro* translated HiBiT peptides containing tested amino acids (Phe: Phenylalanine (blue); 4AcF-CME: 4-acetyl-Phenylalanine-cyano methyl ester (red); OxF-CME: Oxadiazolone-Phenylalanine-cyano methyl ester (purple), after *in vitro* translation of the three HiBiT peptides, using previously reported methods.[1]

| Codon Box | TTT | TAT | TGT | CTT | ATT | GTT | CCT | GCT | TCT | ACT | AAT | AGT | GGT | CAT | CGT | GAT | INI | TGG | CAG |
|---|---|---|---|---|---|---|---|---|---|---|---|---|---|---|---|---|---|---|---|
| | Aromatic | | | Aliphatic | | | | | Polar | | | | | Basic | | Acidic | Cyclization | | covalent |
| 1 | F | Y | W | L | I | V | P | A | S | T | N | S | G | H | R | D | ClAc_F | C | OxF |
| 2 | f | Y | E | L | s | V | P | a | S | T | N | S | G | k | R | D | ClAc_f | C | OxF |
| 3 | CpG | Y | CpmG | L | Sar | V | P | MeA | S | T | N | S | G | Tic | R | D | ClAc_A | C | OxF |
| 4 | HxG | Y | E | L | Sar | V | P | Ahp | S | T | N | S | G | k | R | D | ClAc_A | C | OxF |

**Figure S4.** Designed codon tables for mRNA display libraries. INI: initiator. Lower-case single letters represent D-natural amino acids. f=D-Phe, s=D-Ser, a=D-Ala, k=D-Lys. All non-canonical amino acids are highlighted as red in tables.

| Hit Names | Hit Sequences | Ring Sizes |
|---|---|---|
| FphB-OX-1 | kATbYYYgNC | 9 |
| FphB-OX-2 | kAgNYYDLYYC | 10 |
| FphB-OX-3 | kAgVNYLYNYC | 10 |
| FphB-OX-4 | kAgNYYYLLLL | 10 |
| FphB-OX-5 | kFFDgIC | 6 |
| FphB-OX-6 | kFgIYDC | 6 |
| FphB-OX-7 | kFIFDgC | 6 |
| FphB-OX-8 | kFIFDgNC | 7 |
| FphB-OX-9 | kFgYGYC | 6 |
| FphB-OX-10 | kFYNIgNC | 7 |
| FphB-OX-11 | kFFDgIFC | 7 |
| FphB-OX-12 | kFFgNNFDIC | 9 |
| FphB-OX-13 | kFFDFINgNC | 9 |
| FphB-OX-14 | kfgYNYC | 6 |
| FphB-OX-15 | kfgYDYYC | 7 |
| FphB-OX-16 | kAYDgNC | 6 |
| FphB-OX-17 | kAgYYNC | 6 |
| FphB-OX-18 | kAgNDYDYC | 8 |
| FphB-OX-19 | kFgYFDC | 6 |
| FphB-OX-20 | kfgSaYYC | 7 |
| FphB-OX-21 | kfgsDDYC | 7 |

| | |
|---|---|
| Tic=b | f=D-Phe |
| OxF=g | a=D-Ala |
| ClAc=k | s=D-Ser |

**Figure S5**: Linear sequences of the twenty-one hits from mRNA display screening against FphB. All natural amino acids are annotated using upper case letters. All six non-natural amino acids are annotated using lower case letters. Tic=(*S*)-1,2,3,4-tetrahydroisoquinoline-3-carboxylic acid; OxF=Oxadiazolone phenylalanine; ClAc=Chloroacetic acid; f=D-Phenylalanine; a=D-Alanine; s=D-Serine

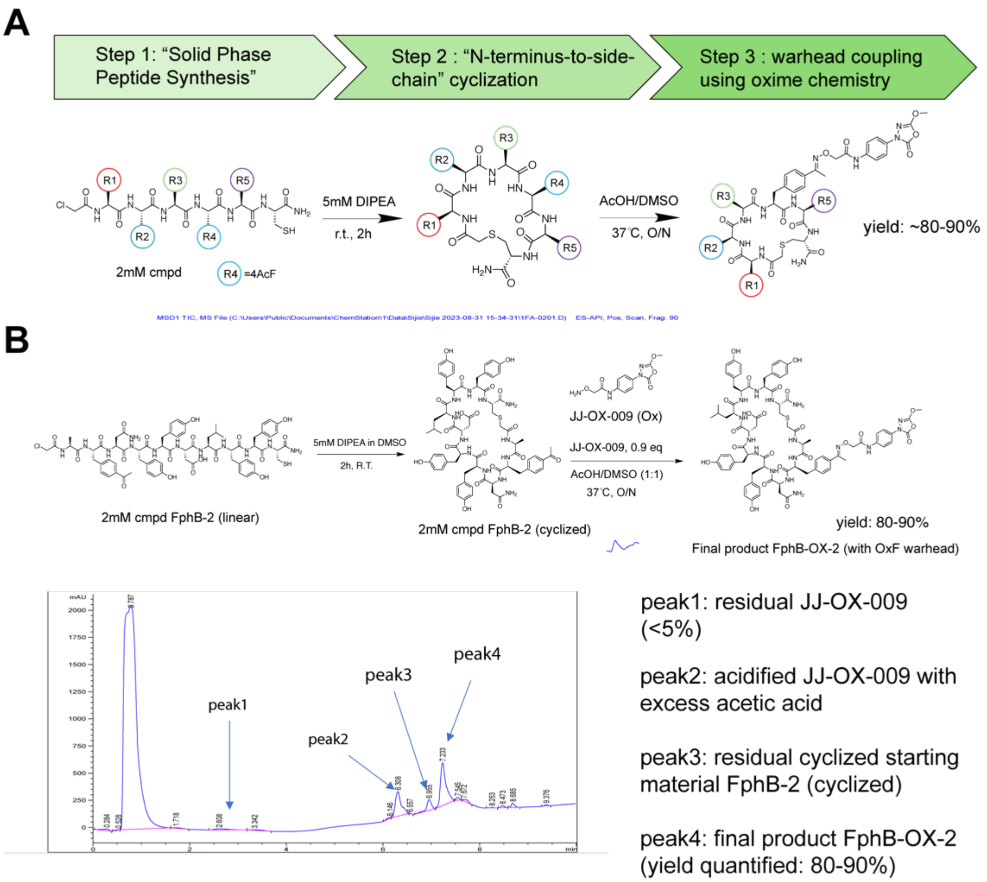


**Figure S6**: Synthesis of 21 compounds and final product yield determination using LCMS. (A) General synthesis scheme used to make the cyclic peptides containing the Ox amino acid. Linear peptides of 21 hits were synthesized using solid-phase peptide synthesis. Crude peptides were dissolved in DMSO at 2 mM concentrations and cyclized through N-terminal chloroacetic acetamide to C-terminal cysteine, using 5 mM DIPEA. The reactions were completed after 2 hours at r.t. In the last step, 0.9 eq of JJ-OX-009 was added with a co-solvent mixture of AcOH/DMSO (1:1 ratio) and incubated overnight at 37 °C to yield the final product in high yield (80-90% yield). (B) Validation of final product yield using a representative compound. Synthesis of compound FphB-OX-2 was accomplished following the chemistry route in (A), followed by the LCMS evaluations of final crude mixtures after reactions. Generally, for the 21 hits synthesized, 4 peaks can be detected and analyzed using LCMS. Peak1: residual unreacted JJ-OX-009; peak2: JJ-OX-009 acidified with excess acetic acid ([M+1] calculated: 323.0 g/mol, [M+1] found: 323.3 g/mol); peak3: residual starting material FphB-2 (cyclized form), ~10%; peak4: final product FphB-OX-2 (yield: 80-90%)

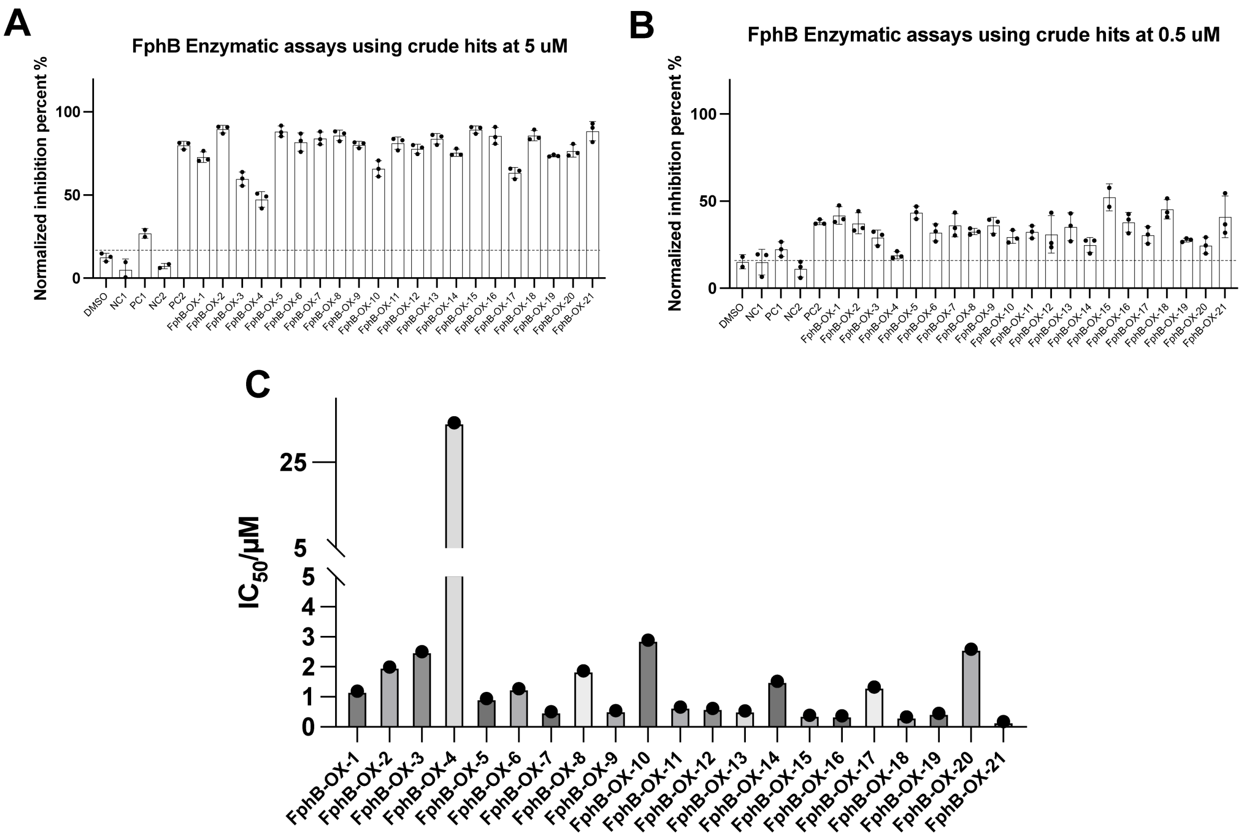


**Figure S7**. Direct enzymatic assay of 21 crude hits. The crude hits, with negative controls (NC1=5mM DIPEA in DMSO, NC2=5mM DIPEA in DMSO/AcOH (1:1) co-solvent) and positive controls (PC1=JJ-OX-009 with 5mM DIPEA in DMSO/AcOH (1:1) co-solvent, PC2=JJ-OX-004 with 5mM DIPEA in DMSO/AcOH (1:1) co-solvent), were flash frozen and lyophilized to complete dryness. After lyophilization, DMSO was used to resuspend the compound to 1mM stock (estimated based on the crude compound yield and purity). Crude compound stocks were tested at (A) 5uM and (B) 0.5uM final concentrations in enzymatic assay using rFphB and 4-MUB substrate (20 μM). Percent inhibition activity was calculated and normalized to a DMSO control, with points and bars representing mean ± standard deviation (n=3). (C) Full dose inhibition activity was quantified for all hits for rFphB. $IC_{50}$ quantifications of compounds in rFphB (100 nM) enzymatic assays, was measured by the cleavage rate of 4-MUB substrate (20 μM). $IC_{50}$ values were determined and plotted as mean value in the graph with points and bars representing mean ± standard deviation (n=3).

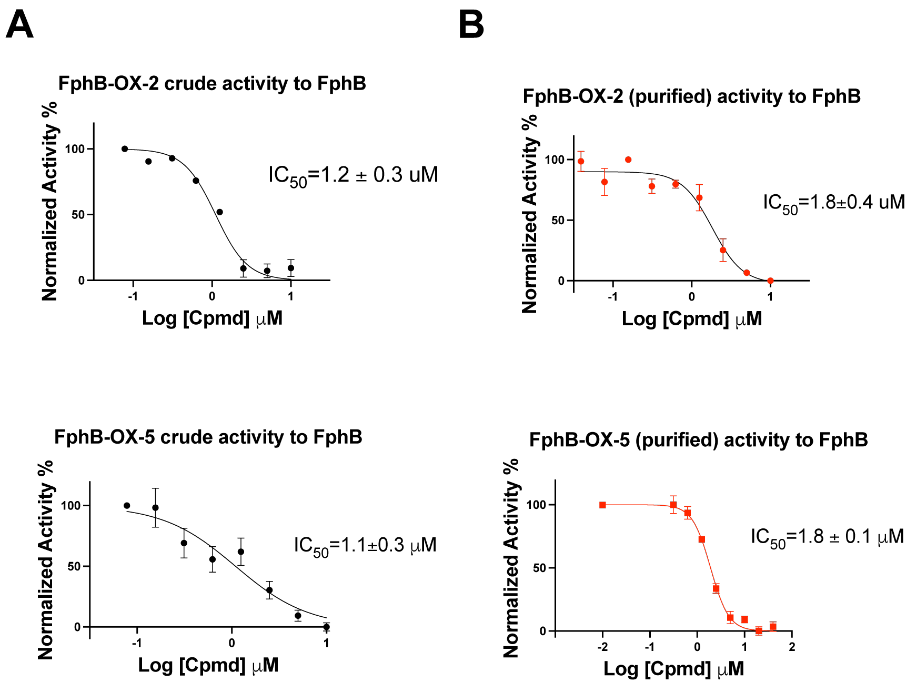


**Figure S8**: Direct comparison of activity of crude and purified compounds in enzymatic assays for rFphB. Plots of crude (A) and purified (B) FphB-OX-2 and FphB-OX-5 peptides in full-dose enzymatic assays against rFphB (100 nM) using the 4-MUB substrate (20 μM). Percent inhibition was calculated and normalized to a DMSO control, with points and bars representing mean ± standard deviation (n=3). $IC_{50}$ for each are shown at inset.

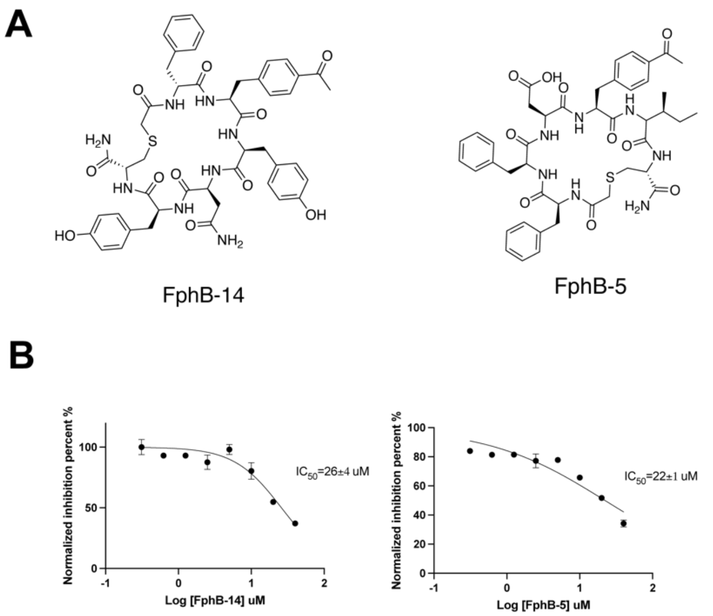


**Figure S9**: Enzymatic assays of non-warhead control molecules FphB-14 and FphB-5. (A) Structures of FphB-14 and FphB-5 (B) FphB-14 and FphB-5 peptides in full-dose enzymatic assays against rFphB (100 nM) using the 4-MUB substrate (20 µM). Percent inhibition was calculated and normalized to a DMSO control, with points and bars representing mean ± standard deviation (n=3). $IC_{50}$ for each are shown at inset.

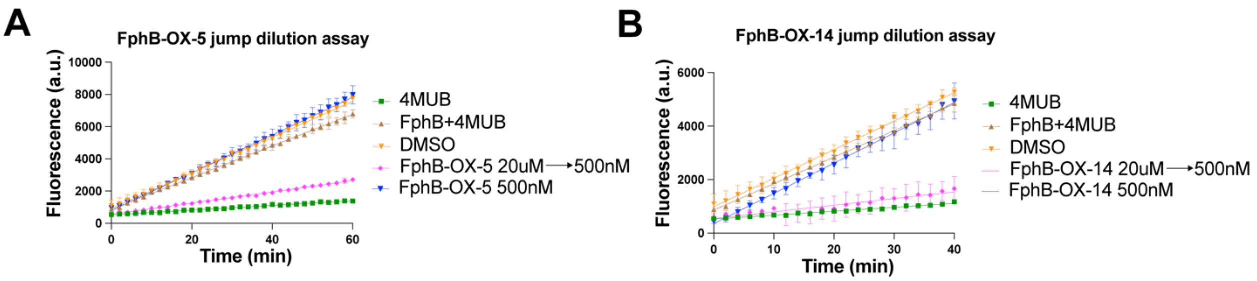


**Figure S10**: Jump dilution assay to determine reversibility of inhibition by Ox-containing cyclic peptides. (A) FphB-OX-5 and (B) FphB-OX-14 at 20 µM (10-fold of $IC_{50}$) was incubated with recombinant rFphB (2 µM) for 1 hr at r.t., followed by 40-fold dilution into a fluorogenic substrate (4-methylumbelliferone butyrate: 4MUB) solution. For the undiluted group (FphB-OX-5 500nM), 50 nM rFphB was incubated with 500 nM compound for 1hr at r.t and substrate added without dilution. Fluorescence progress curves are shown for substrate cleavage over time. Points and bars represent mean ± standard deviation (n=3).

| Compounds | $IC_{50}$ (μM) | | Normalized FphB/FphE index to JJ-OX-009 | Normalized FphB/FphE index to JJ-OX-004 |
|---|---|---|---|---|
| | FphB | FphE | | |
| JJ-OX-009 | 36 | 0.3 | 1 | 0.15 |
| JJ-OX-004 | 0.7 | 0.04 | 6.8 | 1 |
| FphB-OX-5 | 1.8 | 0.7 | 46.7 | 6.8 |
| FphB-OX-5 M1 | 1.1 | 0.9 | 98.2 | 14.3 |
| FphB-OX-5 M2 | 0.7 | 0.6 | 102.9 | 15 |
| FphB-OX-5 M3 | >10 | 1.4 | NA | NA |
| FphB-OX-5 M4 | 0.5 | 0.5 | 120 | 17.5 |
| FphB-OX-14 | 1.3 | 0.016 | 1.5 | 0.22 |
| FphB-OX-14 M1 | 0.09 | 0.022 | 29.3 | 4.3 |
| FphB-OX-14 M2 | 9.4 | 0.20 | 2.5 | 0.4 |
| FphB-OX-14 M3 | 0.5 | 0.014 | 3.36 | 0.5 |
| FphB-OX-14 M4 | 0.11 | 0.015 | 16.4 | 2.4 |

**Figure S11**: Normalized selectivity index for FphB over FphE of the alkyne mutational scanning of FphB-OX-14, FphB-OX-5 compared with JJ-OX-009 and JJ-OX-004. $IC_{50}$ for compounds in the table were determined using enzyme inhibition assays with rFphB (100 nM) and rFphE (1 nM) using the 4-MUB substrate (20 μM). FphB/FphE selectivity index was quantified using the FphE $IC_{50}$/FphB $IC_{50}$ values, and normalizing to the warhead only compound JJ-OX-009 and JJ-OX-004.

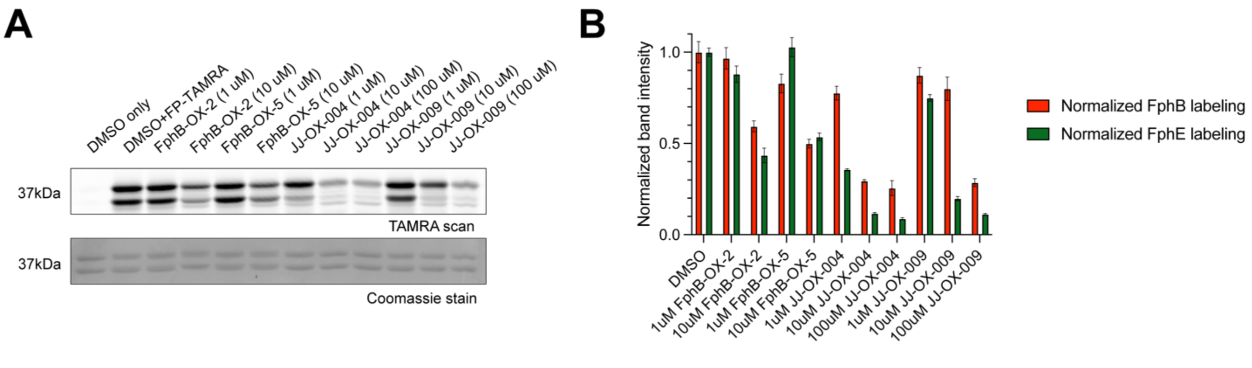


**Figure S12**: Dose-dependent Competition labeling of FphB-OX-5, FphB-OX-2 to rFphB/rFphE mixture. The compounds FphB-OX-2, FphB-OX-5, JJ-OX-004 and JJ-OX-009 were incubated with a protein mixture of rFphB and rFphE (mixed as 1:1 ratio at 1 uM for both proteins) at a range of final doses. Incubation of protein and compounds mixture was performed at room temperature for 1hr in PBST (0.02% triton in PBS), before treating with 2 uM FP-TAMRA probe for 30mins at room temperature. DMSO only: No FP-TAMRA, only DMSO treatment. DMSO+FP-TAMRA: DMSO 1hr treatment followed by 30 min FP-TAMRA treatment. All other samples were treated with compounds at different indicated concentrations for competition labeling, followed by a 30 min FP-TAMRA treatment. Protein competition labeling was measured by (A) SDS-PAGE/fluorescence scan of the gel and quantification of labeled proteins by ImageJ. Protein loading was assessed by coomassie staining. Intensity of bands in panel A were normalized to the DMSO+FP-TAMRA control (the second lane in the TAMRA scan of panel A).

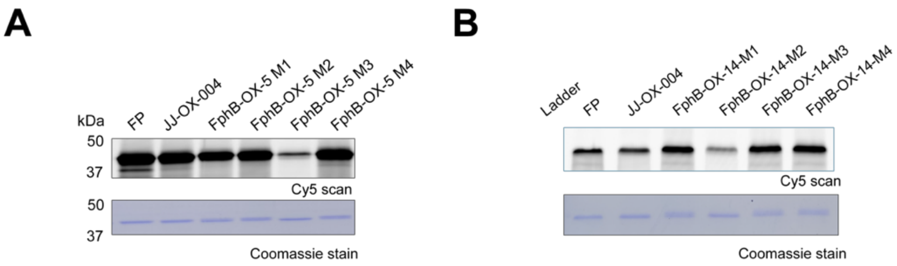


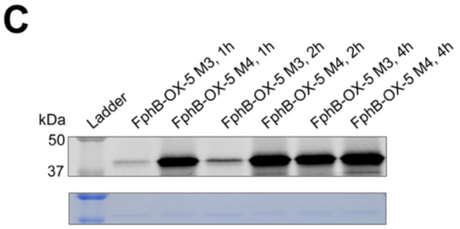


**D**

| Hits | 1h FphB labeling (Y/N, yield) | 4h FphB labeling (Y/N, yield) | 4h FphB S169A labeling (Y/N, yield) |
|---|---|---|---|
| FphB-OX-5 | Y, >80% | Y, >95% | N, 0% |
| FphB-OX-5 M1 | Y, >80% | Y, >95% | N, 0% |
| FphB-OX-5 M2 | Y, >80% | Y, >95% | N, 0% |
| FphB-OX-5 M3 | Y, 20% | Y, >95% | N, 0% |
| FphB-OX-5 M4 | Y, >80% | Y, >95% | N, 0% |

**Figure S13**: In-gel fluorescence labeling of rFphB by FP-alkyne, JJ-OX-004, FphB-OX-5 alkyne mutants and FphB-OX-14 alkyne mutants. (A) and (B) rFphB (1 uM ) was incubated with 1uM of indicated probes for 1h at r.t. in PBS. Labeled samples were clicked with Cy5-azide followed by SDS-PAGE and fluorescence Cy5 imaging scan. (C) Protein labeling of rFphB (1 uM) by FphB-OX-5 and FphB-OX-5 M3 (1uM) for indicated time points (1,2,4 hr) at r.t. in PBS. Labeled samples were clicked with Cy5-azide followed by SDS-PAGE and fluorescence Cy5 imaging scan. (D). Intact rFphB and rFphB (S169A) catalytic dead mutant labeling with FphB-OX-5 and its alkyne derivatives (20 uM) for 1hr or 4 hrs at r.t. followed by mass spectrometry to assess covalent labeling. Approximate labeling yield was quantified by using the total deconvoluted mass ion intensity of both labeled and unlabeled protein.

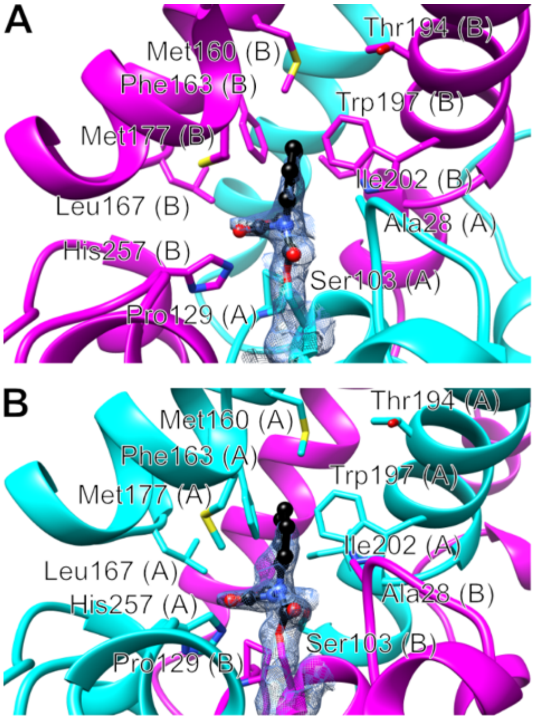


**Figure S14**: Close-up view of FphB-OX-5 covalently bound to FphE-Ser103. The $2F_o{-}F_c$ electron density map for the ligand and Ser103 is shown as a blue mesh at 1 σ. Ligand atoms as balls with carbon atoms in black. Bfactors increase and electron density diminishes with distance to Ser103, the remaining atoms of X beyond what is illustrated were therefore not modelled. Most likely the ligand will reach the surface via a tunnel passing Met160 and Thr194. (A) Covalent binding to FphE chain A (cyan) with active site environment mostly comprised of FphE homodimer copy chain B (magenta) residues. (B) Same as (A) in the corresponding other dimer active site.

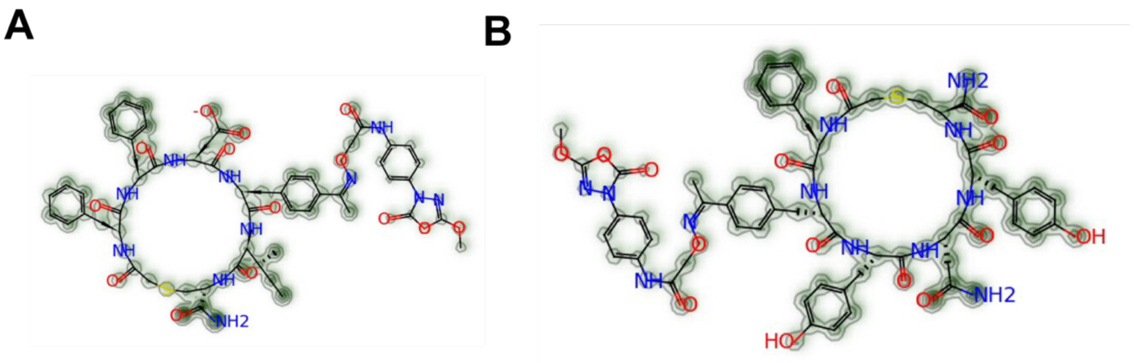

**Figure S15**. Covalent docking using the Alphafold predicted structure of FphB. Models of the FphB active site with (A) FphB-OX-5 per-atom RMSF from molecular dynamics. Contour lines depict increasing RMSF (B) FphB-OX-14 per-atom RMSF from molecular dynamics. Contour lines depict increasing RMSF. (C) Close-up view of opened form oxadiazolone electrophile covalently bound to FphB-Ser169 in the active site pocket.

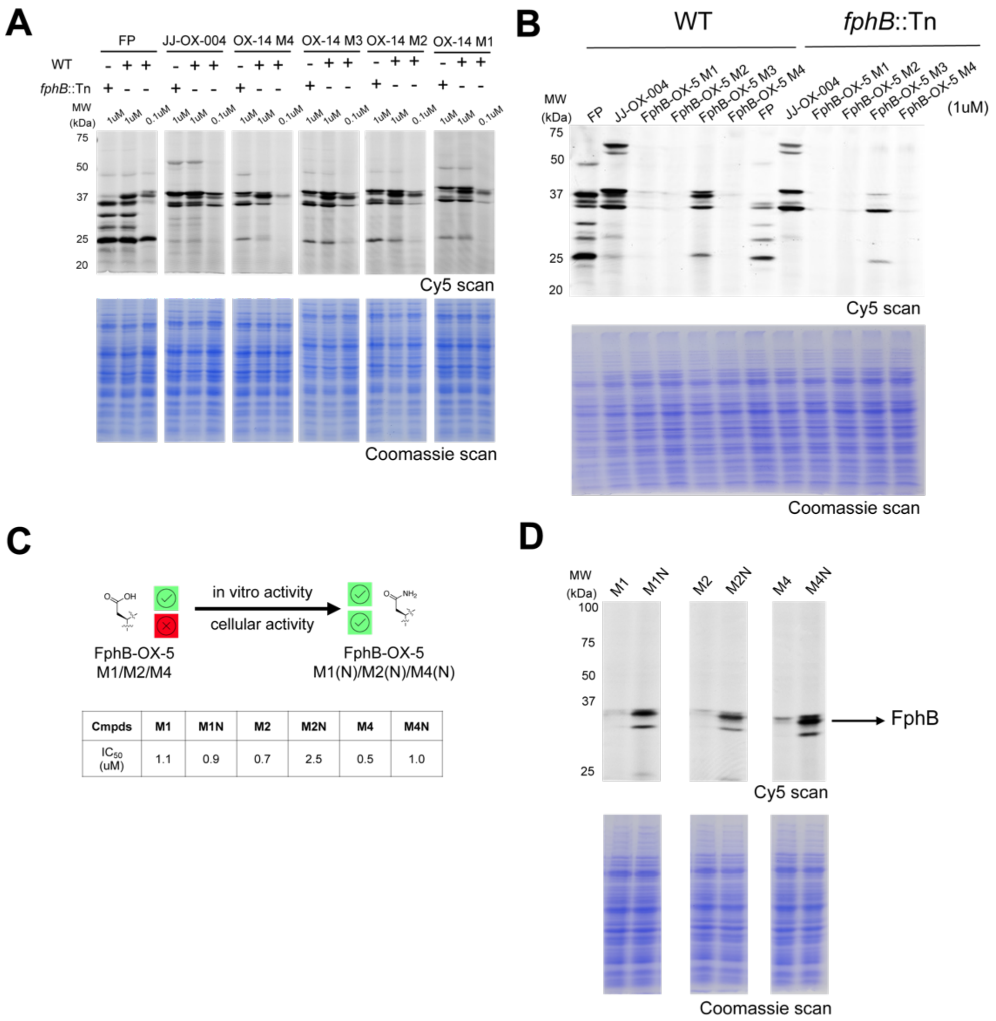


| Cmpds | M1 | M1N | M2 | M2N | M4 | M4N |
|---|---|---|---|---|---|---|
| $IC_{50}$ (uM) | 1.1 | 0.9 | 0.7 | 2.5 | 0.5 | 1.0 |

**Figure S16**: Importance of negative charge for binding of cyclic peptides to Fph targets. (A) SDS-PAGE analysis of live *S.aureus* labeled with FP-alkyne, JJ-OX-004 and FphB-OX-14 M4/M3/M2/M1 mutants. Wild-type (WT) and *fphB* transposon (Tn) mutant USA300 *S.aureus* strains were treated at the indicated concentrations of probes at 37 °C for 1 h. Cells were washed and lysed. The lysate was clicked with Cy5-azide using the CuAAC reaction. Total protein (7 µg) was loaded per lane for SDS-PAGE analysis with fluorescence imaging and coomassie staining. (B) Wild-type (WT) and *fphB* transposon (Tn) mutant USA300 *S.aureus* strains were treated at 1uM of the indicated OX-5 alkyne probes at 37 °C for 1 h and cells were lysed after labeling. Then lysate was clicked with Cy5-azide by the CuAAC reaction. Total protein (7 µg) was loaded per lane for SDS-PAGE analysis with fluorescence imaging and coomassie staining. (C) The aspartic acid residue in FphB-OX-5 M1/M2/M4 cyclic peptides was replaced with asparagine to generate FphB-OX-5 M1N/M2N/M4N. $IC_{50}$ quantifications of compounds were measured using rFphB (100 nM) in enzymatic assays using the 4-MUB substrate (20 µM). (D) Live cell labeling of the six OX-5 M1/M1N/M2/M2N/M4/M4N compounds using Wild-type (WT) USA300 *S.aureus* strain. Cells were treated with 1uM of the six probes at 37 °C for 1 h and lysed after labeling. Lysate was clicked with Cy5-azide by the CuAAC reaction. Total protein (7 µg) was loaded per lane for SDS-PAGE analysis with fluorescence imaging and coomassie staining.

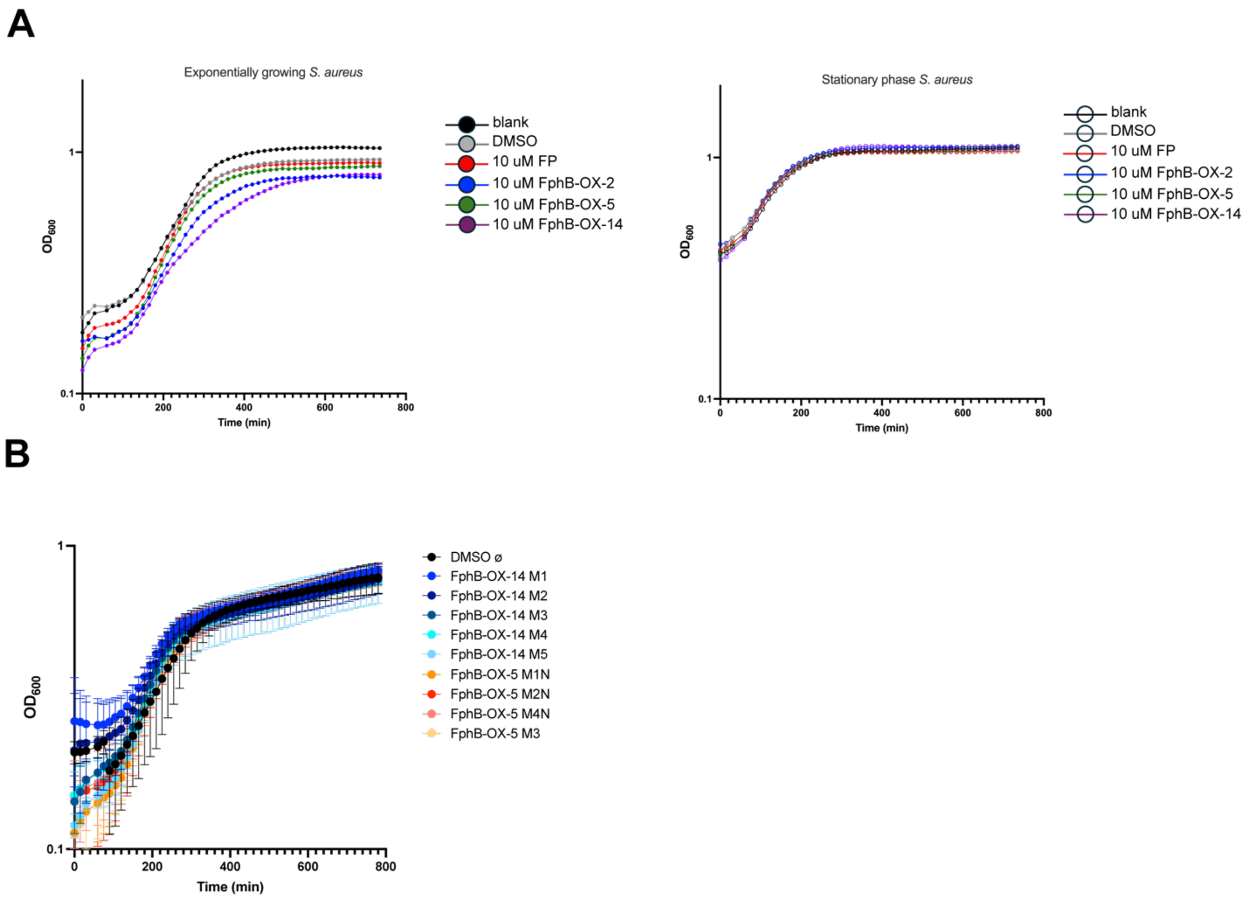


**Figure S17**. bacterial growth inhibition assay. The bacterial strain *Staphylococcus aureus* USA300 was cultured on tryptic soy agar (TSA) plates by overnight incubation at 37 °C. A single colony was picked and inoculated to tryptic soy broth. The primary culture was grown to exponential phase (OD600 = 0.2) at 37 °C and suspension was diluted 1:100. An inoculum of 100 µL was introduced to a treatment plate containing 20 uM (2% DMSO conc.) of compounds in TSB or Mueller-Hinton broth 2 (MHB2, cation adjusted; Millipore Sigma, Bedford, MA) media (100 µL of treatment/well) to give a final total volume of 200 µL/well with final 10 uM (1%DMSO) compound concentration and a final inoculum density of ~5 × $10^5$ CFU/mL. (A) For parental compounds, FphB-OX-14, FphB-OX-5, FphB-OX-2 and FP was tested. The completed assay plate was incubated at 37°C and the growth data were recorded for both exponential phase and stationary phase, with a Cytation 3 Multi-Mode Reader. (B) For all FphB-OX-14 and FphB-OX-5 derivatives, the completed assay plate was incubated at 37°C and the growth data was recorded for exponential phase, with a Cytation 3 Multi-Mode Reader.

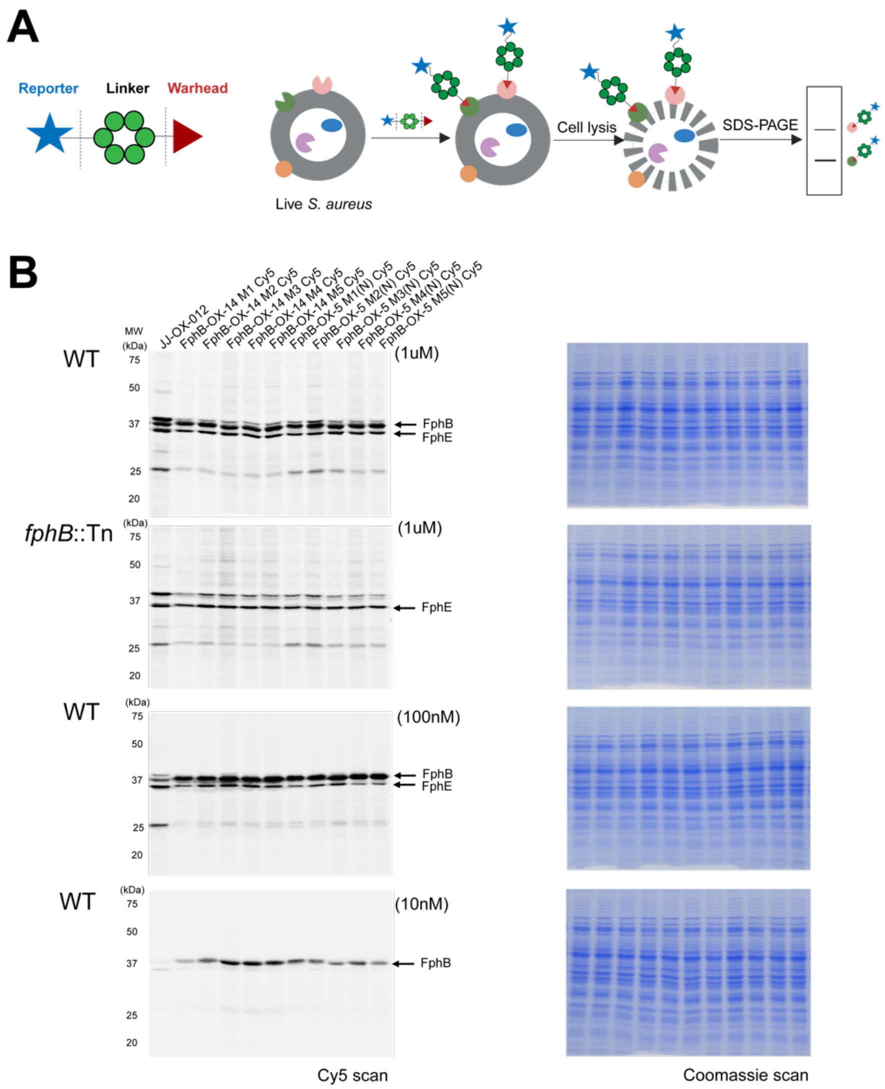


**Figure S18**. Live cell (both wild-type and *fphB* transposon (*fphB*::Tn) mutant strains) labeling using JJ-OX-012, FphB-OX-14 and FphB-OX-5 fluorescent probes. (A) Cartoon of imaging probes and live cell labeling workflow. (B) All fluorescent probes were synthesized by conjugating Cy5 by click reaction, purified and directly used for live cell labeling. Wild-type (WT) and *fphB* transposon (Tn) mutant USA300 *S. aureus* strains were treated at the three indicated concentrations of fluorescent probes (10 nM, 100 nM, 1000 nM) at 37 °C for 1 h before cell lysis. Total protein (7 µg) was loaded per lane for SDS-PAGE analysis with fluorescence imaging and coomassie staining.

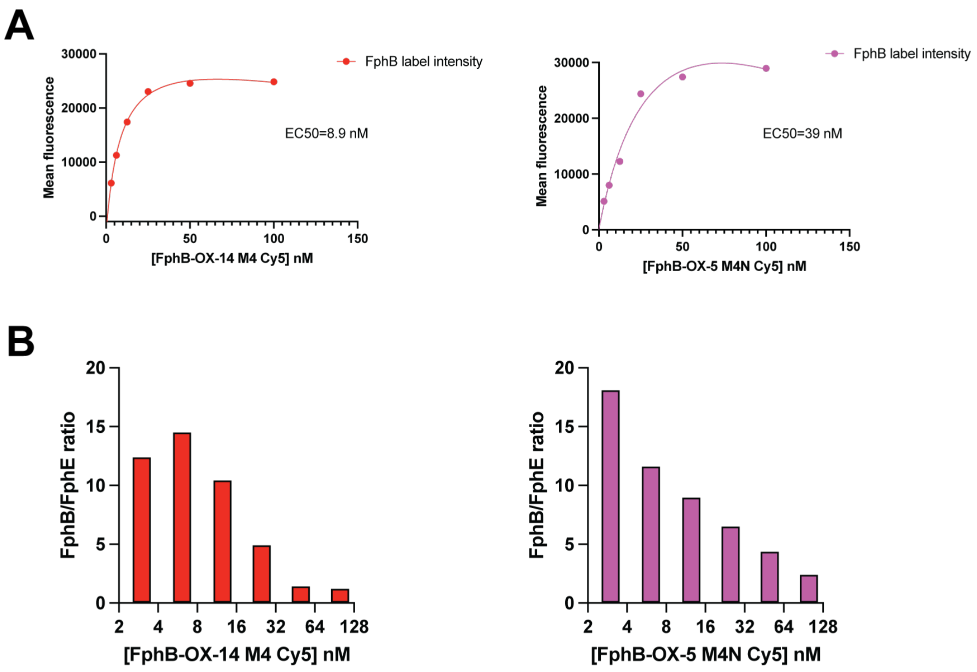


**Figure S19.** Quantifications of dose-dependent live cell labeling using the imaging probes. (A) Quantification of labeling intensity of FphB-OX-14 M4 Cy5, and FphB-OX-5 M4N Cy5 in live *S. aureus* in figure5B using ImageJ. Band intensity (mean fluorescence) of FphB of each compound at each concentration was plotted. $EC_{50}$ was quantitated by fitting the curve using total binding mode in Graphpad 10. (B) Quantification of FphB/FphE labeling ratios of two compounds above in figure5B. Images were scanned at the same time and all band intensity was measured using the same brightness/contrast settings.

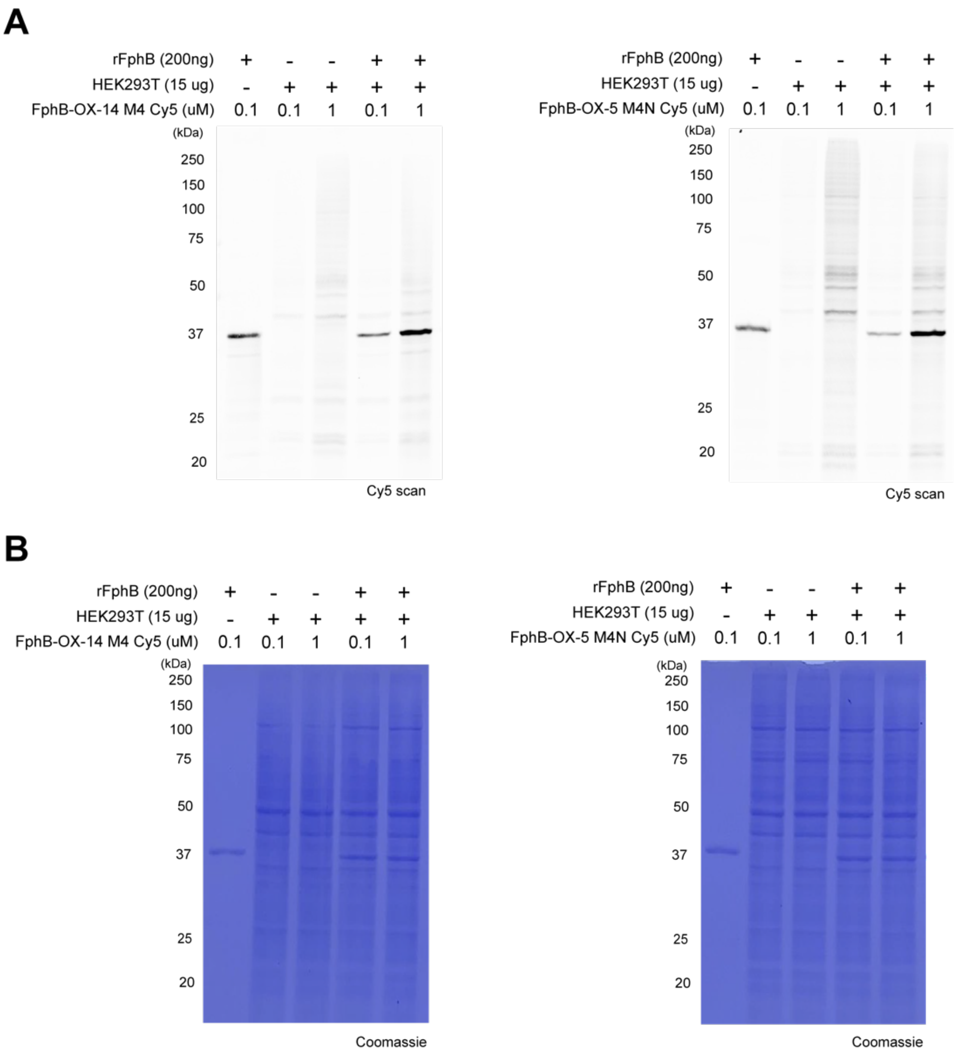


**Figure S20**. SDS-PAGE analysis of HEK293T lysate or intact cell labeling with FphB-OX-14 M4 Cy5 and FphB-OX-5 M4N Cy5. (A) Representative images of HEK293T lysate labeling with FphB-OX-14 M4 Cy5 (Left) and FphB-OX-5 M4N Cy5 (right). HEK293T cell lysate (WT) with or without recombinant FphB (rFphB, 200 ng) was treated with the indicated concentrations of probe at 37 °C for 1 h. Total protein (7 µg) was loaded per lane for SDS-PAGE analysis with fluorescence. (B) Coomassie staining of the gel in A.

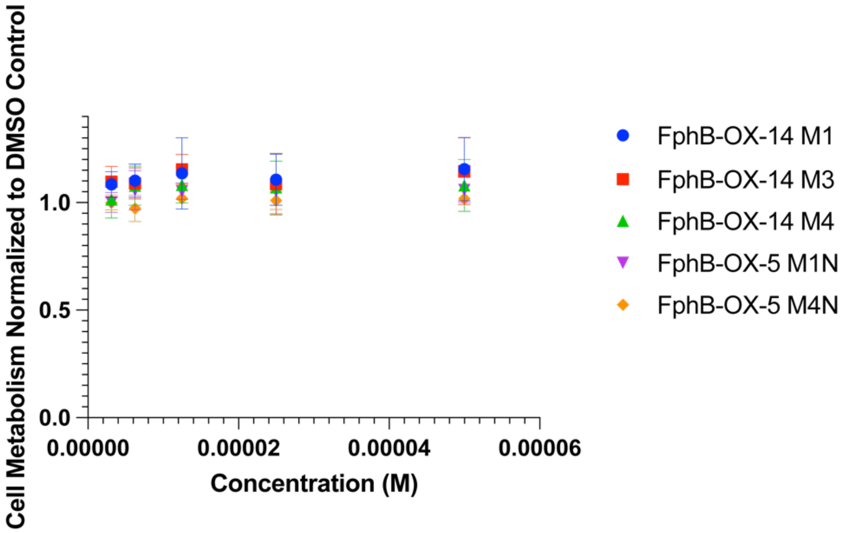


**Figure S21**. The effects of probes on cell viability were determined using a CellTiter-Glo ATP detection system (no. G7573, Promega). HEK293T cells were seeded in $0.1 \times 10^6$ cells/mL density in 96-well clear bottom white microplate (no. 655098, Greiner Bio-One). Cells were treated with compounds at 3.125, 6.25, 12.5, 25, 50 uM for 48 hrs. CellTiter-Glo reagent was added to cells, and incubated with gentle shaking for 15 min in dim light at RT. Luminescence was read on a GloMax microplate reader. Luminescence was normalized to DMSO-treated groups. The data was plotted in GraphPad Prism 10.

#### 2. Materials and Methods

##### FphB cloning, expression and purification

The FphB protein sequence (Uniprot ID: Q2FV90, strain NCTC8325, gene SAOUHSC_02844) was amplified from S. aureus USA300 genome using primers. DNA encoding the S. aureus FphB (27-322, wt or S176A mutant) fused to an N-terminal His6 tag and C-terminal AviTag was cloned into a pET vector and co-transformed into BL212(DE3) cells with a BirA-expressing plasmid for in situ biotinylation. Chemically competent BL21 (DE3) E. coli was transformed by the pET52b-(pETNKI-his-3C-LIC-kan) FphB or mutant FphB (S169A) expression vector, with the co-expression plasmid BirA (677838). An overnight culture of the transformed bacteria in LB (+ carbenicillin 100 μg/mL) selection medium was diluted 1:100 into 1 L of selection medium (ZYM5052) with 12.2 mg/L biotin at 37 °C for 2.5 hrs, and then 20 °C for 24 hrs with shaking (180 rpm). For purification, cells were harvested by centrifugation and bacterial pellets was stored at −80 °C. The following day, each pellet was thawed on ice, resuspended in 115 mL lysis buffer (20 mM Tris pH 8.0, 500 mM NaCl, 10% glycerol, 20 mM imidazole, 1 tablet EDTA free cocktail per 50 mL lysis buffer), and lysed by sonication (3 times, 1 min each: pulse ON 4 sec. pulse OFF 1 sec, 50% amplitude, Branson Sonifier; Branson, Danbury, CT). Lysates were centrifuged at 40,000 rpm in an Avanti JA-17 rotor (Beckman Coulter Life Sciences, Indianapolis, IN) for 1 hr, and filtered through 0.22 uM filter. The supernatant transferred to Ni-NTA resin pre-equilibrated with equilibration buffer (20 mM Tris pH 8.0, 300 mM NaCl, 20 mM imidazole, 10% glycerol) and incubated at 4 °C, rotating for 60 min, then purified by gravity flow. The resin was washed three times with washing buffer (50 mM Tris pH 8.0, 300 mM NaCl, 20 mM imidazole, 10% glycerol) and the His6-tagged protein was eluted in 50 mM Tris pH 8.0, 300 mM NaCl, 250 mM imidazole, 10% glycerol buffer. Eluates were analyzed by SDS-PAGE (Coomassie stain). To further purify the biotinylated proteins, elutes were transferred to streptavidin resin pre-equilibrated with equilibration buffer (20 mM Tris, pH 7.5, 10% glycerol, 300 mM NaCl), and washed three times with equilibration buffer. Biotinylated proteins were eluted in 20 mM Tris, pH 7.5, 10% glycerol, 300 mM NaCl, 50 mM biotin. Pools of fractions were run through SEC chromatography (buffer: 50 mM Tris, pH 7.5, 150 mM NaCl, 10% glycerol).

Protein purity was indicated by SDS-PAGE as ~90%, with 2.587 mg total protein yield, and M.W. was tested as 37742.65 Da, comparable as theoretical M.W.

Sequence of Avitag rFphB:

MAHHHHHHSAALSALQGPGSVKSYMLEQGIRLSRAKRRFMYKEEAMKALEKMAPQTA
GEYEGTNYQFKMPVKVDKHFGSTVYTVNDKQDKHQRVVLYAHGGAWFQDPLKIHFEF
IDELAETLNAKVIMPVYPKIPHQDYQATYVLFEKLYHDLLNQVADSKQIVVMGDSAGG
QIALSFAQLLKEKHIVQPGHIVLISPVLDATMQHPEIPDYLKKDPMVGVDGSVFLAEQW
AGDTPLDNYKVSPINGDLDGLGRITLTVGTKEVLYPDALNLSQLLSAKGIEHDFIPGYYQ
FHIYPVFPIPERRRFLYQVKNIINGGGLNDIFEAQKIEWHE

Sequence of Avitag rFphB (S169A):

MAHHHHHHSAALSALQGPGSVKSYMLEQGIRLSRAKRRFMYKEEAMKALEKMAPQTA
GEYEGTNYQFKMPVKVDKHFGSTVYTVNDKQDKHQRVVLYAHGGAWFQDPLKIHFEF
IDELAETLNAKVIMPVYPKIPHQDYQATYVLFEKLYHDLLNQVADSKQIVVMGDAAGG
QIALSFAQLLKEKHIVQPGHIVLISPVLDATMQHPEIPDYLKKDPMVGVDGSVFLAEQW
AGDTPLDNYKVSPINGDLDGLGRITLTVGTKEVLYPDALNLSQLLSAKGIEHDFIPGYYQ
FHIYPVFPIPERRRFLYQVKNIINGGGLNDIFEAQKIEWHE

**Luminescence-based assay evaluating in vitro translation efficiency of non-canonical amino acids**

Luminescence monitoring of *in vitro* translation of non-canonical amino acids using NanoBiT system was carried as previously reported.[1] *In vitro* translation reactions were carried out using in vitro translation mix, with the 0.04 μM DNA template or the 2 μM RNA template, 0.2 mM each of the amino acids necessary for translation, and 250 μM each aminoacyl-aa-tRNA, including tested non-natural amino acids, acylated-OxF-tRNA, acylated-4AcF-tRNA, and a positive benchmark control acylated-Phe-tRNA, and incubated at 37 °C for up to 2 h. For IVT quantification using luminescence as readout, the reaction was supplemented with LgBiT protein and the furimazine substrate (Promega). The translation reaction mixture was incubated at 37 °C for up to 2h, and luminescence was read via a plate reader.

**MALDI-TOF Mass Spectrometry of *in Vitro* Translated Peptides**

MAIDI validation of translated peptide in NanoBiT assay was performed as previously described.[1] After *in vitro* translation, the reaction volume (10−20 μL) was acidified to 0.1% trifluoroacetic acid and desalted on a $ZipTip_{C18}$ (Millipore) into 70% acetonitrile/water saturated with α-cyano-4-hydroxycinnamic acid as the MALDI carrier. MALDI-TOF MS analysis was performed using a RAPID-FLEX (Bruker) in reflector/positive mode.

**Synthesis of Aminoacyl-amino acids-tRNA**

As previously described,[1] 20 μL of aminoacylation reactions were carried out as follows: 20 μM tRNA and 20 μM flexizyme were combined in 0.1 M buffer, heated at 95 ˚C for 3 min and cooled to room temperature over 5 min. 600 mM (for eFx reactions) or 20 mM $MgCl_2$ (dFx) was added and the mixture was chilled on ice for 5 min. The reaction was initiated by addition of 25 mM of the substrate amino acid in DMSO and incubated on ice for the listed times. After acylation reaction, 0.3 M NaOAc (pH 5.2) and 100% EtOH were added and centrifuged for 15 min at 10,000g to pellet the aminoacyl-aa-tRNA. The pellet was rinsed with 70% EtOH aqueous solution containing 0.1 M NaOAc (pH 5.2) and then dried.

**Ribosomal synthesis of peptide macrocycle library**

Reagents and protocols for this *in vitro* translation system were performed as described.[1–3] The *in vitro* translation system of recombinant *E. coli* was reconstituted in 50 mM HEPES-KOH (pH 7.6), 10mM $Mg(OAc)_2$, 100 mM KOAc, 1 mM DTT, 2 mM spermidine, 20 mM creatine phosphate, 2mM ATP, 2 mM GTP, 1 mM CTP, 1 mM UTP, and 1.5 mg/mL *E. coli* total tRNA with final protein concentrations of 0.03 μM ArgRS, 0.38 μM AsnRS, 0.13 μM AspRS, 0.09 μM GlyRS, 0.02 μM LeuRS, 0.68 μM PheRS, 0.16 μM ProRS, 0.04 μM SerRS, 0.09 μM ThrRS, 0.02 μM TyrRS and 0.02 μM ValRS[MOU1] , 0.6 μM MTF, 2.7 μM IF1, 3 μM IF2, 1.5 μM IF3, 0.1 μM EF-G, 20 μM EF-Tu/Ts, 0.25 μM RF2, 0.17 μM RF3, 0.5 μM RRF, 1 μM T7 RNA polymerase, 4 μg/mL creatine kinase, 1.2 μM ribosome, 2 μM RNA template and 0.2 mM each of all necessary amino acids. For initiation of libraries, either N-chloroacetyl L-phenylalanine, N-chloroacetyl D-phenylalanine, or N-chloroacetyl L-alanine were acylated onto the initiator tRNA ($tRNA^{fMet}$). In all libraries, Cys was reprogrammed to the TGG codon via acylation onto $tRNA^{Asn}{}_{CCA}$ and OxF-CME was acylated onto $tRNA^{Asn}{}_{CUG}$.

10μM mRNA library was hybridized with a puromycin linker (11 μM) at RT for 3 min. The mRNA library was translated at 37°C for 30 min in the reprogrammed in vitro translation system to generate the peptide-mRNA fusion library.[4] Each reaction mixture contained 2 μM mRNA-peptide-linker conjugate and 25μM each tRNA aminoacylated with the specified noncanonical/canonical amino acids. After the translation, the reaction was quenched with 17mM EDTA. The product was subsequently reverse transcribed using RNase H minus reverse transcriptase (Promega) at 42°C for 30 min, and buffer was exchanged for phosphate buffered saline with 0.01% Tween-20 (PBST).

**Affinity selection of mRNA display libraries against FphB**

For affinity selection, the peptide-mRNA/cDNA solution was incubated with 250nM biotinylated FphB for 60 min at room temperature, and the Neutravidin coated beads (Cytiva) were further added and incubated for 20 min to isolate FphB binders. The beads were washed once with cold PBST buffer, the cDNA was eluted from the beads by heating for 5 min at 95°C, and fractional recovery from the affinity selection step was assessed by quantitative PCR using Sybr green I on a LightCycler thermal cycler (Roche). Beginning in round 2, we performed negative selections by pre-incubating the library with Neutravidin beads before incubation with FphB. From Round 3 onwards we added post-selection denaturing washes with 2x 5M guanidine-HCl. Finally, for the final rounds (5-6) we performed parallel selections on WT and S176A catalytic-dead FphB. Sequencing of the final enriched cDNA was carried out using a MiSeq next-generation sequencer (Illumina).

**Enzyme inhibition assay**

The enzyme activity of the purified enzymes was tested using 4-methylumbelliferyl butyrate (4-MUB) fluorogenic substrate. Briefly, 100 nM of WT rFphB were added 20 μM solution of each substrate mixed in 1X phosphate buffered saline (PBS; Corning, Manasas, VA) with 0.02% TritonX-100 (Fisher Scientific, Fairlawn, NJ). Fluorescence (λex = 365 nm and λem = 455 nm) was then read at 30 °C in 2-min intervals for 60 min using a Cytation 3 Multi-Mode Reader (BioTek, Winooski, VT). Turnover rates in the linear phase of the reaction (10–20 min) were calculated using GraphPad Prism9 as RFU/min. Rates were normalized by subtracting background hydrolysis rates measured for each substrate in reaction buffer in the absence of protein. The inhibitory activity of chemical probes was determined by incubating WT rFphB (100 nM) with a series of the inhibitor doses and quantifying residual activity using the 20 μM 4MUB. Inhibitors at various concentrations were preincubated with rFphB (1% final DMSO) at room temperature for 1 h, and rFphB activity was measured by monitoring the change in fluorescence intensity after adding 4-MUB for 1 h using a Cytation 3 Multi-Mode Reader (λex = 365 nm and λem = 455 nm). $IC_{50}$ Calculations were performed following dose response-inhibition mode using GraphPad prism 10 software.

For $IC_{50}$ shift assay, the inhibition of rFphB by FphB-OX-5 and FphB-OX-5 M3 (0, 0.15625, 0.3125, 0.625, 1.25, 2.5, 5, 10 and 20 μM) was investigated at three different incubation time points (0, 30 min, 1 hr, 2 hr, 3 hr, and 4 hr). rFphB activity was measured by monitoring the change in fluorescence intensity after adding 4-MUB. $IC_{50}$ values were estimated by dose-response equation as mentioned above. To determine $k_{inact}/K_I$, the reaction velocity for each well was determined by dividing the relative fluorescence units (RFU) by the reaction time. Inhibitor occupancy was calculated by fitting the reaction velocities into Equation 1 below.

$$Occupancy = \left(1 - \frac{v_x}{v_0}\right) * 100\ \%$$

Where $v_x$ is the reaction velocity in the presence of inhibitors, $v_0$ is the reaction velocity in the absence of inhibitors.

Then the inactivation rate constant was calculated by fitting the occupancy into equation 2 below,

$$Y = 100 * (1 - e^{-k_{obs} * t})$$

Where Y is the occupancy at different time, $k_{obs}$ is the observed inactivation rate constant, t is the incubation time of enzyme and inhibitors.

Then the $k_{obs}$ values acquired at different inhibitor concentrations were fitted into equation 3 below for $k_{inact}$ and $K_I$ values.

$$k_{obs} = k_{inact} * \frac{[I]}{[I] + K_I}$$

Where $k_{inact}$ is the concentration-independent inactivation rate-constants, and $K_I$ is the concentration of inhibitor required for half of the maximum potential rate of covalent bond formation.

**In-Gel fluorescence labeling of purified recombinant proteins**

Purified WT (1 μM) and S169A rFphB (1 μM) were incubated with the indicated concentrations of each chemical probe for 1 h at r.t.. For samples treated with alkyne probes, 20 μL of resulting mixtures were added 2.16 μL of freshly prepared click mix (0.5 μL of 50 mM CuSO4 in H2O, 1.16 μL of 100 mM BTTAA in DMSO, and 0.5 μL of 1 mM N3-TAMRA in DMSO) and 1.16 μL of 300 mM sodium ascorbate solution in $H_2O$. After incubation for 30 min at 37 ºC, 8 μL of 4X SDS loading buffer was added and samples were boiled at 100 ºC for 5 min. The samples treated with fluorescent probe was directly subjected by 4X SDS loading buffer without click reaction and boiled at the same condition with alkyne probes. Then, samples were analyzed by SDS-PAGE (12%) running at 120 V in an electrophoresis chamber under the ambient temperature. In-gel fluorescence was visualized using a GE Typhoon FLA 9000 (GE Healthcare, Pittsburgh, PA) followed by staining using Coomassie.

**Mass spectrometry validation of covalent FphB labeling with chemical probes**

1 μM WT rFphB was incubated with 20 μM of probes in 100 μL 1X PBS (pH=7.4) with 1% (v/v) DMSO, at room temperature for labeling. After 1 h, 10-20 μL of the labeled sample was directly used and analyzed on an Agilent 1200 HPLC equipped with an Agilent Zorbax SB-C3 column (1.8 μm, 2.1 x 150 mm, pore size 300 Å) coupled to an Agilent 6125B Single Quad Mass Spectrometer (Agilent Technologies, Santa Clara, CA). For LC-MS conditions, 95% solvent A (0.1% formic acid in LC-MS grade water), 5% solvent B (0.1% formic acid in LC-MS grade acetonitrile) at 0.6 mL/min was used for 2 min to remove excess of salts into waste before a linear 6-minute gradient from 80% solvent A (20% solvent B) to 20% solvent A (80% solvent B), followed by a one-minute 95% solvent A (5% solvent B). Protein labeling efficiency was quantified by the total intensity. The acquired mass of protein was deconvoluted using the deconvolution software in Agilent Bioanalysis Software package.

**Jump dilution assay**

The assay was performed by incubating 20 uM of compound FphB-OX-5 or FphB-OX-14 (DMSO as control) with 2 uM WT rFphB in assay buffer (0.02% TritonX-100 in 1X PBS) for 1 hr at ambient temperature. Pre-incubated sample was then diluted 40-fold (final concentration of 50 nM rFphB, 500 nM compound) into a solution containing 20 μM of 4-MU butyrate substrate. Enzyme activity was recorded by monitoring the change in fluorescence intensity for 1 h using a Cytation 3 Multi-Mode Reader (λex = 365 nm and λem = 455 nm). As control groups, compounds were also tested at 500 nM with 50 nM rFphB and 20 μM of 4-MU butyrate. The total volume/well, final concentrations of enzyme and substrates, and DMSO concentrations were all kept constant between the controls and jump dilution samples. Fluorescence intensities were plotted against time using GraphPad Prism 10.

**FphE X-ray Crystallography**

As previously described,[5] FphE was broad screened for crystallization using commercially available screens and hits were further optimized manually. For the unbound FphE structure 0.2 μL of 15 mg/mL FphE (9 mM HEPES pH 7.5, 87 mM NaCl, 13% DMSO) were mixed with 0.2 μL of reservoir solution. Sitting drop reservoir contained 25 μL of 0.18 M magnesium chloride, 0.1 M Tris pH 7.5, 22.5% w/v polyethylene glycol monomethyl ether 2000. The crystal was frozen in a solution of ~25% glycerol, 75% reservoir. For compound FphB-OX-5 bound, 10 μL of 15 mg/mL FphE (10 mM HEPES pH 7.5, 100 mM NaCl) were mixed with 4 μL compound FphB-OX-5 (10 mM in DMSO) and incubated at 18 °C overnight. 0.15 μL FphE compound FphB-OX-5 solution was mixed with 0.3 μL of reservoir solution. Sitting drop reservoir contained 25 μL of 0.18 M magnesium chloride, 0.1 M MES pH 8.5, 22.5% w/v polyethylene glycol monomethyl ether 2000. The crystal was frozen in a solution of ~25% ethylene glycerol, 75% reservoir. X-ray diffraction data were collected at the Australian synchrotron MX18 and MX29 beamline. Datasets were processed with XDS, merging and scaling were performed using AIMLESS. Phases were initially solved with Phenix Phaser molecular replacement using a model form Alphafold via Uniprot3 for FphE S. aureus strain USA300 AF-Q2FDS6-F1. Model building and refinement were conducted in COOT and Phenix. Structure figures, analysis and alignments were created with UCSF Chimera and LigPlot.

**FphE data collection, processing, refinement, deposition, and analysis**

X-ray diffraction data were collected at the Australian synchrotron and MX2 beamline. Datasets were processed with XDS, merging and scaling were performed using AIMLESS. Phases were initially solved with Phenix Phaser molecular replacement using the previously determined FphE structure PDB ID 8T87. Model building and refinement were conducted in COOT and Phenix.

The final structure was deposited to the worldwide protein databank PDB ID 9DRO. Statistics for the datasets are listed in Table SX and SY. Structure figures and analysis were created with UCSF Chimera.

**FphB structure modeling**

The starting structure for FphB was generated using Alphafold2 (1). The cosolvent simulation was prepared using CosolvKit (2) and run for 1 ms using the default protocol included in the package with 0.1M oxadiazolone as the cosolvent**.** The resulting trajectory was aligned with cpptraj (3) and the catalytic pocket was analyzed using MDpocket (4) to quantify pocket volume and mean localized hydrophobic density of the course of the trajectory. We then visually inspected individual frames from the trajectory which maximized those two values before picking one for further experiments.

**FphB-OX-5 and FphB-OX-14 Model Prediction using Covalent Docking**

Prior to covalent docking, FphB-OX-5 and FphB-OX-14 macrocycle backbone conformations were pre-sampled with molecular dynamics. Prior to simulation, the handles bearing the oxadiazolone warheads were truncated to phenylalanine residues. For FphB-OX-5, initial macrocycle conformations were generated with OpenFF Toolkit[6] and parameterized for molecular dynamics simulations with Espaloma 0.3.2.[7] The macrocycle was solvated in a TIP3P water box with 15 Å buffer on all sides. $Na^+$ and $Cl^-$ ions were added to neutralize charge and reach an ionic strength of 0.1 M. Simulations were performed with OpenMM v8.1.1[8] Following energy minimization, the system was gradually heated to 300 K and then equilibrated for 1 ns prior to the production run of 1  ms. The simulation was run with an NVT ensemble at 300 K and 1.0 ATM with periodic boundary conditions. A Langevin Middle integrator/thermostat was used with a friction coefficient of 1.0/ps and 2 fs timestep. The Particle Mesh Ewald method[9] was used for electrostatic interactions with a nonbonded cutoff of 1.0 nm and a tolerance of 0.0005. Frames from the simulation were written to a trajectory file every 10,000 time steps. The resulting trajectory was analyzed with MDAnalysis in Python.[10]

For FphB-OX-14, the macrocycle was simulated using CosolvKit for 1 ms (2 fs timestep) with 0.1M ethanol cosolvent in a 20Å radius TIP3P water box. The resulting trajectory was analyzed with MDAnalysis in Python to track Φ/Ψ torsions of each macrocycle residue. For FphB-OX-5, only the torsions for M1-M3 were found to exhibit distinct states; therefore, only these 6 torsions were used in further analysis. The distance from each point in Ramachandran space to each of three reference points (α-helix: -90°, -20°; β-sheet: -110°, 140°; Lα: 65°, 20°) was then calculated resulting in a dim vector for FphB-OX-5 and a dim vector for FphB-OX-14. A 6-component Gaussian Mixture Model (GMM) implemented with Scikit-learn was fitted on these distances for each macrocycle. These models were then used to group frames from the trajectories as having similar macrocycle backbone conformations. A frame was finally selected from each cluster to minimize the mean deviation from the median Ramachandran angles for M1-M3 for FphB-OX-5 or all Ramachandran angles for FphB-OX-14.

Six macrocycle conformations were selected for FphB-OX-5 and one for FphB-OX-14. We then used RDKit (https://www.rdkit.org) to the covalent handle.. We used Meeko (https://github.com/forlilab/Meeko) to prepare the ligands for covalent docking with Ser 176.[11] The macrocycle backbone was kept rigid for each selected conformation while sidechains were allowed to sample rotatable torsions freely. Reactive docking simulations were run with AutoDock-GPU[12] with 25,000,000 evaluations per LGA run. Output poses with the oxadiazolone warhead positioned within ~3-4Å of the OG of Ser 176 were selected for further simulation with molecular dynamics.

Molecular dynamics simulations were conducted using the OpenMM 8.1 software suite. The protein was parameterized with the Amber ff14SB force field, while the ligand parameters were obtained from Espaloma 0.3.2. The system was solvated in a dodecahedral box of TIP3P-FB waters, and sufficient NaCl was to reach 0.15 M salt concentration. Langevin dynamics simulations were performed with periodic boundary conditions and nonbonded interactions were calculated using the particle mesh Ewald (PME) method, with a cutoff of 10 Å and a smoothing function applied from 9 Å. Hydrogen bond constraints were enforced via the SHAKE algorithm, and hydrogen masses were adjusted to 3 u.m.a. Following energy minimization, systems were gradually heated to 300K in the NVT ensemble over 1000 ps with a 1 fs timestep, before switching to NPT conditions by adding a Monte Carlo barostat. The equilibration comprised 15 stages, summing up 2.5 ns, where harmonic restraints applied to all ligand and protein-heavy atoms were slowly released while increasing the timestep to 4 fs. For each system, production runs comprised three replicas of 100 ns.

**Bacterial culture and bacterial growth inhibition assay**

The bacterial strain *Staphylococcus aureus* USA300 was cultured on tryptic soy agar (TSA) plates by overnight incubation at 37 °C. A single colony was picked and inoculated to tryptic soy broth (TSB; Sigma-Aldrich, St. Louis, MO). The primary culture was grown to exponential phase (OD600 = 0.2) at 37 °C and suspension was diluted 1:100. An inoculum of 100 μL was introduced to a treatment plate containing two-fold serial dilutions of tested compounds in TSB or Mueller-Hinton broth 2 (MHB2, cation adjusted; Millipore Sigma, Bedford, MA) media (100 μL of treatment/well) to give a final total volume of 200 μL/well and a final inoculum density of ~5 × $10^5$ CFU/mL. The completed assay plate was incubated at 37°C and the growth data were recorded, with a Cytation 3 Multi-Mode Reader.

**Live *S. aureus* or lysate labeling**

Wild-type (WT) and *fphB* transposon mutant (*fphB*::Tn) USA300 *S. aureus* cells were cultured on TSA plates by overnight (~16 hr) incubation at 37 °C. Three colonies from each strain were picked and transferred to TSB media (three 50 mL falcon tubes, 20 mL each tube). The culture was grown to stationary phase shaking at 37 °C. Bacterial cultures were spun down at 3,000 g for 10 min and the supernatant was removed. The pellets were washed twice in 1X PBS and resuspended in the same buffer (5 mL). The resuspended bacteria was aliquoted into 400 μL for each sample.

Live cell labeling: Cells were preincubated with the indicated concentration of probes for 1 hr at 37 ºC. After washing twice with 1X PBS, cells were resuspended in 100 μL 1X PBS with 0.1% TritonX-100 and lysed by bead-beating at 4 ºC for 1 hr. For alkyne probes, after lysis, 20 μL of lysed cells in buffer were added 2.16 μL of freshly prepared click mix (0.5 μL of 50 mM CuSO4 in $H_2O$, 1.16 μL of 100 mM BTTAA in DMSO, and 0.5 μL of 1 mM $N_3$-Cy5 in DMSO) and 1.16 μL of 300 mM sodium ascorbate solution in $H_2O$. After incubation for 30 min at 37 ºC, 20 μL of 4X SDS loading buffer was added and samples were boiled at 100 ºC for 10 min. The prepared samples labeled by Cy5 probes were directly added by 4X SDS loading buffer without click reaction and boiled at 100 ºC for 10 min.

Lysate labeling: Cells were resuspended in 100 μL 1X PBS with 0.1% TritonX-100 and lysed by bead-beating at 4 ºC for 1 hr. After lysis, lysate was diluted using 1X PBS to 400 μL, and incubated with probes for 1 hr at 37 ºC. After labeling, lysate was either used for click reaction for alkyne probes or directly mixed with 4X SDS loading buffer and boiled at 100 ºC for 10 min for gel electrophoresis and imaging.

SDS-PAGE and imaging: The denatured samples were allowed to cool to ambient temperature and analyzed by SDS-PAGE (12%) running at 120 V in an electrophoresis chamber. Protein concentrations for each sample were quantified by using the BCA assay kit (Pierce/Thermo Scientific, Rockford, IL), and the equal amounts of total protein loaded were adjusted based BCA assay into each well. In-gel fluorescence (Cy5) was visualized using a GE Typhoon FLA 9000, followed by staining using Coomassie.

**Confocal microscopy**

*S. aureus* USA300 wild-type and *S. aureus* USA300 *fphB*::Tn strains containing pMC29 (which contains GFP under control of a constitutively-expressed promoter) were grown shaking at 37˚C in 10 mL TSB with 10 μg/ml chloramphenicol to maintain the plasmid. After overnight growth, samples were pelleted at 3,000 rpm for 10 minutes and washed twice with phosphate buffered saline (PBS) pH 7.4. Samples were resuspended in 2 mL of PBS. 100 nM of either JJ-OX-12, FphB-OX5-M4(N), or FphB-OX14-M4 was added to a 100 μL aliquot of each sample and incubated for 1 h at 37 ˚C. Samples were washed three times in PBS + 1% DMSO and subsequently resuspended in 50 μL PBS. Agarose pads were placed on glass slides and 2 μL of sample added on top of the pad immediately prior to imaging. Confocal microscopy was performed using a Zeiss LSM 780 microscope. GFP imaging was performed with excitation at 488 nm, power 20%, and gain 650. Cy5 imaging was performed with excitation at 633 nm, power 20%, and gain 650. Brightfield images were obtained simultaneously via the T-PMT channel. Post-imaging adjustments to brightness and contrast in the Cy5 channel were performed in ImageJ with the same adjustments applied equally to all images. Signal-to-noise analysis was performed using ImageJ. Individual bacteria were circled based on localization in the GFP channel and the mean fluorescent signal within this region of interest (ROI) was measured in the Cy5 channel. 150 ROI from three representative images per sample were measured. Similar measurements were obtained from regions not containing bacteria (background regions). The signal-to-noise ratio (SNR) was calculated as mean fluorescence from ROI divided by mean fluorescence from background regions.

**Quantitative analysis of microscopy images**

Method 1: Images were converted to 8-bit and analyzed in ImageJ. Individual cells were localized in the GFP channel, and a line was drawn, spanning the central axis of the cell. Fluorescence intensity along this line in the Cy5 channel was analyzed using the ImageJ "Plot Profile" tool. Measurements were taken for 20 representative cells per condition.

Method 2: A multi-step methodology including image pre-processing, segmentation for cell image analysis was employed. The initial step involves reading the image and converting it to grayscale, followed by the application of Otsu thresholding for segmentation and generating a binary image. Otsu's method, an automatic thresholding technique, selects an optimal threshold by maximizing the variance between two classes of pixels in a grayscale image. To enhance segmentation, morphological operations are performed, including the removal of small objects, closing gaps in cell boundaries, and filling holes within cells. Subsequently, the watershed transform is applied to the binary image to effectively separate each cell. The watershed transforms, a computer vision image segmentation technique, treats pixel intensities as elevations in a topographical landscape, allowing for the delineation of object boundaries based on the dynamics of water flowing into catchment basins. The resulting segmented cells are displayed by overlaying blue boundaries onto the original image. For the cell analysis, the major axis is calculated and plotted for each detected cell individually, visualizing it on the overlayed image.

**HEK293T cell labeling**

HEK293T cells (ATCC CRL-11268) were grown in Dulbecco's Modified Eagle Medium (DMEM; Gibco 11995065) supplemented with 10% fetal animal serum (FAS). Cells were grown in a humidified incubator at 37 ºC and 5% $CO_2$ atmosphere. For intact cell labeling, HEK293T cells were seeded at 40% confluency in six-well dishes. After 24 h, the cell growth medium was removed and replaced with fresh medium containing either DMSO vehicle or Cy5 probes (100 nM). Following incubation for 2 h, media was removed, and each well was rinsed once with 1X PBS. Cells were harvested by scraping into pre-cooled 1.5-mL Eppendorf tubes. Cells were pelleted by centrifugation (5 min, 2,000 g, 4 ºC). Pellets were washed two times with ice cold 1X PBS at which point cell pellets were resuspended in lysis buffer (1% SDS in 1X PBS) and lysed via tip sonication (3 times, pulse ON 5 sec and pulse OFF 5 sec). Lysate was cleared by centrifugation (10 min, 10,000 g). The supernatant was moved to a fresh tube and protein concentration was determined by BCA assay kit. Gel samples were prepared at a concentration of 2 mg/mL with appropriate volumes of 1% SDS buffer and 4X reducing loading buffer and boiled for 5 min at 95 ºC. For each sample, 3.5 µL (7 µg) were separated on a 12% SDS-PAGE gel. Fluorescence was visualized using a GE Typhoon FLA 9000 followed by staining using Coomassie. For lysate cell labeling, HEK293T cells grown in 10-mL dishes. The cell media was aspirated, and cells were washed with 1X PBS before being harvested by scraping into pre-cooled 15-mL Falcon tubes. Cells were pelleted by centrifugation (5 min, 2,000 g, 4 ºC) and washed two times with ice cold 1X PBS. Cells were then resuspended in ice cold PBS and lysed via tip sonication (3 times, pulse ON 5 sec and pulse OFF 5 sec). Lysate was centrifuged (10 min, 10,000 g) and the supernatant was moved to a fresh tube on ice. Protein concentration was determined by BCA assay. Labeling was conducted under the following three conditions: (1) HEK293T lysate (15 µg) with rFphB (200 ng), (2) HEK293T lysate (15 µg) alone, or (3) rFphB (200 ng) alone. For (1) and (2), 20 and 100 nM of Cy5 probes were added and proteins were incubated at 37 ºC for 2 hours. For (3), 100 nM of Cy5 probes was added and proteins were incubated at 37 ºC for 2 hours. After which, gel samples were prepared by adding 4X reducing loading buffer and boiling for 5 minutes at 95 ºC. Each sample was separated on a 12% SDS-PAGE gel. Fluorescence was visualized using a GE Typhoon FLA 9000 followed by staining using Coomassie.

**HEK293T cell toxicity assay**

The effects of probes on cell viability were determined using a CellTiter-Glo ATP detection system (no. G7573, Promega). HEK293T cells were seeded in 0.1 × 106 cells/mL density in 96-well clear bottom white microplate (no. 655098, Greiner Bio-One). Cells were treated with compounds at 3.125, 6.25, 12.5, 25, 50 uM for 48 hrs. CellTiter-Glo reagent was added to cells, and incubated with gentle shake for 15 min in dim light at RT. Luminescence was read on a GloMax microplate reader. Luminescence was normalized to DMSO-treated groups. The data was plotted in GraphPad Prism 10.

#### 3. Chemical Synthesis

##### General methods

Reagents and solvents were purchased from commercial suppliers and were used without further purification. All reactions were performed under argon atmosphere and stirring unless otherwise noted. Reaction flasks were dried overnight at 100 °C in an oven. Flash column chromatography was carried out using SiliaFlash® P60 (230–400 mesh, SiliCycle) with the indicated solvents of reagent-grade. Thin-layer chromatography (TLC) was performed using pre-coated 0.25 mm silica gel plates (Merck). $^{1}$H and $^{13}$C NMR spectra were recorded on BRUKER AVANCE III 500 spectrometer as solutions in the indicated solvents at room temperature (rt). Chemical shifts were quoted in parts per million (ppm, δ) downfield from tetramethylsilane (TMS) and were referenced to the deuterated solvent. $^{1}$H NMR data were reported in the order of chemical shift, multiplicity (s, singlet; brs, broad singlet; d, doublet; dd, doublet of doublets; t, triplet; td, triplet of doublets; q, quartet; qd, quartet of doublets; quint, quintet; m, multiplet and/or multiple resonance), number of protons, and coupling constant in hertz (Hz). High-resolution mass spectra were analyzed by LC-ESI/MS on a Waters Acquity UPLC system coupled to a Thermo Exploris 240 Orbitrap mass spectrometer. The following abbreviations for reagents and solvents are used: *N,N*-dimethylformamide (DMF), dimethyl sulfoxide (DMSO), *N*-methyl-2-pyrrolidone (NMP), petroleum ether (PE), tetrahydrofuran (THF).

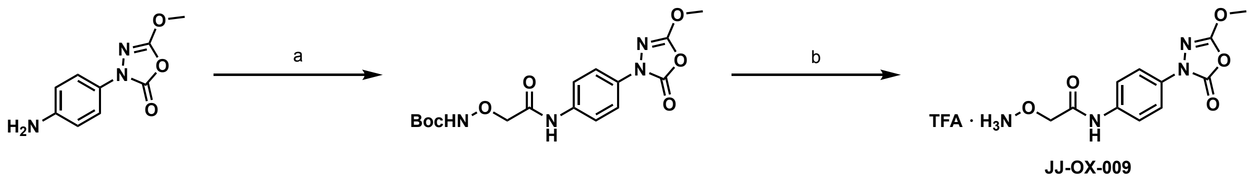


**Scheme S1**. Synthesis of JJ-OX-009. Reagent and conditions: (a) (Boc-aminooxy)acetic acid, EDC·HCl, HOBt, THF; (b) TFA, $CH_2Cl_2$, 56% for 2 steps.

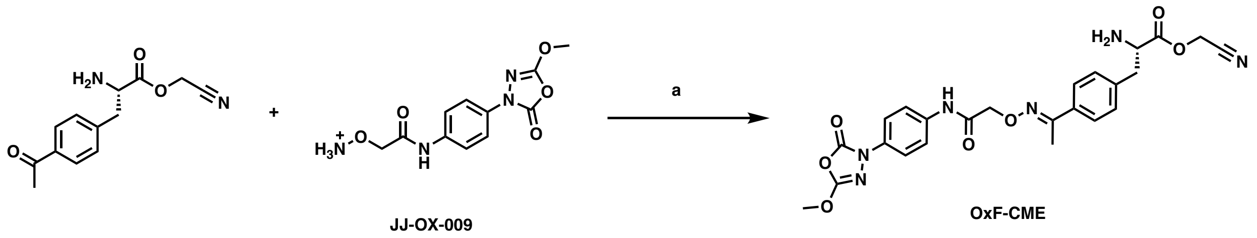


**Scheme S2**. Synthesis of OxF-CME. Reagent and conditions: (a) AcOH, DMSO;

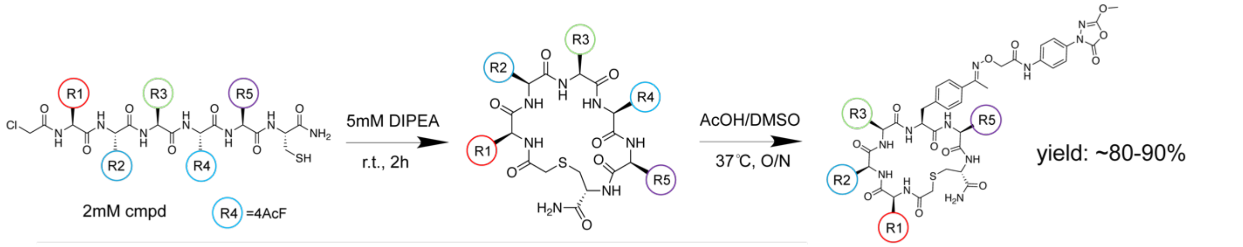


**Scheme S3**. Synthesis of macrocyclic hits from mRNA screening (general scheme. First step: cyclization; Second step: warhead coupling using oxime chemistry). Reagent and conditions: first step: DIPEA, r.t.; second step: AcOH, DMSO, 37 °C

**Synthesis of JJ-OX-009**

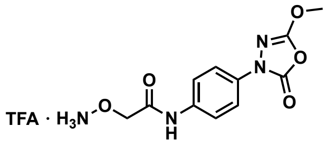


JJ-OX-009

To a stirred solution of 3-(4-aminophenyl)-5-methoxy-1,3,4-oxadiazol-2(3*H*)-one (100 mg, 0.483 mmol) in THF (5.07 mL) was added (Boc-aminooxy)acetic acid (90.4 mg, 0.473 mmol), EDC·HCl (97.2 mg, 0.507 mmol), and HOBt (52.3 mg, 0.507 mmol) at ambient temperature. After being stirred for 16 h at the same temperature, the reaction mixture was evaporated and diluted with $CH_2Cl_2$, and washed with brine. The organic layer was dried over $Na_2SO_4$ and concentrated in vacuo. The resulting residue was purified by flash column chromatography on silica gel (EtOAc/*n*-hexane = 1:1.5). Resulting intermediate was diluted in $CH_2Cl_2$ (2.0 mL) and added TFA (2.0 mL) at ambient temperature. After being stirred for 1 h, the reaction mixture was concentrated in vacuo. The resulting residue was purified by reverse-phase column chromatography C18 column (5–95% acetonitrile in water with 0.1% formic acid) to give JJ-OX-009 (104 mg, 56% for two steps) as a white solid: $^{1}H$ NMR (500 MHz, DMSO-$d_6$) δ 10.3 (s, 1H), 7.74 – 7.71 (m, 2H), 7.68 – 7.66 (m, 2H), 4.61 (s, 2H), 4.06 (s, 3H); $^{13}C$ NMR (126 MHz, DMSO-$d_6$) δ 166.1, 158.7, 158.5, 158.2, 155.4, 148.0, 135.6, 131.7, 120.1, 119.7, 118.5, 117.3, 115.0, 72.1, 58.1.

**Synthesis of OxF-CME**

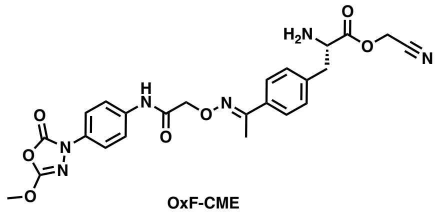

OxF-CME

Cyanomethyl (*S*,*E*)-2-amino-3-(4-(1-((2-((4-(5-methoxy-2-oxo-1,3,4-oxadiazol-3(2*H*) yl)phenyl)amino)-2-oxoethoxy)imino)ethyl)phenyl)propanoate

To a stirred solution of cyanomethyl (S)-3-(4-acetylphenyl)-2-aminopropanoate (984 mg, 4.0 mmol) in DMSO (10.0 mL), JJ-OX-009 (1124 mg, 4.0 mmol) and AcOH (1144 uL, 20.0 mmol) were added at ambient temperature. After being stirred for 4 h at 37 °C, the reaction mixture was HPLC purified to afford 1245 mg (~70% yield) of OxF-CME as a colorless oil: $^{1}$H NMR (500 MHz, DMSO-$d_6$) δ 10.06 (s, 1H), 8.42 (d, 2H), 7.77 – 7.70 (d, 2H), 7.70-7.60 (d, 4H), 7.34-7.28 (d, 2H), 5.15-5.09 (s, 2H), 4.78-4.72 (d, 2H), 4.07 (m, 1H), 3.16-3.10 (d, 2H), 2.55 (s, 6H); $^{13}$C NMR (126 MHz, DMSO-$d_6$) δ 168.1, 167.6, 155.6, 155.3, 155.4, 148.0, 136.0, 135.6, 134.8, 131.5, 126.4-126.30, 120.3-120.0, 115.2, 72.8, 52.8, 50.2, 35.5, 12.9

#### Synthesis of macrocyclic hits

*Solid-Phase Peptide Synthesis (SPPS)*

Peptides were prepared using traditional SPPS methods. All couplings and deprotections were monitored by ninhydrin tests. Briefly, peptides were synthesized on rink amide resin (30 mg, loading 100-200 mesh, 0.68 meq/g, 1% DVB) (Chem-Impex Int'L INC, Cat. # 02900) using standard Fmoc chemistry. A Syro II (Biotage) fully automated parallel peptide synthesizer with standard reactor block with 2 mL reaction vessel (PP-Reactor, 2 mL, with PE Frit, Cat. # V020PE051) (2mL plunger, Cat. # V020ST020) were used for synthesis in addition to manual synthesis for larger purified peptide stocks. General procedures for linear peptide synthesis follow the general procedure A (Fmoc deprotection), general procedure B (amide coupling), and general procedure C (washing steps). For the last step chloroacetic acid coupling, 5eq chloroacetic acid, 5 eq HBTU, and 5eq collidine were pre-mixed for activation before manually loading onto the peptide resin for reactions. The procedure was repeated twice. After the last step coupling, 6 times washing using DMF was performed before cleavage. Peptides were cleaved from resin using a mixture of 95% trifluoroacetic acid (Chem-Impex Int'L INC, Cat. # 00289), 2.5 % triisopropylsilane (Sigma Aldrich, Cat # 233781), 2.5% MilliQ water for 2 hours at RT, with occasional shaking. The cleavage mixture was drained and collected. The resin was then washed with additional cleavage mixture, drained, and collected. TFA was concentrated through evaporation with air stream in a ventilated hood. The residual cleavage mixture was precipitated in diethyl ether and allowed to cool at -20°C for 2 hours. The ether was then removed, and this process was completed three times. After the third ether wash, the residual ether was allowed to evaporate, and the compounds were dissolved in DMSO into 2mM stocks for next step cyclization.

*General Procedure A*: Fmoc Deprotection. Peptides were deprotected using a 20% piperidine (TCI, Cat # 203-642-1) in DMF solution. Peptides were deprotected for 15 min at RT two times.

*General Procedure B*: Amide bond coupling for amino acids. The coupling reagent (2-(1Hbenzotriazol-1-yl)-1,1,3,3-tetramethyluronium hexafluorophosphate (HBTU) (Peptides International, Cat # KHB-1065-PI) was pre-dissolved in DMF. The coupling reagent 2,4,6-collidine (Alfa Aesar, Cat # A11058) was pre-dissolved in DMF. All coupling reagent amino acids were predissolved in DMF. 5 equivalents of amino acid and 5 equivalents HBTU, and 10.0 equivalents of 2,4,6-collidine were preactivated and added to the reaction vessel using minimal DMF. The coupling reaction was allowed to react for 2x30min at RT.

*General Procedure C*: Washing Step. The reaction vessel was drained followed by addition of DMF and allowed to sit at RT for 1 min, this process was repeated two more times. The reaction vessel was drained followed by addition of DCM and allowed to sit at RT for 1 min, this process was repeated two more times. The reaction vessel was drained followed by addition of DMF and allowed to sit at RT for 1 min, this process was repeated two more times. Finally, the reaction vessel was drained.

*Peptide Cyclization*

Cyclization of peptides was achieved by adding 5mM DIPEA into 2mM peptide stocks above, and reactions were performed for 2 hours at r.t. with shaking. Cyclization of compounds was monitored and cyclization product yield was quantified by the LCMS.

*Warhead Coupling Using Oxime Chemistry*

Warhead JJ-OX-009 was coupled onto cyclized peptides through oxime chemistry. After cyclization, cyclized peptides in DMSO were added 1eq volume of acetic acid, and 0.9 eq of warhead JJ-OX-009. Oxime reaction was performed at 37°C for overnight with constant shaking at 700 rpm. Reaction progress and yield was monitored by LCMS, and purified using HPLC after overnight reactions.

**Synthesis of fluorescent imaging probes**

After macrocycle hits synthesized, 1 eq. (10 umols) of purified macrocycle peptide with alkyne handle was incubated with 1.5 eq (15 umols) of Cy5 azide (Broadpharm, catlog# BP-23908) and click cocktail (0.1 eq $CuSO_4$, 0.5 eq BTTAA, 20 eq. sodium ascorbate), 37°C, 700 rpm. Reaction was carried out in dark for 8 hrs and yield was monitored by LCMS. Reaction crude was purified by HPLC, dried and redissolved in DMSO to obtain the final fluorescent compound stocks (Cy5 conjugated imaging probes).

#### 4. LCMS of OxF-CME, JJ-OX-009 and FphB chemical probes

Compound Summary Table

| Compound name | Calculated MW $[M+1]^+$ | Found MW $[M+1]^+$ |
|---|---|---|
| JJ-OX-009 | 281.09 | 281.0 |
| OxF-CME | 509.17 | 509.0 |
| JJ-OX-12 | 936.55 | 936.2 |
| FphB-OX-5 | 1134.43 | 1133.9 |
| FphB-OX-5 M1 | 1082.40 | 1081.9 |
| FphB-OX-5 M2 | 1082.40 | 1082.0 |
| FphB-OX-5 M3 | 1114.44 | 1114.0 |
| FphB-OX-5 M4 | 1115.38 | 1115.9 |
| FphB-OX-5 M1N | 1081.40 | 1080.9 |
| FphB-OX-5 M2N | 1081.40 | 1080.9 |
| FphB-OX-5 M4N | 1115.38 | 1114.9 |
| FphB-OX-5 M5N | 1228.46 | 1227.9 |
| FphB-OX-5 M3 A1 | 1156.46 | 1155.9 |
| FphB-OX-5 M3 A2 | 1147.5 | 1147.0 |
| FphB-OX-5 M3 A3 | 1148.44 | 1147.9 |
| FphB-OX-5 M3 A4 | 1133.44 | 1132.9 |
| FphB-OX-5 M3 A5 | 1076.42 | 1075.9 |
| FphB-OX-5 M3 A6 | 1090.44 | 1090.0 |
| FphB-OX-5 M3 A7 | 1205.48 | 1204.9 |
| FphB-OX-5 M3 A8 | 1147.46 | 1146.9 |
| FphB-OX-5 M3 A9 | 1182.46 | 1181.9 |
| FphB-OX-5 M3 A10 | 1120.45 | 1119.9 |
| FphB-OX-5 M1N Cy5 | 1645.78 | 1645.6/823.3 |
| FphB-OX-5 M2N Cy5 | 1645.78 | 1645.6/823.3 |
| FphB-OX-5 M3 Cy5 | 1678.80 | 1678.6/839.8 |

| | | |
|---|---|---|
| FphB-OX-5 M4N Cy5 | 1679.76 | 1681.0/840.3 |
| FphB-OX-5 M5N Cy5 | 1792.85 | 1792.6/896.8 |
| FphB-OX-14 | 1199.42 | 1198.9 |
| FphB-OX-14 M1 | 1147.39 | 1146.9 |
| FphB-OX-14 M2 | 1131.39 | 1131.1 |
| FphB-OX-14 M3 | 1180.41 | 1179.8 |
| FphB-OX-14 M4 | 1131.39 | 1130.9 |
| FphB-OX-14 M5 | 1294.45 | 1293.8 |
| FphB-OX-14 M1 Cy5 | 1711.75 | 1711.6/856.3 |
| FphB-OX-14 M2 Cy5 | 1695.76 | 1695.6/848.3 |
| FphB-OX-14 M3 Cy5 | 1744.78 | 1744.6/872.8 |
| FphB-OX-14 M4 Cy5 | 1695.76 | 1695.6/848.3 |
| FphB-OX-14 M5 Cy5 | 1858.82 | 1858.6/929.8 |

LCMS traces

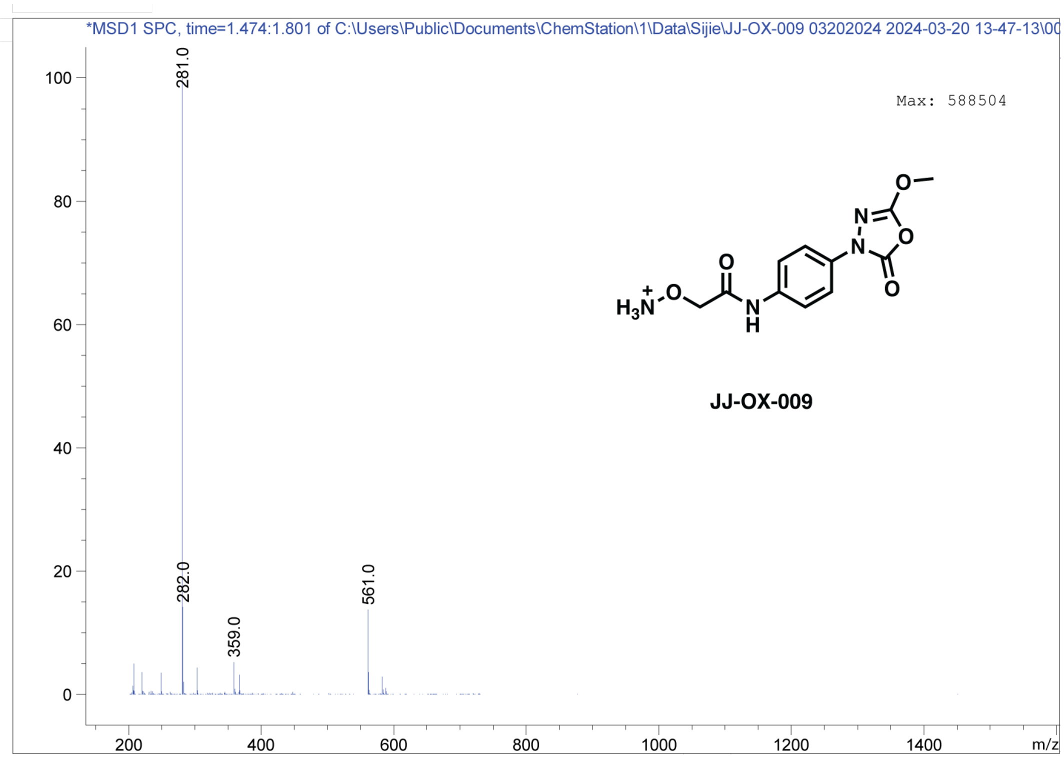
*MSD1 SPC, time=1.474:1.801 of C:\Users\Public\Documents\ChemStation\1\Data\Sijie\JJ-OX-009 03202024 2024-03-20 13-47-13\00
Max: 588504
281.0
282.0
359.0
561.0
JJ-OX-009
m/z

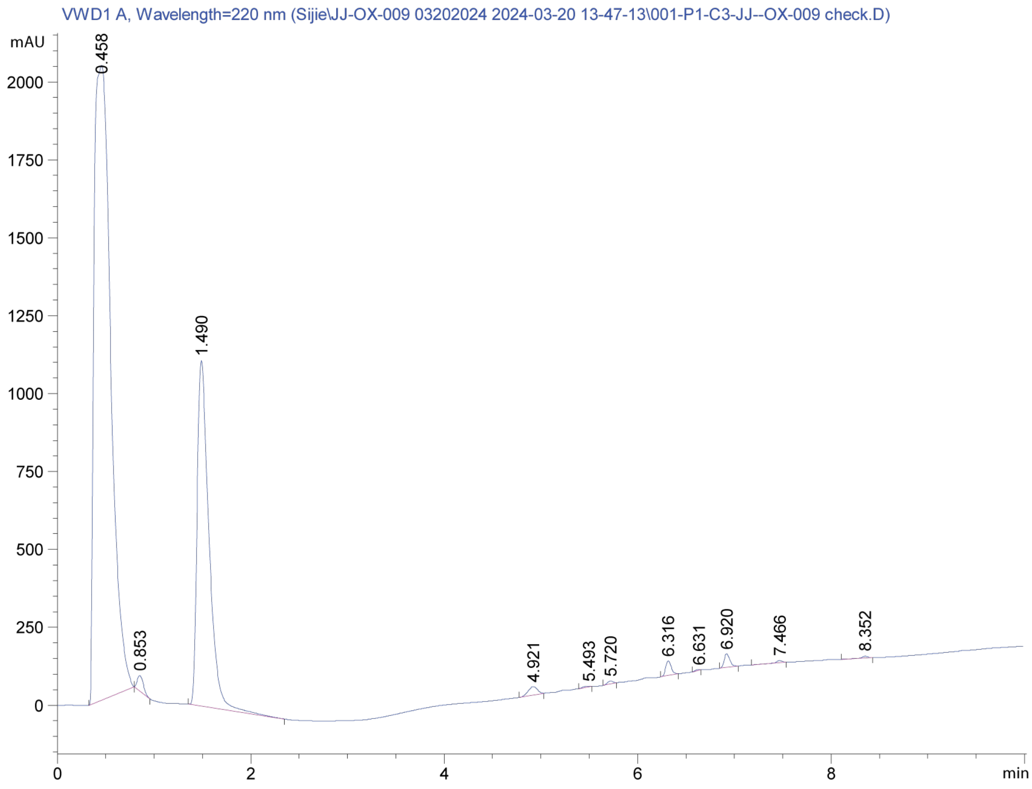
VWD1 A, Wavelength=220 nm (Sijie\JJ-OX-009 03202024 2024-03-20 13-47-13\001-P1-C3-JJ--OX-009 check.D)
mAU
0.458
0.853
1.490
4.921
5.493
5.720
6.316
6.631
6.920
7.466
8.352
min

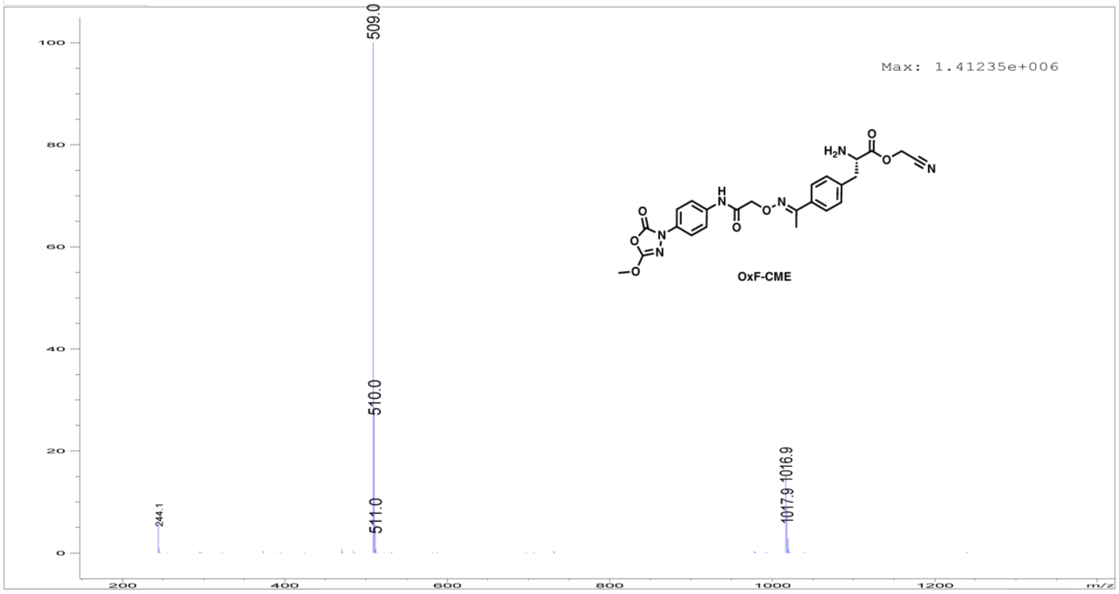

509.0
Max: 1.41235e+006
OxF-CME
510.0
244.1
511.0
1017.9 1016.9
m/z


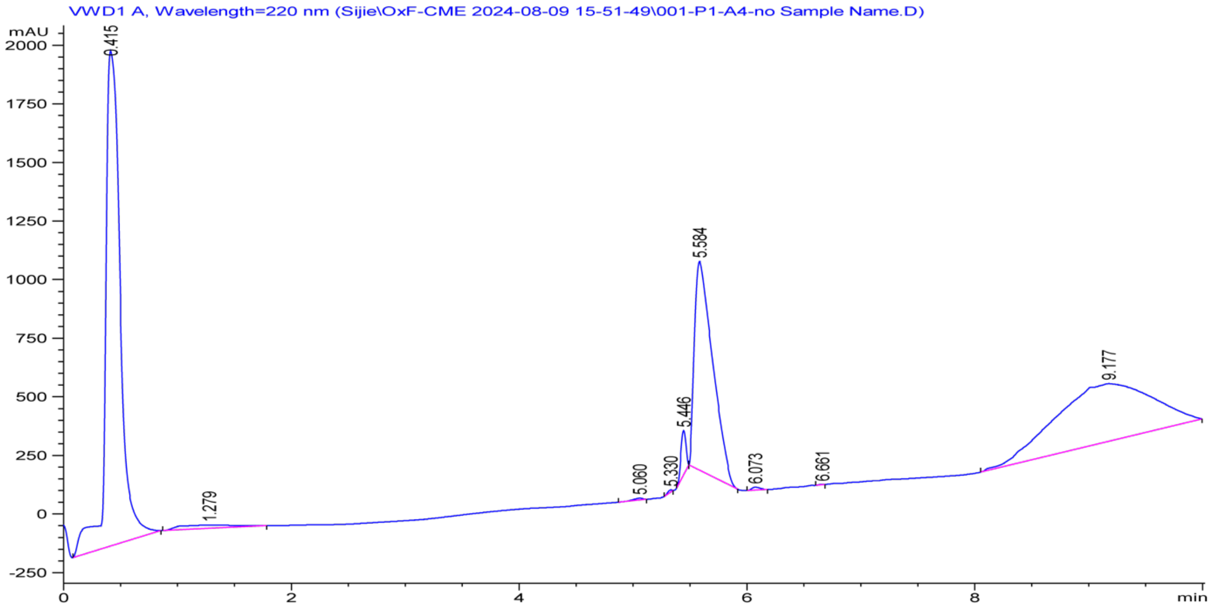

VWD1 A, Wavelength=220 nm (Sijie\OxF-CME 2024-08-09 15-51-49\001-P1-A4-no Sample Name.D)
mAU
0.415
1.279
5.060
5.330
5.446
5.584
6.073
6.661
9.177
min

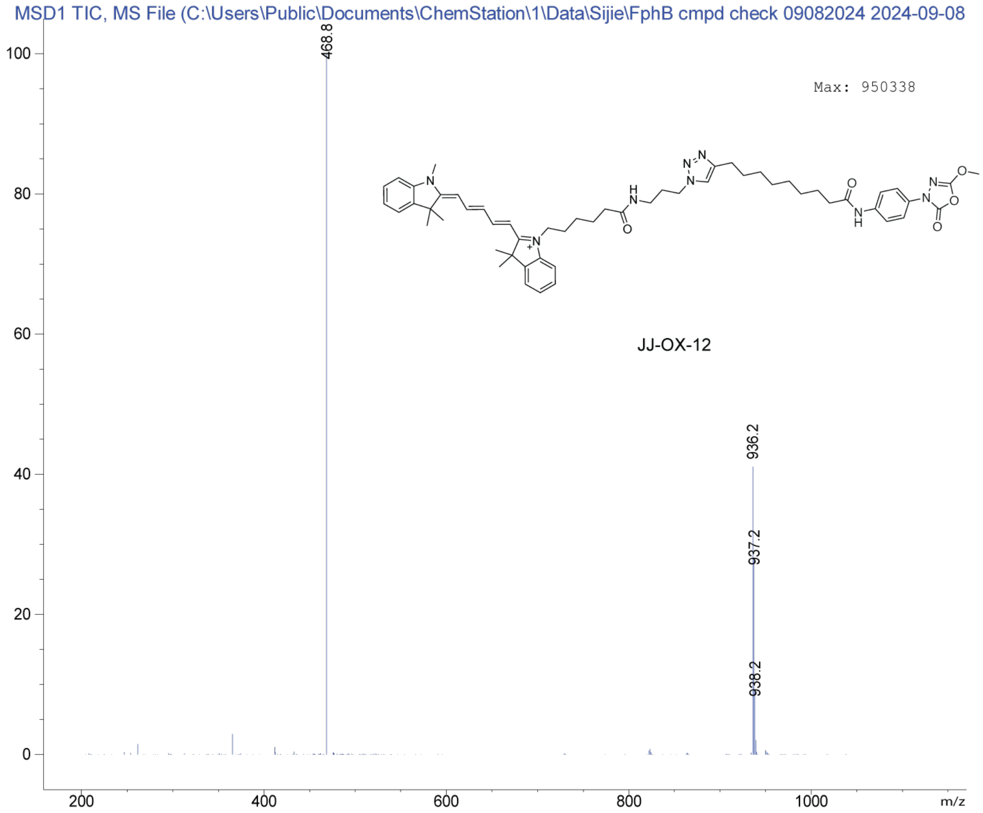
MSD1 TIC, MS File (C:\Users\Public\Documents\ChemStation\1\Data\Sijie\FphB cmpd check 09082024 2024-09-08
Max: 950338
468.8
936.2
937.2
938.2
JJ-OX-12
m/z

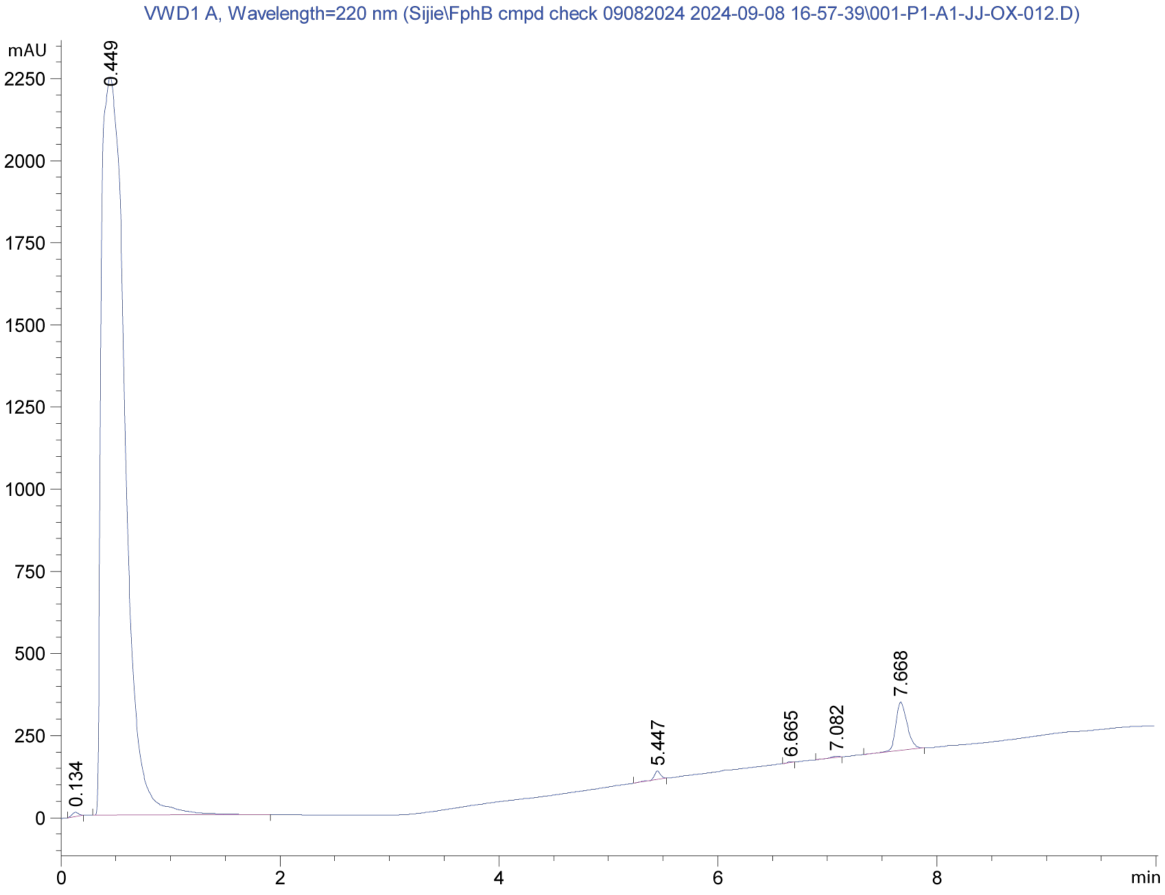
VWD1 A, Wavelength=220 nm (Sijie\FphB cmpd check 09082024 2024-09-08 16-57-39\001-P1-A1-JJ-OX-012.D)
mAU
0.134
0.449
5.447
6.665
7.082
7.668
min

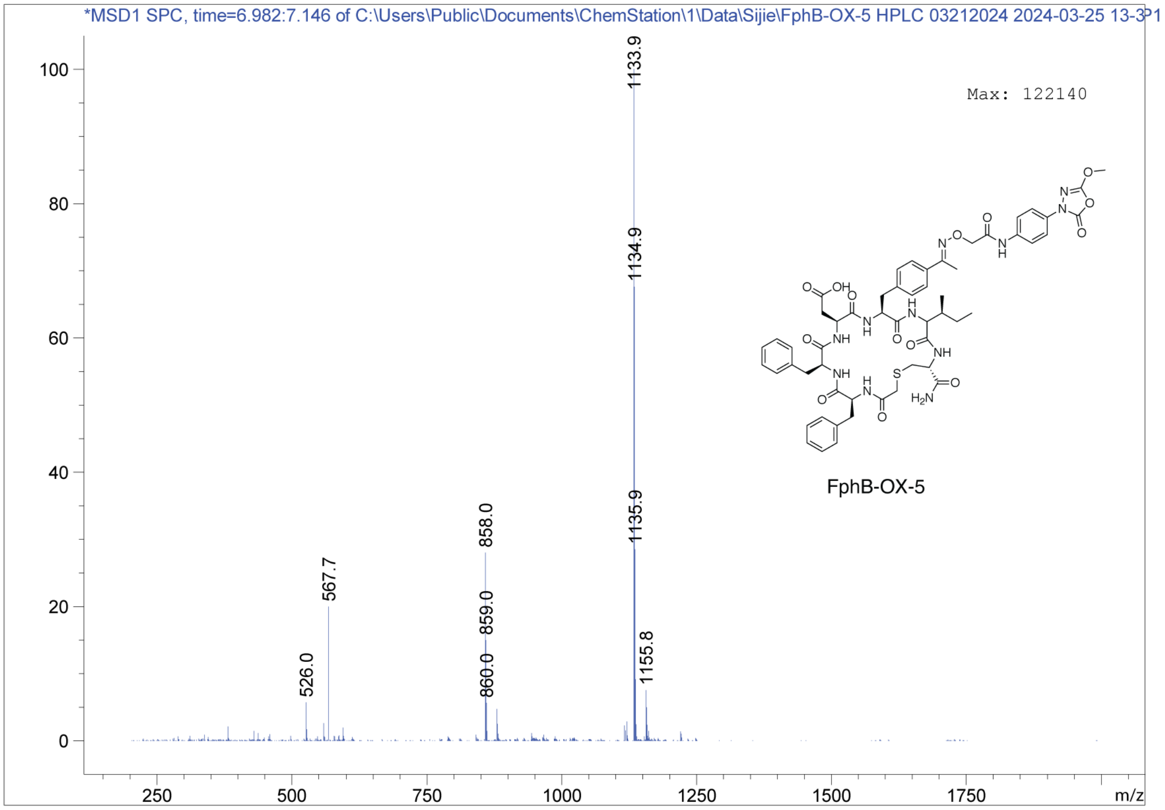
*MSD1 SPC, time=6.982:7.146 of C:\Users\Public\Documents\ChemStation\1\Data\Sijie\FphB-OX-5 HPLC 03212024 2024-03-25 13-3P1
Max: 122140
1133.9
1134.9
1135.9
1155.8
858.0
859.0
860.0
567.7
526.0
FphB-OX-5
m/z

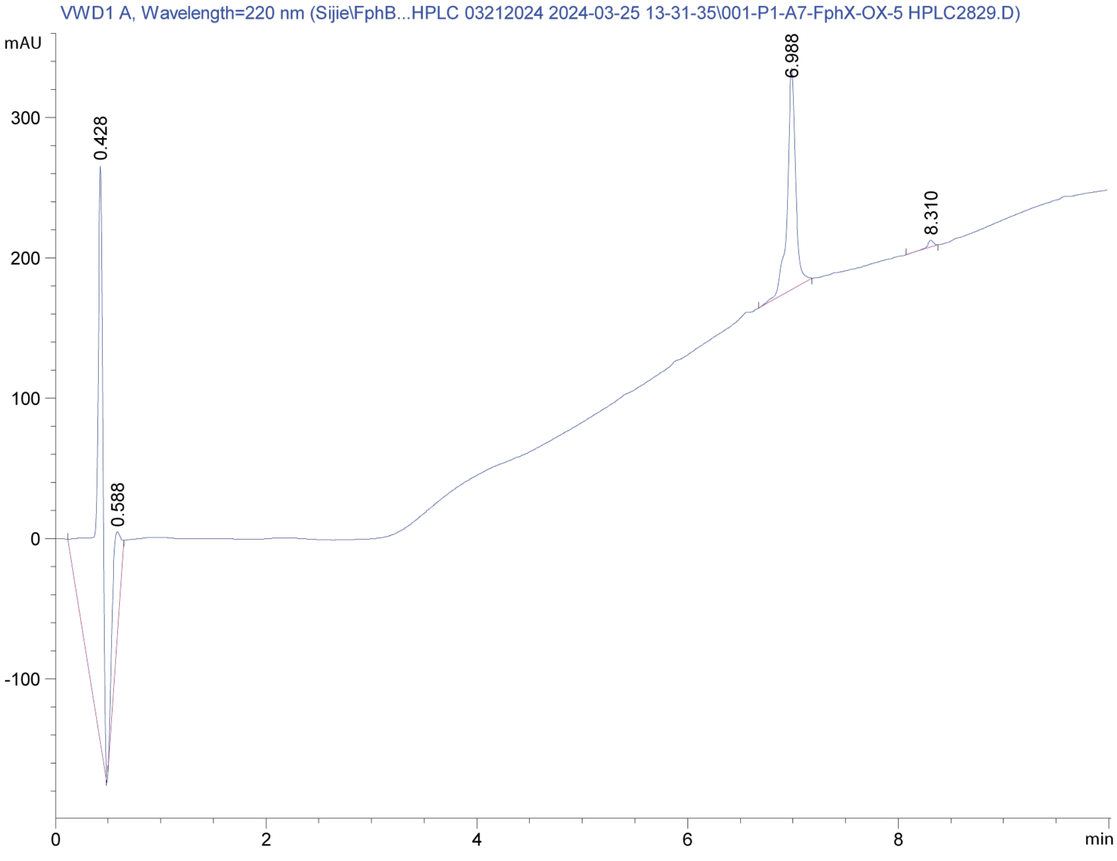
VWD1 A, Wavelength=220 nm (Sijie\FphB...HPLC 03212024 2024-03-25 13-31-35\001-P1-A7-FphX-OX-5 HPLC2829.D)
mAU
0.428
0.588
6.988
8.310
min

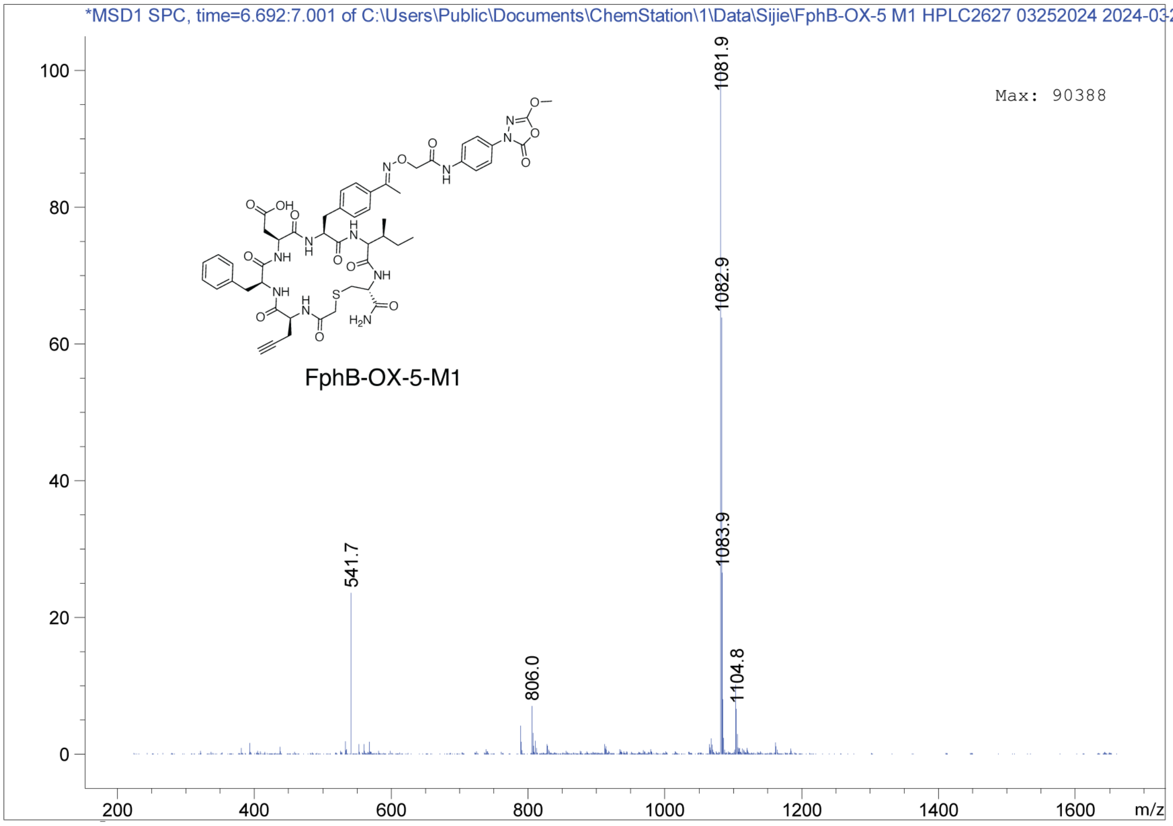
*MSD1 SPC, time=6.692:7.001 of C:\Users\Public\Documents\ChemStation\1\Data\Sijie\FphB-OX-5 M1 HPLC2627 03252024 2024-03-2
Max: 90388
1081.9
1082.9
1083.9
541.7
806.0
1104.8
FphB-OX-5-M1
0
20
40
60
80
100
200
400
600
800
1000
1200
1400
1600
m/z

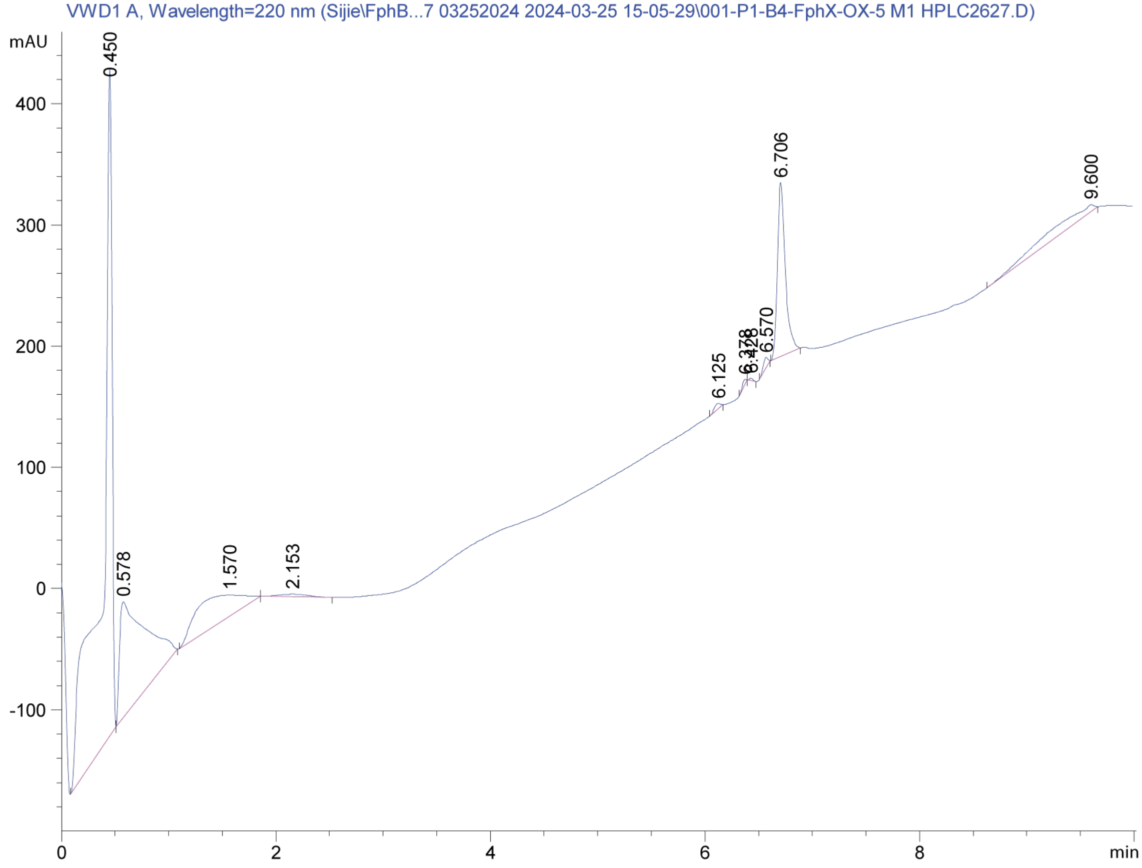
VWD1 A, Wavelength=220 nm (Sijie\FphB...7 03252024 2024-03-25 15-05-29\001-P1-B4-FphX-OX-5 M1 HPLC2627.D)
mAU
0.450
0.578
1.570
2.153
6.125
6.378
6.428
6.570
6.706
9.600
400
300
200
100
0
-100
0
2
4
6
8
min

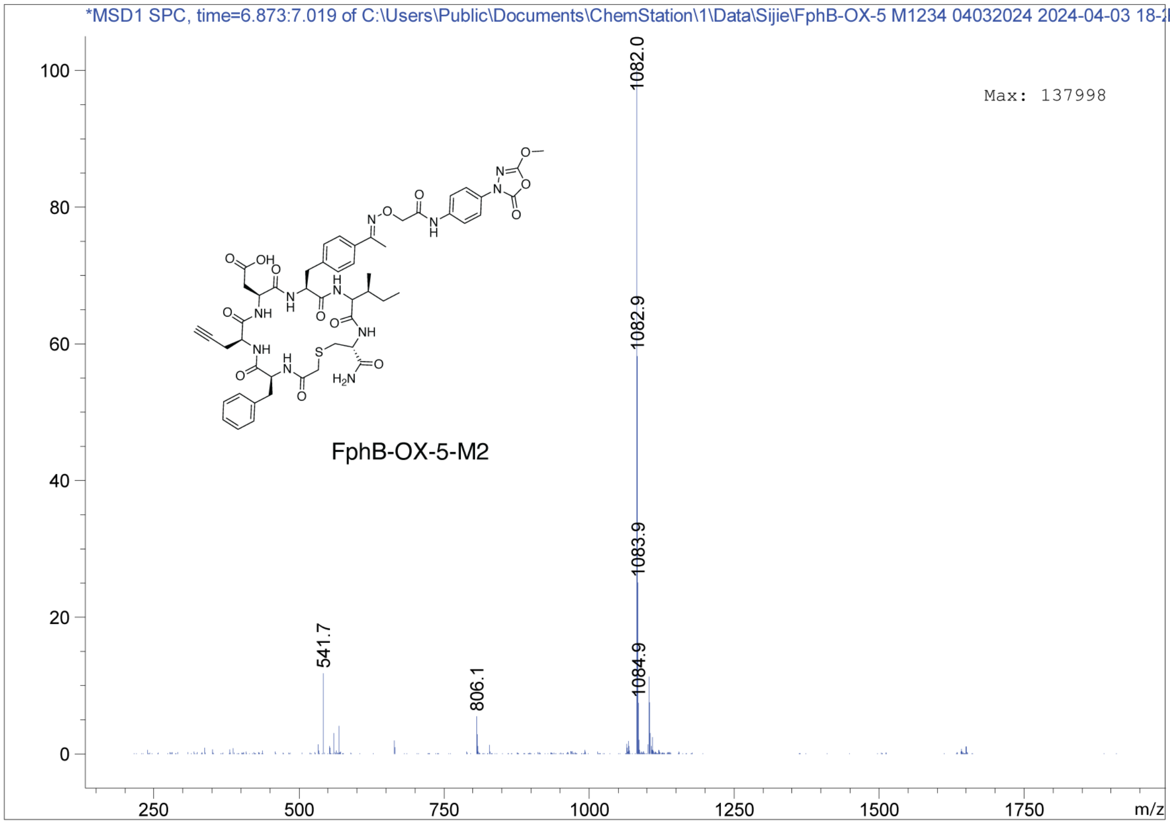
*MSD1 SPC, time=6.873:7.019 of C:\Users\Public\Documents\ChemStation\1\Data\Sijie\FphB-OX-5 M1234 04032024 2024-04-03 18-2P
Max: 137998
1082.0
1082.9
1083.9
1084.9
541.7
806.1
FphB-OX-5-M2
m/z

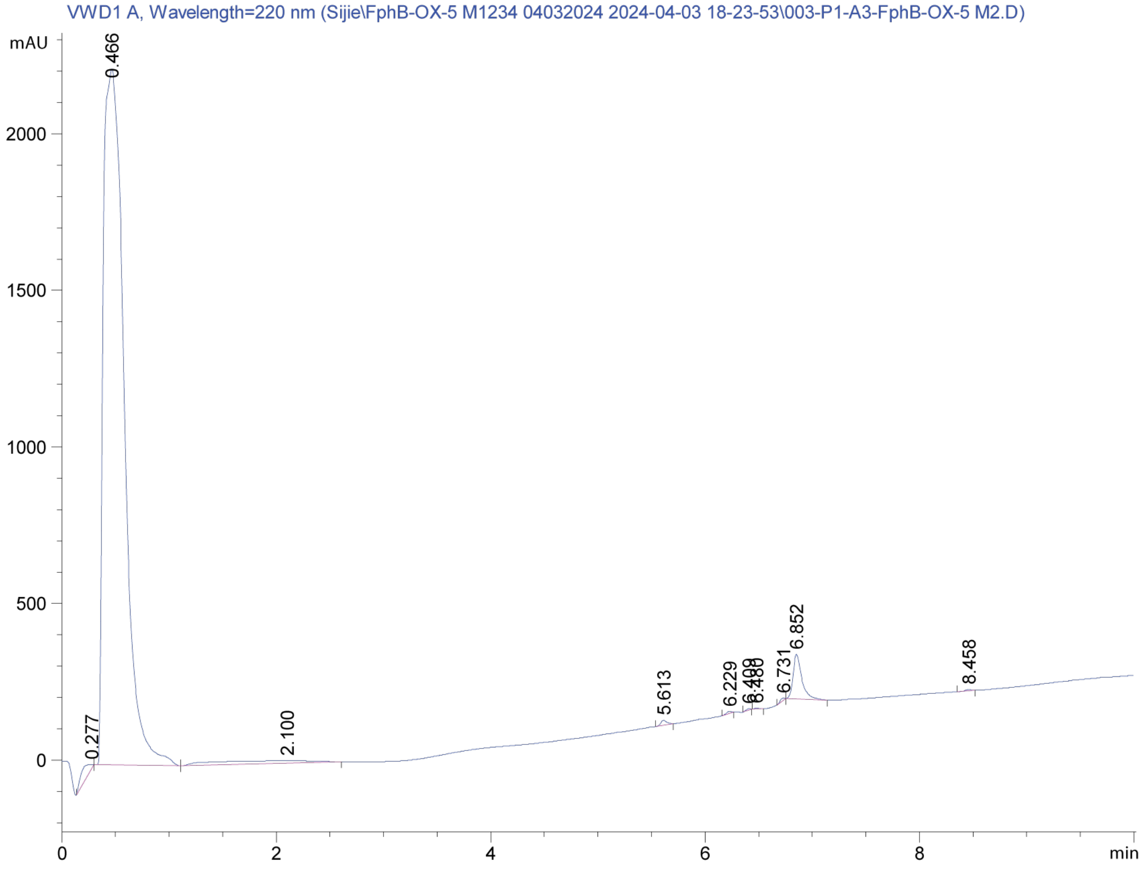
VWD1 A, Wavelength=220 nm (Sijie\FphB-OX-5 M1234 04032024 2024-04-03 18-23-53\003-P1-A3-FphB-OX-5 M2.D)
mAU
0.466
0.277
2.100
5.613
6.229
6.852
6.731
8.458
min

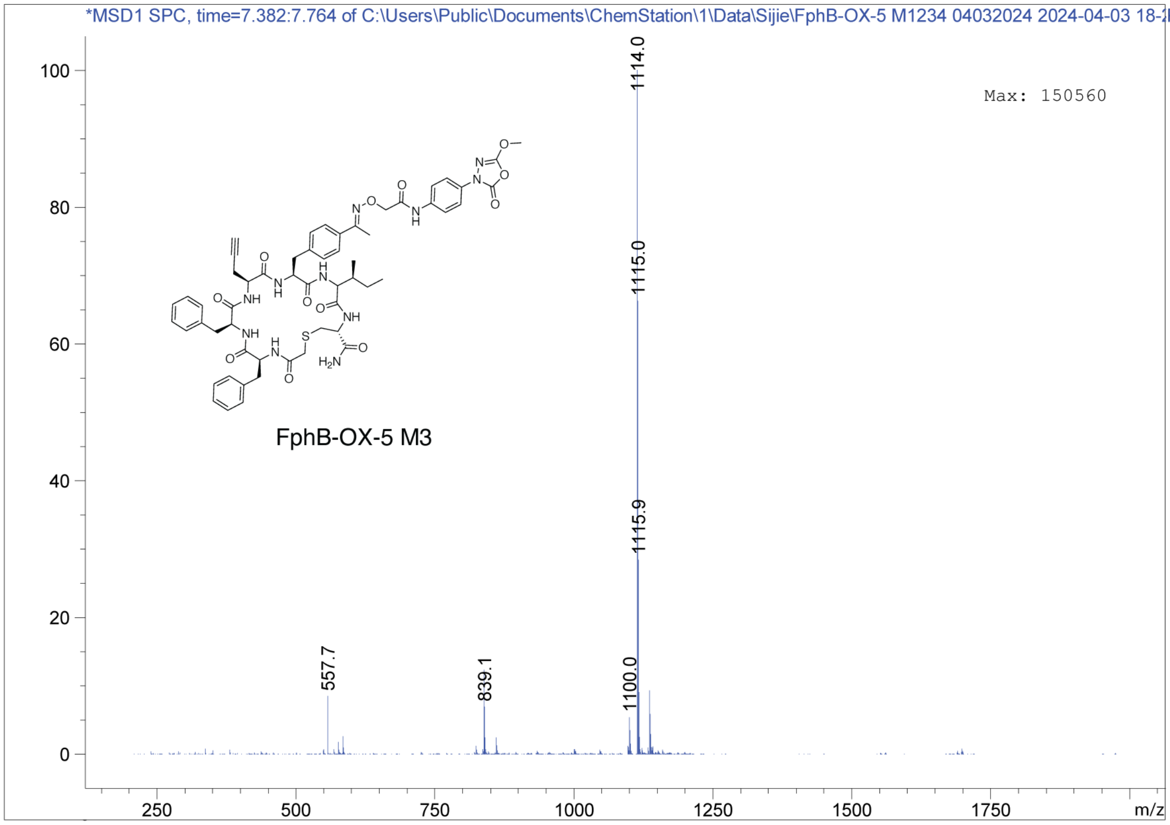
*MSD1 SPC, time=7.382:7.764 of C:\Users\Public\Documents\ChemStation\1\Data\Sijie\FphB-OX-5 M1234 04032024 2024-04-03 18-2P
Max: 150560
1114.0
1115.0
1115.9
557.7
839.1
1100.0
FphB-OX-5 M3
100
80
60
40
20
0
250
500
750
1000
1250
1500
1750
m/z

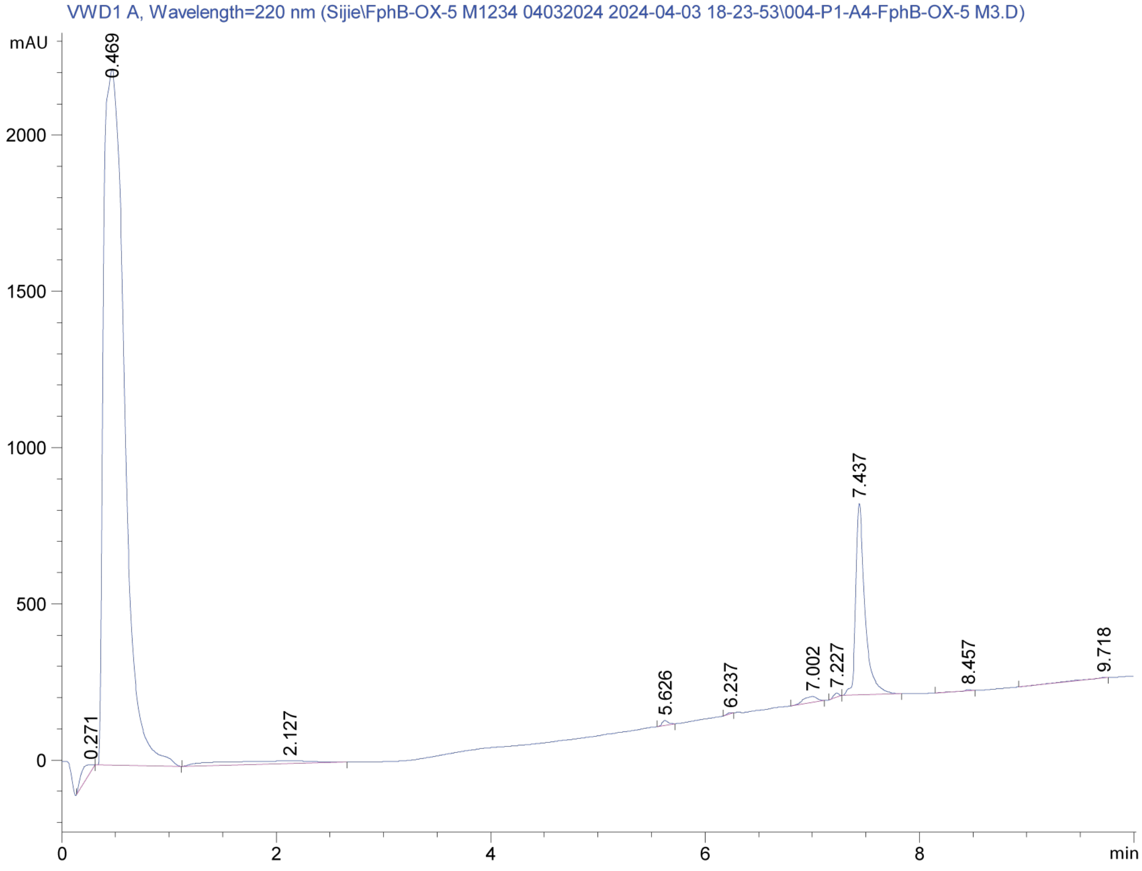
VWD1 A, Wavelength=220 nm (Sijie\FphB-OX-5 M1234 04032024 2024-04-03 18-23-53\004-P1-A4-FphB-OX-5 M3.D)
mAU
0.469
0.271
2.127
5.626
6.237
7.002
7.227
7.437
8.457
9.718
2000
1500
1000
500
0
0
2
4
6
8
min

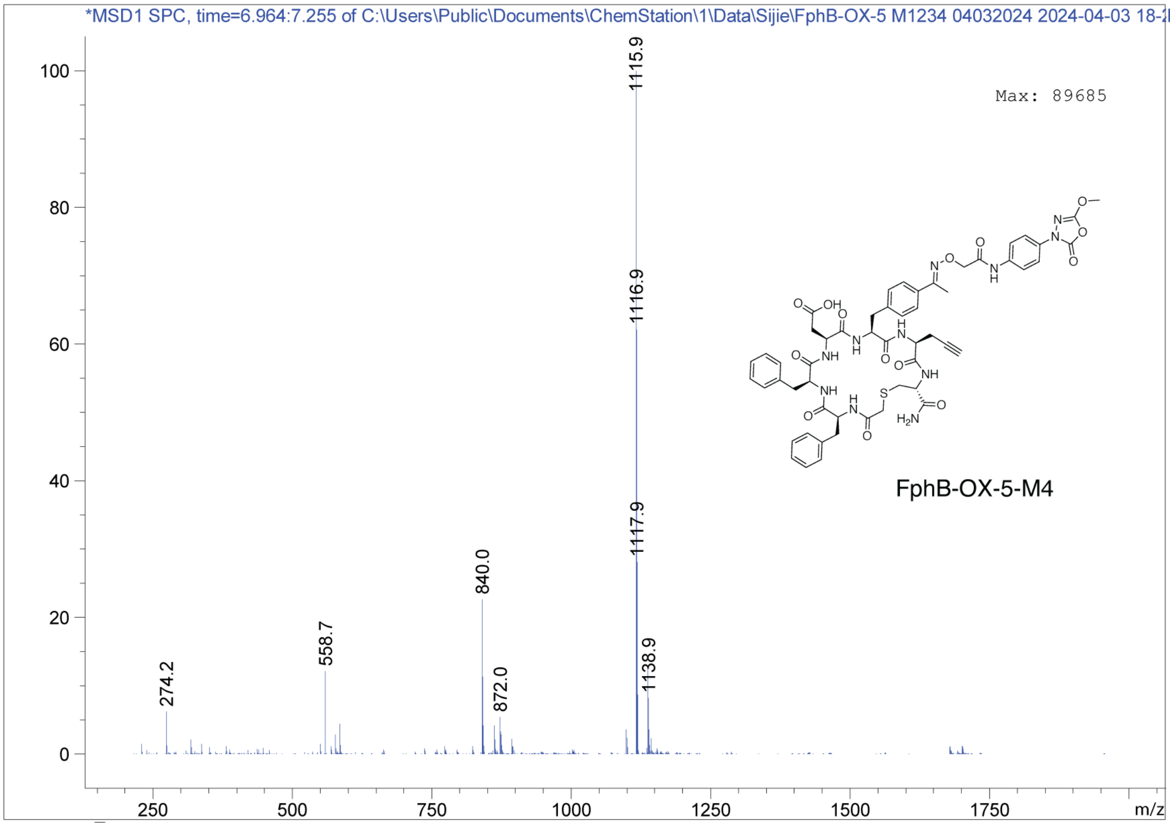
*MSD1 SPC, time=6.964:7.255 of C:\Users\Public\Documents\ChemStation\1\Data\Sijie\FphB-OX-5 M1234 04032024 2024-04-03 18-2P
Max: 89685
1115.9
1116.9
1117.9
840.0
558.7
274.2
872.0
1138.9
FphB-OX-5-M4
m/z

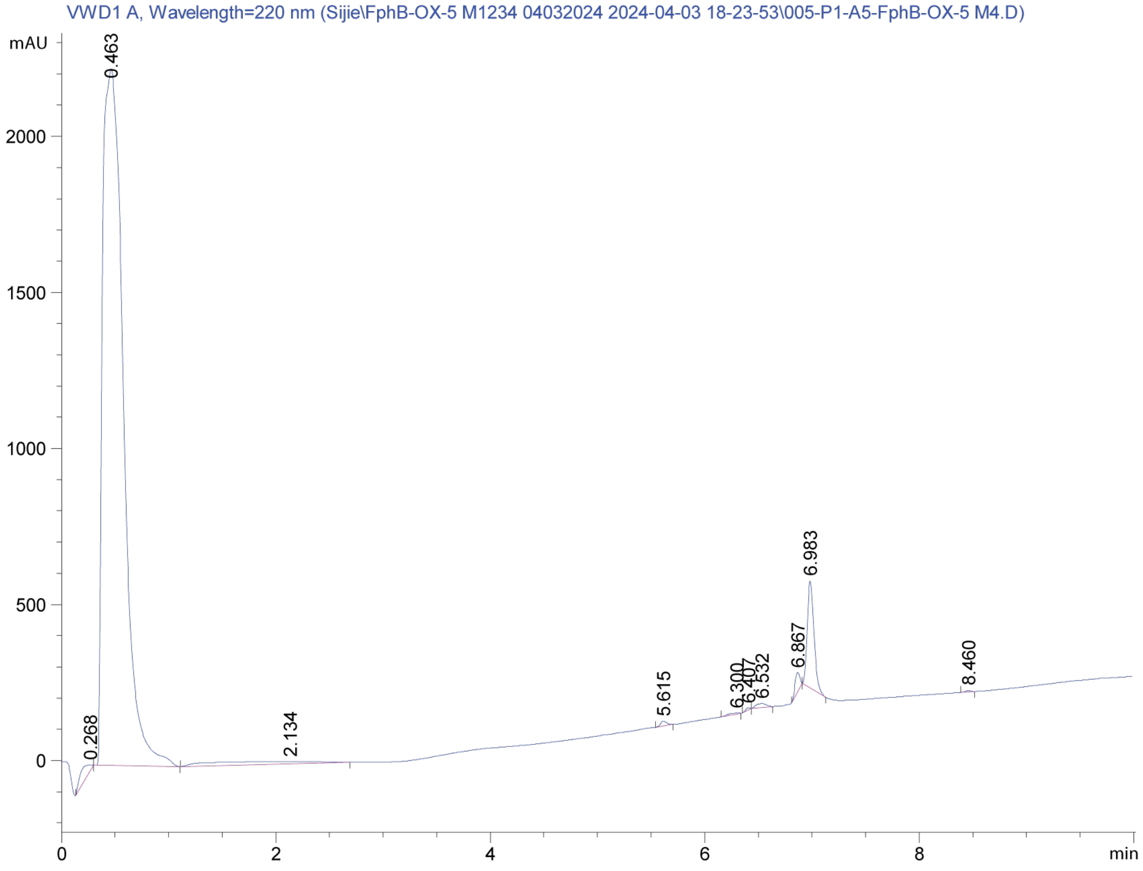
VWD1 A, Wavelength=220 nm (Sijie\FphB-OX-5 M1234 04032024 2024-04-03 18-23-53\005-P1-A5-FphB-OX-5 M4.D)
mAU
0.463
0.268
2.134
5.615
6.300
6.407
6.532
6.867
6.983
8.460
min

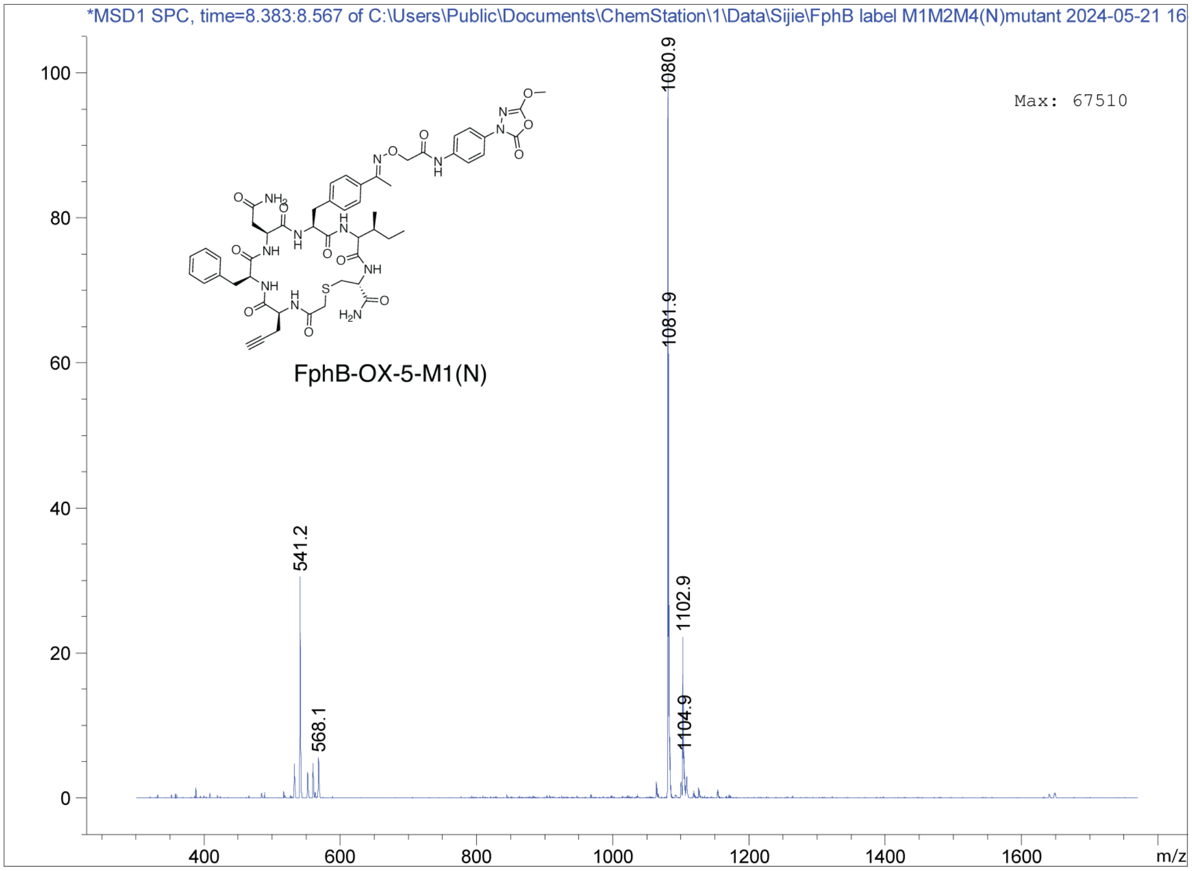
*MSD1 SPC, time=8.383:8.567 of C:\Users\Public\Documents\ChemStation\1\Data\Sijie\FphB label M1M2M4(N)mutant 2024-05-21 16
Max: 67510
1080.9
1081.9
541.2
1102.9
1104.9
568.1
FphB-OX-5-M1(N)
m/z

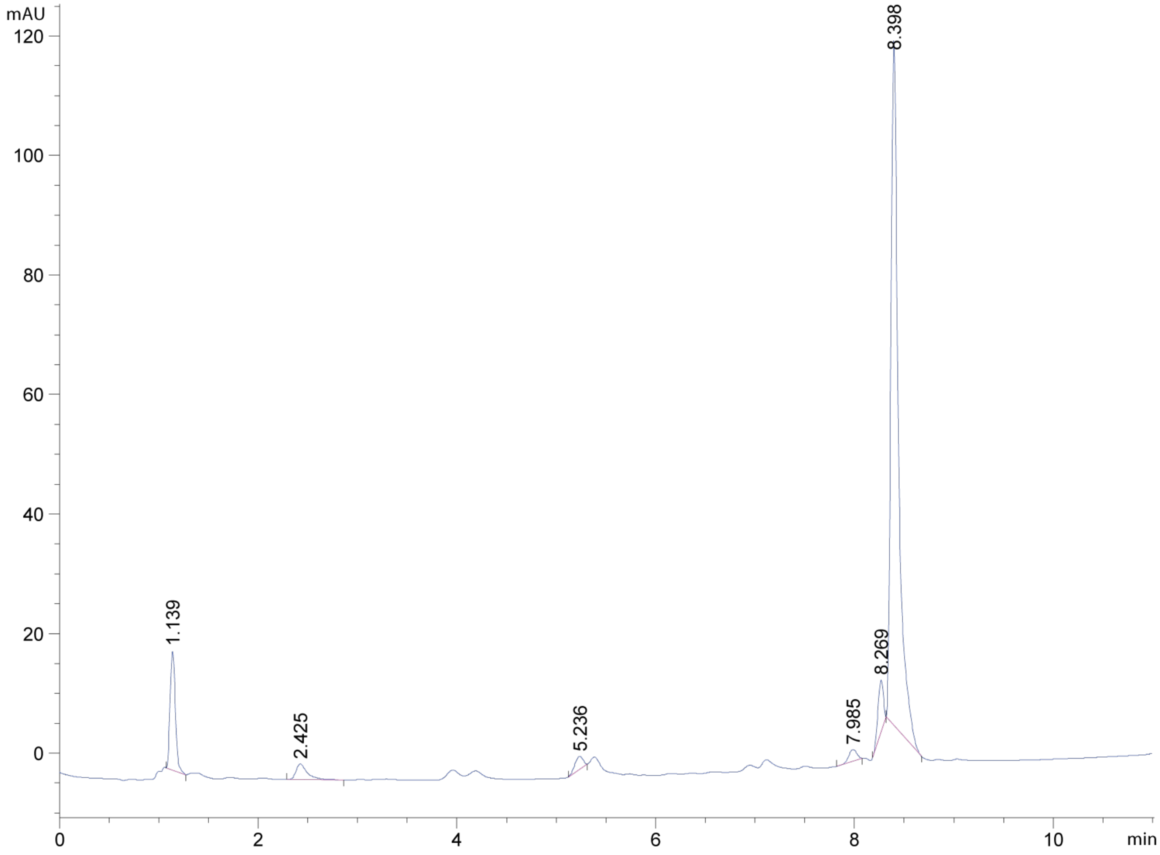
mAU
8.398
1.139
2.425
5.236
7.985
8.269
min

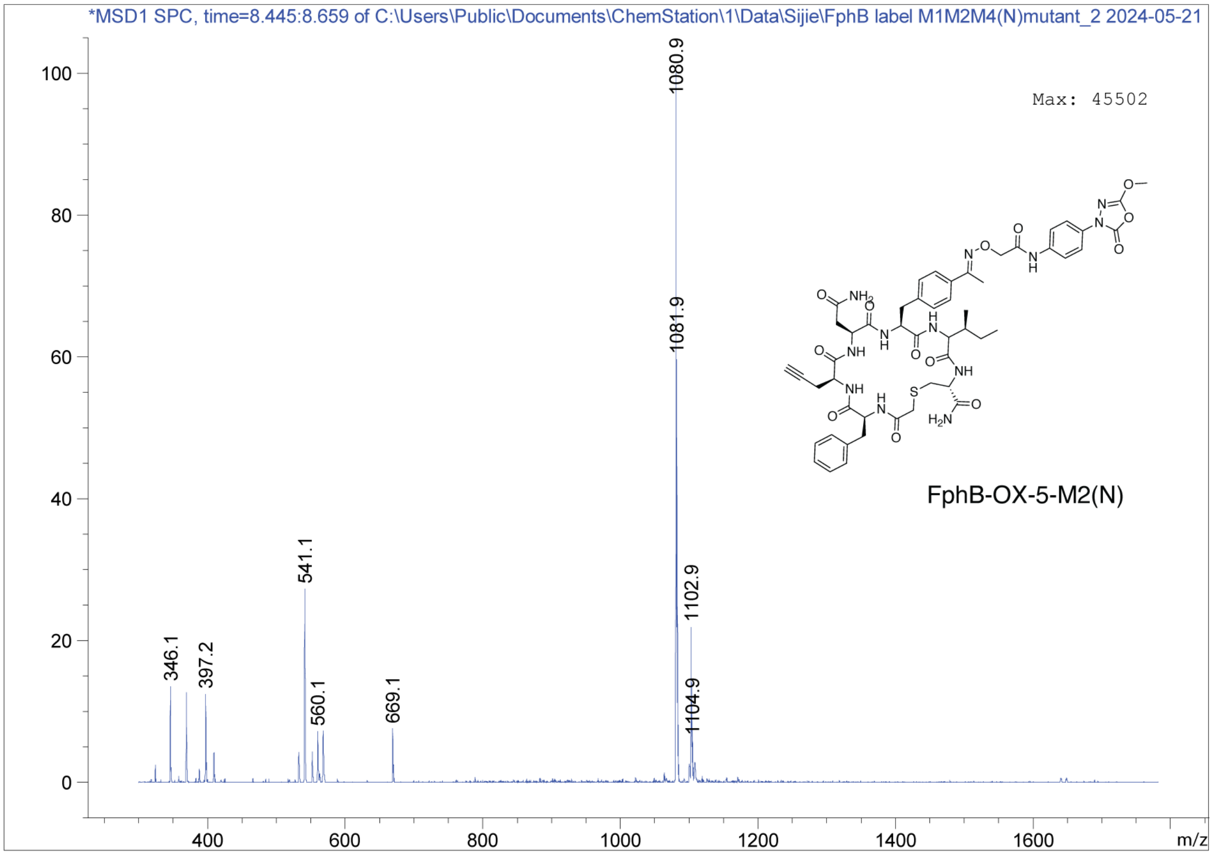
*MSD1 SPC, time=8.445:8.659 of C:\Users\Public\Documents\ChemStation\1\Data\Sijie\FphB label M1M2M4(N)mutant_2 2024-05-21
Max: 45502
1080.9
1081.9
541.1
1102.9
346.1
397.2
560.1
669.1
1104.9
m/z
FphB-OX-5-M2(N)

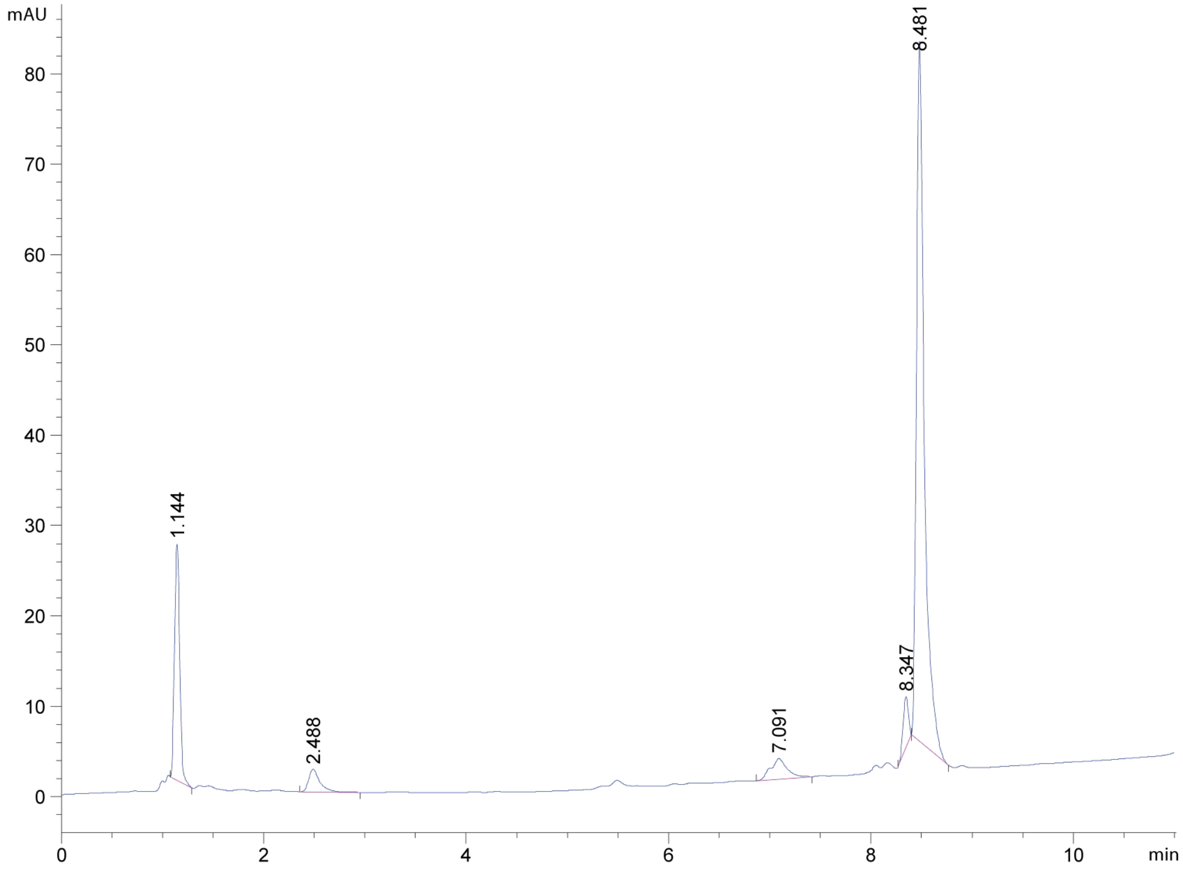
mAU
8.481
1.144
2.488
7.091
8.347
min

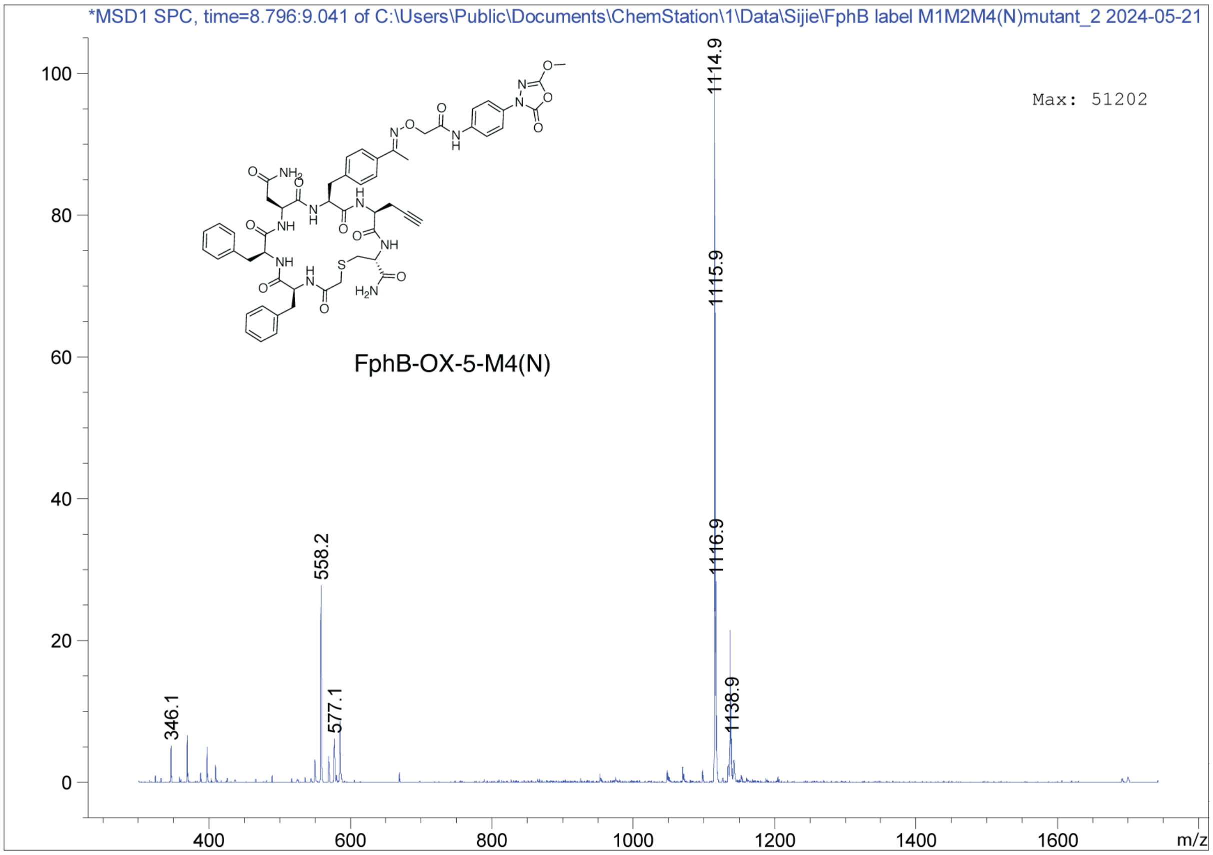
*MSD1 SPC, time=8.796:9.041 of C:\Users\Public\Documents\ChemStation\1\Data\Sijie\FphB label M1M2M4(N)mutant_2 2024-05-21
Max: 51202
FphB-OX-5-M4(N)
1114.9
1115.9
1116.9
558.2
346.1
577.1
1138.9
100
80
60
40
20
0
400
600
800
1000
1200
1400
1600
m/z

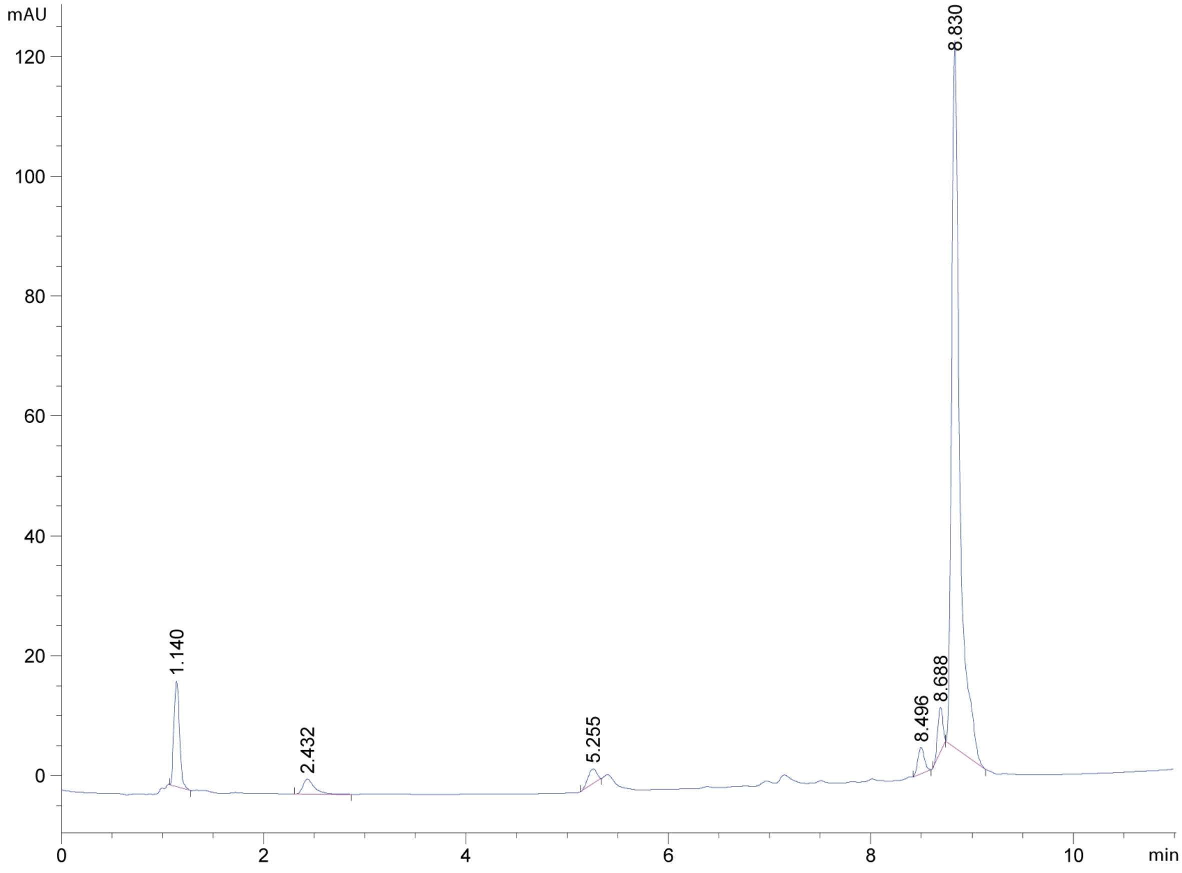
mAU
120
100
80
60
40
20
0
8.830
1.140
2.432
5.255
8.496
8.688
0
2
4
6
8
10
min

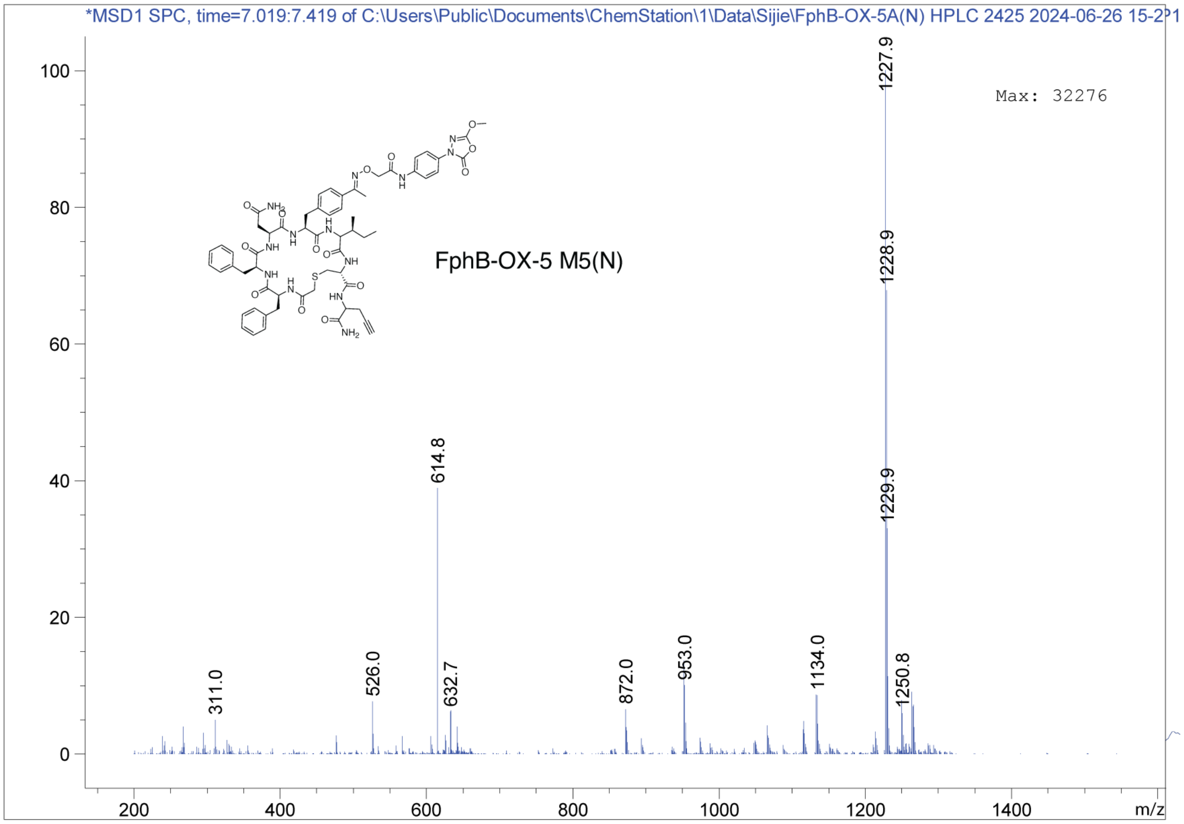
*MSD1 SPC, time=7.019:7.419 of C:\Users\Public\Documents\ChemStation\1\Data\Sijie\FphB-OX-5A(N) HPLC 2425 2024-06-26 15-2?1
Max: 32276
FphB-OX-5 M5(N)
1227.9
1228.9
1229.9
614.8
311.0
526.0
632.7
872.0
953.0
1134.0
1250.8
m/z

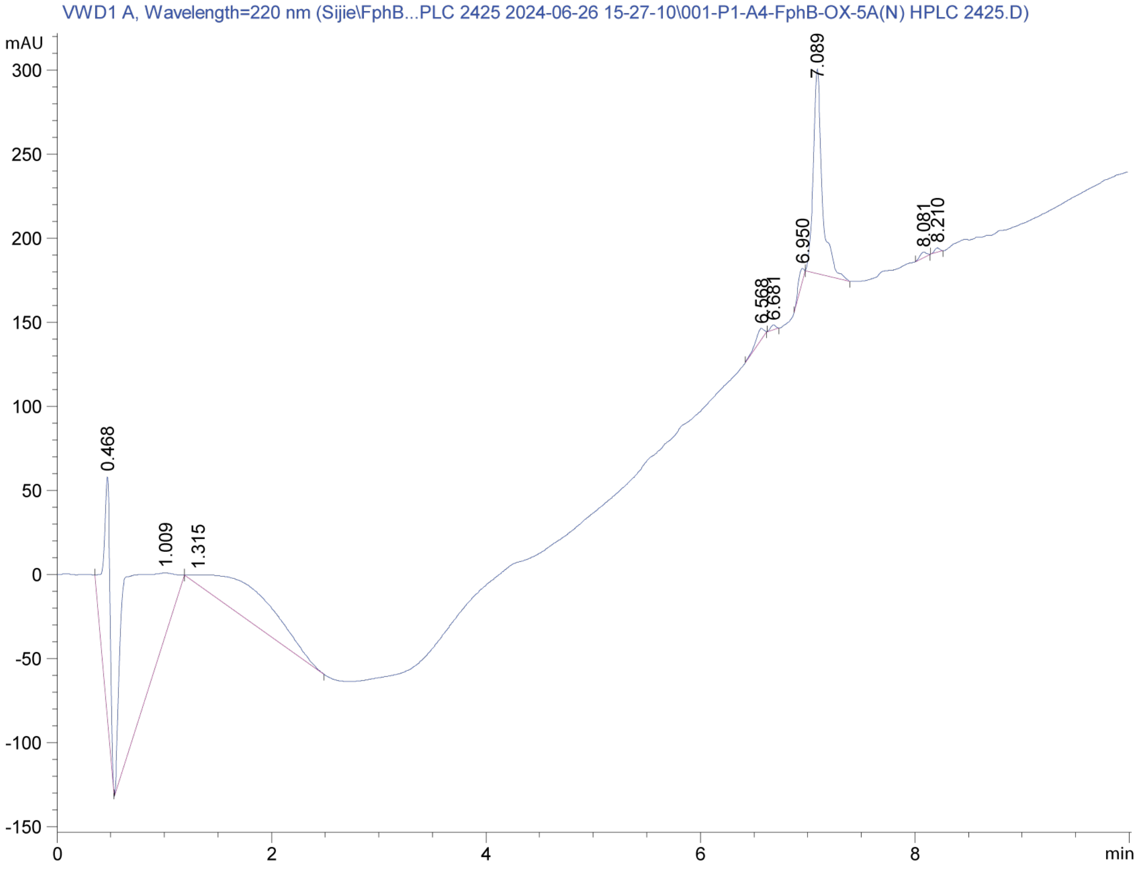
VWD1 A, Wavelength=220 nm (Sijie\FphB...PLC 2425 2024-06-26 15-27-10\001-P1-A4-FphB-OX-5A(N) HPLC 2425.D)
mAU
0.468
1.009
1.315
6.568
6.681
6.950
7.089
8.081
8.210
min

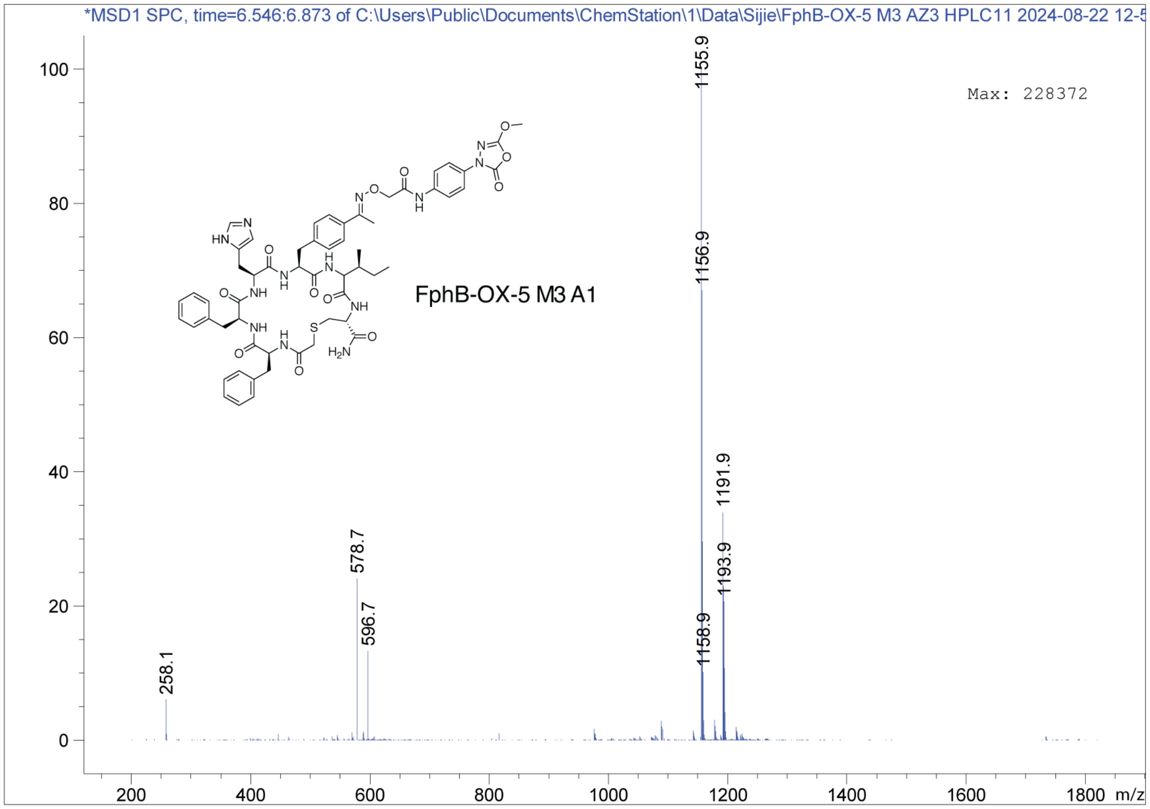
*MSD1 SPC, time=6.546:6.873 of C:\Users\Public\Documents\ChemStation\1\Data\Sijie\FphB-OX-5 M3 AZ3 HPLC11 2024-08-22 12-5
Max: 228372
FphB-OX-5 M3 A1
1155.9
1156.9
1191.9
1193.9
1158.9
578.7
596.7
258.1
m/z

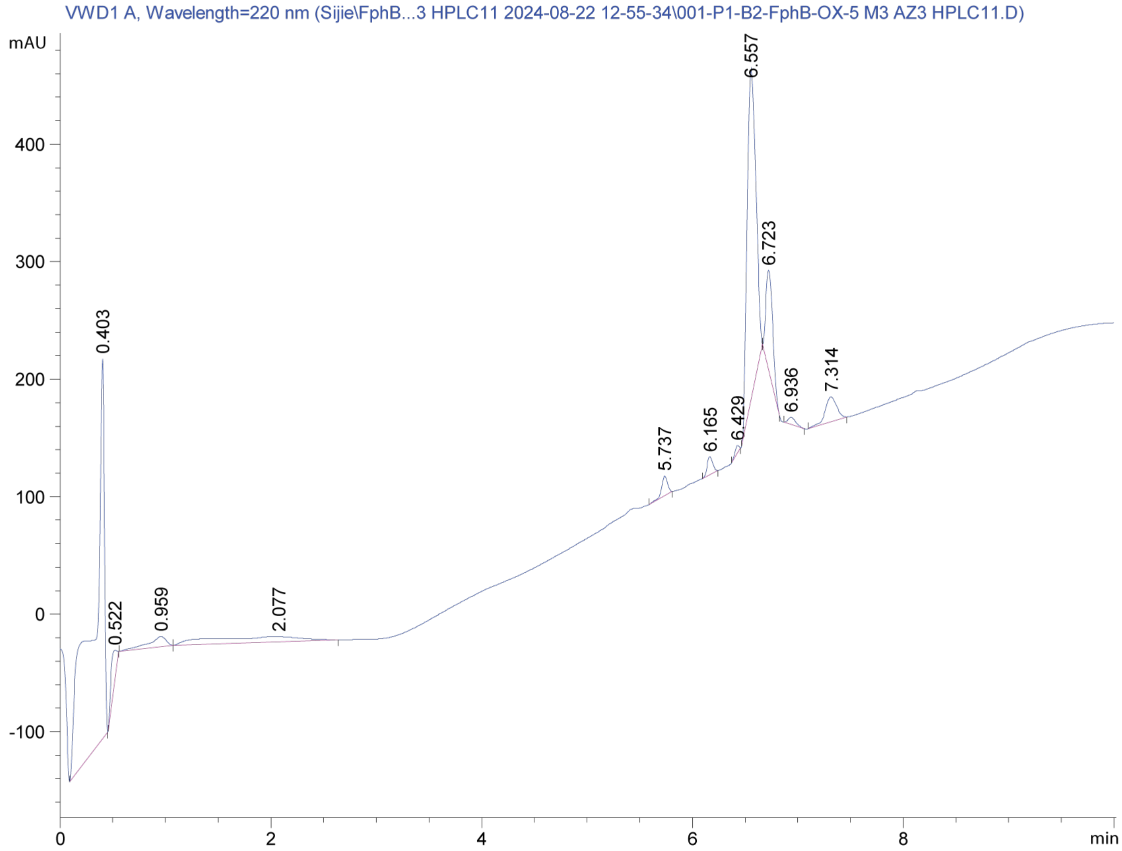
VWD1 A, Wavelength=220 nm (Sijie\FphB...3 HPLC11 2024-08-22 12-55-34\001-P1-B2-FphB-OX-5 M3 AZ3 HPLC11.D)
mAU
6.557
6.723
0.403
7.314
6.936
6.429
6.165
5.737
0.522
0.959
2.077
min

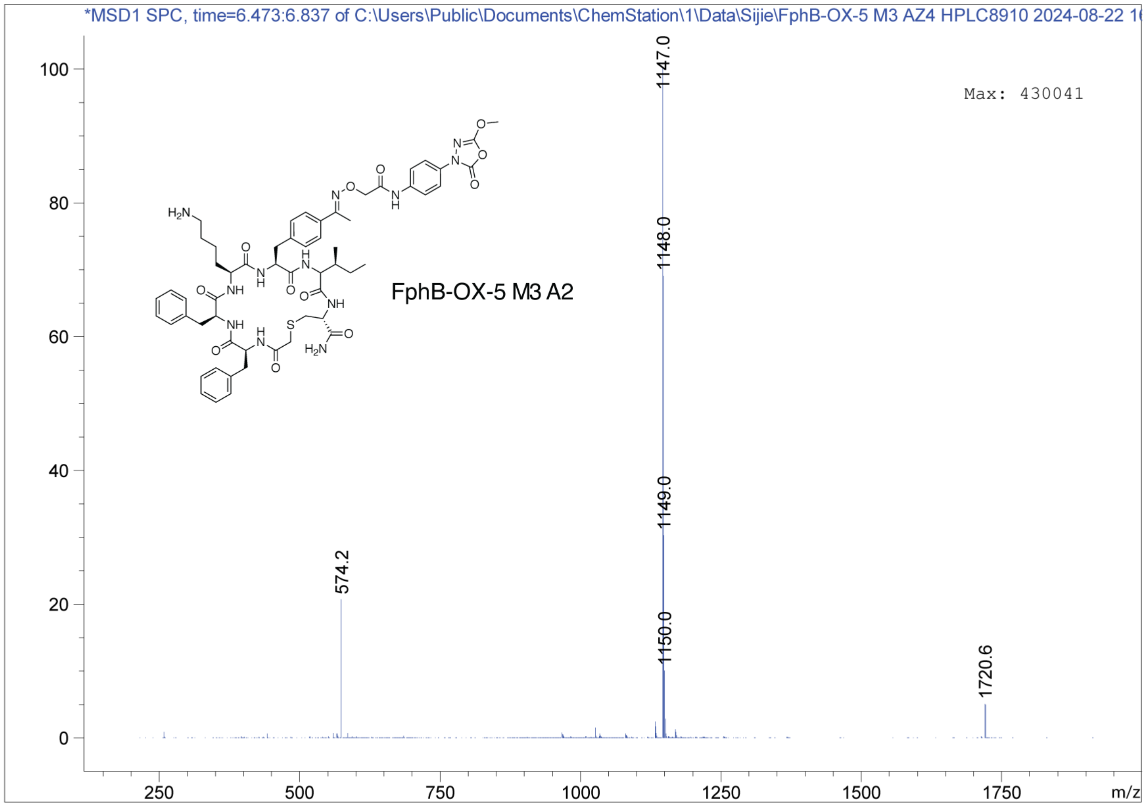
*MSD1 SPC, time=6.473:6.837 of C:\Users\Public\Documents\ChemStation\1\Data\Sijie\FphB-OX-5 M3 AZ4 HPLC8910 2024-08-22 1
Max: 430041
FphB-OX-5 M3 A2
1147.0
1148.0
1149.0
1150.0
574.2
1720.6
m/z

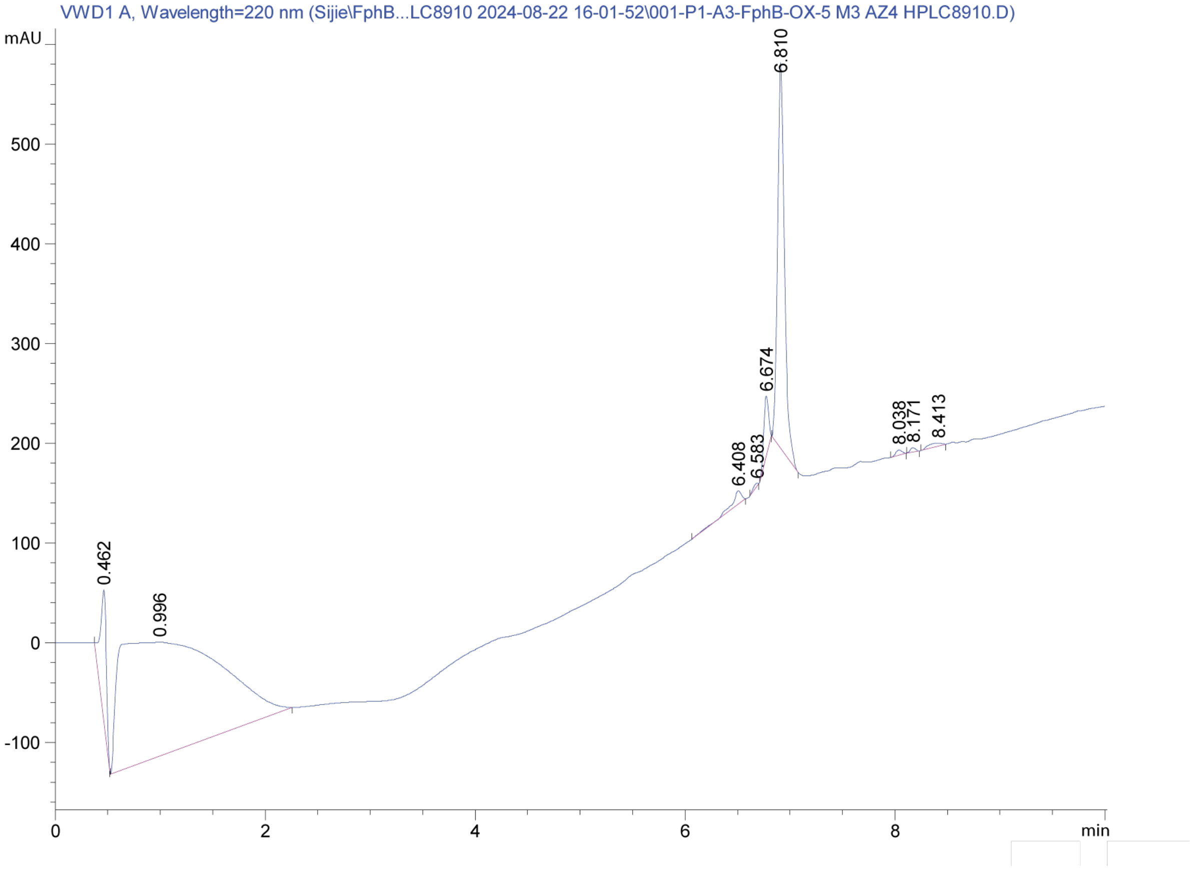
VWD1 A, Wavelength=220 nm (Sijie\FphB...LC8910 2024-08-22 16-01-52\001-P1-A3-FphB-OX-5 M3 AZ4 HPLC8910.D)
mAU
6.810
6.674
6.408
6.583
8.038
8.171
8.413
0.462
0.996
min

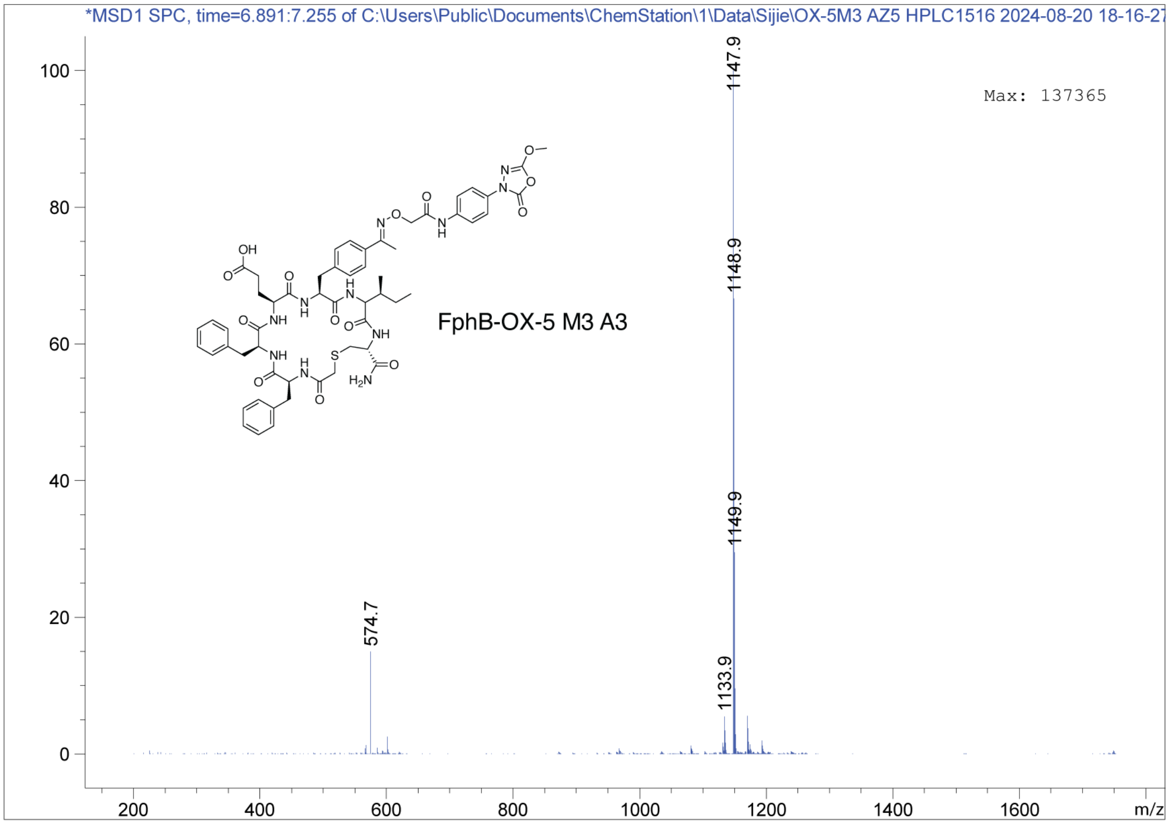
*MSD1 SPC, time=6.891:7.255 of C:\Users\Public\Documents\ChemStation\1\Data\Sijie\OX-5M3 AZ5 HPLC1516 2024-08-20 18-16-27
Max: 137365
FphB-OX-5 M3 A3
1147.9
1148.9
1149.9
1133.9
574.7
m/z

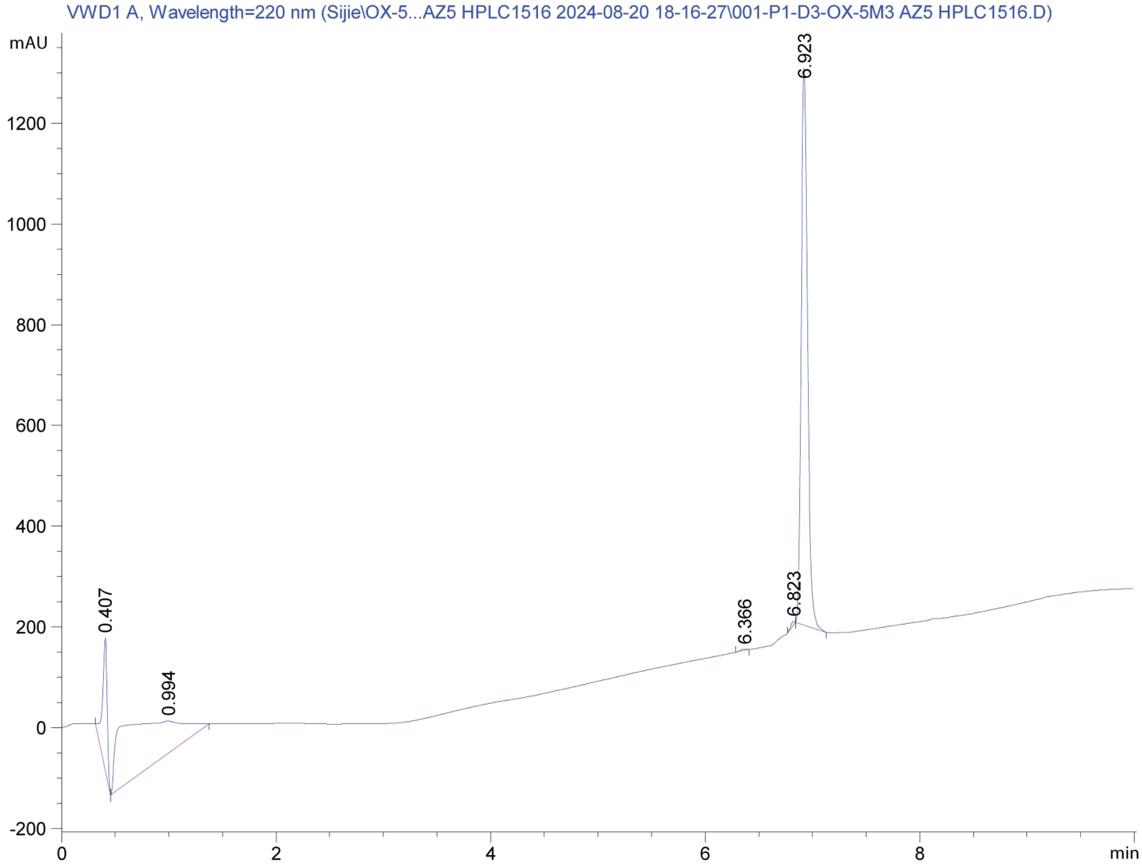
VWD1 A, Wavelength=220 nm (Sijie\OX-5...AZ5 HPLC1516 2024-08-20 18-16-27\001-P1-D3-OX-5M3 AZ5 HPLC1516.D)
mAU
6.923
6.823
6.366
0.407
0.994
min

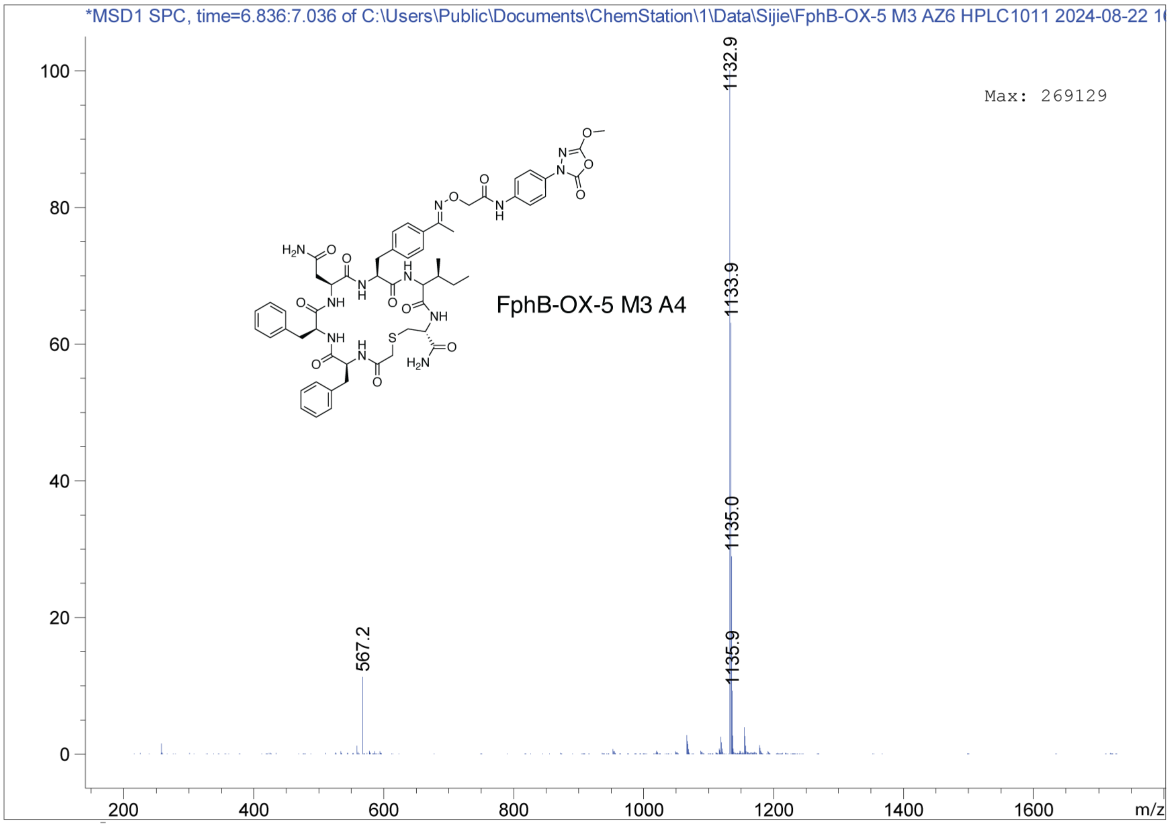
*MSD1 SPC, time=6.836:7.036 of C:\Users\Public\Documents\ChemStation\1\Data\Sijie\FphB-OX-5 M3 AZ6 HPLC1011 2024-08-22 1
Max: 269129
1132.9
1133.9
1135.0
1135.9
567.2
FphB-OX-5 M3 A4
m/z

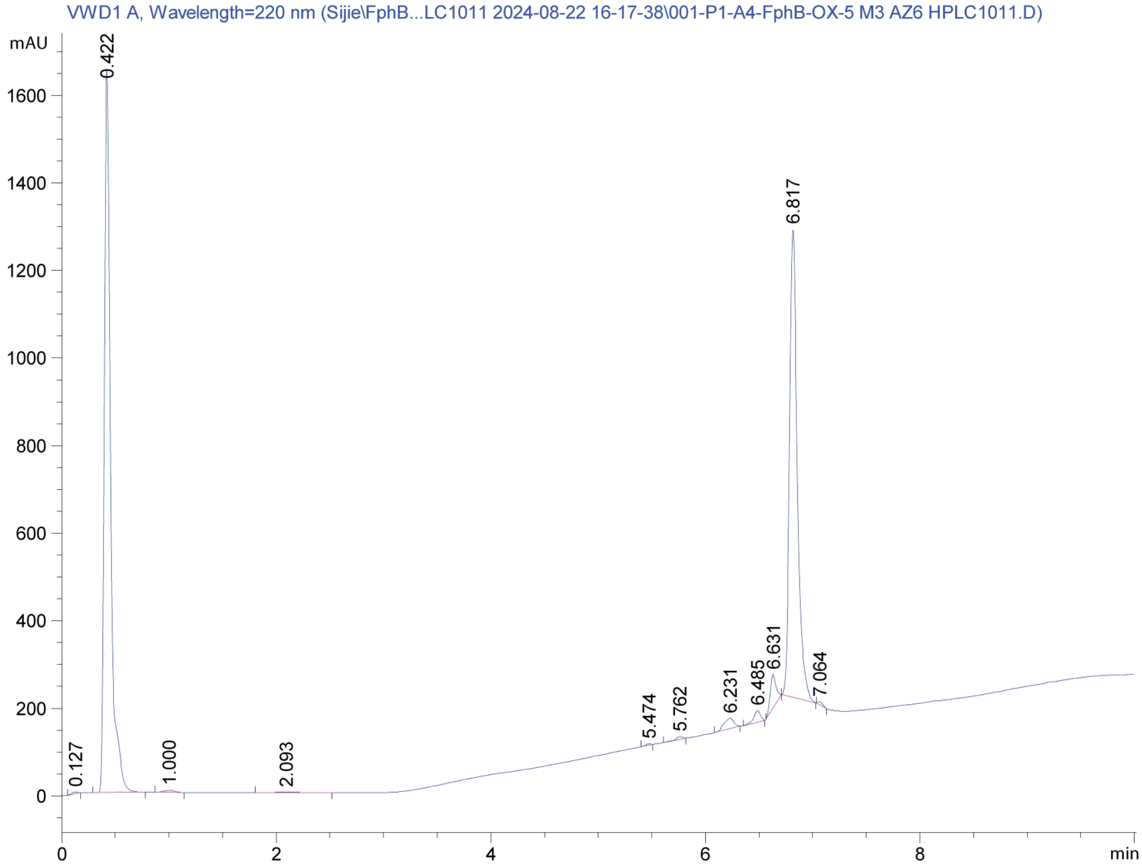
VWD1 A, Wavelength=220 nm (Sijie\FphB...LC1011 2024-08-22 16-17-38\001-P1-A4-FphB-OX-5 M3 AZ6 HPLC1011.D)
mAU
0.422
6.817
0.127
1.000
2.093
5.474
5.762
6.231
6.485
6.631
7.064
min

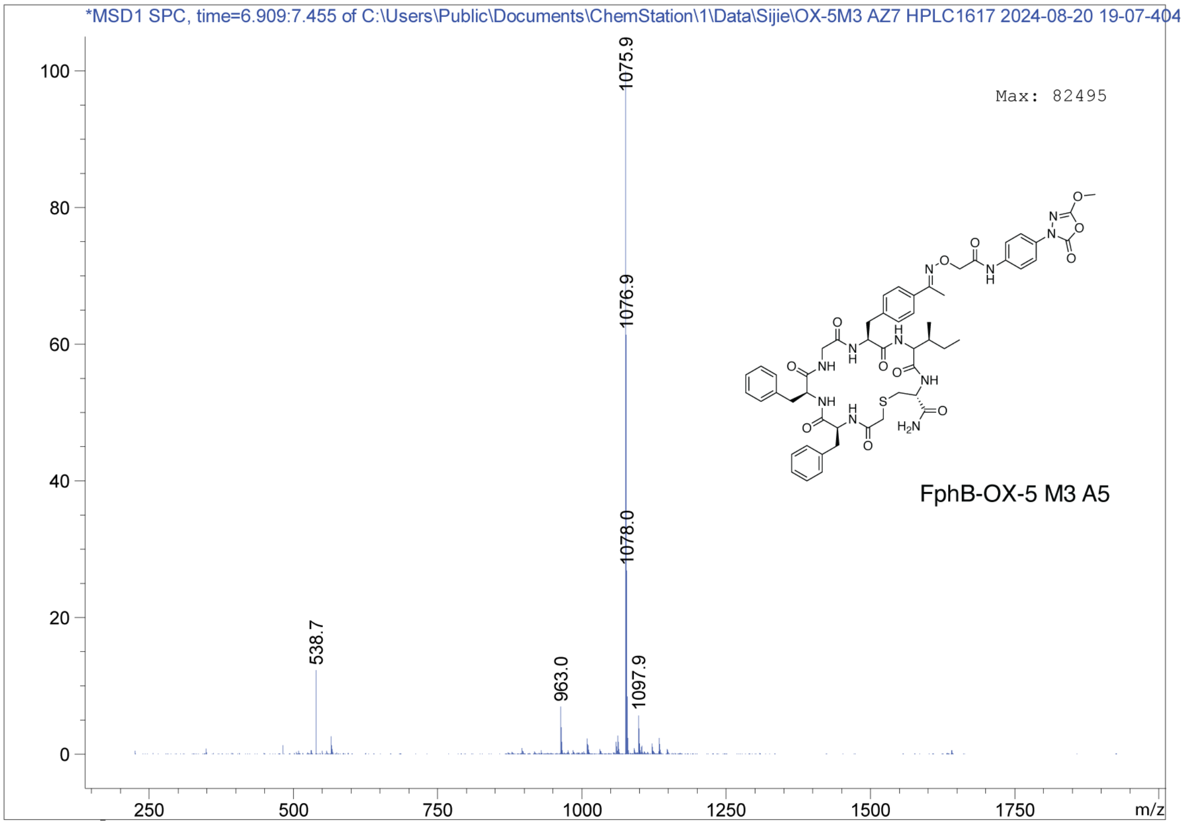
*MSD1 SPC, time=6.909:7.455 of C:\Users\Public\Documents\ChemStation\1\Data\Sijie\OX-5M3 AZ7 HPLC1617 2024-08-20 19-07-404
Max: 82495
1075.9
1076.9
1078.0
538.7
963.0
1097.9
FphB-OX-5 M3 A5
m/z

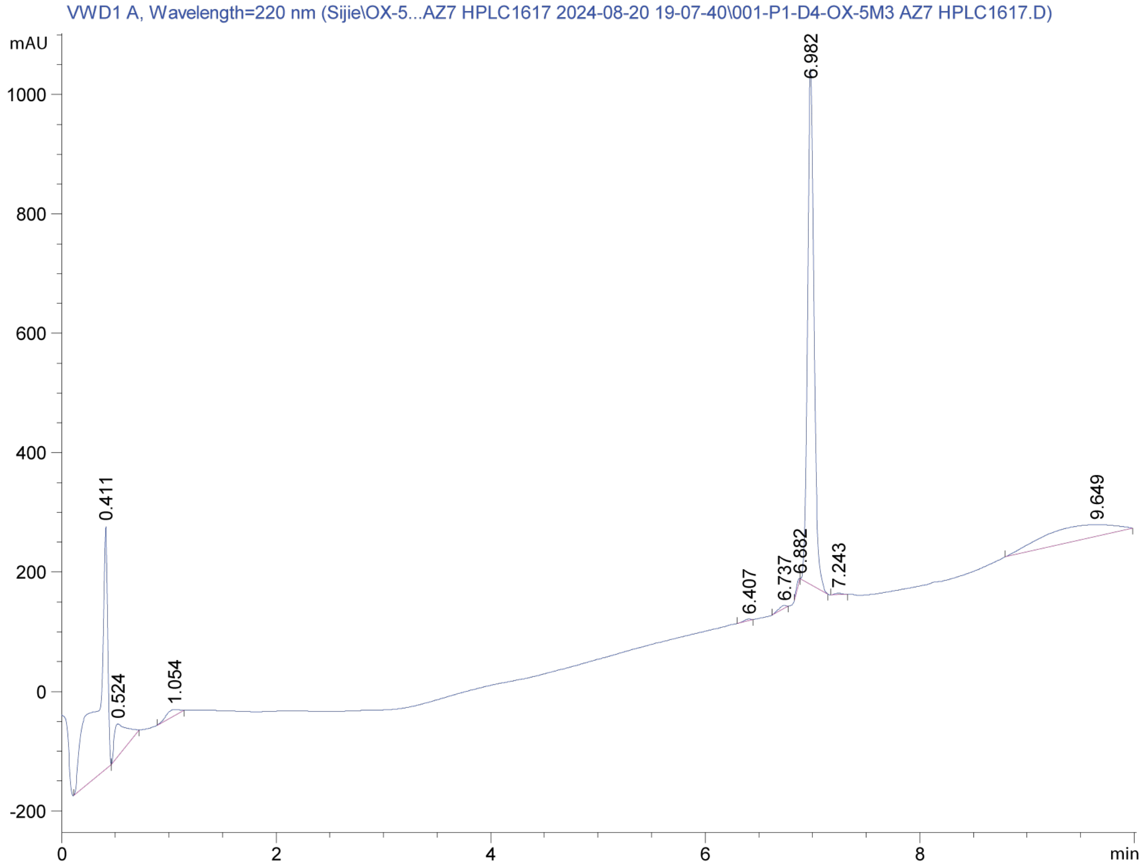
VWD1 A, Wavelength=220 nm (Sijie\OX-5...AZ7 HPLC1617 2024-08-20 19-07-40\001-P1-D4-OX-5M3 AZ7 HPLC1617.D)
mAU
6.982
0.411
0.524
1.054
6.407
6.737
6.882
7.243
9.649
min

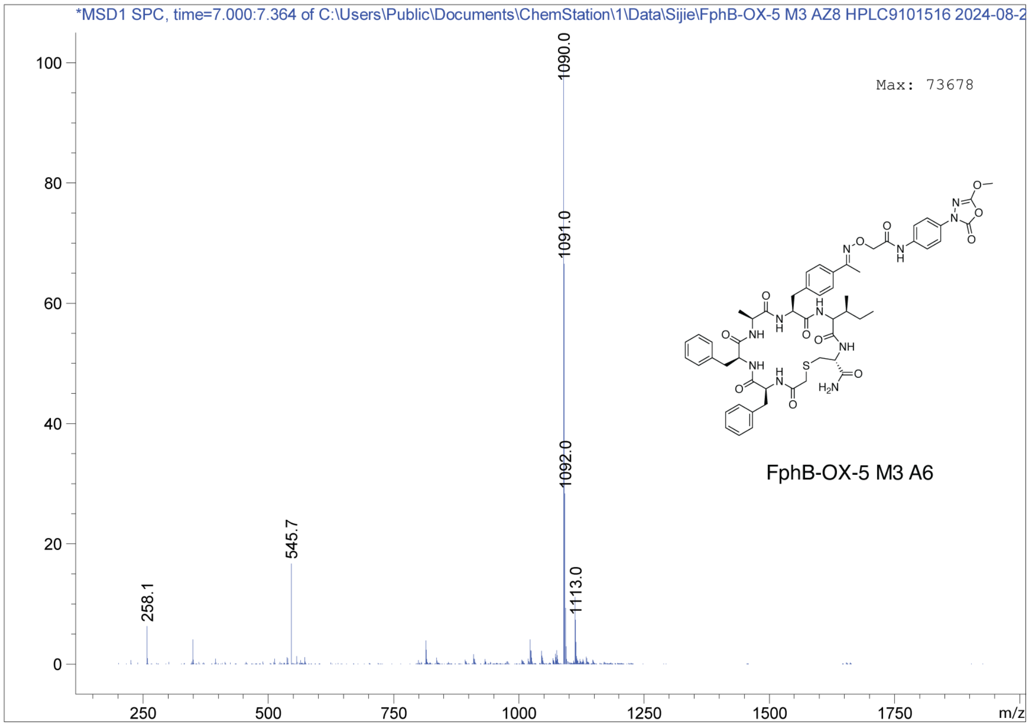
*MSD1 SPC, time=7.000:7.364 of C:\Users\Public\Documents\ChemStation\1\Data\Sijie\FphB-OX-5 M3 AZ8 HPLC9101516 2024-08-2
Max: 73678
1090.0
1091.0
1092.0
545.7
258.1
1113.0
FphB-OX-5 M3 A6
m/z

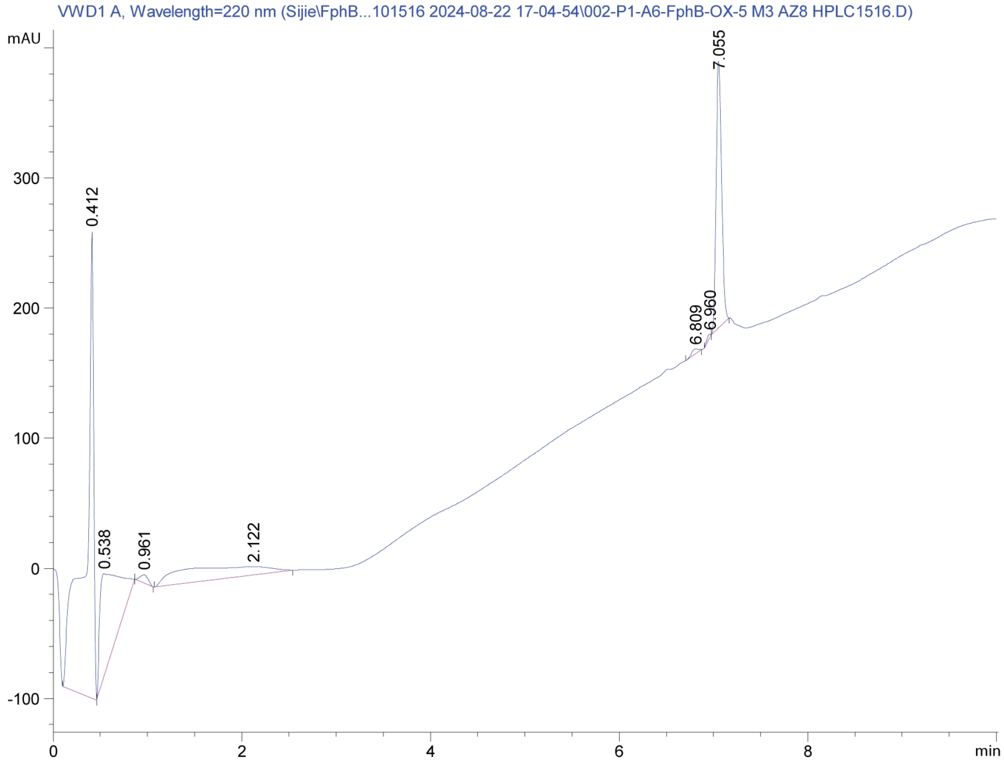
VWD1 A, Wavelength=220 nm (Sijie\FphB...101516 2024-08-22 17-04-54\002-P1-A6-FphB-OX-5 M3 AZ8 HPLC1516.D)
mAU
7.055
0.412
6.809
6.960
0.538
0.961
2.122
min

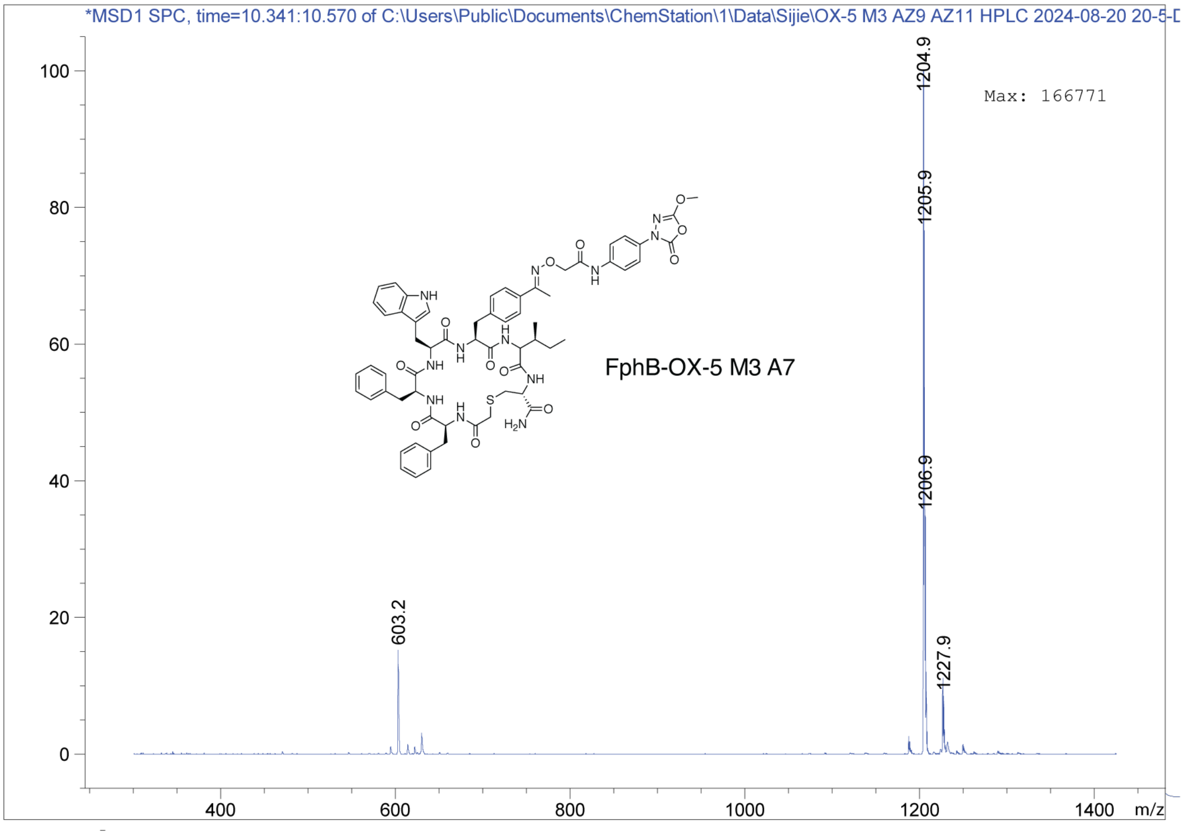
*MSD1 SPC, time=10.341:10.570 of C:\Users\Public\Documents\ChemStation\1\Data\Sijie\OX-5 M3 AZ9 AZ11 HPLC 2024-08-20 20-5
Max: 166771
FphB-OX-5 M3 A7
603.2
1204.9
1205.9
1206.9
1227.9
m/z

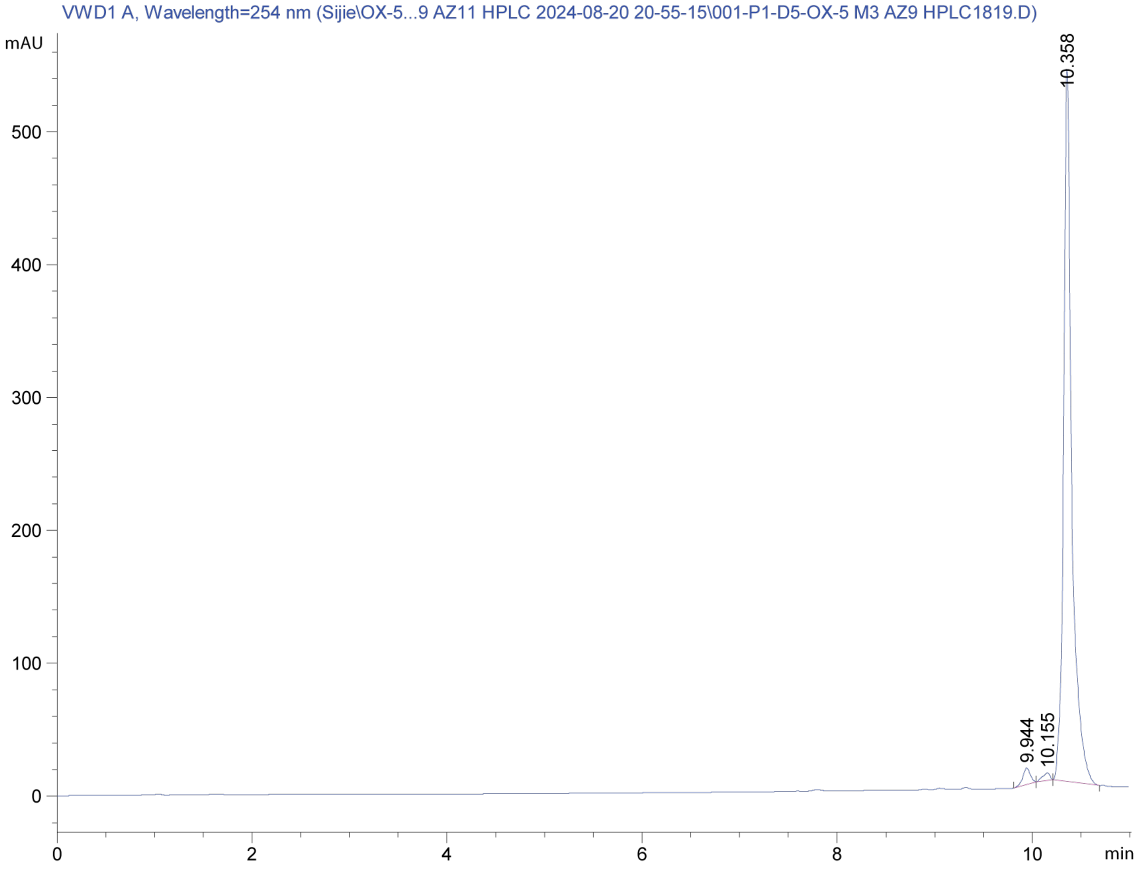
VWD1 A, Wavelength=254 nm (Sijie\OX-5...9 AZ11 HPLC 2024-08-20 20-55-15\001-P1-D5-OX-5 M3 AZ9 HPLC1819.D)
mAU
10.358
9.944
10.155
min

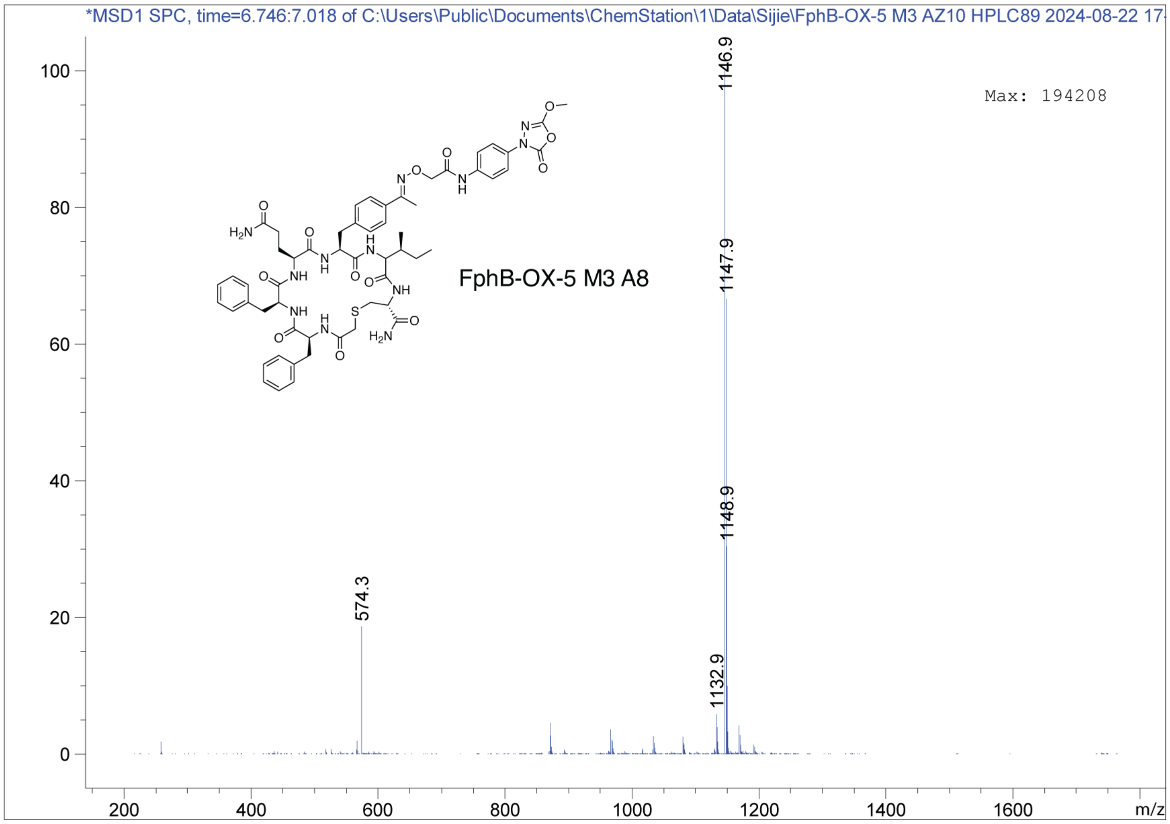
*MSD1 SPC, time=6.746:7.018 of C:\Users\Public\Documents\ChemStation\1\Data\Sijie\FphB-OX-5 M3 AZ10 HPLC89 2024-08-22 17-
Max: 194208
FphB-OX-5 M3 A8
1146.9
1147.9
1148.9
574.3
1132.9
m/z

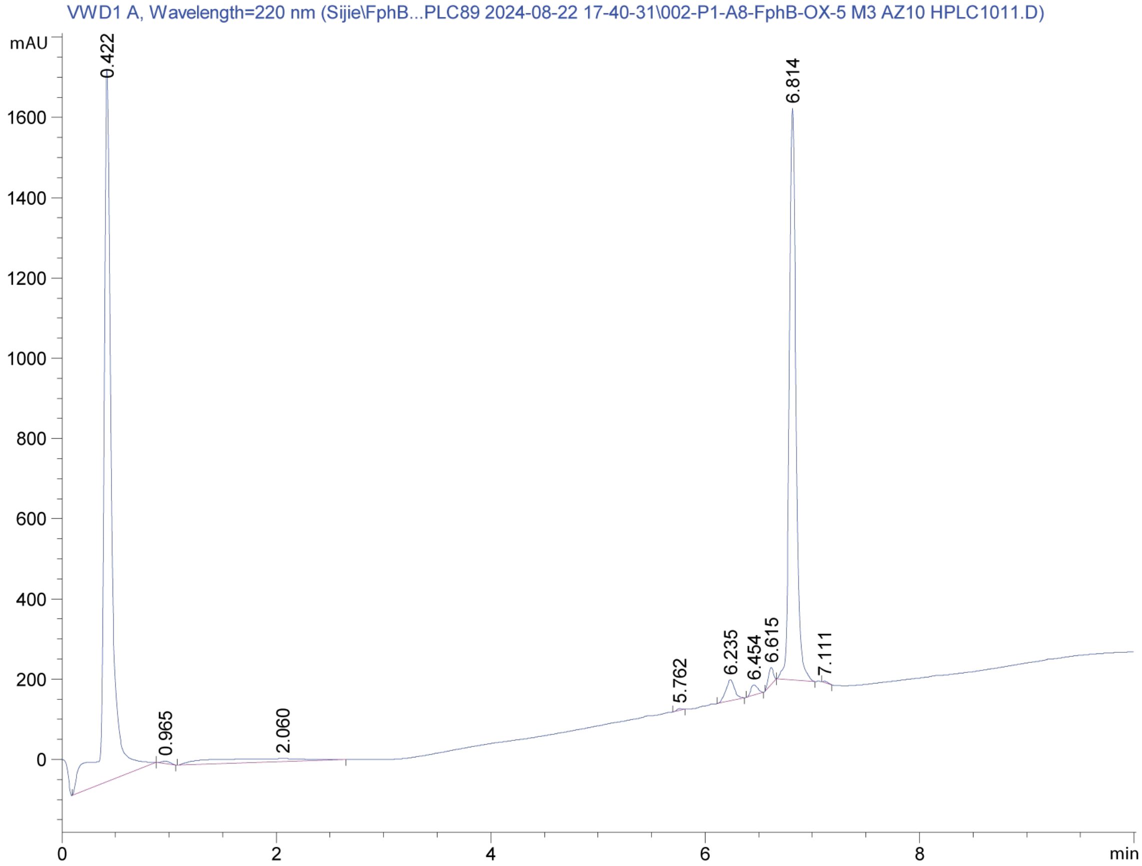
VWD1 A, Wavelength=220 nm (Sijie\FphB...PLC89 2024-08-22 17-40-31\002-P1-A8-FphB-OX-5 M3 AZ10 HPLC1011.D)
mAU
0.422
6.814
0.965
2.060
5.762
6.235
6.454
6.615
7.111
min

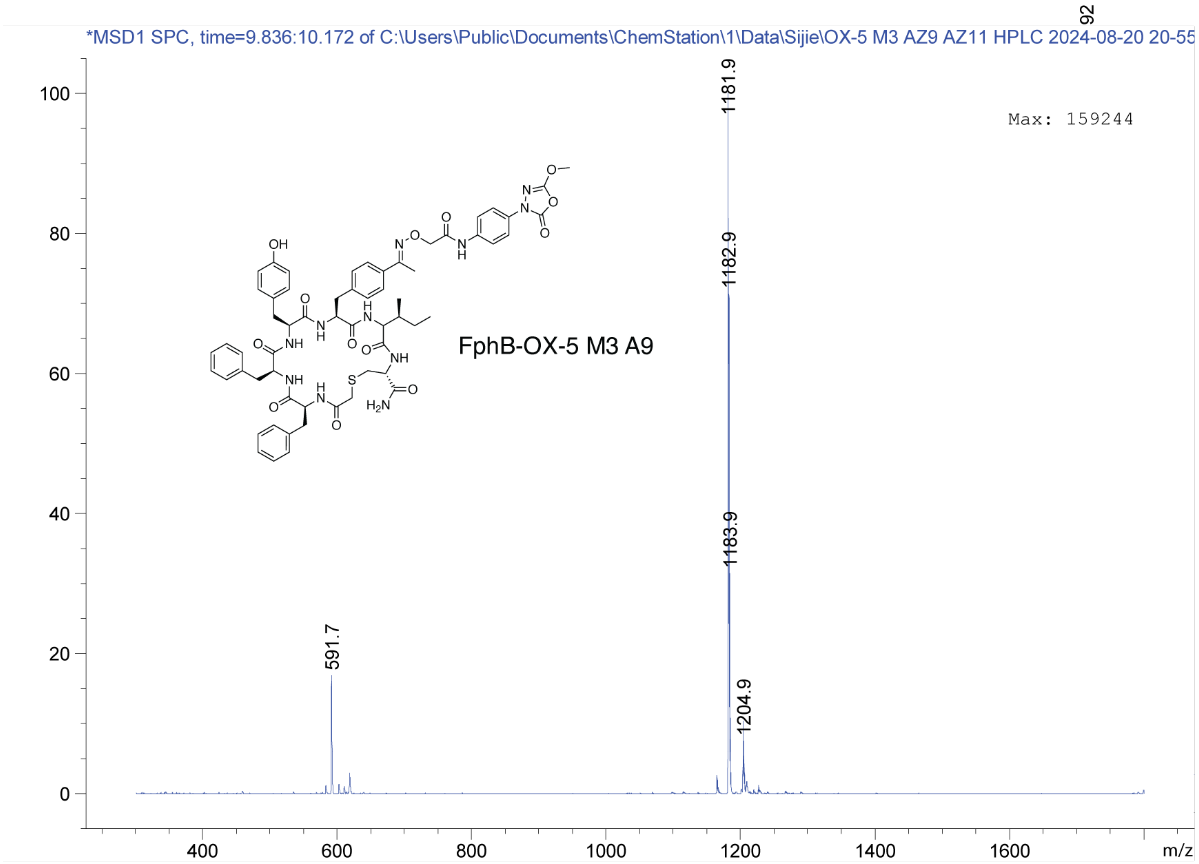
*MSD1 SPC, time=9.836:10.172 of C:\Users\Public\Documents\ChemStation\1\Data\Sijie\OX-5 M3 AZ9 AZ11 HPLC 2024-08-20 20-55
92
Max: 159244
1181.9
1182.9
1183.9
591.7
1204.9
FphB-OX-5 M3 A9
m/z

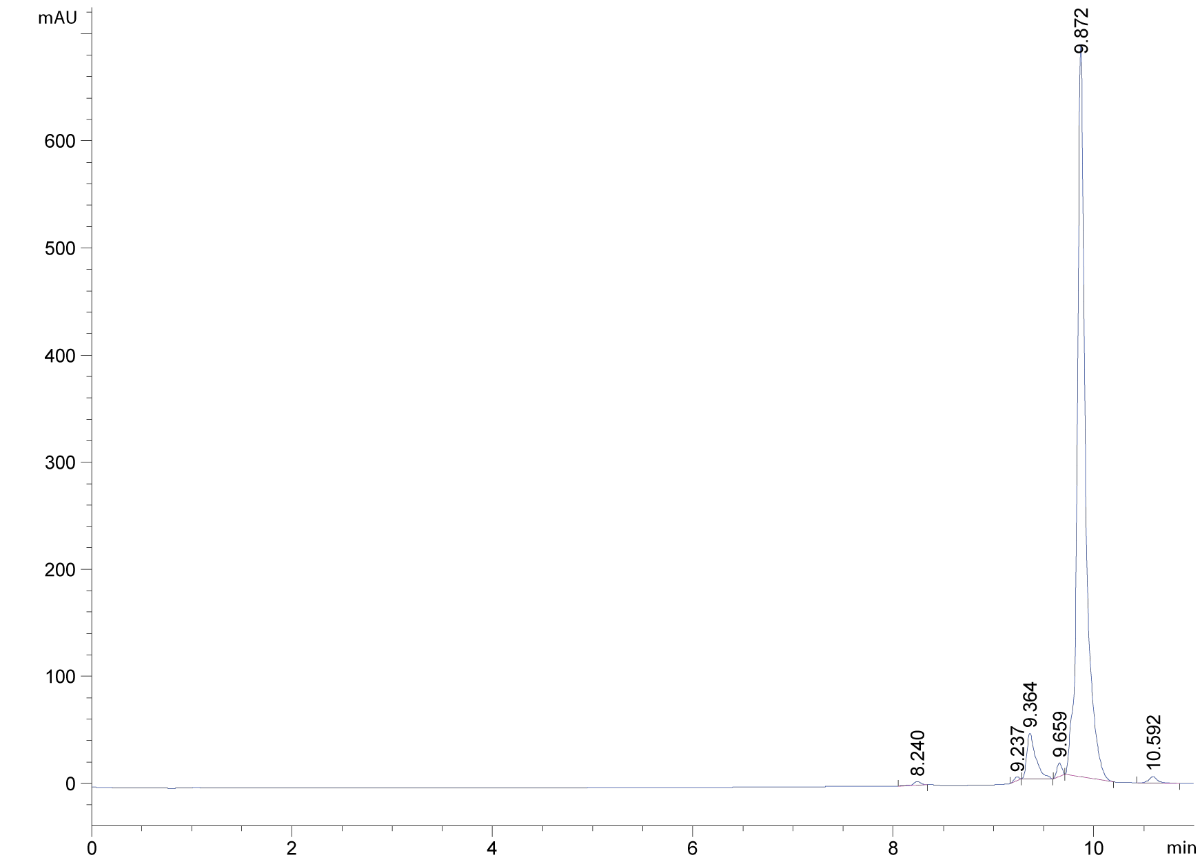
mAU
9.872
8.240
9.237
9.364
9.659
10.592
min

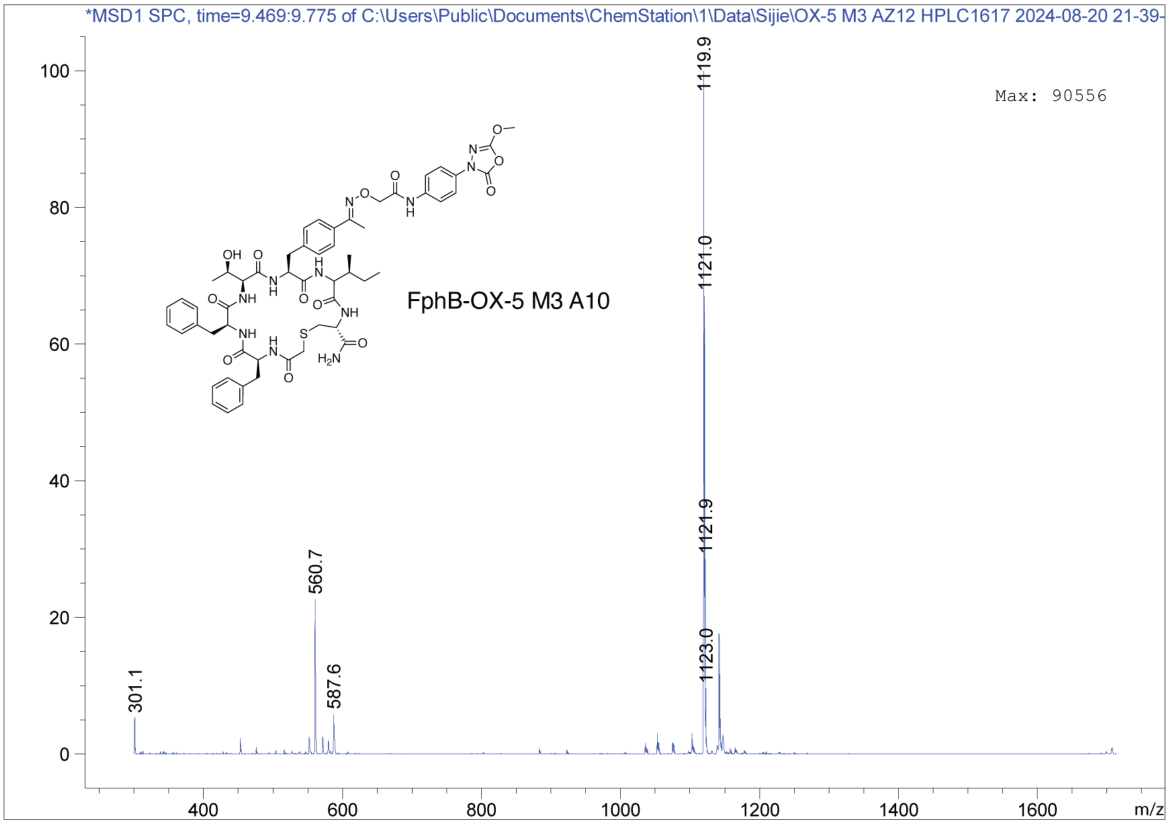
*MSD1 SPC, time=9.469:9.775 of C:\Users\Public\Documents\ChemStation\1\Data\Sijie\OX-5 M3 AZ12 HPLC1617 2024-08-20 21-39-
Max: 90556
FphB-OX-5 M3 A10
1119.9
1121.0
1121.9
1123.0
560.7
587.6
301.1
m/z

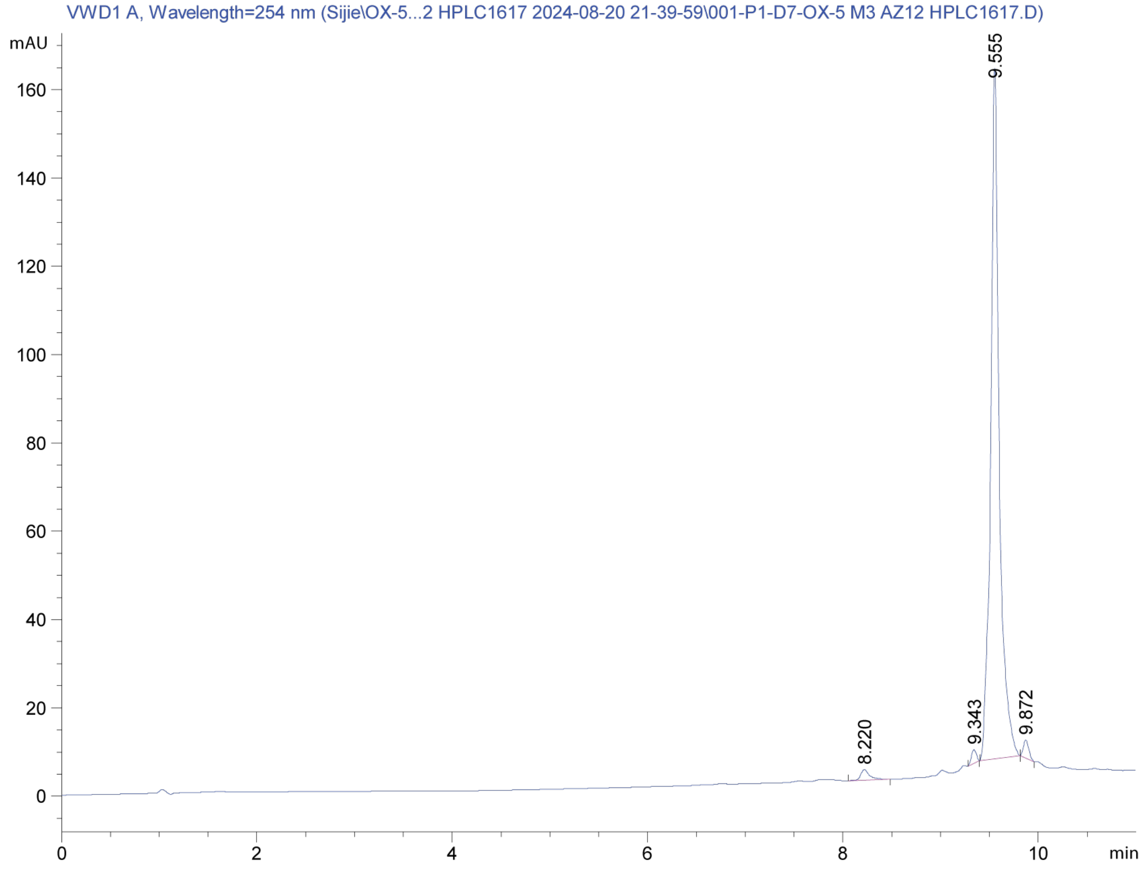
VWD1 A, Wavelength=254 nm (Sijie\OX-5...2 HPLC1617 2024-08-20 21-39-59\001-P1-D7-OX-5 M3 AZ12 HPLC1617.D)
mAU
9.555
8.220
9.343
9.872
min

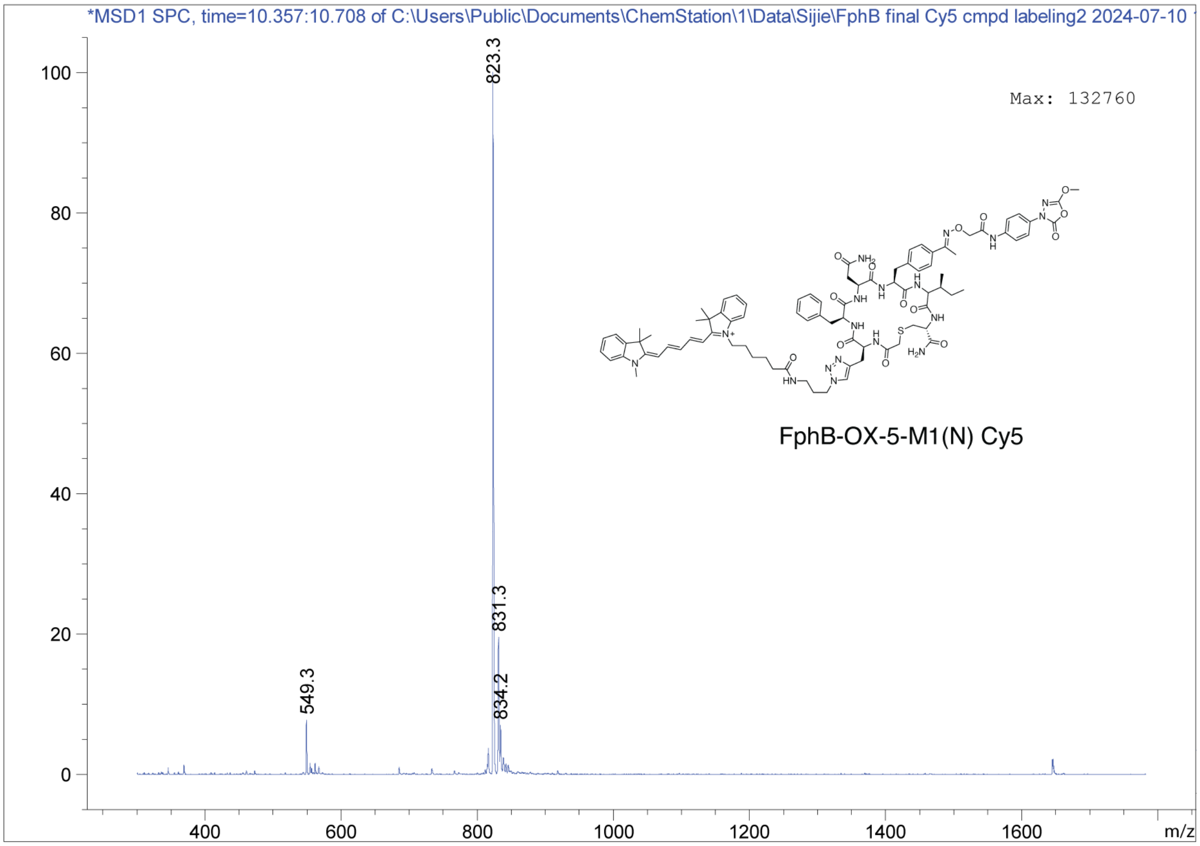
*MSD1 SPC, time=10.357:10.708 of C:\Users\Public\Documents\ChemStation\1\Data\Sijie\FphB final Cy5 cmpd labeling2 2024-07-10
Max: 132760
823.3
831.3
834.2
549.3
FphB-OX-5-M1(N) Cy5
0
20
40
60
80
100
400
600
800
1000
1200
1400
1600
m/z

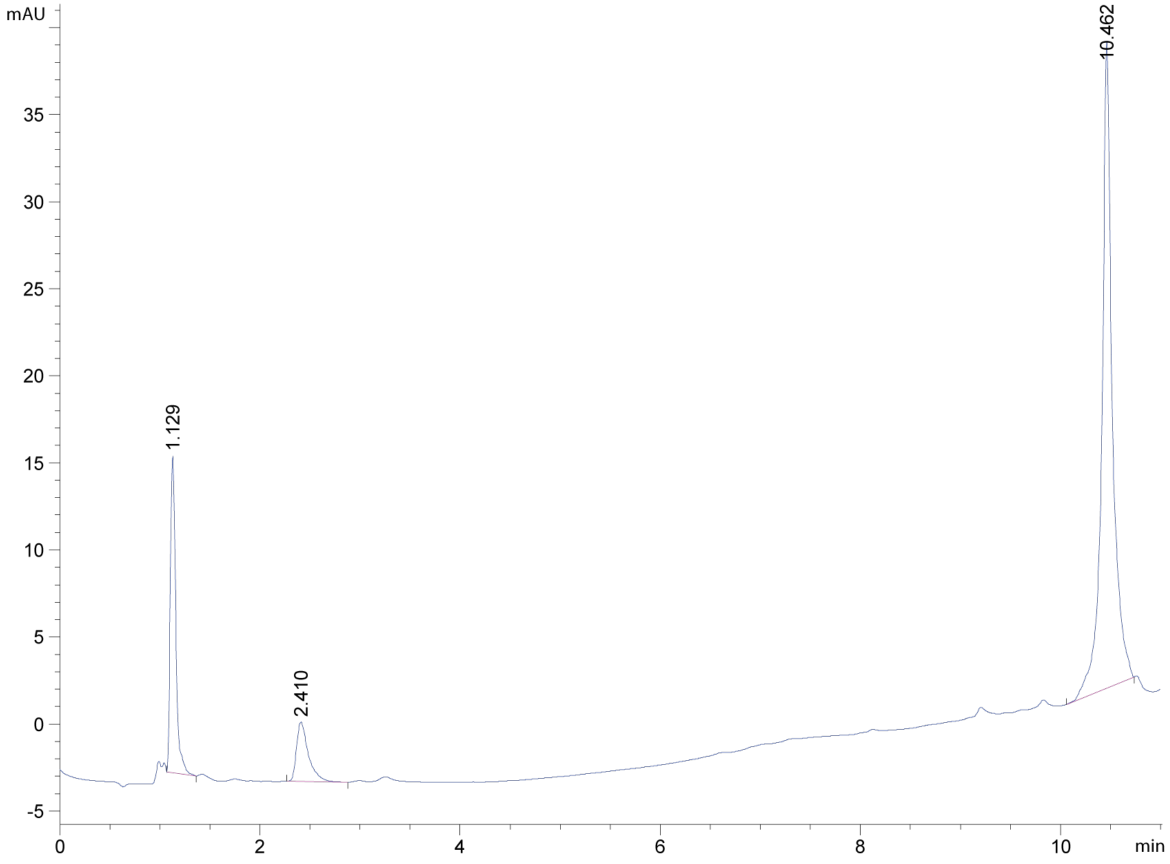
mAU
10.462
1.129
2.410
-5
0
5
10
15
20
25
30
35
0
2
4
6
8
10
min

MS Spectrum

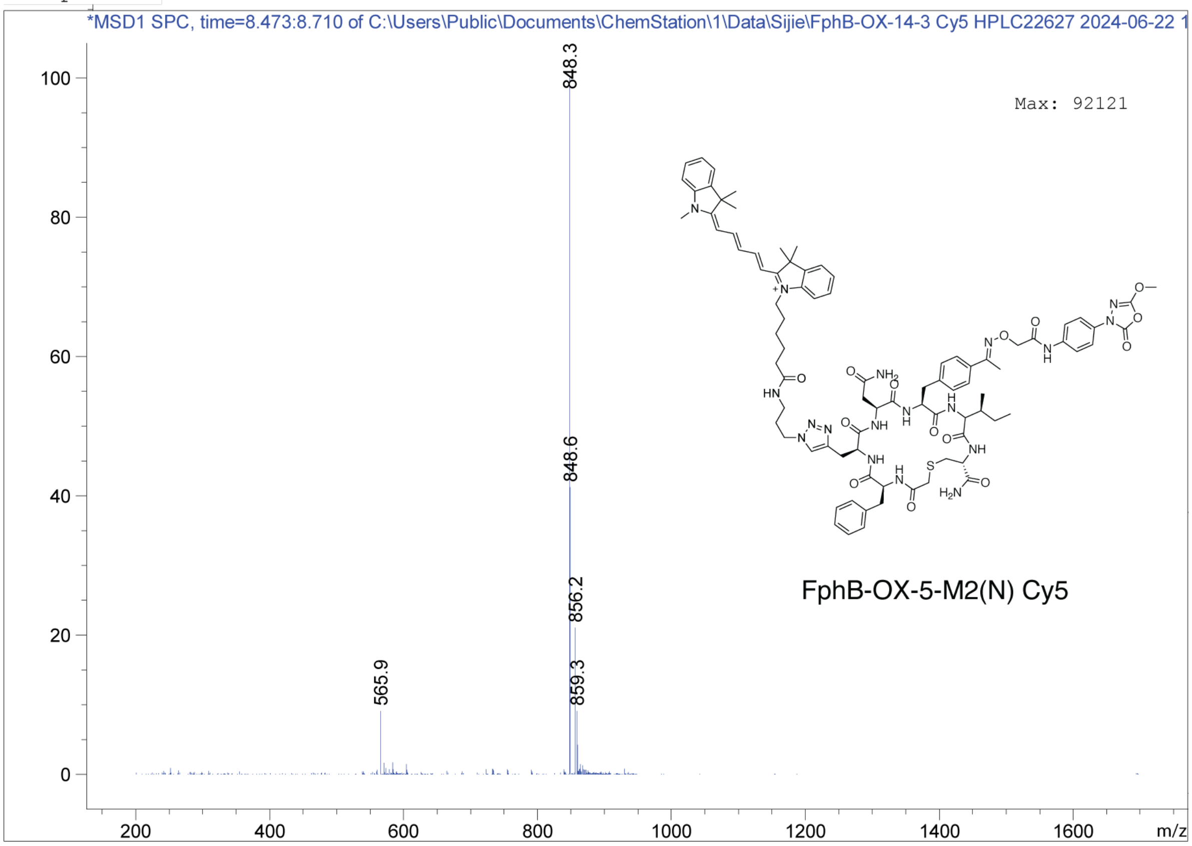
*MSD1 SPC, time=8.473:8.710 of C:\Users\Public\Documents\ChemStation\1\Data\Sijie\FphB-OX-14-3 Cy5 HPLC22627 2024-06-22 1
Max: 92121
848.3
848.6
856.2
859.3
565.9
FphB-OX-5-M2(N) Cy5
m/z

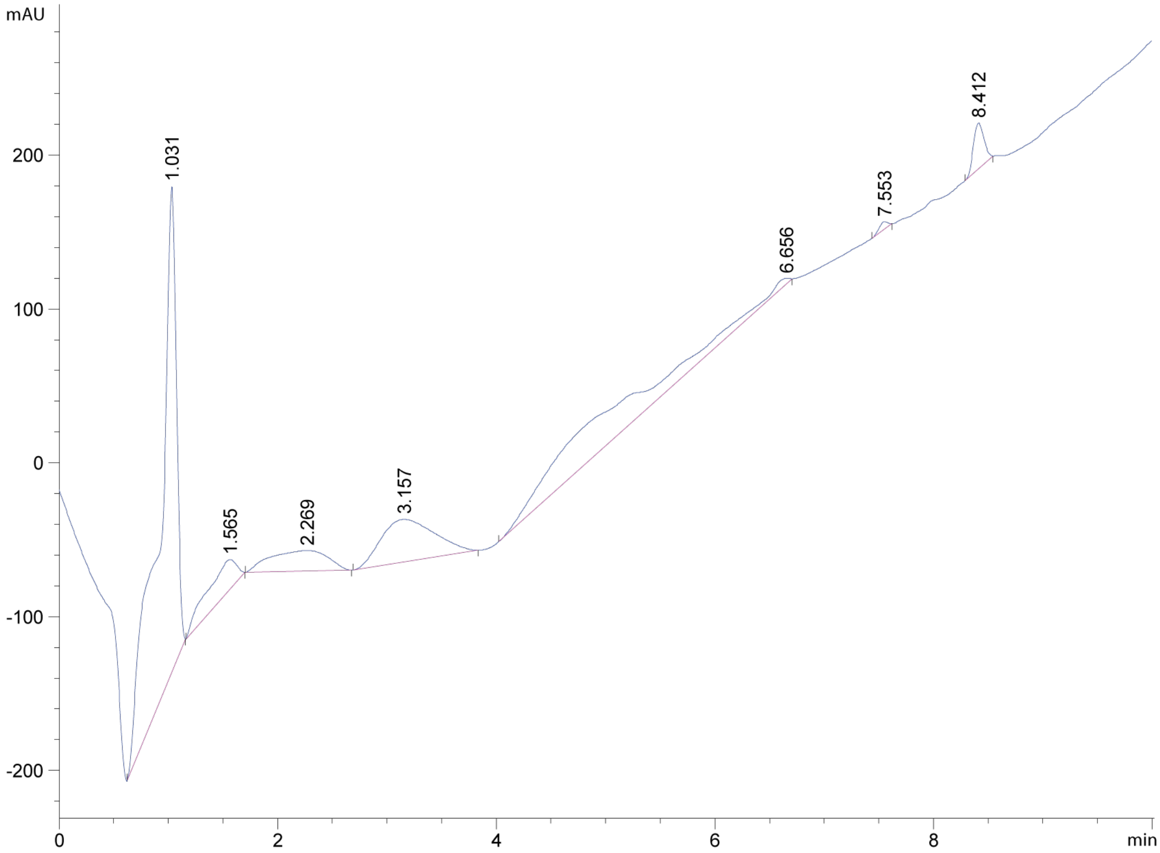
mAU
1.031
1.565
2.269
3.157
6.656
7.553
8.412
min

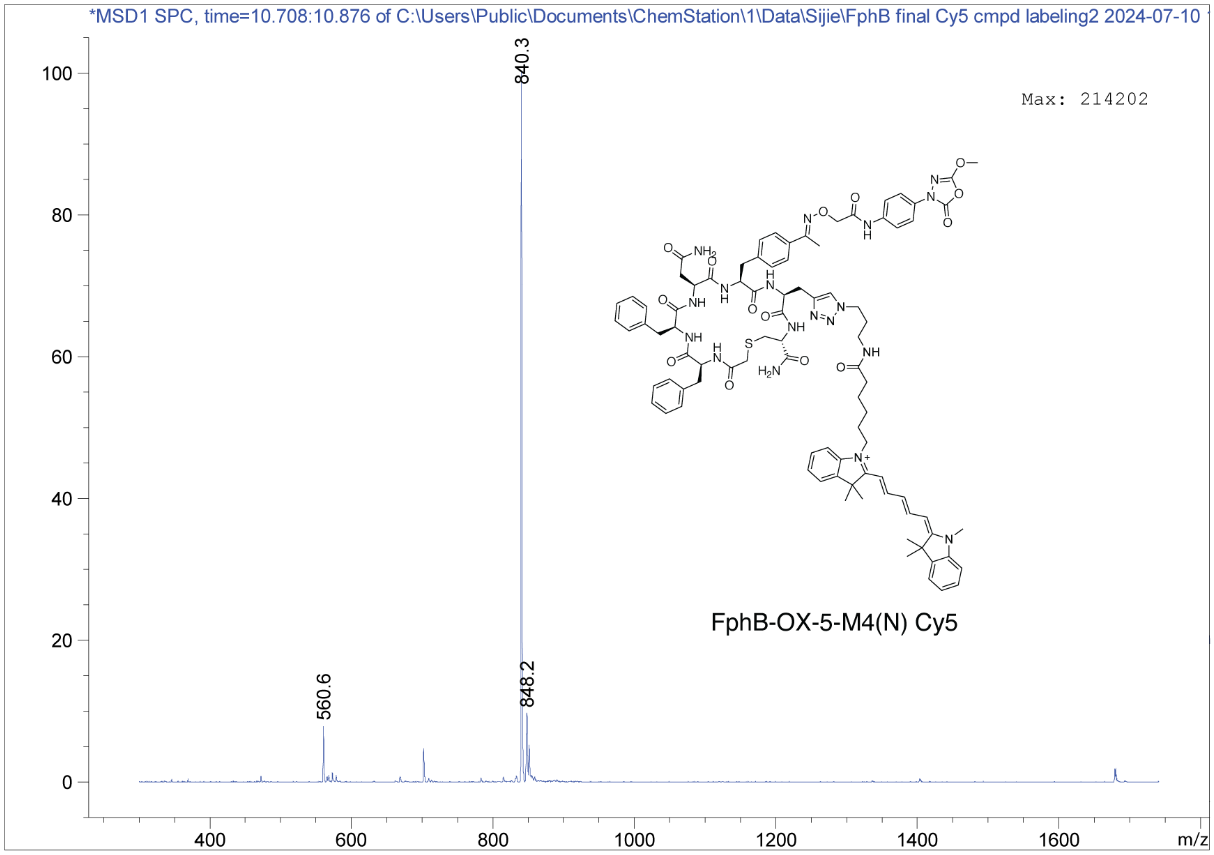
*MSD1 SPC, time=10.708:10.876 of C:\Users\Public\Documents\ChemStation\1\Data\Sijie\FphB final Cy5 cmpd labeling2 2024-07-10
Max: 214202
840.3
848.2
560.6
FphB-OX-5-M4(N) Cy5
m/z

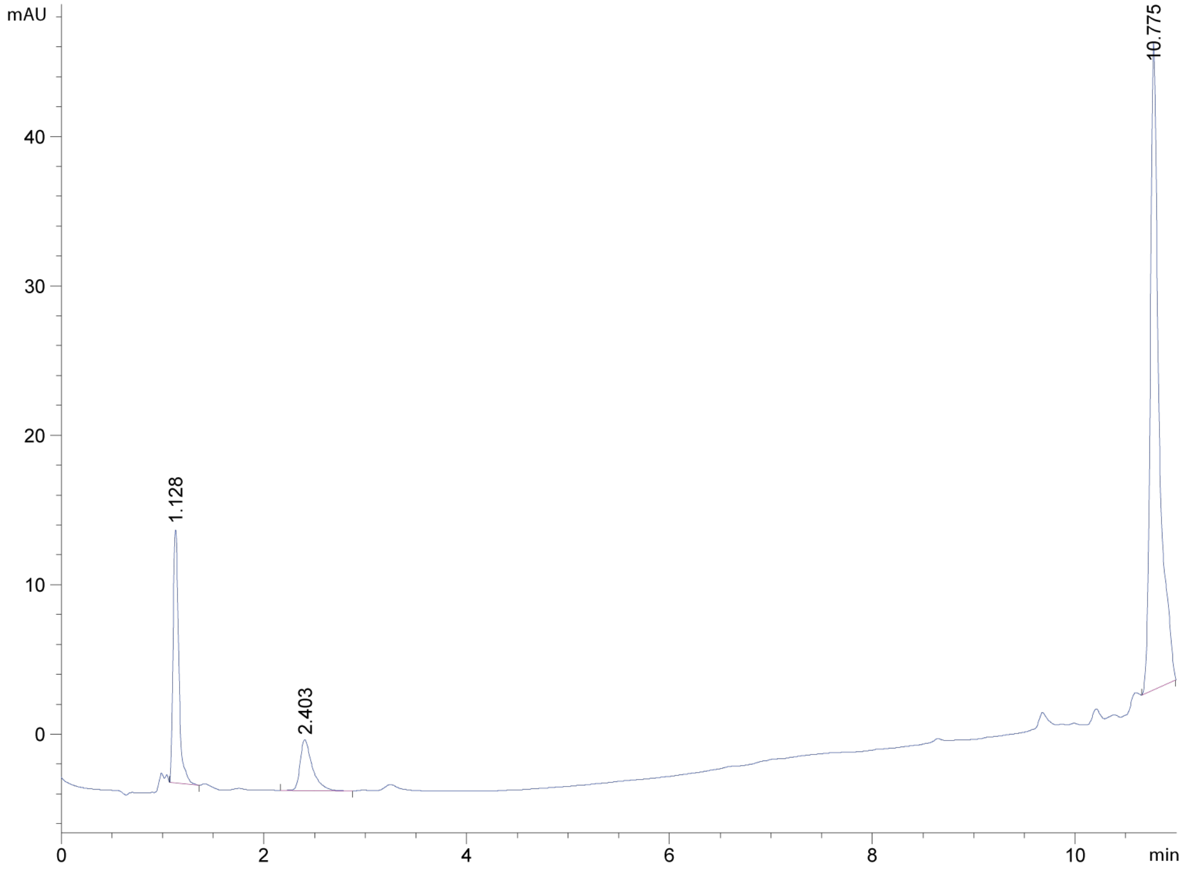
mAU
10.775
1.128
2.403
min

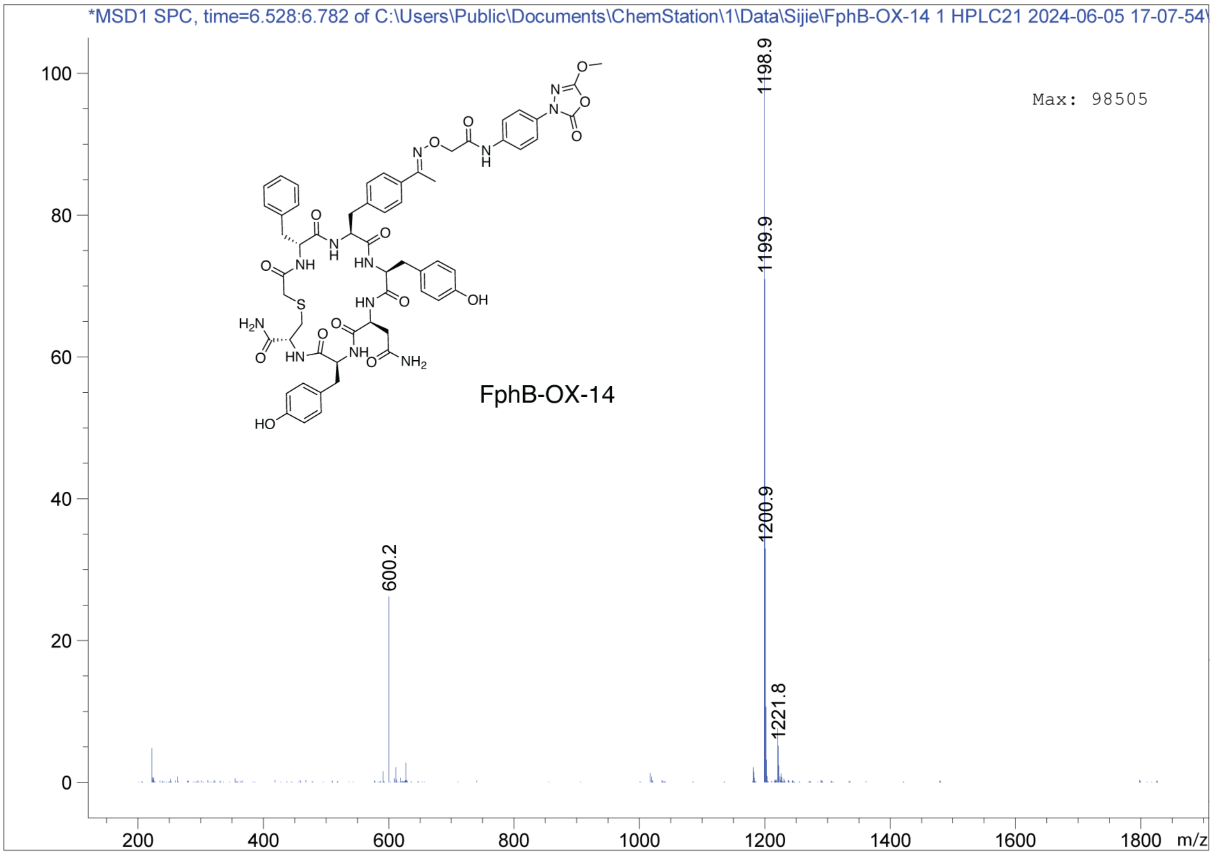
*MSD1 SPC, time=6.528:6.782 of C:\Users\Public\Documents\ChemStation\1\Data\Sijie\FphB-OX-14 1 HPLC21 2024-06-05 17-07-54\
Max: 98505
FphB-OX-14
1198.9
1199.9
1200.9
600.2
1221.8
m/z

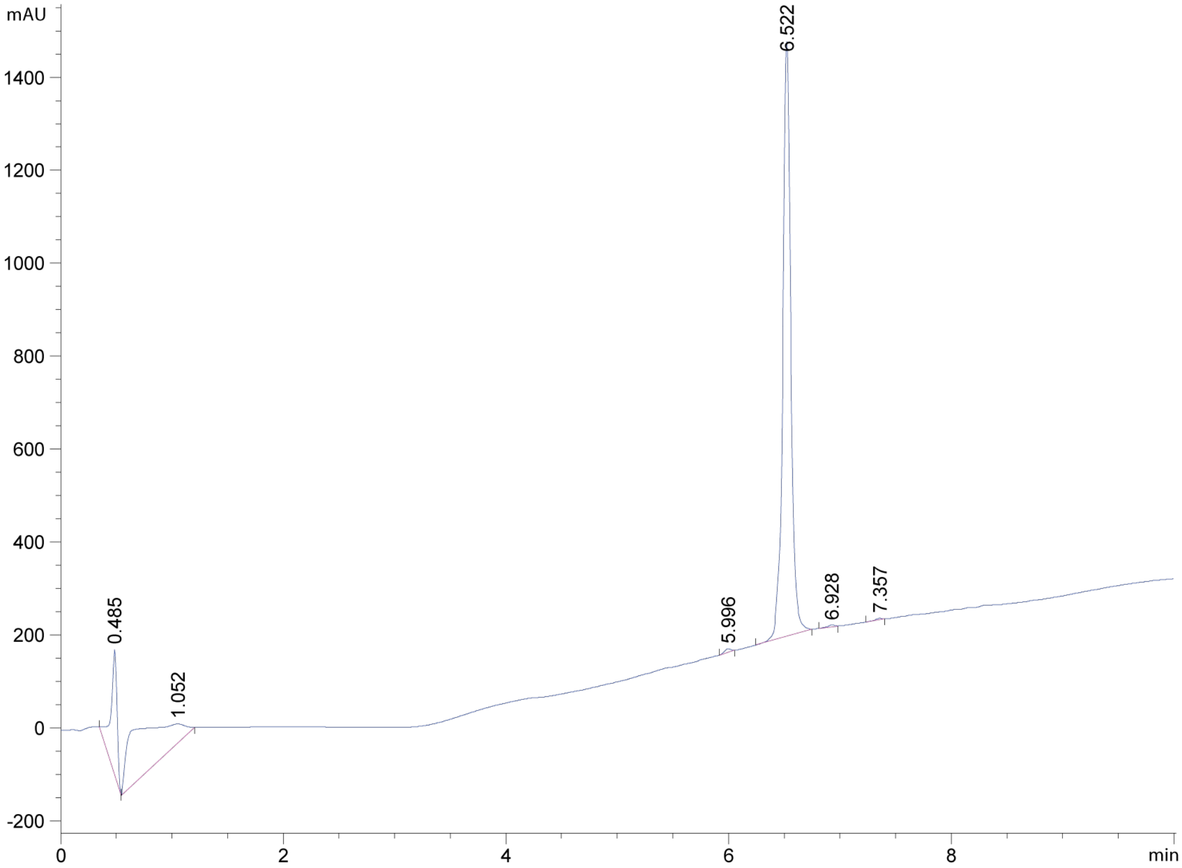
mAU
6.522
0.485
1.052
5.996
6.928
7.357
min

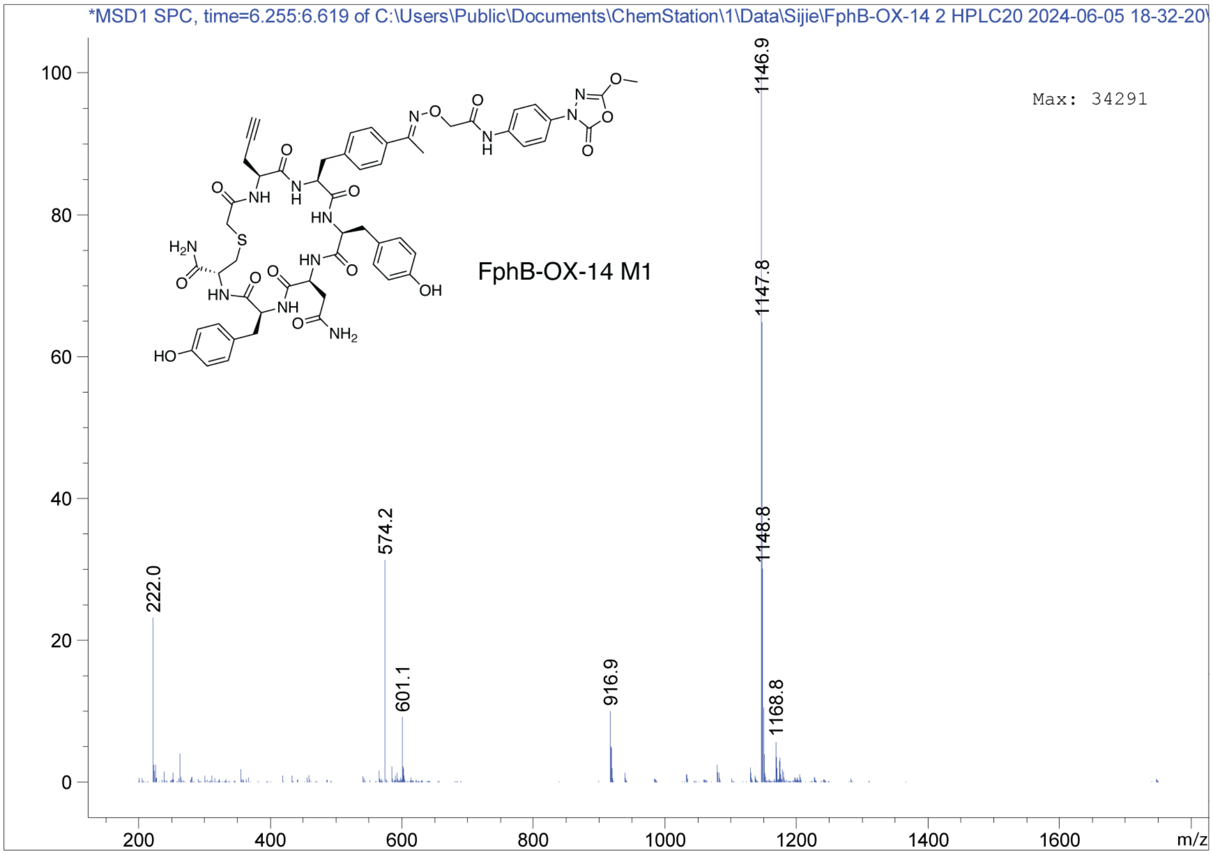
*MSD1 SPC, time=6.255:6.619 of C:\Users\Public\Documents\ChemStation\1\Data\Sijie\FphB-OX-14 2 HPLC20 2024-06-05 18-32-20\
Max: 34291
FphB-OX-14 M1
1146.9
1147.8
1148.8
574.2
222.0
601.1
916.9
1168.8
m/z

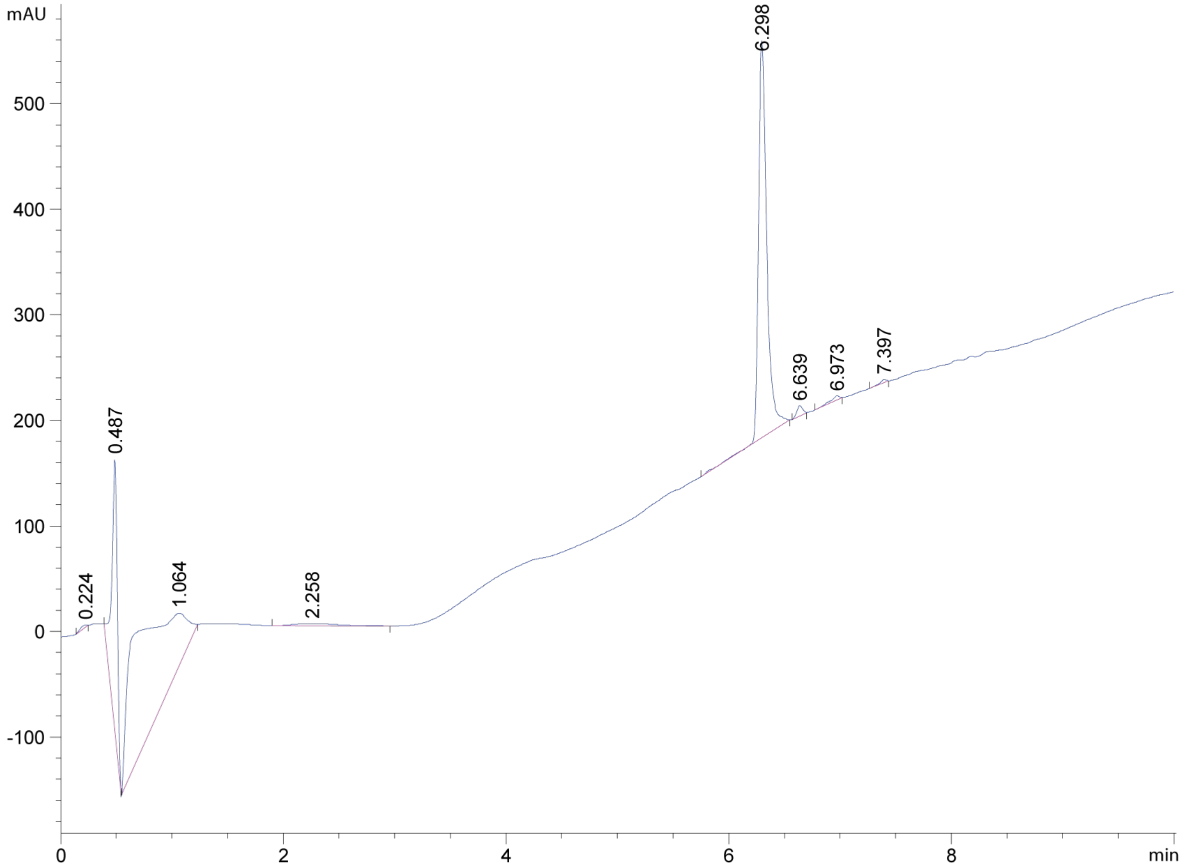
mAU
6.298
0.487
0.224
1.064
2.258
6.639
6.973
7.397
min

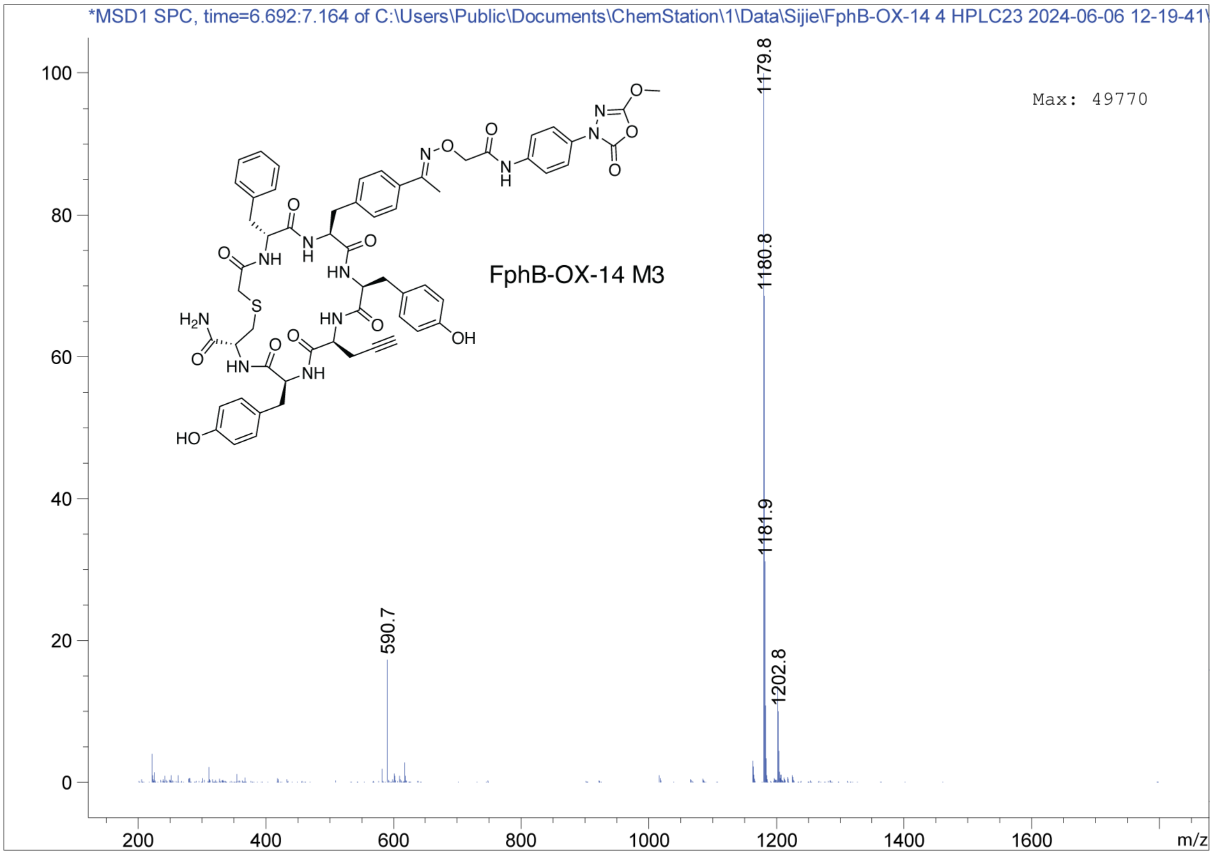
*MSD1 SPC, time=6.692:7.164 of C:\Users\Public\Documents\ChemStation\1\Data\Sijie\FphB-OX-14 4 HPLC23 2024-06-06 12-19-41\
Max: 49770
FphB-OX-14 M3
1179.8
1180.8
1181.9
1202.8
590.7
m/z

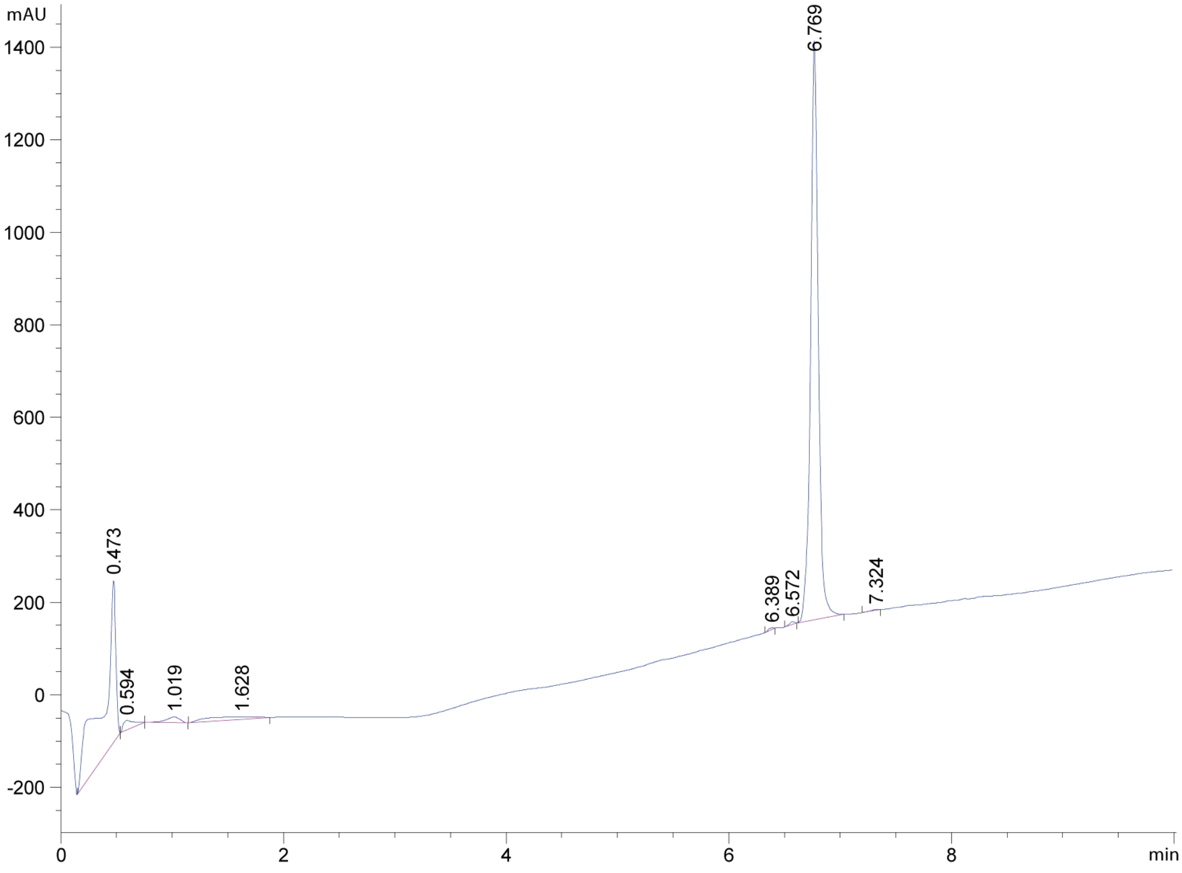
mAU
6.769
0.473
0.594
1.019
1.628
6.389
6.572
7.324
min

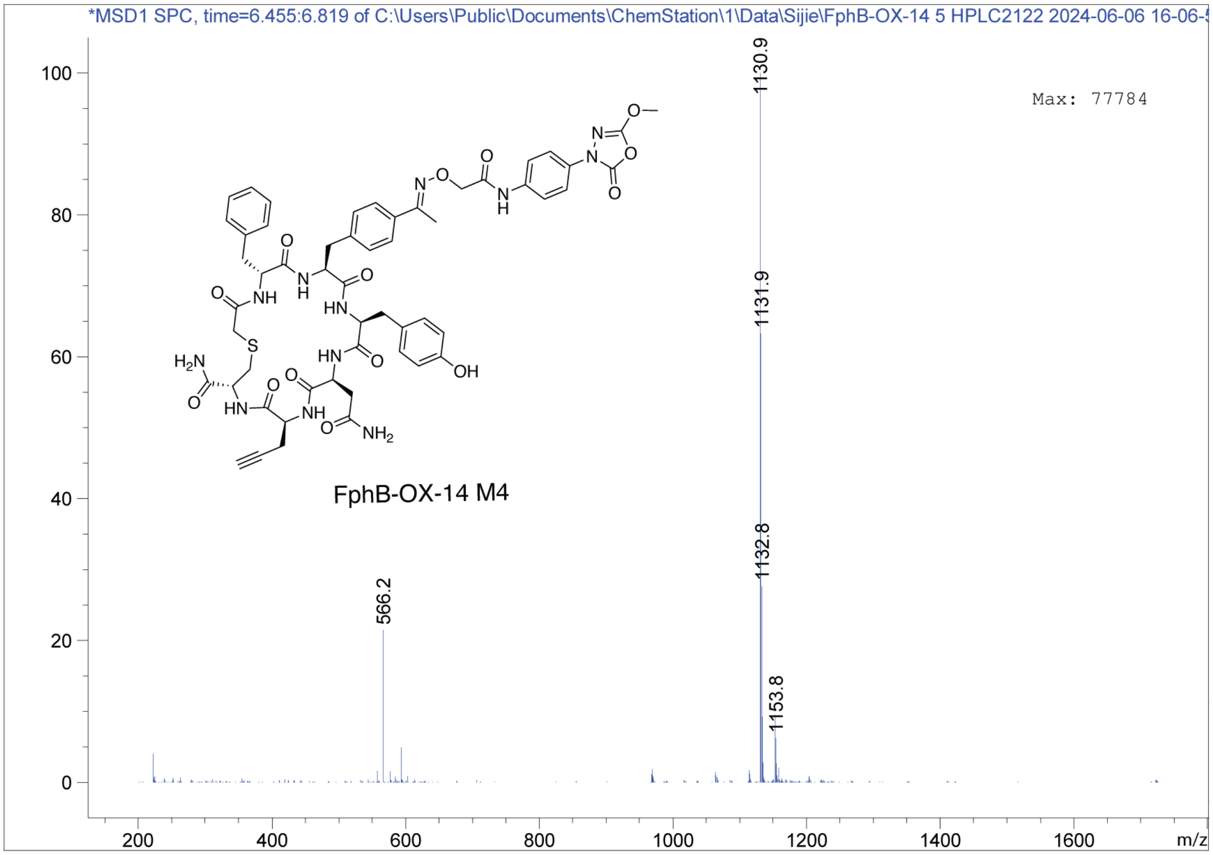
*MSD1 SPC, time=6.455:6.819 of C:\Users\Public\Documents\ChemStation\1\Data\Sijie\FphB-OX-14 5 HPLC2122 2024-06-06 16-06-
Max: 77784
FphB-OX-14 M4
1130.9
1131.9
1132.8
566.2
1153.8
m/z

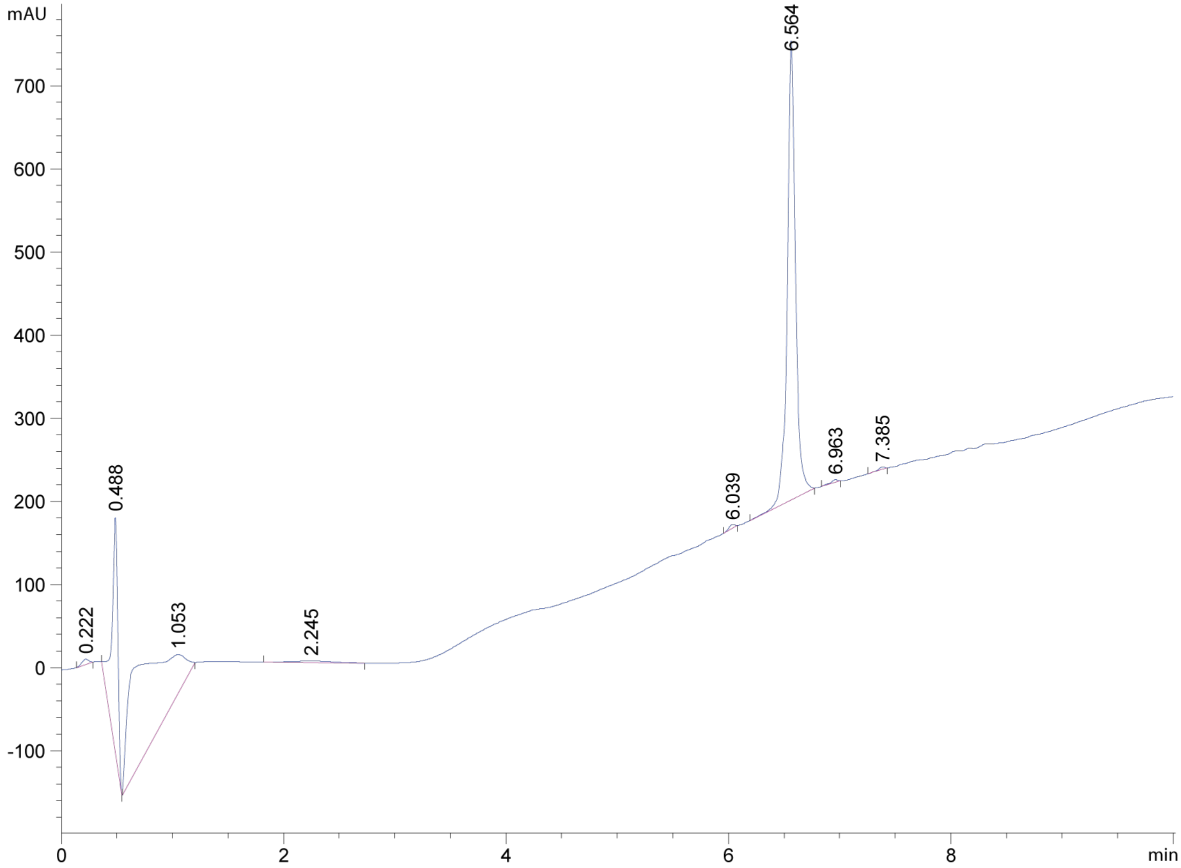
mAU
6.564
0.488
0.222
1.053
2.245
6.039
6.963
7.385
min

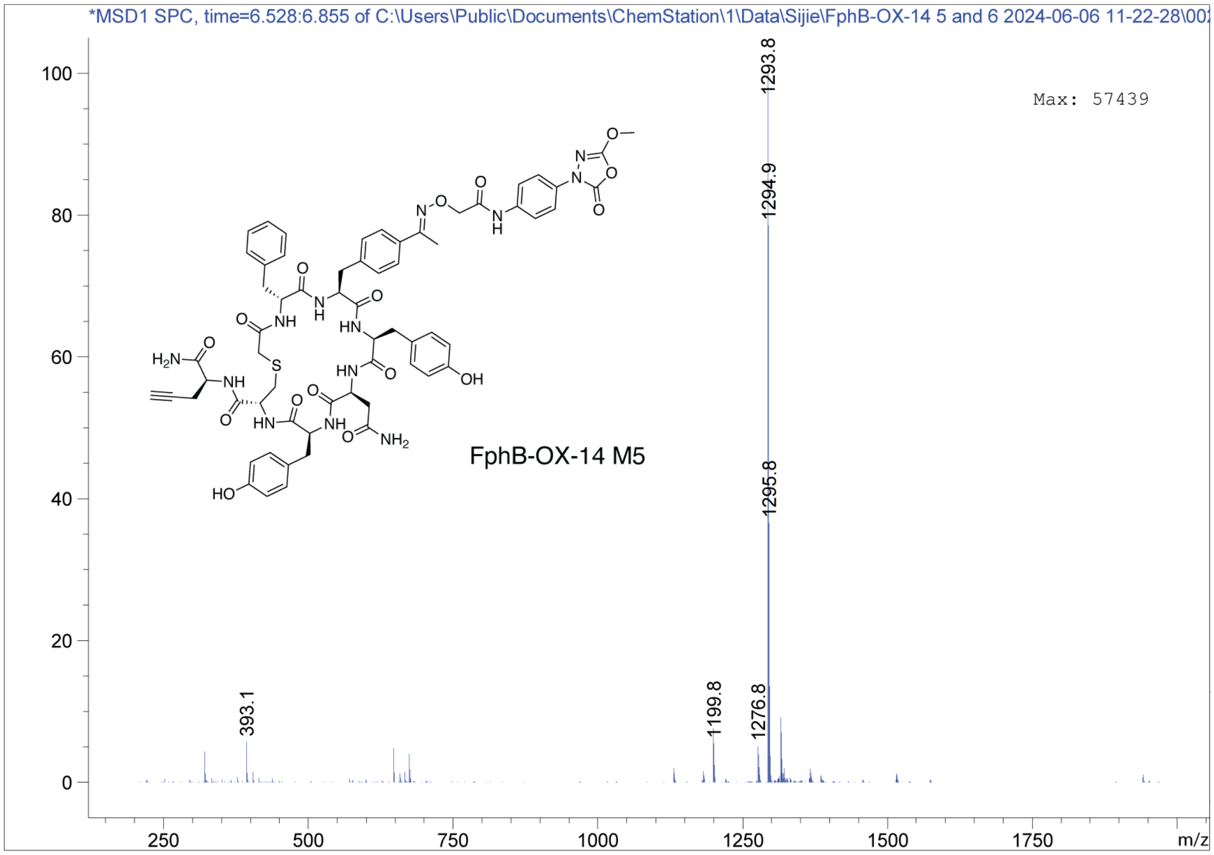
*MSD1 SPC, time=6.528:6.855 of C:\Users\Public\Documents\ChemStation\1\Data\Sijie\FphB-OX-14 5 and 6 2024-06-06 11-22-28\00
Max: 57439
FphB-OX-14 M5
1293.8
1294.9
1295.8
1276.8
1199.8
393.1
m/z

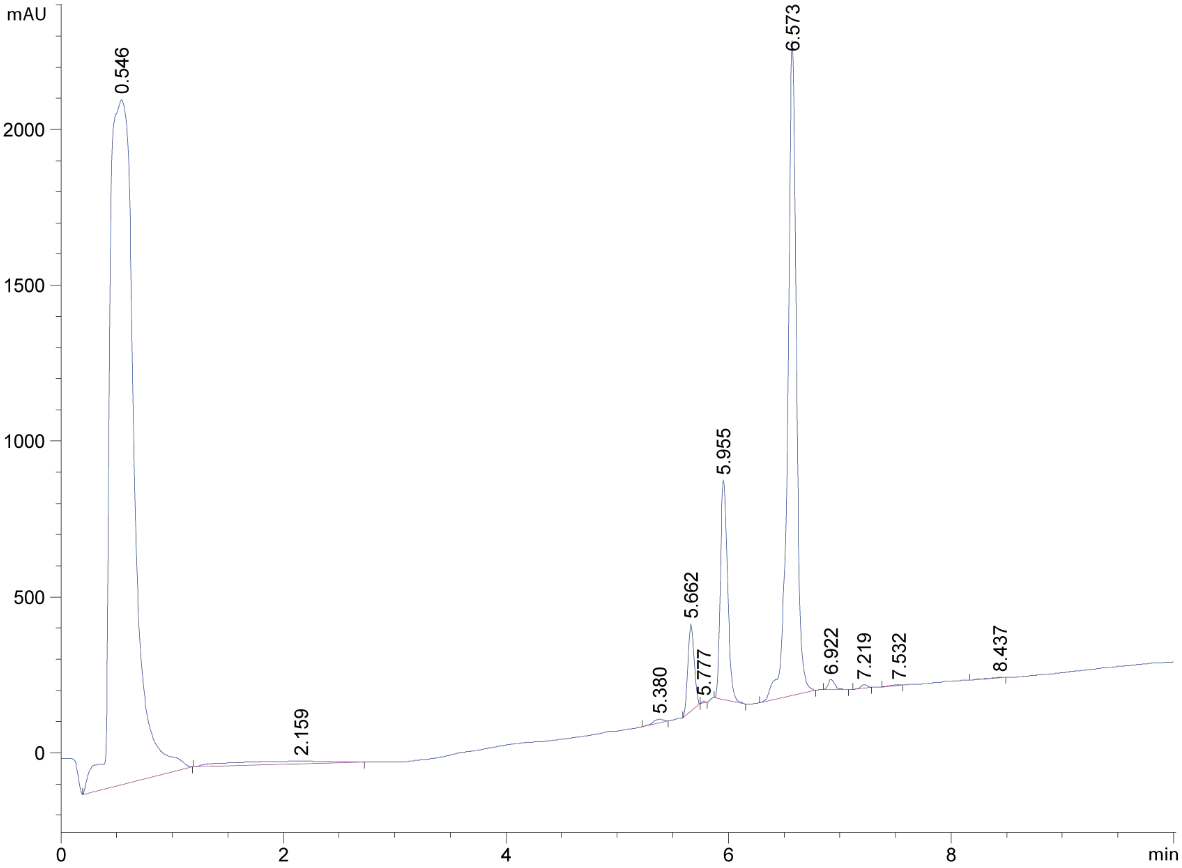
mAU
0.546
6.573
5.955
5.662
5.777
5.380
2.159
6.922
7.219
7.532
8.437
min

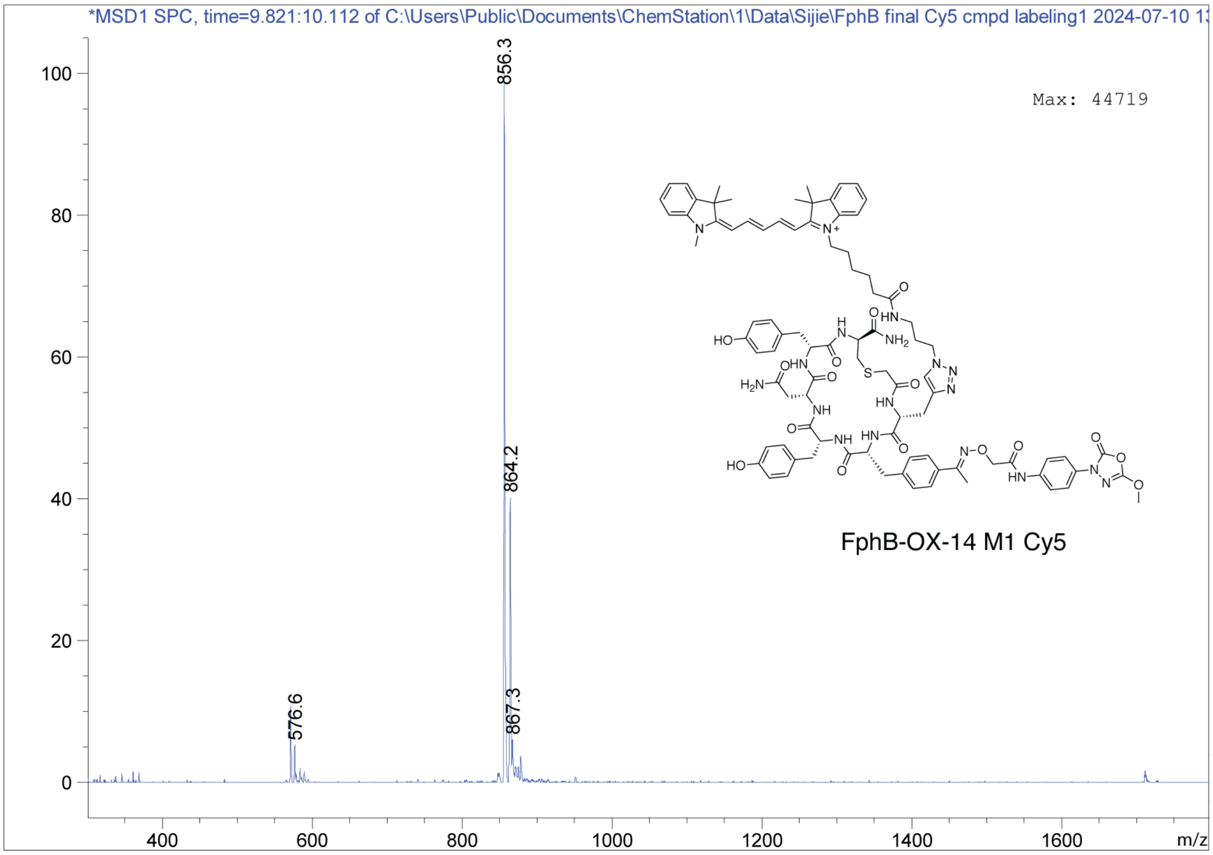
*MSD1 SPC, time=9.821:10.112 of C:\Users\Public\Documents\ChemStation\1\Data\Sijie\FphB final Cy5 cmpd labeling1 2024-07-10 1
Max: 44719
856.3
864.2
867.3
576.6
FphB-OX-14 M1 Cy5
m/z

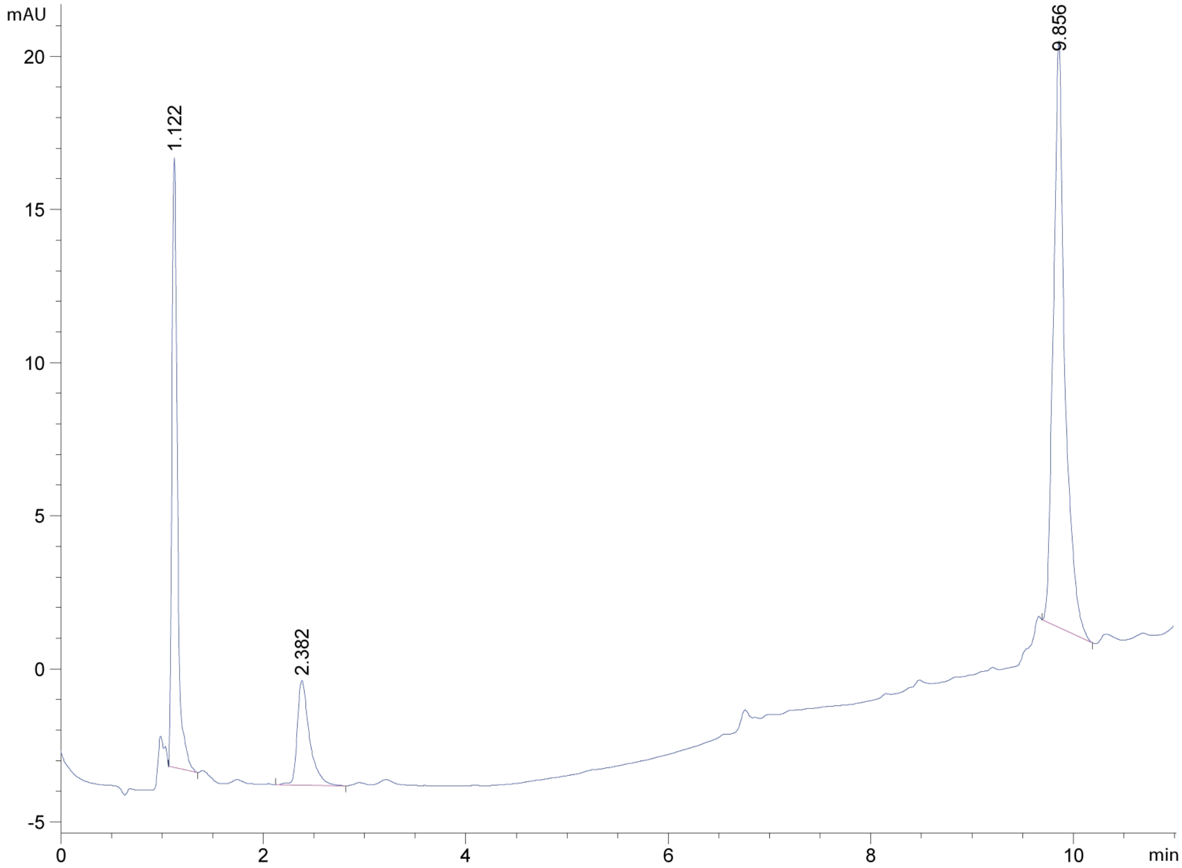
mAU
1.122
2.382
9.856
min

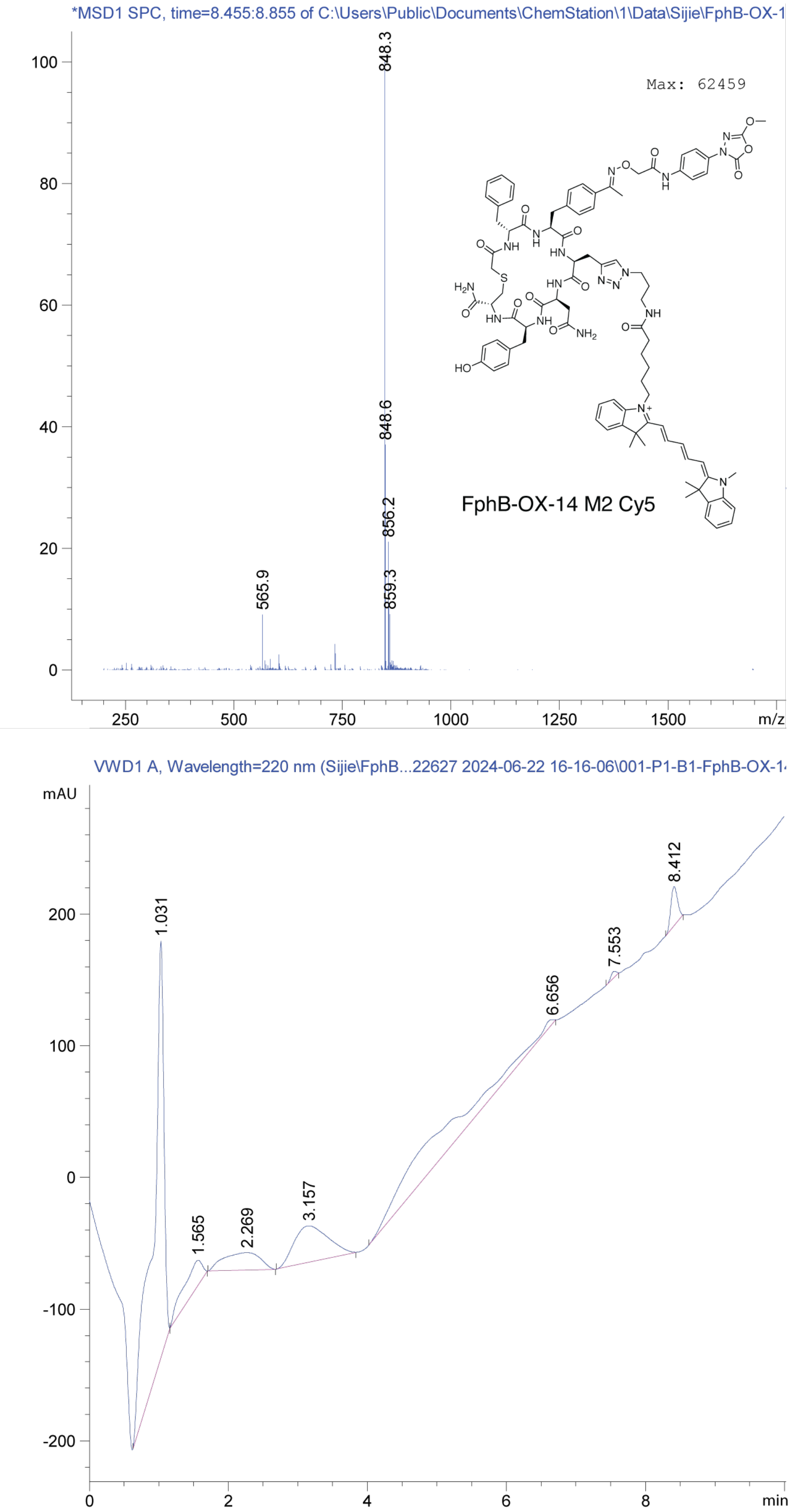
*MSD1 SPC, time=8.455:8.855 of C:\Users\Public\Documents\ChemStation\1\Data\Sijie\FphB-OX-1
Max: 62459
848.3
848.6
856.2
859.3
565.9
FphB-OX-14 M2 Cy5
m/z
VWD1 A, Wavelength=220 nm (Sijie\FphB...22627 2024-06-22 16-16-06\001-P1-B1-FphB-OX-1
mAU
1.031
1.565
2.269
3.157
6.656
7.553
8.412
min

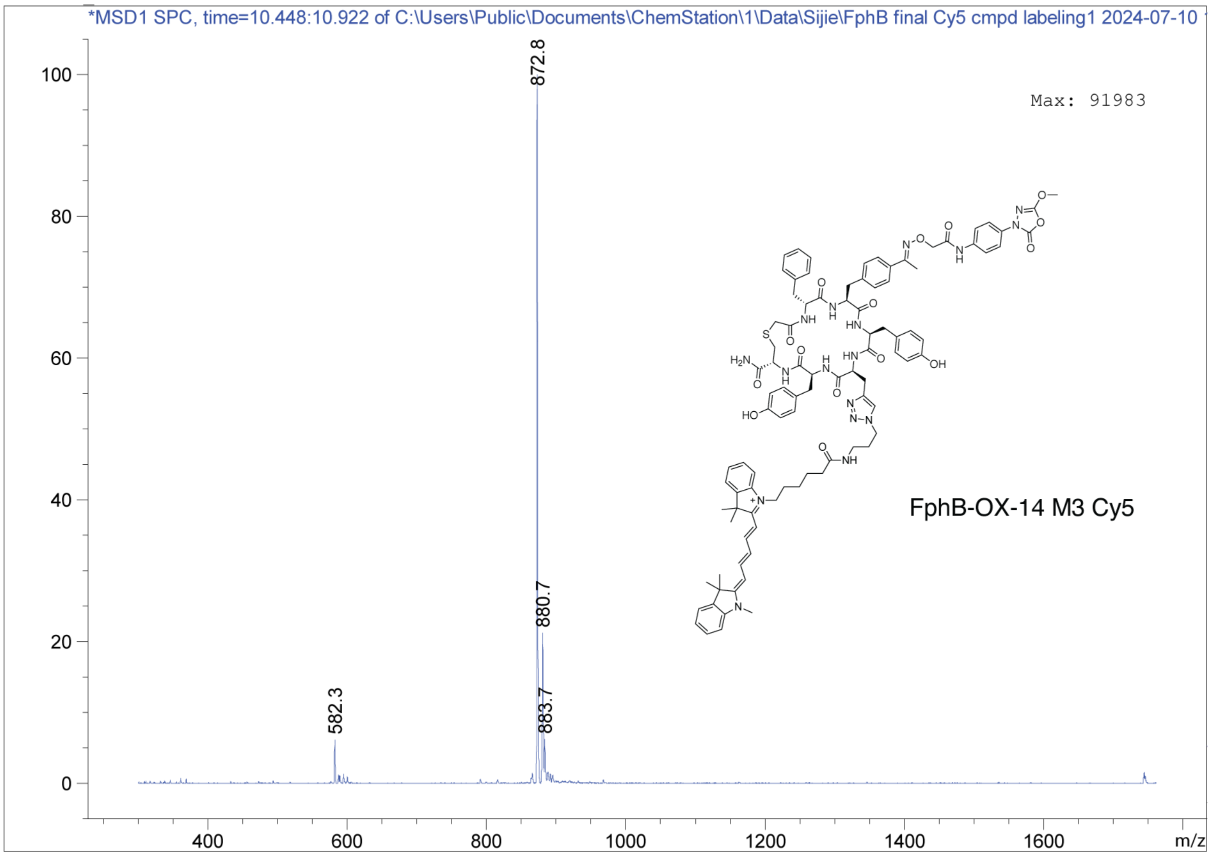
*MSD1 SPC, time=10.448:10.922 of C:\Users\Public\Documents\ChemStation\1\Data\Sijie\FphB final Cy5 cmpd labeling1 2024-07-10
Max: 91983
872.8
880.7
883.7
582.3
FphB-OX-14 M3 Cy5
0
20
40
60
80
100
400
600
800
1000
1200
1400
1600
m/z

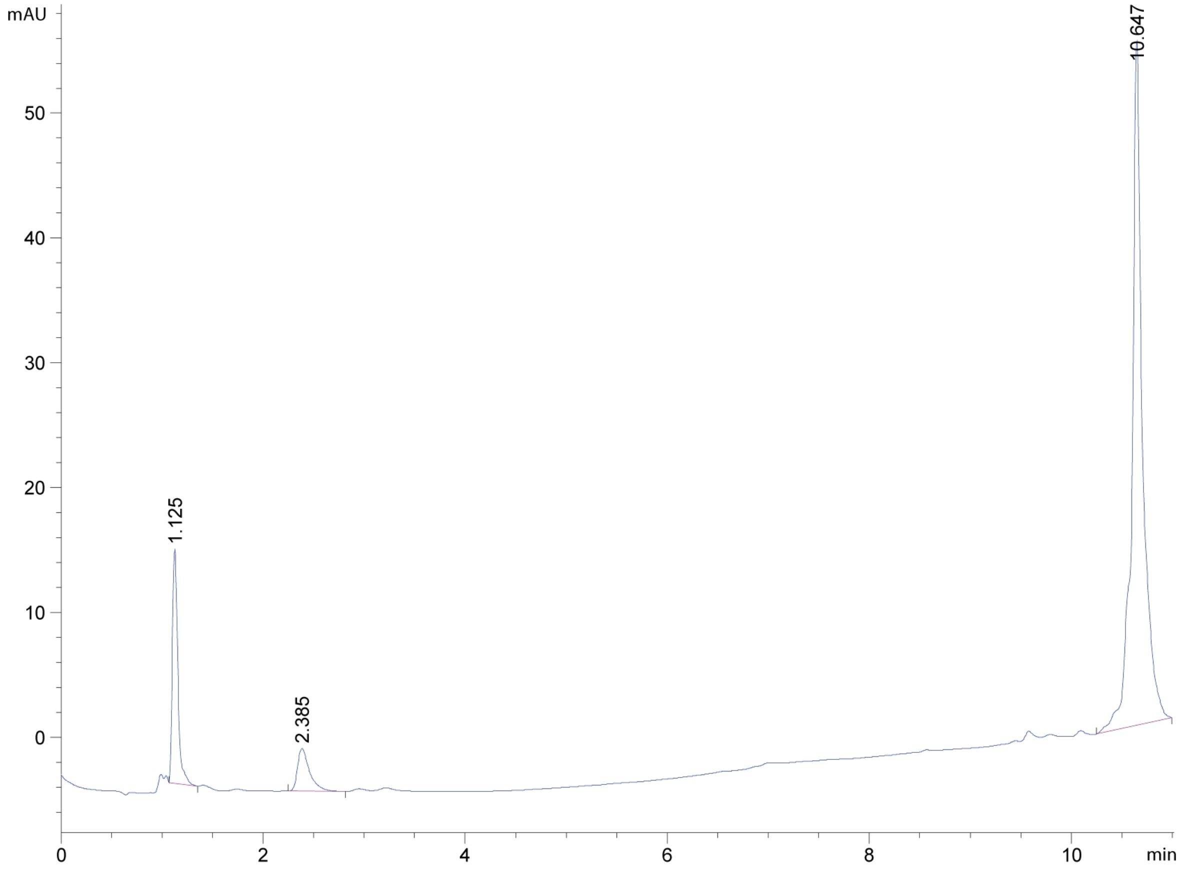
mAU
10.647
1.125
2.385
0
10
20
30
40
50
0
2
4
6
8
10
min

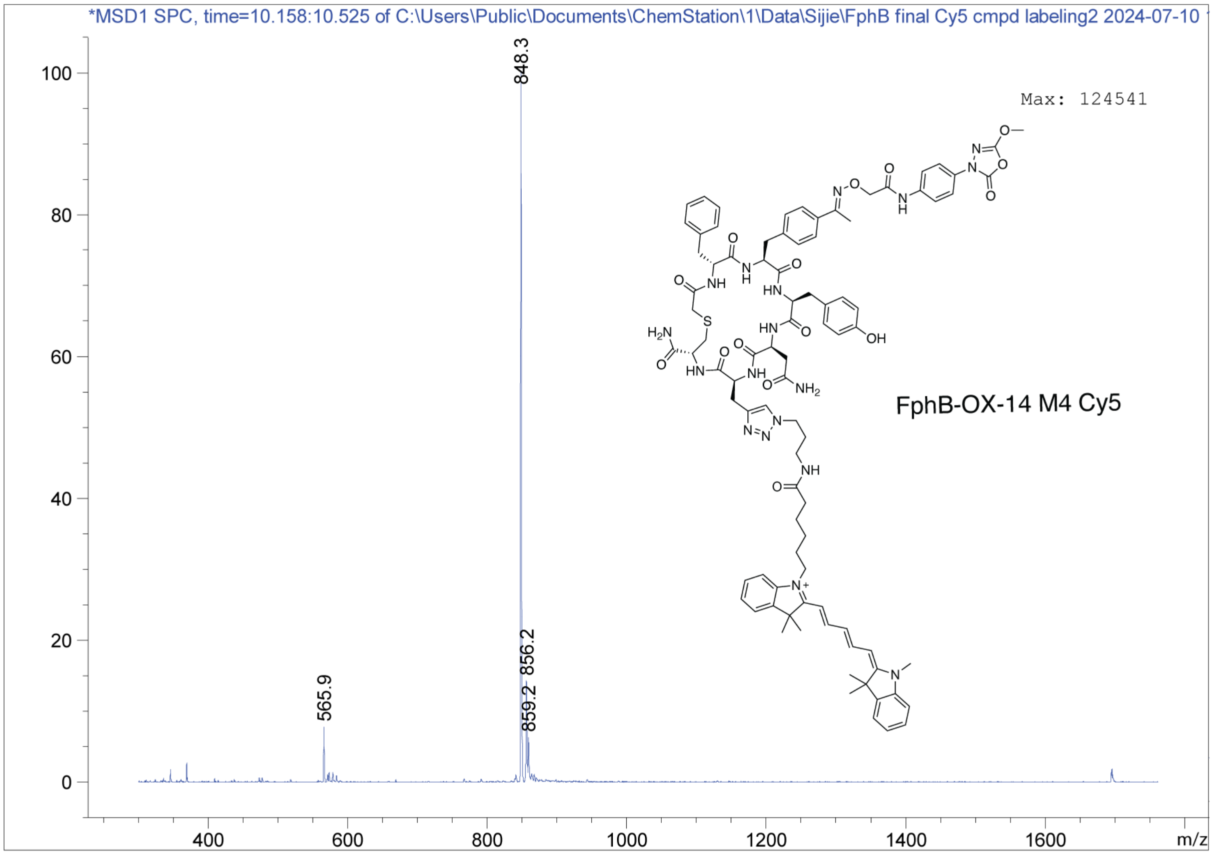
*MSD1 SPC, time=10.158:10.525 of C:\Users\Public\Documents\ChemStation\1\Data\Sijie\FphB final Cy5 cmpd labeling2 2024-07-10
Max: 124541
848.3
856.2
859.2
565.9
FphB-OX-14 M4 Cy5
m/z

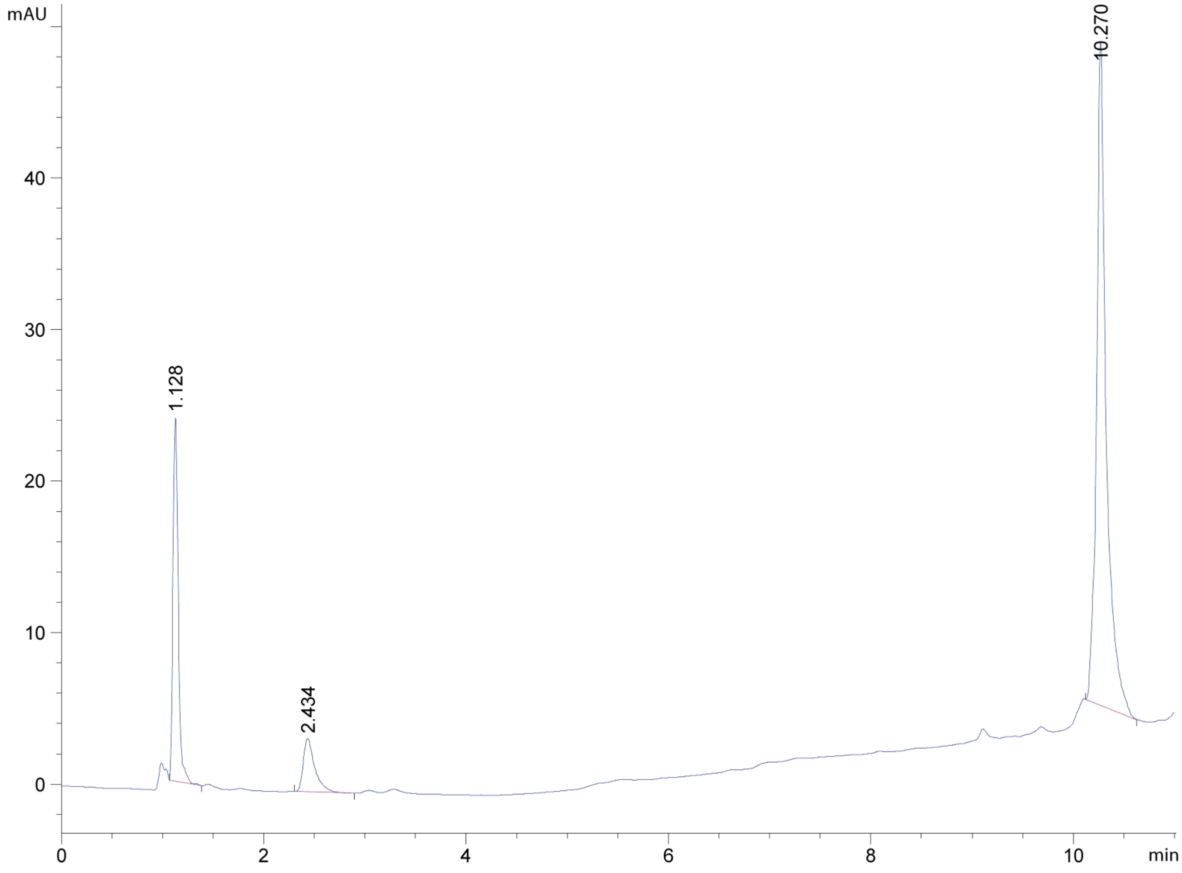
mAU
10.270
1.128
2.434
min

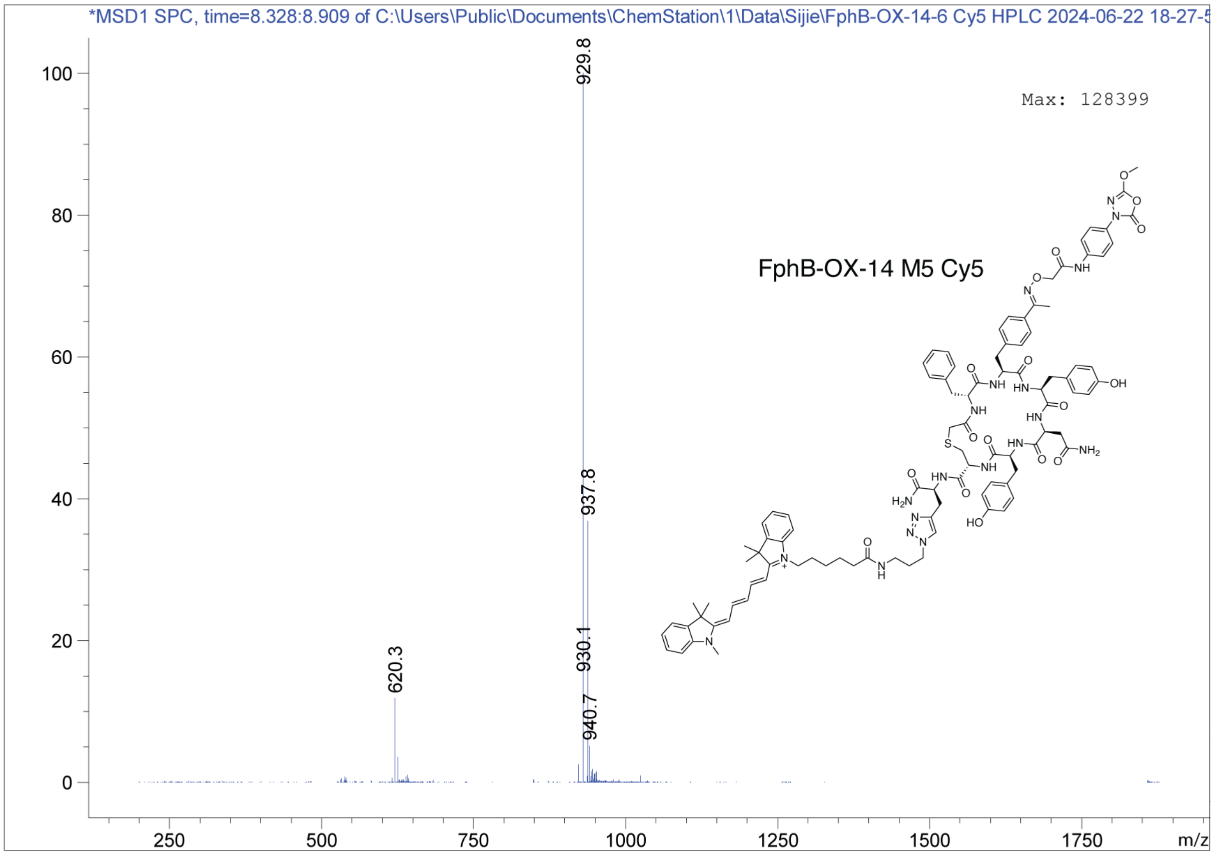
*MSD1 SPC, time=8.328:8.909 of C:\Users\Public\Documents\ChemStation\1\Data\Sijie\FphB-OX-14-6 Cy5 HPLC 2024-06-22 18-27-5
Max: 128399
FphB-OX-14 M5 Cy5
929.8
937.8
930.1
940.7
620.3
m/z

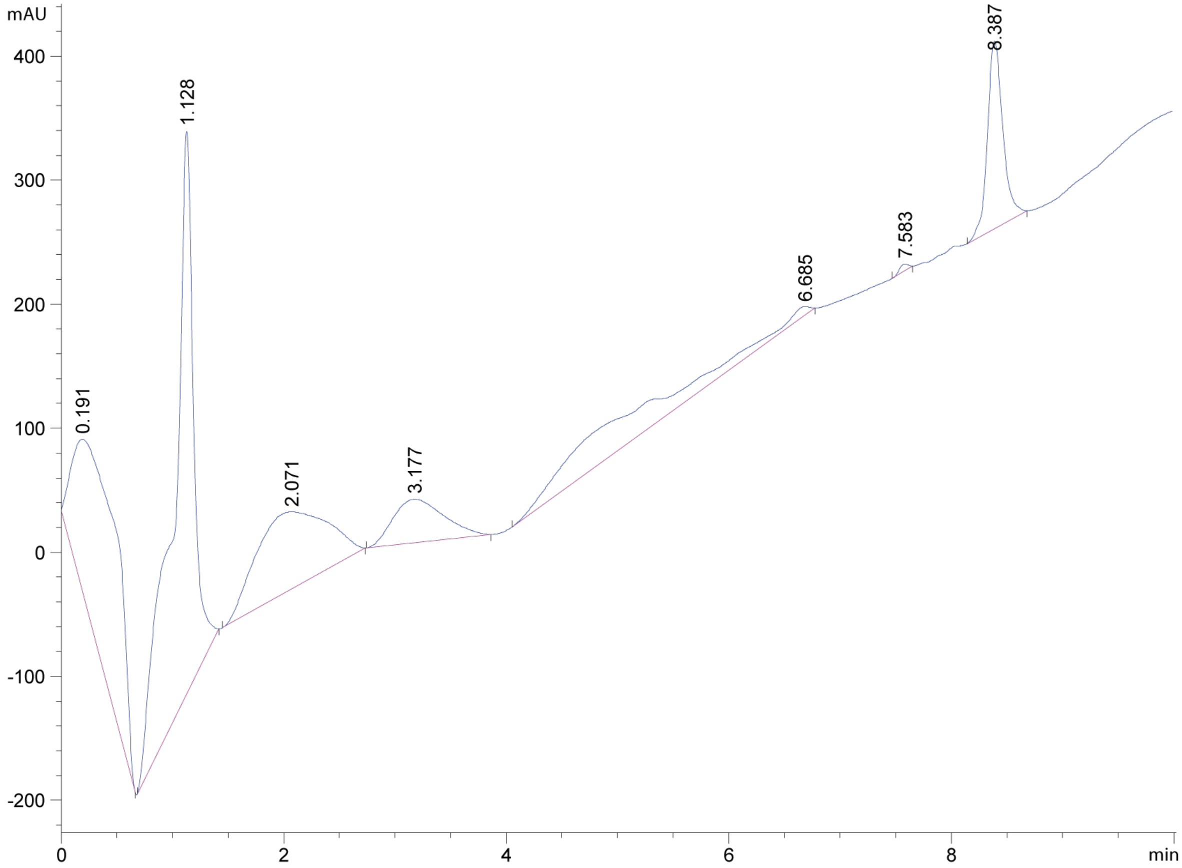
mAU
0.191
1.128
2.071
3.177
6.685
7.583
8.387
min

### 5. $^{1}$H NMR Spectra

$^{1}$H NMR of compound JJ-OX-009 (500 MHz, DMSO-$d_6$)

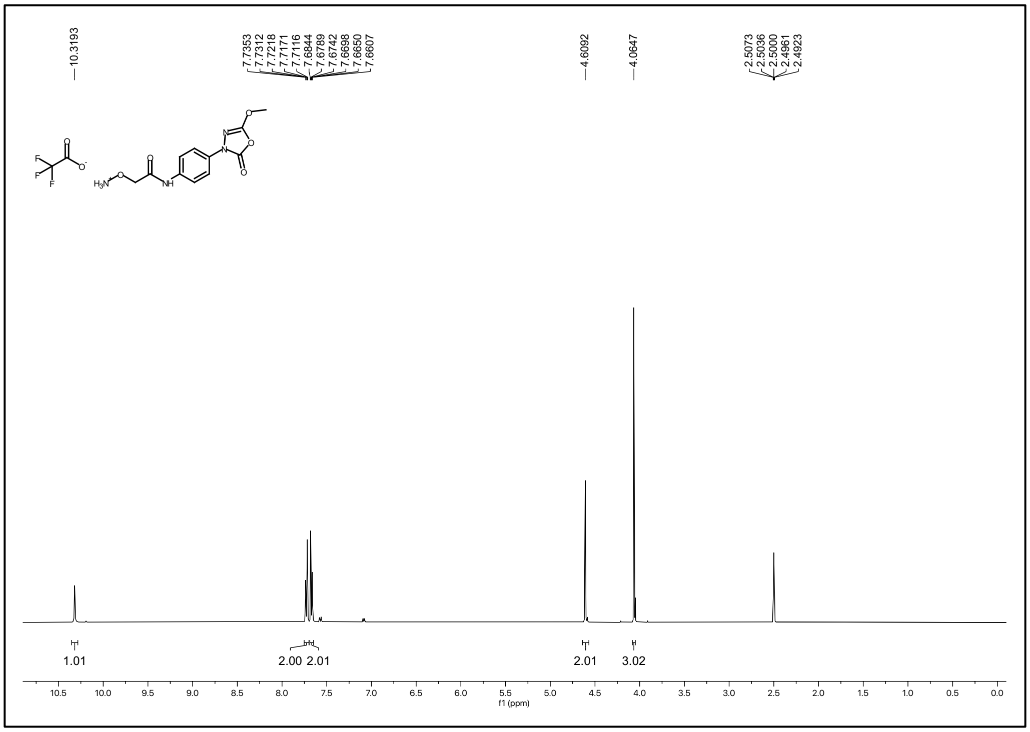

$^{1}$H NMR of compound OxF-CME (500 MHz, DMSO-$d_6$)

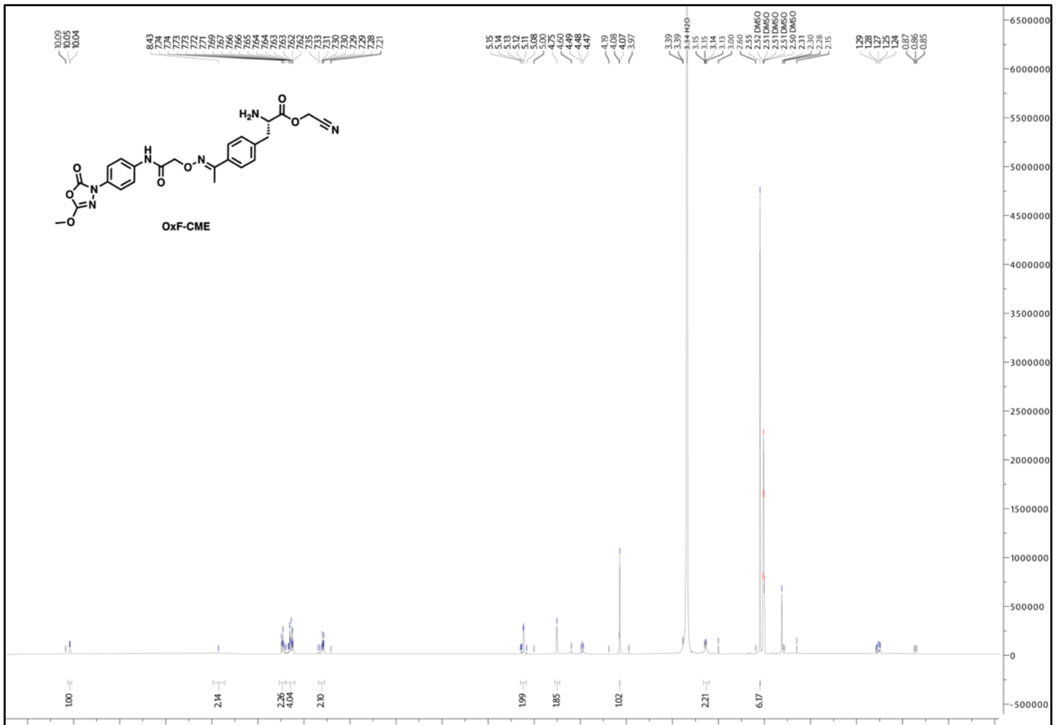

#### 6. $^{13}$C NMR Spectra

$^{13}$C NMR of compound JJ-OX-009 (500 MHz, DMSO-$d_6$)

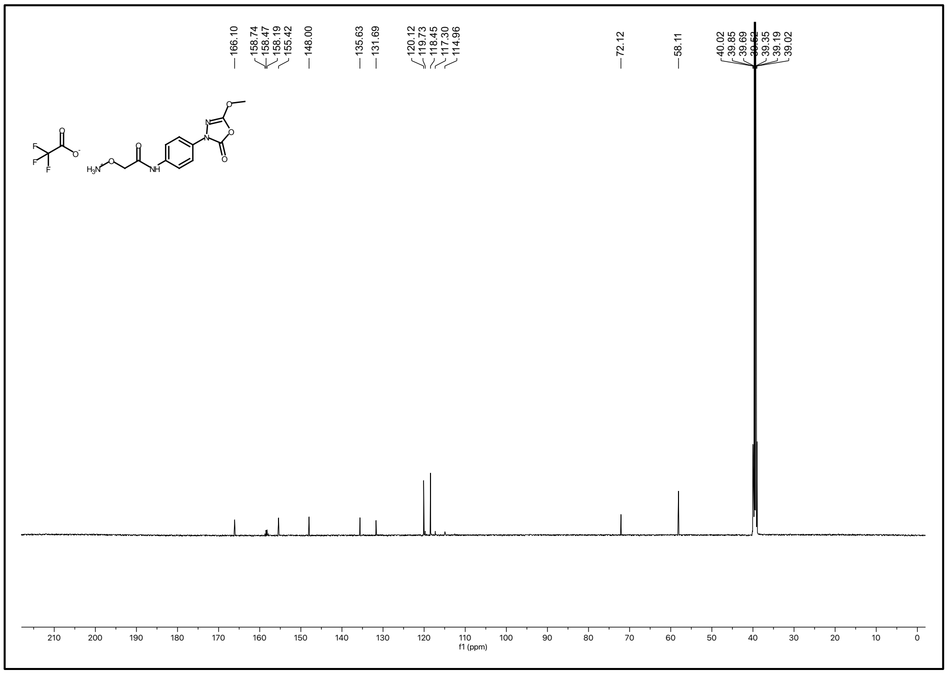

$^{13}$C NMR of compound OxF-CME (500 MHz, DMSO-$d_6$)

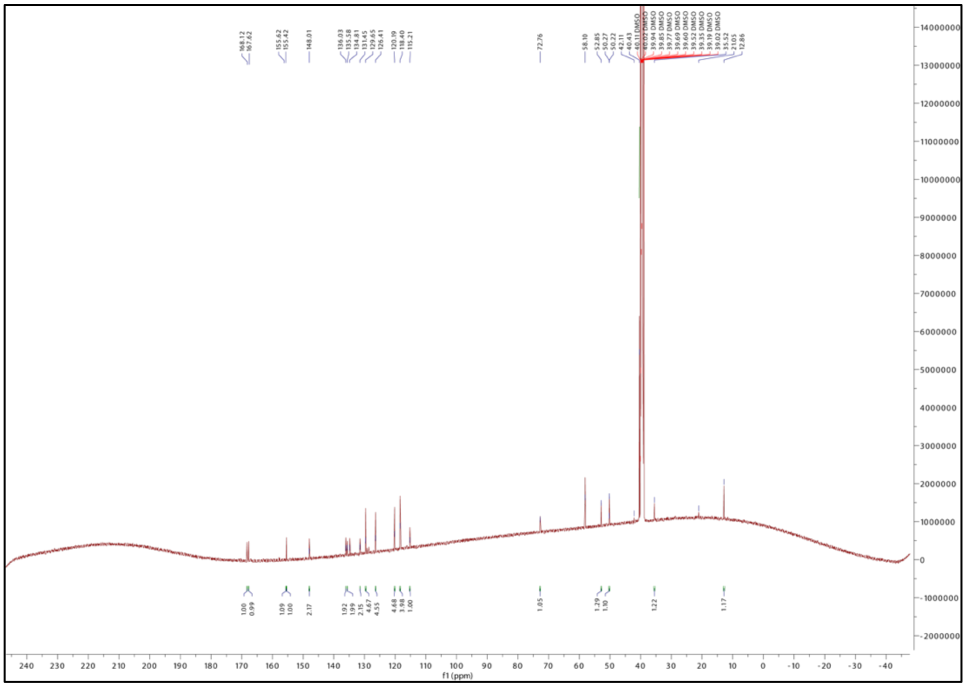

#### 7. REFERENCES

(1) Chan, A. I.; Sawant, M. S.; Burdick, D. J.; Tom, J.; Song, A.; Cunningham, C. N. Evaluating Translational Efficiency of Noncanonical Amino Acids to Inform the Design of Druglike Peptide Libraries. *ACS Chem. Biol.* **2023**, acschembio.2c00712. https://doi.org/10.1021/acschembio.2c00712.

(2) Adaligil, E.; Song, A.; Hallenbeck, K. K.; Cunningham, C. N.; Fairbrother, W. J. Ribosomal Synthesis of Macrocyclic Peptides with B2- and B2,3-Homo-Amino Acids for the Development of Natural Product-Like Combinatorial Libraries. *ACS Chem. Biol.* **2021**, *16* (6), 1011–1018. https://doi.org/10.1021/acschembio.1c00062.

(3) Goto, Y.; Katoh, T.; Suga, H. Flexizymes for Genetic Code Reprogramming. *Nat Protoc* **2011**, *6* (6), 779–790. https://doi.org/10.1038/nprot.2011.331.

(4) Ishizawa, T.; Kawakami, T.; Reid, P. C.; Murakami, H. TRAP Display: A High-Speed Selection Method for the Generation of Functional Polypeptides. *J. Am. Chem. Soc.* **2013**, *135* (14), 5433–5440. https://doi.org/10.1021/ja312579u.

(5) Jo, J.; Upadhyay, T.; Woods, E. C.; Park, K. W.; Pedowitz, N. J.; Jaworek-Korjakowska, J.; Wang, S.; Valdez, T. A.; Fellner, M.; Bogyo, M. Development of Oxadiazolone Activity-Based Probes Targeting FphE for Specific Detection of Staphylococcus Aureus Infections. *J. Am. Chem. Soc.* **2024**, *146* (10), 6880–6892. https://doi.org/10.1021/jacs.3c13974.

(6) Mobley, D. L.; Bannan, C. C.; Rizzi, A.; Bayly, C. I.; Chodera, J. D.; Lim, V. T.; Lim, N. M.; Beauchamp, K. A.; Slochower, D. R.; Shirts, M. R.; Gilson, M. K.; Eastman, P. K. Escaping Atom Types in Force Fields Using Direct Chemical Perception. *J. Chem. Theory Comput.* **2018**, *14* (11), 6076–6092. https://doi.org/10.1021/acs.jctc.8b00640.

(7) Wang, Y.; Fass, J.; Kaminow, B.; E. Herr, J.; Rufa, D.; Zhang, I.; Pulido, I.; Henry, M.; Macdonald, H. E. B.; Takaba, K.; D. Chodera, J. End-to-End Differentiable Construction of Molecular Mechanics Force Fields. *Chemical Science* **2022**, *13* (41), 12016–12033. https://doi.org/10.1039/D2SC02739A.

(8) Eastman, P.; Swails, J.; Chodera, J. D.; McGibbon, R. T.; Zhao, Y.; Beauchamp, K. A.; Wang, L.-P.; Simmonett, A. C.; Harrigan, M. P.; Stern, C. D.; Wiewiora, R. P.; Brooks, B. R.; Pande, V. S. OpenMM 7: Rapid Development of High Performance Algorithms for Molecular Dynamics. *PLOS Computational Biology* **2017**, *13* (7), e1005659. https://doi.org/10.1371/journal.pcbi.1005659.

(9) Essmann, U.; Perera, L.; Berkowitz, M. L.; Darden, T.; Lee, H.; Pedersen, L. G. A Smooth Particle Mesh Ewald Method. *The Journal of Chemical Physics* **1995**, *103* (19), 8577–8593. https://doi.org/10.1063/1.470117.

(10) Michaud-Agrawal, N.; Denning, E. J.; Woolf, T. B.; Beckstein, O. MDAnalysis: A Toolkit for the Analysis of Molecular Dynamics Simulations. *Journal of Computational Chemistry* **2011**, *32* (10), 2319–2327. https://doi.org/10.1002/jcc.21787.

(11) Bianco, G.; Forli, S.; Goodsell, D. S.; Olson, A. J. Covalent Docking Using Autodock: Two-Point Attractor and Flexible Side Chain Methods. *Protein Science* **2016**, *25* (1), 295–301. https://doi.org/10.1002/pro.2733.

(12) Santos-Martins, D.; Solis-Vasquez, L.; Tillack, A. F.; Sanner, M. F.; Koch, A.; Forli, S. Accelerating AutoDock4 with GPUs and Gradient-Based Local Search. *J. Chem. Theory Comput.* **2021**, *17* (2), 1060–1073. https://doi.org/10.1021/acs.jctc.0c01006.
